# S-Trap eliminates cell culture media polymeric surfactants for effective proteomic analysis of mammalian cell bioreactor supernatants

**DOI:** 10.1101/2020.02.17.951798

**Authors:** Lucia F. Zacchi, Dinora Roche Recinos, Ellen Otte, Campbell Aitken, Tony Hunt, Vanessa Sandford, Yih Yean Lee, Benjamin L. Schulz, Christopher B. Howard

## Abstract

Proteomic analysis of bioreactor supernatants can inform on cellular metabolic status, viability, and productivity, as well as product quality, which can in turn help optimize bioreactor operation. Incubating mammalian cells in bioreactors requires the addition of polymeric surfactants such as Pluronic F68, which reduce the sheer stress caused by agitation. However, these surfactants are incompatible with mass spectrometry proteomics and must be eliminated during sample preparation. Here, we compared four different sample preparation methods to eliminate polymeric surfactants from filtered bioreactor supernatant samples: organic solvent precipitation; filter-assisted sample preparation (FASP); S-Trap; and single-pot, solid-phase, sample preparation (SP3). We found that SP3 and S-Trap substantially reduced or eliminated the polymer(s), but S-Trap provided the most robust clean-up and highest quality data. Additionally, we observed that SP3 sample preparation of our samples and in other published datasets was associated with partial alkylation of cysteines, which could impact the confidence and robustness of protein identification and quantification. Finally, we observed that several commercial mammalian cell culture media and media supplements also contained polymers with similar mass spectrometry profiles, and we suggest that proteomic analyses in these media will also benefit from the use of S-Trap sample preparation.

## Introduction

The biopharmaceutical market continues to grow, with current and emerging highly post-translationally modified proteins driving research and development to optimize production efficiency while controlling quality ^1-3^. Bioreactor operation is critical for optimizing production ^4-7^, with changes in bioreactor operational conditions leading to changes in product yield and quality ^7-10^. In addition to the product, characterizing the co-secreted proteome is especially important, since the complexity and content of the secreted proteome can advise on the metabolic status and needs of the cells, as well as impacting the efficiency of downstream product purification and characterization ^11-13^.

Liquid chromatography coupled to tandem mass spectrometry (LC-MS/MS) is a versatile and effective tool to perform qualitative and quantitative measurements of proteins in a sample ^14-15^. Recent years have seen remarkable progress both in the hardware and software underpinning proteomics ^15^. Mass spectrometry proteomics can now provide a detailed overview and quantification of the proteins and their post-translational modifications in a wide variety of samples ^16-22^. However, even the highest-performing mass spectrometry instruments require effective and robust sample preparation to enrich analytes of interest and deplete contaminants. Many methods have been developed to aid in the preparation of samples for mass spectrometry proteomics, with the optimal selection depending on the specific sample content and experimental questions at hand ^23-40^.

Culturing mammalian cells in bioreactors requires the use of surfactants such as Pluronic F68 to protect the cells from hydrodynamic damage caused by agitation ^41-42^. However, these compounds can interfere with LC-MS/MS analysis and are best eliminated during sample preparation ^43^. Here we compared four different methods for mass spectrometry proteomics sample preparation: organic solvent precipitation ^44-46^, filter-assisted sample preparation (FASP) ^33-34^, S-Trap ^29, 32, 36-37, 47-48^, and single-pot, solid-phase, sample preparation (SP3) ^27-28^. We compared the ability of these methods to eliminate Pluronic F68 and similar polymers, and the number of proteins and peptides identified with each method. We found that FASP using Amicon columns was not an adequate method to prepare samples that contained these type of polymers (including bioreactor samples and commercial laboratory CHO medium). We also found that organic solvent precipitation, S-Trap, and SP3 were able to reduce the amount of polymer in the samples, but that SP3 and S-Trap performed better than precipitation. We also found that samples prepared with SP3 showed partial alkylation, a suboptimal feature for proteomic experiments. Together, S-Trap provided the most consistent polymer removal for robust, reproducible, and comprehensive proteomic analysis.

## Experimental section

### Sample collection

A variety of samples from fresh or spent mammalian cell culture media, fresh mammalian cell media supplements, and *Saccharomyces cerevisiae* whole cell extract were tested. The fresh media and media supplements tested included CD CHO medium (10743029, ThermoFisher), CHO CD EfficientFeed A (A1023401, ThermoFisher), CHO CD EfficientFeed B (A1024001, ThermoFisher), and typical mammalian cell culture media DMEM high glucose (11965092, ThermoFisher), EMEM (51412C, Merck), and Optimem (31985088, ThermoFisher) supplemented or not with 1 g/L Pluronic F68 (P1300, Sigma), anti-clumping agent (ACA) (0010057DG, ThermoFisher), and Glutamax (35050061, ThermoFisher).

Spent media from CHO cells incubated in flasks were obtained as follows. Chinese Hamster Ovary (CHO)-S cells were grown in suspension in CD CHO medium supplemented with 200 mM Glutamax. Cells were seeded at a density of 0.2×10^5^ cells/mL and cultured at 37 °C, 7.5% CO_2_, and 130 rpm shaking. Cells were cultured for 3-5 days until reaching a density of 1-2×10^6^ cells/mL. Cells were pelleted at 700 rpm for 10 min and the supernatant was removed for analysis as spent medium.

The spent media was obtained from CHO cell fed batch culture. A CHO K1SV cell line stably expressing a modified version of human Coagulation factor IX (rFIX, accession number P00740 UniProtKB, with Q_2_G and P_44_V amino acid substitutions) integrated using the glutamine synthetase expression system ^49^, and also expressing the protease PACE/Furin (Accession number P09958, UniProtKB) (provided by CSL, Marburg, Germany) was seeded at 0.3×10^6^ cells/mL in 3 L of CD CHO medium supplemented with 30 mg/L reduced menadione sodium bisulphite. Bioreactors were fed EfficientFeedB as a daily bolus starting on working day (WD) 3 until WD10, up to the equivalent of 40% (1.2 L) to a total working volume of 4.2 L in the bioreactor, following manufacturer’s instructions. Samples of 15 mL were collected, centrifuged at 2,000 rcf for 10 min, filtered through a 0.2 µm polyethersulfone filter (Pall), dispensed in matrix tubes to a volume of 0.5 mL and stored at −80 °C.

*Saccharomyces cerevisiae* were obtained as follows. *S. cerevisiae* strain YBS10 ^50^ was grown at 30 °C overnight with shaking at 200 rpm in YPD (1% yeast extract, 2% peptone, 2% glucose). 3 mL of saturated culture was centrifuged at 18,000 rcf for 3 min at room temperature. The medium was discarded and the cells were resuspended in 200 µL of ice-cold 50 mM HEPES buffer pH 7.4 containing 1 x protease inhibitor cocktail (Roche) and 1 mM phenylmethylsulfonyl fluoride. Cells were lysed by glass bead beating for 20 min at 4 °C. The lysate was transferred to a new microfuge tube and was clarified by centrifugation at 18,000 rcf for 1 min at room temperature. The supernatant was transferred to a protein LoBind tube (Eppendorf) and frozen at −20 °C.

### Mass spectrometry sample preparation

Four different sample preparation techniques were used to prepare the samples: organic solvent precipitation, Filter Aided Sample Preparation (FASP) using 0.5 mL 10 kDa or 30 kDa cut-off Amicon columns (UFC503024 and UFC503096, Millipore), S-Trap columns (S-Trap C02-mini, Protifi), and Single-Pot Solid-Phase enhanced Sample Preparation (SP3) using paramagnetic beads (GEHE45152105050250 Sera-Mag SpeedBead Carboxylate-Modified Magnetic Particles (Hydrophilic, GE Healthcare) and GEHE65152105050250 Sera-Mag SpeedBead Carboxylate-Modified Magnetic Particles (Hydrophobic, GE Healthcare)).

The organic solvent precipitation protocol was performed as previously described ^51^, except as noted. Briefly, 300 µL of sample was clarified by centrifugation at 18,000 rcf for 3 min at room temperature, and 25 µL was transferred to a protein LoBind tube containing 200 µL of denaturation buffer (6 M guanidine hydrochloride, 50 mM Tris HCl buffer pH 8, 10 mM dithiothreitol (DTT)). Samples were incubated at 30 °C for 30 min in a MS100 Thermoshaker at 1500 rpm, acrylamide was added to a final concentration of 25 mM, and samples were incubated at 30 °C for 1 h in a Thermoshaker at 1500 rpm. DTT was added to an additional final concentration of 5 mM to quench excess acrylamide, and samples were precipitated by addition of 4 volumes of 1:1 methanol:acetone and incubation at −20 °C for 16 h. Samples were centrifuged at 21,000 rcf for 10 min at room temperature, the supernatant was removed, samples were centrifuged again for at 21,000 rcf for 1 min, and the remaining solvent was removed. Samples were air dried for ∼15 min at room temperature, resuspended in 50 mM ammonium bicarbonate containing 0.5 µg of trypsin (T6567, Sigma) and incubated at 37 °C for 16 h in a Thermoshaker at 1500 rpm. All samples were desalted by ziptipping with C18 ZipTips (ZTC18S960, Millipore). For comparison of the extent of alkylation using SP3 or precipitation, the denaturation, reduction, and alkylation steps were performed identically to the SP3 protocol as described below, with a heat denaturation incubation step of 95 °C for 10 min.

The FASP protocol was performed as previously described ^33-35^. Briefly, samples were centrifuged, denatured, reduced, and alkylated as described above for the precipitation. Briefly, 300 µL of sample was clarified by centrifugation at 18,000 rcf for 3 min at room temperature, and 25 µL was transferred to a protein LoBind tube containing 200 µL of denaturation buffer (6 M guanidine hydrochloride, 50 mM Tris HCl buffer pH 8, 10 mM dithiothreitol (DTT)). Samples were incubated at 30 °C for 30 min in a MS100 Thermoshaker at 1500 rpm, acrylamide was added to a final concentration of 25 mM, and samples were incubated at 30 °C for 1 h in a Thermoshaker at 1500 rpm. DTT was added to an additional final concentration of 5 mM to quench excess acrylamide. Samples were then loaded onto a 10 or a 30 kDa cut-off Amicon column and centrifuged at 10,000 rcf for at least 10 min. The filter was washed twice with 500 µL of 50 mM ammonium bicarbonate, and 150 µL of ammonium bicarbonate solution containing 0.5 µg of trypsin was added to the column. The column was then incubated overnight at 37 °C in a rotatory shaker. The column was transferred to a new collection tube, and the liquid was eluted by centrifugation at 10,000 rcf for at least 10 min at room temperature. To recover the digested peptides that were still trapped in the filter, 50 µL of 50 mM ammonium bicarbonate was added to the column, followed by centrifugation at 10,000 rcf for at least 10 min at room temperature. Samples were desalted with C18 ZipTips.

The S-Trap protocol was performed following the manufacturer’s instructions (www.protifi.com). Briefly, 25 µL of centrifuged supernatant samples were transferred to protein LoBind tubes containing 25 µL of 2 times lysis buffer (10 % sodium dodecyl sulfate (SDS), 100 mM Tris HCl buffer pH 7.55, 20 mM DTT). Samples were incubated at 90 °C for 10 min and cooled to room temperature before adding acrylamide to a final concentration of 25 mM and incubation at 30 °C for 1 h in a Thermoshaker at 1500 rpm. Excess acrylamide was quenched by adding DTT to an additional final concentration of 5 mM. Samples were acidified by adding phosphoric acid to 1.2% v/v final concentration, and were diluted 1:7 with S-Trap binding buffer (90% methanol, 100 mM Tris HCl buffer pH 7.1), loaded onto the S-Trap mini columns, and centrifuged at 4,000 rcf at room temperature. The samples were washed 5 times with 400 µL of S-Trap binding buffer. Samples were then resuspended in 125 µL of 50 mM ammonium bicarbonate with 1 µg of trypsin, and columns were incubated at 37 °C for 15 h in a humidified chamber, without agitation. To recover the peptides, the column was first rehydrated with 80 µL of 50 mM ammonium bicarbonate, and after 15 min of incubation at room temperature the columns were centrifuged at 1,000 rcf for 1 min at room temperature. This was followed by subsequent elution with 80 µL of 0.1% formic acid followed by 80 µL of 50% acetonitrile and 0.1% formic acid. All elutions were pooled, samples were dried in a MiVac Sample concentrator (SP Scientific), resuspended in 30 µL of 0.1% formic acid, and desalted with C18 ZipTips.

The SP3 protocol was performed as previously described ^24^. Briefly, 25 µL of centrifuged supernatant samples were transferred to a protein LoBind tube (Eppendorf) and 200 µL of buffer SDS (1% SDS, 50 mM HEPES pH 7.5, 10 mM DTT, and 1 x cOmplete mini EDTA-free Protease Inhibitor Cocktail (4603159001, Roche)) was added. Samples were incubated at 60 °C for 30 min or 95 °C for 10 min and then cooled to room temperature before acrylamide was added to a final concentration of 25 mM. After incubation at 30 °C for 1 h in a multivortex, DTT was added to a final concentration of 5 mM, samples were incubated at room temperature for 5 min. SP3 beads were prepared by mixing equal volumes of hydrophobic and hydrophilic beads into one Eppendorf tube, the tube was placed in a magnetic rack (Invitrogen Dynal), beads were pelleted, the liquid was removed by pipetting, and the beads were washed 4 times with 500 µL of water. Beads were reconstituted in water to a concentration of 20 µg/µL. Beads were kept at 4 °C until use. Denatured and reduced/alkylated protein samples were transferred to regular Eppendorf tubes. 5 µL of bead mix was added and mixed by pipetting, and 100% ethanol was added to a final concentration of 50% v/v. The mix was incubated at room temperature in a thermomixer at 1000 rpm for 25 min. Tubes were centrifuged briefly to collect the liquid and beads, and the tubes were placed in a magnetic rack and incubated for 2 min until the beads settled. The supernatant was removed and discarded and beads were washed 3 times with 200 µL of 80% ethanol. 50 µL of 50 mM ammonium bicarbonate with 0.5 µg of trypsin was added to each tube. Tubes were sonicated at low intensity at 17 °C for 15 s in a water bath to reconstitute the beads, and were incubated overnight at 37 °C in a Thermoshaker at 1000 rpm. The tubes were centrifuged at 20,000 rcf for 1 min to pellet the beads, and placed in a magnetic rack for 2 min. When the beads settled the supernatant was recovered into a protein LoBind tube and peptides were desalted with C18 ZipTips.

Details of the methods, samples, and replicates are provided in Supplementary information.

### Mass spectrometry data acquisition and analysis

Desalted peptides were analyzed by liquid chromatography electrospray ionization tandem mass spectrometry (LC-ESI-MS/MS) using a Prominence nanoLC system (Shimadzu) and a TripleTof 5600 mass spectrometer with a Nanospray III interface (SCIEX) essentially as described ^52-53^. Samples were desalted on an Agilent C18 trap (0.3 × 5 mm, 5 µm) at a flow rate of 30 µL/min for 3 min, followed by separation on a Vydac Everest C18 (300 Å, 5 µm, 150 mm × 150 µm) column at a flow rate of 1 µL/min. A gradient of 10-60% buffer B over 45 min where buffer A = 1% ACN / 0.1% FA and buffer B = 80% ACN / 0.1% FA was used to separate peptides. Gas and voltage settings were adjusted as required. An MS TOF scan across 350-1800 *m/z* was performed for 0.5 s followed by data dependent acquisition (DDA) of up to 20 peptides with intensity greater than 100 counts, across 100-1800 *m/z* (0.05 s per spectra) using a collision energy (CE) of 40 +/- 15 V. The mass spectrometry data is available in the ProteomeXchange Consortium (http://proteomecentral.proteomexchange.org) ^54^ via the PRIDE partner repository ^55^ with the dataset identifier PXD017214.

DDA data was analyzed using ProteinPilot v5.0.1 (SCIEX) and Preview (v2.13.17, Protein Metrics). The following parameters were used to search in ProteinPilot: Sample type: identification; cysteine alkylation: acrylamide; digestion: trypsin; instrument: TripleTOF 5600; ID focus: biological modifications; search effort: thorough; FDR analysis: 1% global. The search database included the entire *Cricetulus griseus* proteome (UP000001075 downloaded from UniProtKB on May 14^th^ 2017 containing a total of 23,884 proteins) and human Coagulation Factor IX wild-type amino acid sequence (P00740, UniProtKB). The following parameters were used to search in Preview: Modifications: Cysteine fixed +71.037114 (propionamide), +57.021464 (carbamidomethylation), or unknown; Cleavage site: RK, C-terminal; Initial search specificity: fully specific (fastest); Fragmentation type: CID/HCD; Protein database: the *Cricetulus griseus* proteome UP000001075 downloaded from UniProtKB on April 20^th^ 2018 containing a total of 23884 proteins, the *Saccharomyces cerevisiae* S288C proteome (UP000002311 downloaded on April 20^th^ 2018 containing a total of 6,049 proteins), or the *Homo sapiens* proteome (UP000005640 downloaded on April 20^th^ 2018 containing a total of 20,303 proteins). The number of proteins and distinct peptides shown correspond to ProteinPilot searches, considering 1% global FDR and a minimum of one confident peptide per protein.

## Results and discussion

Our goal was to develop a simple, efficient, and robust method to prepare samples from bioreactor supernatant for mass spectrometry proteomic analysis that eliminated polymeric contaminants and achieved high performance protein identification and characterization. We first tested the FASP method using 0.5 mL 10 kDa cut-off Amicon columns. FASP is a commonly used sample preparation method for bottom-up proteomics that provides consistent and reproducible results for a variety of samples ^33-35^, including bioreactor supernatants ^56-60^. We processed 20 µL of filtered supernatant from a fed batch culture in CD CHO medium supplemented with CHO CD EfficientFeed B and vitamin K. We identified 526 proteins and 2441 distinct peptides in the sample (1% Global FDR) (Fig. 1A). However, we also observed the presence of an abundant polymer (curved dotted lines showing increasing molecular weight (MW) and retention time (RT), Fig. 1A). MS/MS of diverse MW forms of this polymer produced fragment ions with *m/z* of 45.04, 89.06, 133.08, and 177.10 (Fig. 1B). We therefore concluded that there was a polymer in the bioreactor samples that the regular FASP protocol was not able to eliminate.

**Figure 1.**
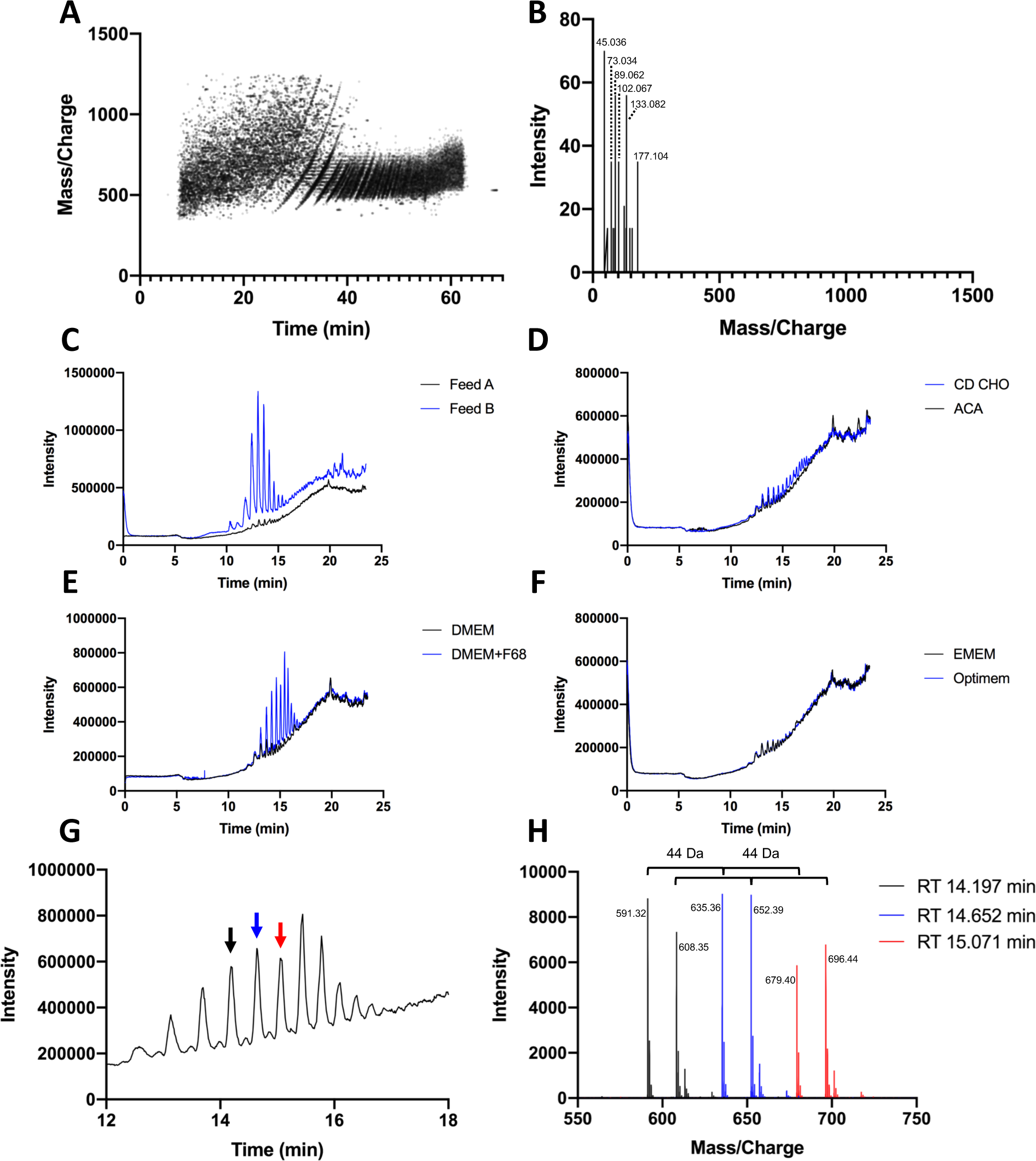
Presence of a polymeric contaminant in a variety of mammalian cell media and media supplements. (**A**) Representation of all fragmented precursors identified in DDA mode in a sample of filtered supernatant from CHO bioreactor culture grown in CD CHO medium supplemented with EfficientFeed B and vitamin K, and prepared with FASP using a 10 kDa cut-off Amicon column. (**B**) Representative MS/MS spectrum of the polymeric compound. Typical ions observed included *m/z* of 45.04, 89.06, 133.08, and 177.10. Total ion chromatograms (TIC) of fresh media and media supplements prepared using FASP with a 10 kDa cut-off Amicon column of (**C**) EfficientFeed A and B, (**D**) CD CHO and anti-clumping agent (ACA), (**E**) DMEM ± 1 g/L Pluronic F68, and (**F**) EMEM and Optimem. (**G**) TIC of DMEM + 1 g/L Pluronic F68 between retention time (RT) 12-18 min showing multiple peaks corresponding to the polymer. The peaks at RT 14.197 min, 14.652 min, and 15.071 min are highlighted with black, blue, and red arrows, respectively, and their corresponding MS spectra showing 44 Da repeating units are shown in (**H**).

The polymer in the bioreactor sample was likely Pluronic F68, which is a common media additive. However, we were intrigued by our observation that a polymer with similar characteristics was also present in filtered spent CD CHO medium supplemented with anti-clumping agent (ACA) from transient CHO cell transfections (Supplementary Figure S1). We therefore tested multiple fresh mammalian cell culture media or media supplements for the presence of a similar polymer (Fig. 1). Indeed, a polymeric contaminant with a similar chromatography and fragmentation profile was observed in fresh CHO CD EfficientFeed B and CD CHO, and less abundantly in DMEM (Fig. 1C, D, and E). CHO CD EfficientFeed A, ACA, EMEM, and Optimem showed little or no polymeric contaminant (Fig. 1C, D, and F). The polymer observed in all fresh and spent media (transient lab transfection or bioreactor) consisted of repeating units of ∼44.04 Da that eluted throughout the LC gradient (Fig. 1H). Common contaminants with repeating units of ∼44 Da that interfere with MS detection are polyethylene glycol and related compounds such as the detergents Triton and Tween ^43^. Due to the proprietary nature of the chemical composition of commercial media, the source and identity of the polymer in fresh media and media supplements tested was unclear. When we supplemented fresh DMEM medium with 1 g/L Pluronic F68, we observed a considerable increase in the polymeric signal detected (Fig. 1E), with the same MS and MS/MS ion patterns as in the bioreactor sample, suggesting the polymer observed in the bioreactor samples was Pluronic F68 (Fig. 1B and H, and Supplementary Figure S2). Together, our results showed that several commercial mammalian cell media contained a polymer with 44 Da repeating units that was similar to Pluronic F68, and that could not be removed from samples using the regular FASP protocol.

We attempted to optimize the FASP method by changing the buffers used in the washing step. Amicon columns do not tolerate high concentration of detergents, so we tested 6 M guanidinium chloride in 50 mM Tris HCl buffer pH 8, or 8 M urea in 50 mM Tris HCl buffer pH 8, two common protein denaturants used in bottom up proteomics. Performing two washes with these solutions did not successfully eliminate the polymer (Supplementary Figure S3). Next, we tested increasing the number of washes with the standard 50 mM ammonium bicarbonate (from one to seven washes) using 10 kDa or 30 kDa cut-off Amicon columns. Increasing the number of washes reduced the amount of polymer contaminant remaining in the samples but did not eliminate the polymer completely, and it also reduced the number of proteins identified (Supplementary Figure S1). Microcon columns (Millipore) also did not successfully eliminate the polymer (Supplementary Figure S3). We concluded that FASP using Amicon or Microcon columns was not suitable to prepare bioreactor supernatant samples for proteomic analysis.

To change strategy, we compared the FASP protocol to three different bottom-up proteomics sample preparation methods: organic solvent protein precipitation ^44-46^, S-Trap columns ^29, 32, 36-37, 47-48^, and SP3 with paramagnetic beads (Fig. 2 and Supplementary Table S1) ^23-24, 26-31^. The organic solvent used here was a 1:1 mixture of methanol and acetone ^44-45^. All methods have been previously shown to eliminate detergents from diverse samples ^24-25, 46-48^, and organic solvent precipitation ^61-64^ and FASP ^56-60^ have been previously used to prepare bioreactor supernatant samples for LC-MS/MS proteomics. We found that all three methods could remove the polymer(s) from the samples and that SP3 and S-Trap were better than precipitation (Fig. 2B-E). SP3 was recently shown to eliminate polymeric contaminants such as Pluronic F68 from samples for mass spectrometry proteomics ^25^, and we corroborated this result, although in our hands we observed inconsistent polymer removal (Fig. 2C and Supplementary Fig. S4). On the other hand, S-Trap consistently achieved almost complete polymer removal from the samples over multiple trials (Fig. 2D). We also found that all three methods showed comparable total ion chromatograms (TICs), which had higher intensities than the FASP TICs (Fig. 2A, orange) even though the same amount of starting material was used (Fig. 2A). S-Trap also identified the same number or more proteins than the other three methods (Fig. 2F, G). Indeed, S-Trap has also been shown to provide better protein coverage and more robust bottom-up proteomic results than FASP for other sample types, such as mammalian whole cell lysates and milk fat globules ^32, 36^. The comparative protein and peptide identification results for all methods were consistent with prior publications, showing that S-Trap outperformed FASP and precipitation in terms of the number of proteins identified ^23, 29, 36^ (Fig. 2F). Further, S-Trap, SP3, and precipitation allowed identification of more peptides than FASP (Fig. 2G). Together, due to the more consistent polymer removal and improved proteomic results, we concluded that S-Trap was the method of choice to eliminate Pluronic F68 from bioreactor samples and to eliminate similar polymeric contaminants present in mammalian cell media and media supplements for mass spectrometry proteomic applications.

**Figure 2.**
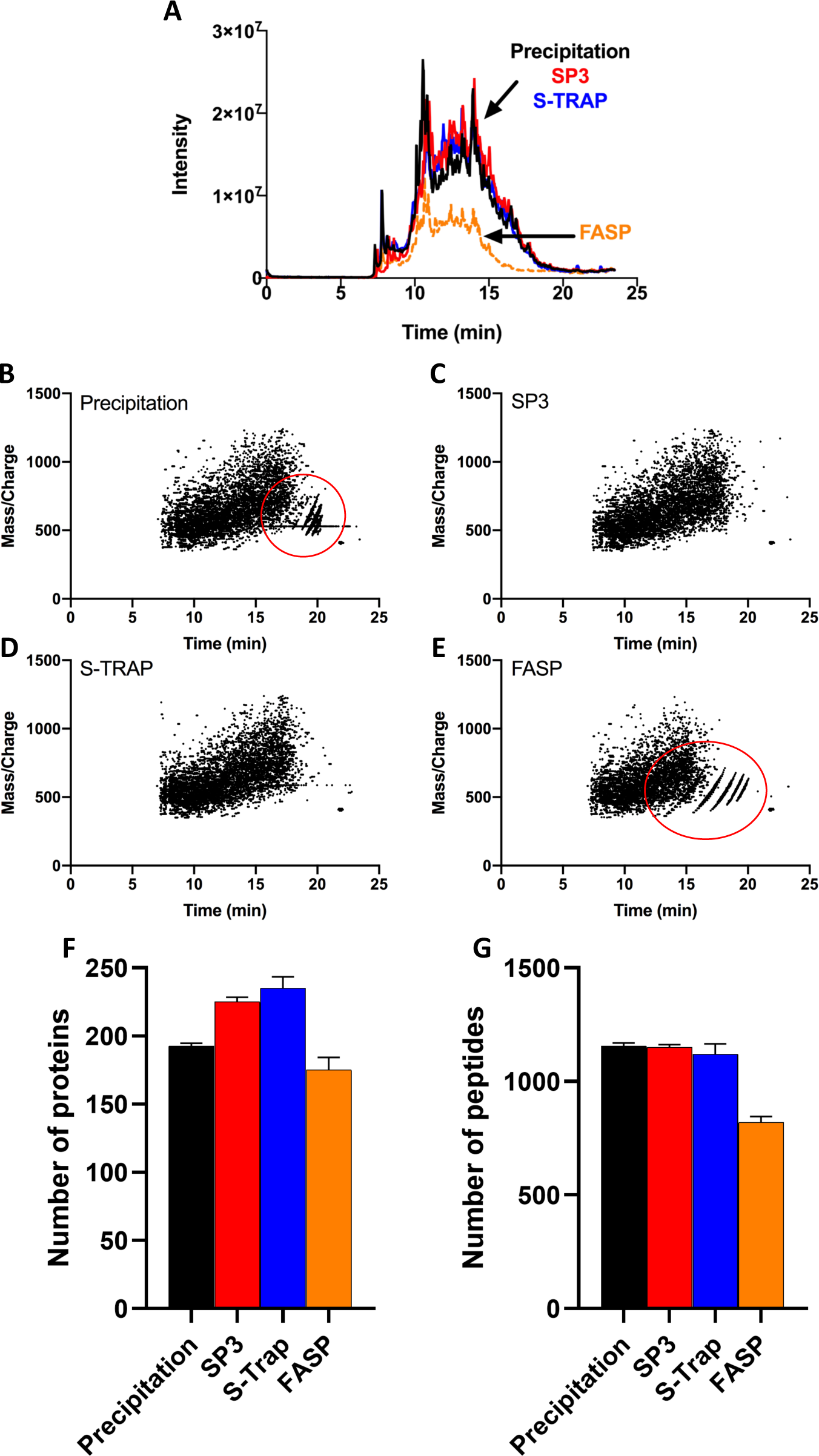
Comparison of sample preparation methods for bioreactor supernatant samples. 20 μL of filtered bioreactor supernatant sample from CHO cells incubated in CD CHO medium supplemented with EfficientFeed B were prepared using four different methods. (**A**) TIC obtained with organic solvent precipitation (black line), SP3 (red line), S-Trap (blue line), or FASP using 30 kDa Amicon columns (orange dotted line). Representations of all fragmented precursors identified in DDA mode for (**B**) precipitation, (**C**) SP3, (**D**) S-Trap, and (**E**) FASP. (**F**) Number of proteins (with at least 1 confident peptide match) and (**G**) peptides identified with each method in ProteinPilot, considering 1% global FDR (Mean +/- SD, n=2). Red circles in (**B**) and (**E**) denote the presence of the polymer.

SP3 is a recently developed sample preparation method for bottom-up proteomics ^23-31^, and we verified that it was easy to perform and that it provided superior results to FASP (Fig. 2). However, during the course of our analyses we uncovered a potential limitation of the SP3 technology: partial peptide alkylation (Fig. 3). A key step during sample preparation for bottom-up proteomics is reduction and alkylation of cysteine residues. Cysteine alkylation ensures that disulfide bonds do not re-form after reduction, facilitating peptide identification and measurement. Normally, cysteine alkylation is set as a fixed parameter during software identification searches under the assumption that alkylation has proceeded to completion and with the expectation that non-modified peptides will not be identified. However, most database searching software provides flexibility in these search parameters, allowing discovery of unexpected or missing modifications. Using ProteinPilot (SCIEX) and Preview (Protein Metrics) we observed that while the mammalian cell culture samples processed using S-Trap, precipitation, and FASP showed complete alkylation, samples processed with the SP3 method consistently showed partial alkylation (Fig. 3, Supplementary Figure S5, and Supplementary Tables S1 and S2). In order to rule out an effect of polymeric surfactants or sample type on the partial-alkylation observed with SP3 we also tested yeast whole cell extracts from yeast grown in polymer-free YPD medium (Fig. 3). To control for the potential impact of differences in the sample preparation methods prior to loading the sample into the beads or columns, the same buffers and procedure to reduce and alkylate the yeast whole cell extract sample were used for the precipitation and SP3 methods (Fig. 3 and Supplementary Table S4). Analyses using Preview showed that duplicate yeast whole cell extract SP3 samples showed 61% and 90% of peptides alkylated compared to 100% for the same samples processed with S-Trap or precipitation. Notably, the abundance of some non-alkylated peptide variants in the SP3 samples was higher than the abundance of the equivalent alkylated peptides (Fig. 3A and D). Figure 3 shows two examples of partially alkylated peptides from the two most abundant proteins identified in yeast whole cell extracts prepared using SP3: peptide Y_414_RPNCPIILVTR_425_ from Pyruvate Kinase 1 (P00549, Fig. 3 A-C) and peptide I_244_GLDCASSEFFK_255_ from Enolase 2 (P00925, Fig. 3 D-F). Samples prepared with SP3 showed abundant non-alkylated peptides (YRPNCPIILVTR: 482.2746, *z*=3 and RT ∼17 min; and IGLDCASSEFFK: 658.8132, *z*=2, RT ∼20 min, Fig. 3A and D and Supplementary Fig. S5) compared to their alkylated counterparts (YRPNC(+71)PIILVTR: 505.9537, *z*=3 and RT ∼15.1 min, and IGLDC(+71)ASSEFFK: 694.3318, *z*=2, RT ∼19 min, Fig. 3A-D and Supplementary Fig. S5). The same yeast whole cell extract samples prepared and analyzed in parallel with S-Trap (Fig. 3B and E) or precipitation (Fig. 3C and F, Supplementary Table ST2) showed complete alkylation. The partial alkylation observed with SP3 sample preparation was not due to the type of sample, since we observed a similar partial alkylation in bioreactor supernatants, spent CD CHO medium from transfected CHO cells, and yeast whole cell extract. The partial alkylation was also not due to the presence of a polymer in the samples (yeast whole cell extract samples were prepared in the absence of any obvious polymeric compound), and was observed when the denaturation incubation condition was 60 °C for 30 min or 95 °C for 10 min (Fig. 3, Supplementary Tables S1 and S2). Finally, the partial alkylation was not due to differences in the denaturation, reduction, and alkylation steps between the protocols, because for analysis of yeast samples the precipitation and SP3 sample preparation steps were performed identically and simultaneously. To exclude the possibility that the partial alkylation we observed was a consequence of our sample preparation technique, we inspected published data that had been obtained using SP3 (PXD008698 ^24^, Supplementary Table S3). In agreement with our data, Preview analyses of this published data also showed partial alkylation in SP3 samples. For example, Preview found alkylation of only 53%, 76%, and 86% of 72, 63, and 59 peptides analyzed in three raw data files from this dataset. The partial alkylation observed with SP3 is intriguing, since the denaturation, reduction, and alkylation steps happen before the proteins are mixed with the beads (see the Experimental Section). Partial alkylation is problematic for the quality of protein identification, and especially for the robustness of peptide quantification. In addition, partial alkylation could be a symptom of other unanticipated (and at this point unclear) sample preparation shortcomings associated with SP3.

**Figure 3.**
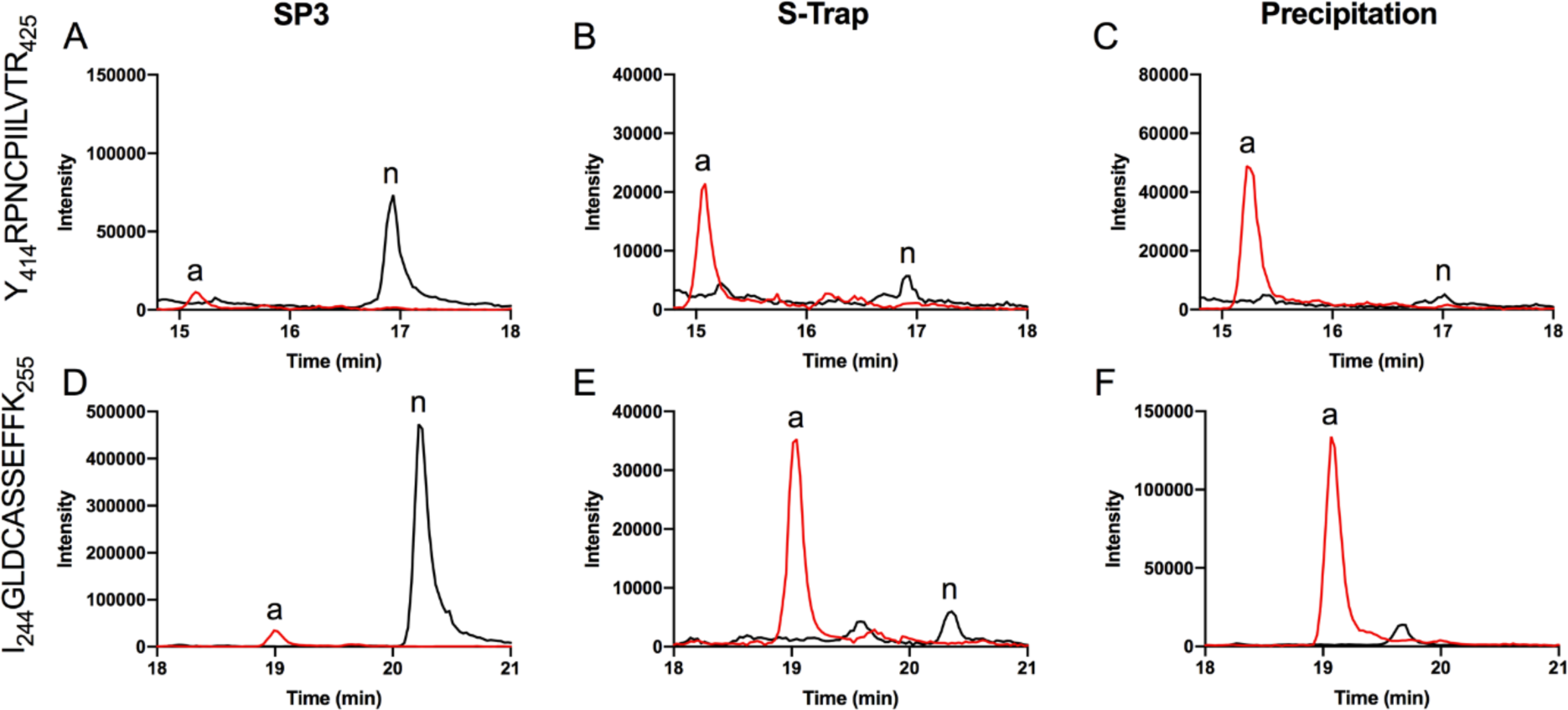
Partial alkylation of cysteine residues with SP3 sample preparation. Yeast whole cell extract was prepared using three different methods. Shown are the extracted ion chromatograms of two cysteine containing peptides identified in the DDA MS data. Alkylated (red line, (a)) and non-alkylated (black line, (n)) Y_414_RPNCPIILVTR_425_ peptide from Pyruvate Kinase 1 (P00549, UniProtKB) when samples were prepared with (**A**) SP3, (**B**) S-Trap, or (**C**) organic solvent precipitation. Alkylated and non-alkylated I_244_GLDCASSEFFK_255_ peptide from Enolase 2 (P00925, UniProtKB) when samples were prepared with (**D**) SP3, (**E**) S-Trap, or (**F**) organic solvent precipitation.

## Conclusions

We compared four bottom-up proteomics sample preparation techniques to identify the optimal method for use with mammalian cell bioreactor culture spent media. We found that several commercial mammalian cell media and media supplements, including CD CHO medium and EfficientFeed B media supplement, contained a polymer with similar MS profiles to Pluronic F68. We found that FASP was unable to remove polymeric surfactants from the samples, that precipitation could partially remove polymeric surfactants, and that SP3 could inconsistently remove polymeric surfactants. During the course of our analyses we observed that SP3 sample preparation was associated with partial alkylation of cysteines, and confirmed this result by analyzing previously published data from other laboratories. While the cause of partial alkylation is unclear and was not investigated in this work, it is a caveat of this technique that should be considered when choosing SP3 as a method for quantitative proteomics. We demonstrate that sample preparation using S-Trap gives consistent, robust, high quality mass spectrometry proteomic results, achieving effective removal of Pluronic F68 and other polymeric contaminants present in mammalian cell culture media.

## Acknowledgements

We thank Dr Amanda Nouwens and Peter Josh from the Mass Spectrometry Facility at the School of Chemistry and Molecular Biosciences of The University of Queensland for their help and advice. LFZ was funded by a Promoting Women Fellowship from the University of Queensland, Australia. BLS was funded by an Australian National Health and Medical Research Council RD Wright Biomedical (CDF Level 2) Fellowship APP1087975. This work was funded by an Australian Research Council Discovery Project DP160102766 to BLS and an Australian Research Council Industrial Transformation Training Centre IC160100027 to BLS, CBH, and YYL.

## Graphical Abstract (For TOC Only)

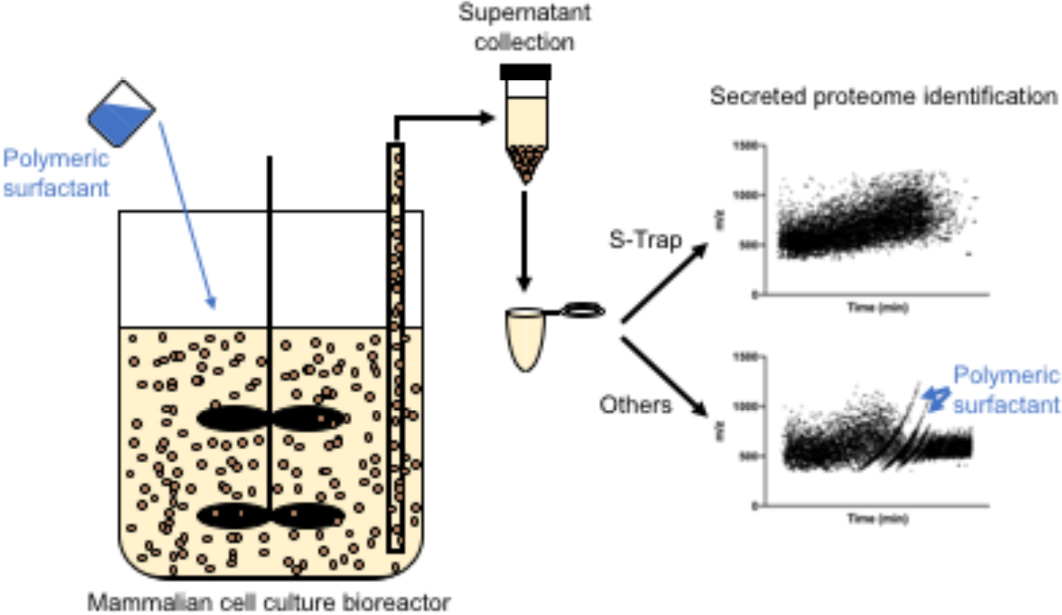

## Supporting information

The following supporting information is available.

**Supplementary Figure S1.** Ineffective removal of polymer with FASP.

**Supplementary Figure S2.** Presence of a polymer in several common mammalian cell media and media supplements.

**Supplementary Figure S3.** Ineffective removal of polymer with FASP in Amicon or Microcon columns.

**Supplementary Figure S4.** Example of a CHO bioreactor supernatant sample prepared with SP3 displaying polymeric contaminant.

**Supplementary Figure S5.** MS/MS spectra of peptides described in Figure 3.

**Supplementary Table S1.** ProteinPilot output of DDA data analysis (Protein and distinct peptides) from bioreactor supernatant samples processed with FASP, S-Trap, SP3, or precipitation.

**Supplementary Table S2.** ProteinPilot output of DDA data analysis from yeast whole cell extract samples processed with S-Trap, SP3, or precipitation.

**Supplementary Table S3.** Re-analysis of published dataset PXD008698 using SP3 sample preparation ^24^ using Preview.

**Supplementary Table S4.** Summary of the different proteomic workflows tested in this work.

