## Supporting Information for "S-Trap eliminates cell culture media polymeric surfactants for effective proteomic analysis of mammalian cell bioreactor supernatants"

**Table of contents**

| Name of file | Page # |
|---|---|
| **Supplementary Figure S1.** Ineffective removal of polymer with FASP. | 2 |
| **Supplementary Figure S2.** Presence of a polymer in several common mammalian cell media and media supplements. | 3 |
| **Supplementary Figure S3.** Ineffective removal of polymer with FASP in Amicon or Microcon columns. | 4 |
| **Supplementary Figure S4.** Example of a sample prepared with SP3 displaying polymeric contaminant. | 5 |
| **Supplementary Figure S5.** MS/MS spectra of peptides described in Figure 3 | 6 |
| **Supplementary Table S1.xls.** ProteinPilot output of DDA data analysis (Protein and distinct peptides) from bioreactor supernatant samples processed with FASP, S-Trap, SP3, or precipitation. | |
| **Supplementary Table S2.xls**. ProteinPilot output of DDA data analysis from yeast whole cell extract samples processed with S-Trap, SP3, or precipitation. | |
| **Supplementary Table S3.pdf**. Re-analysis of published dataset PXD008698 using SP3 sample preparation using Preview. | |
| **Supplementary Table S4.pdf.** Summary of the different proteomic workflows tested in this work. | |

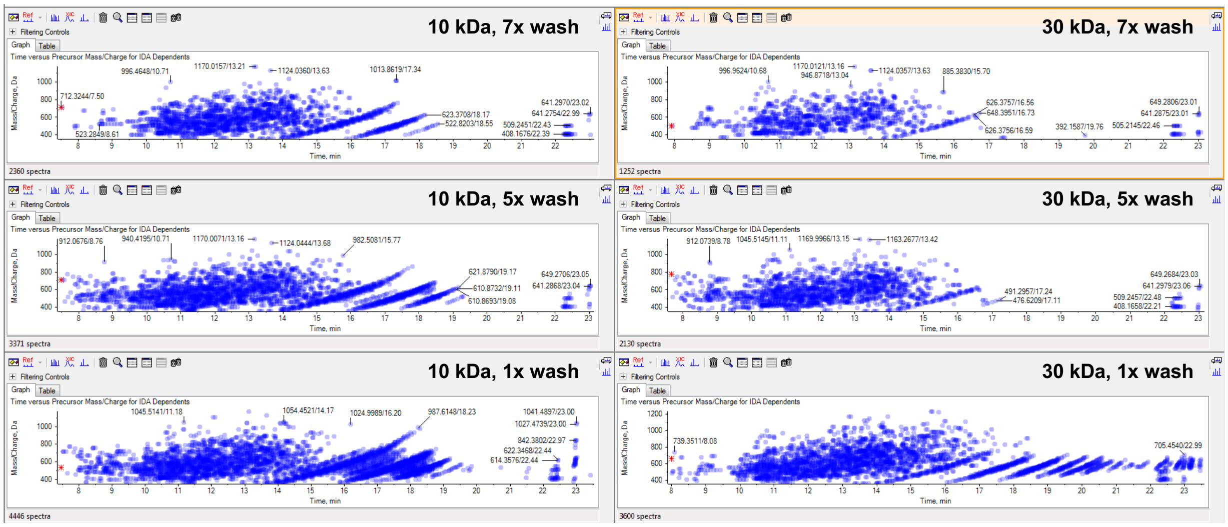


| Sample | Nr of Proteins (>0.05, 10%) | Nr of peptides |
|---|---|---|
| 10kDa, 1x wash | 130 | 538 |
| 10kDa, 5x wash | 128 | 585 |
| 10kDa, 7x wash | 112 | 474 |
| 30kDa, 1x wash | 119 | 701 |
| 30kDa, 5x wash | 114 | 509 |
| 30kDa, 7x wash | 90 | 334 |

**Supplementary Figure S1. Increasing the number of washes of Amicon columns with 50 mM ammonium bicarbonate during FASP does not effectively remove polymeric surfactants but does reduce the number of proteins and peptides identified.** 25 µL of cell-free spent CD CHO medium supplemented with anti-clumping agent and Glutamax was processed using the FASP protocol as described in the Experimental Section, with one, five, or seven washes with 500 µL of 50 mM ammonium bicarbonate. Representation of the fragmented precursors across time (*m/z vs* min). The table below shows the number of proteins and distinct peptides identified with ProteinPilot in each sample.

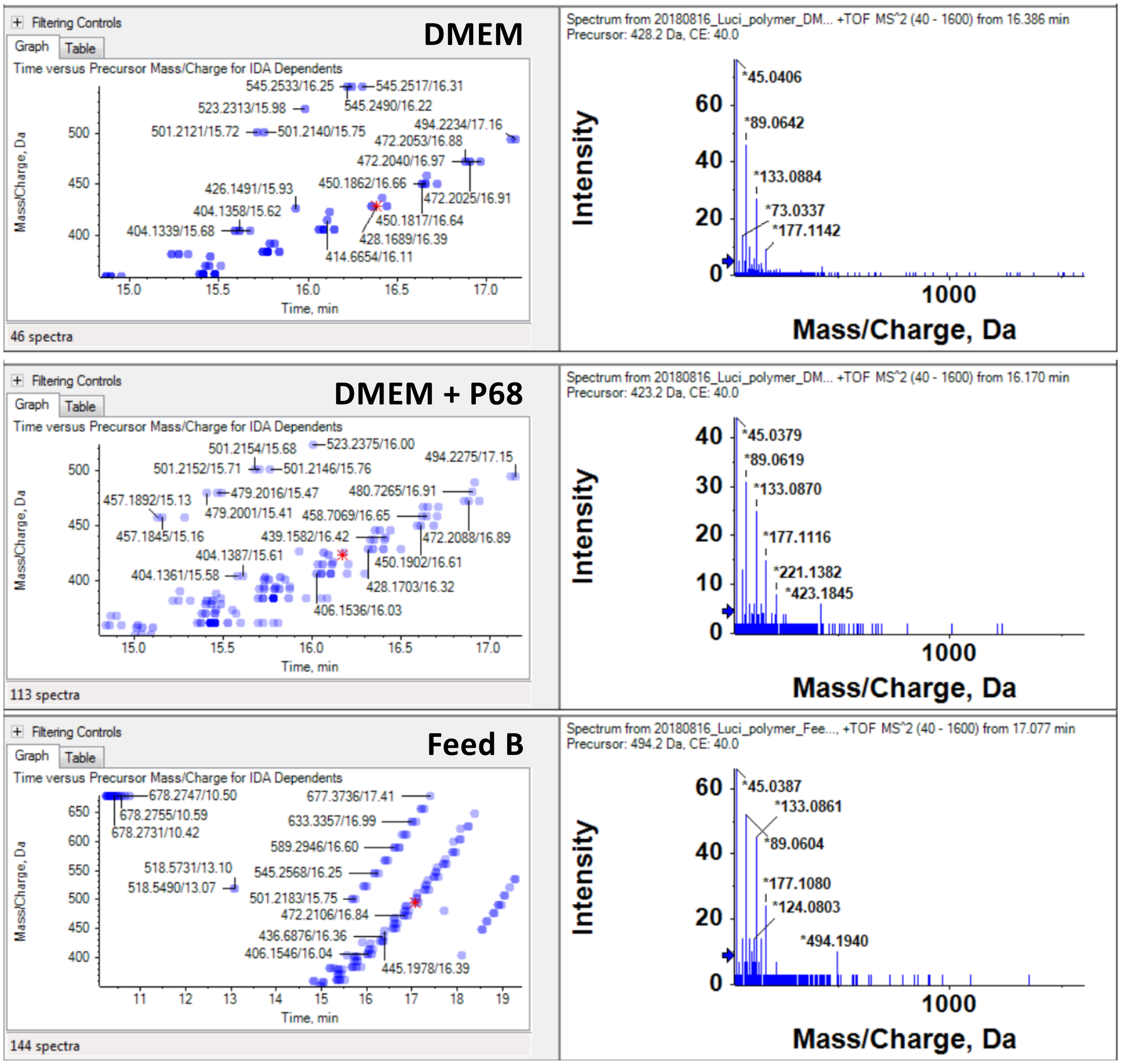


**Supplementary Figure S2. Presence of a polymer with similar MS/MS spectra in DMEM, DMEM supplemented with Pluronic F68, and EfficientFeed B.** DMEM, DMEM supplemented with 1 g/L Pluronic F68 (DMEM + P68), and EfficientFeed B (Feed B) were prepared using the FASP method. Left, representations of the fragmented precursors across time (*m/z vs* min); right, representative MS/MS spectra of the polymeric contaminants.

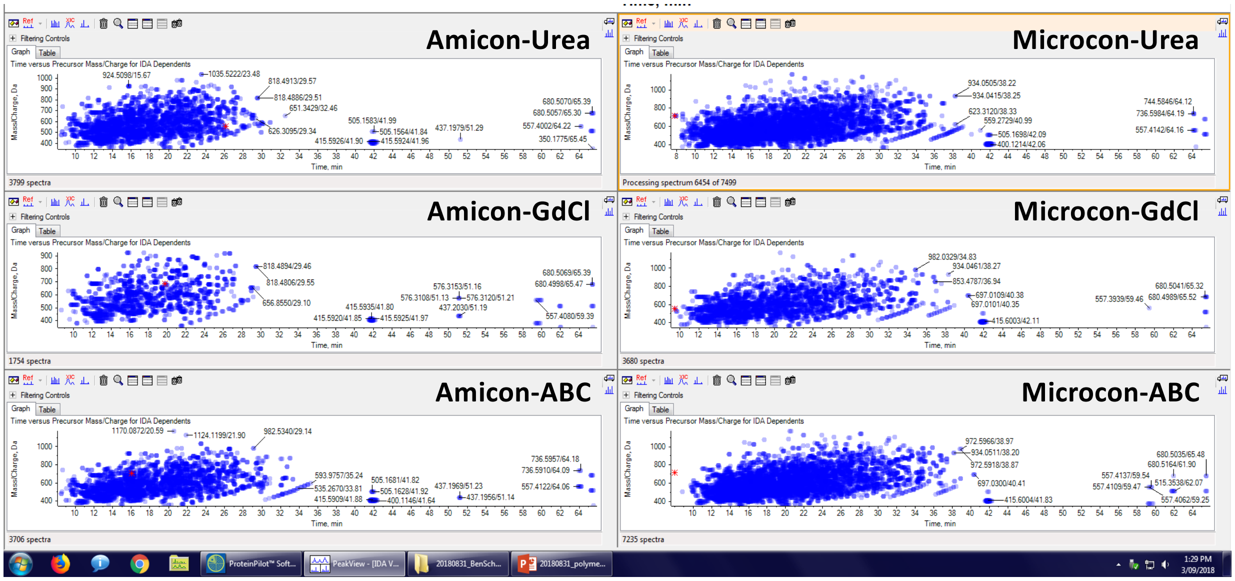


| Sample | Proteins | Peptides |
|---|---|---|
| 11.Amicon_2xABC | 73 | 549 |
| 12.Amicon_2xGdCl | 49 | 253 |
| 13.Amicon_2xUrea | 74 | 510 |
| 14.Microcon_2xABC | 94 | 765 |
| 15.Microcon_2xGdCl | 39 | 316 |
| 16.Microcon_2xUrea | 91 | 793 |

**Supplementary Figure S3. Washing denatured protein samples with strong denaturating buffers does not remove polymer in 10 kDa cut-off Amicon or Ultracel PL-30 Microcon columns.** 25 µL of cell-free spent CD CHO medium supplemented with anti-clumping agent and Glutamax were processed using the FASP protocol as described in the Experimental Section, with the columns washed 2 times with 500 µL of 6 M guanidinium chloride in 50 mM Tris pH 8 (GdCl), 8 M Urea in 50 mM Tris pH 8 (urea), or 50 mM ammonium bicarbonate (ABC) followed by 500 µL ABC before trypsinization. The graphs show the fragmented precursors across time (*m/z vs* min). The table shows the number of proteins and peptides identified in each sample.

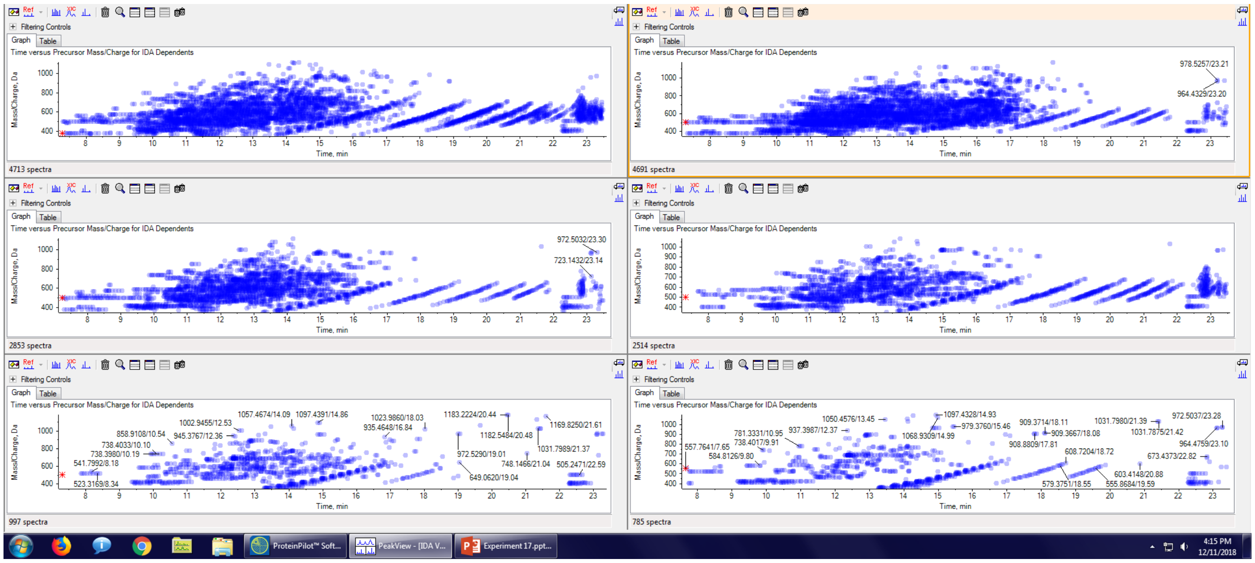

**Supplementary Figure S4. Example of a sample prepared with SP3 displaying polymeric contaminant.** 20 µL (bottom row), 100 µL (middle row), or 200 µL (top row) of filtered bioreactor supernatant sample from CHO cells incubated in CD CHO medium supplemented with EfficientFeed B and 1 gr/L pluronic F68 were prepared using SP3 method. Representations of all fragmented precursors identified in DDA mode for two technical replicates.

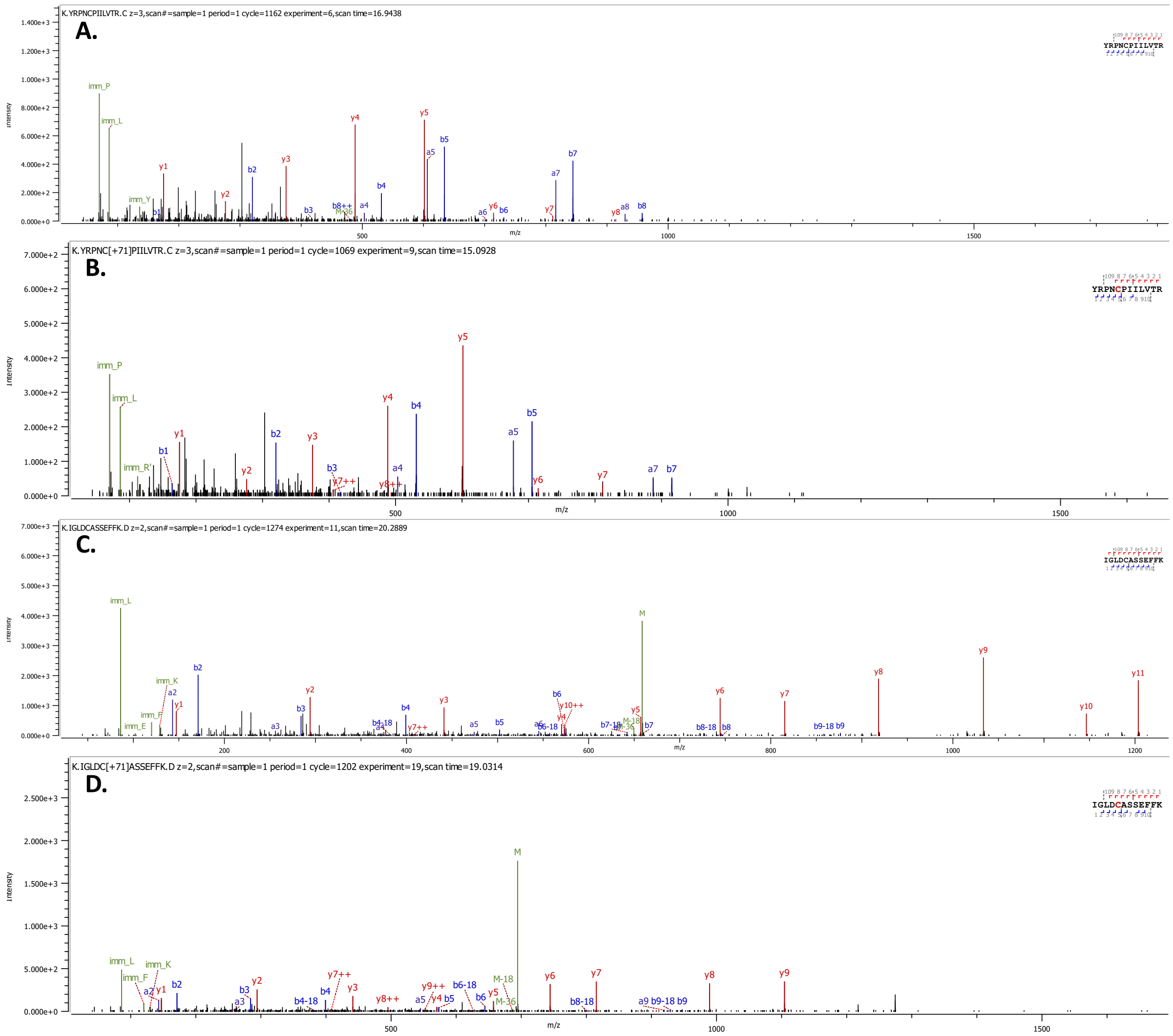


**Supplementary Figure S5. MS/MS spectra of peptide** $Y_{414}RPNCPIILVTR_{425}$ **from protein yeast Pyruvate Kinase 1** (P00549, UniProtKB) **and peptide** $I_{244}GLDCASSEFFK_{255}$ **from yeast protein Enolase 2** (P00925, UniProtKB)**.** Yeast whole cell extracts were prepared with S-Trap or SP3 methods and the samples analysed with a TripleTof 5600 instrument in DDA mode. Shown are the MS/MS spectra of the non-alkylated and alkylated (+71) versions of the peptides identified in the SP3 (A,C) or S-Trap (B,D) samples, respectively. The peptides shown are: (**A**) YRPNCPIILVTR: 482.2746 *m/z*, *z*=3, (**B**) YRPNC(+71)PIILVTR: 505.9537 *m/z*, *z*=3, (**C**) IGLDCASSEFFK: 658.8132 *m/z*, *z*=2, and (**D**) IGLDC(+71)ASSEFFK: 694.3318 *m/z*, *z*=2.

**Supplementary Tables (.xls)**

**Supplementary Table S1.** ProteinPilot output of DDA data analysis (Protein and distinct peptides) from bioreactor supernatant samples (working day 12, CHO cells grown in CD CHO medium supplemented with CHO CD EfficientFeed B) processed with FASP, S-Trap, SP3, or precipitation.

**Supplementary Table S2.** ProteinPilot output of DDA data analysis (Protein and distinct peptides) from yeast whole cell extract samples (yeast grown overnight in YPD medium) processed with S-Trap, SP3, or precipitation. SP3 samples were prepared using a denaturation step of 95 °C for 10 min.

**Supplementary Table S3.** Re-analysis of published data using SP3 sample preparation [24] using Preview. Three samples from the published dataset PXD008698 were downloaded and analyzed in Preview and Byonic.

**Supplementary Table S4.** Summary of the different proteomic workflows tested in Figure 1, 2, and 3 in this work.
