## Supplementary Table S3 for "S-Trap eliminates cell culture media polymeric surfactants for effective proteomic analysis of mammalian cell bioreactor supernatants"

ch_03Feb2018_SP3-Solvents_ethanol-ph8-100p_1.raw_PMi - Preview results 1
ch_03Feb2018_SP3-Solvents_acetone-ph8-100p_1.raw_PMi - Preview results 12
ch_06Feb2018_SP3-Binding-ethanol-100p_mix-yes_rinse-yes_1.raw_PMi - Pre 25

### Preview™ by PROTEIN METRICS Inc.

Summary | Detail

#### Table of Contents (ch_03Feb2018_SP3-Solvents_ethanol-ph8-100p_1.raw) (Schulz_CP_Homo_sapiens_proteomeUP000005640reveiwed_Download_20180420_proteins_20303.fasta)


1. Representative proteins
2. Score distribution
3. Mass measurement errors
4. Cysteine
5. Nonspecific cleavage
6. Oxidation
7. Chemical modifications
8. Posttranslational modifications
9. Computation information


#### Representative proteins

Right click (or option-click) and save link: peptide and protein identifications (in csv format). Or, access the folder.

| Rank | ProtScore | # Spectra ID'd | # Unique Peptides | Database Sequence # | # aa's | Protein Name |
|---|---|---|---|---|---|---|
| 1 | 289.09 | 31 | 29 | 29794 | 1960 | >sp\|P35579\|MYH9_HUMAN Myosin-9 OS=Homo sapiens OX=9606 GN=MYH9 PE=1 SV=4 |
| 2 | 268.01 | 33 | 29 | 14876 | 724 | >sp\|P08238\|HS90B_HUMAN Heat shock protein HSP 90-beta OS=Homo sapiens OX=9606 GN=HSP90AB1 PE=1 SV=4 |
| 3 | 246.87 | 28 | 23 | 5194 | 654 | >sp\|P11021\|BIP_HUMAN Endoplasmic reticulum chaperone BiP OS=Homo sapiens OX=9606 GN=HSPA5 PE=1 SV=2 |
| 4 | 237.74 | 32 | 24 | 12650 | 641 | >sp\|P0DMV9\|HS71B_HUMAN Heat shock 70 kDa protein 1B OS=Homo sapiens OX=9606 GN=HSPA1B PE=1 SV=1 |
| 5 | 225.42 | 34 | 28 | 10117 | 858 | >sp\|P13639\|EF2_HUMAN Elongation factor 2 OS=Homo sapiens OX=9606 GN=EEF2 PE=1 SV=4 |
| 6 | 196.76 | 25 | 20 | 11486 | 646 | >sp\|P11142\|HSP7C_HUMAN Heat shock cognate 71 kDa protein OS=Homo sapiens OX=9606 GN=HSPA8 PE=1 SV=1 |
| 7 | 195.66 | 26 | 24 | 41440 | 466 | >sp\|P08670\|VIME_HUMAN Vimentin OS=Homo sapiens OX=9606 GN=VIM PE=1 SV=4 |
| 8 | 180.20 | 25 | 19 | 34832 | 445 | >sp\|P68371\|TBB4B_HUMAN Tubulin beta-4B chain OS=Homo sapiens OX=9606 GN=TUBB4B PE=1 SV=1 |
| 9 | 177.02 | 29 | 23 | 17426 | 2639 | >sp\|P21333-2\|FLNA_HUMAN Isoform 2 of Filamin-A OS=Homo sapiens OX=9606 GN=FLNA |
| 10 | 172.69 | 19 | 19 | 38994 | 806 | >sp\|P55072\|TERA_HUMAN Transitional endoplasmic reticulum ATPase OS=Homo sapiens OX=9606 GN=VCP PE=1 SV=4 |
| 11 | 163.23 | 19 | 18 | 27671 | 2101 | >sp\|Q14980-2\|NUMA1_HUMAN Isoform 2 of Nuclear mitotic apparatus protein 1 OS=Homo sapiens OX=9606 GN=NUMA1 |
| 12 | 162.70 | 20 | 17 | 26540 | 586 | >sp\|P20700\|LMNB1_HUMAN Lamin-B1 OS=Homo sapiens OX=9606 GN=LMNB1 PE=1 SV=2 |
| 13 | 162.47 | 22 | 18 | 10608 | 434 | >sp\|P06733\|ENOA_HUMAN Alpha-enolase OS=Homo sapiens OX=9606 GN=ENO1 PE=1 SV=2 |
| 14 | 161.87 | 24 | 16 | 34144 | 375 | >sp\|P60709\|ACTB_HUMAN Actin; cytoplasmic 1 OS=Homo sapiens OX=9606 GN=ACTB PE=1 SV=1 |
| 15 | 154.76 | 33 | 31 | 3268 | 5890 | >sp\|Q09666\|AHNK_HUMAN Neuroblast differentiation-associated protein AHNAK OS=Homo sapiens OX=9606 GN=AHNAK PE=1 SV=2 |
| 16 | 153.92 | 15 | 14 | 6958 | 652 | >sp\|Q92841-3\|DDX17_HUMAN Isoform 4 of Probable ATP-dependent RNA helicase DDX17 OS=Homo sapiens OX=9606 GN=DDX17 |
| 17 | 145.58 | 13 | 13 | 39061 | 545 | >sp\|P49368\|TCPG_HUMAN T-complex protein 1 subunit gamma OS=Homo sapiens OX=9606 GN=CCT3 PE=1 SV=4 |
| 18 | 144.27 | 16 | 15 | 35062 | 488 | >sp\|P78371-2\|TCPB_HUMAN Isoform 2 of T-complex protein 1 subunit beta OS=Homo sapiens OX=9606 GN=CCT2 |
| 19 | 140.50 | 19 | 19 | 17366 | 2511 | >sp\|P49327\|FAS_HUMAN Fatty acid synthase OS=Homo sapiens OX=9606 GN=FASN PE=1 SV=3 |
| 20 | 137.69 | 18 | 16 | 27856 | 1014 | >sp\|P09874\|PARP1_HUMAN Poly [ADP-ribose] polymerase 1 OS=Homo sapiens OX=9606 GN=PARP1 PE=1 SV=4 |
| 21 | 130.82 | 16 | 11 | 40882 | 248 | >sp\|P06753-5\|TPM3_HUMAN Isoform 5 of Tropomyosin alpha-3 chain OS=Homo sapiens OX=9606 GN=TPM3 |
| 22 | 130.78 | 17 | 14 | 12628 | 806 | >sp\|Q00839-2\|HNRPU_HUMAN Isoform 2 of Heterogeneous nuclear ribonucleoprotein U OS=Homo sapiens OX=9606 GN=HNRNPU |
| 23 | 128.60 | 22 | 19 | 17521 | 803 | >sp\|P14625\|ENPL_HUMAN Endoplasmin OS=Homo sapiens OX=9606 GN=HSP90B1 PE=1 SV=1 |
| 24 | 128.58 | 15 | 15 | 38834 | 529 | >sp\|P50990-2\|TCPQ_HUMAN Isoform 2 of T-complex protein 1 subunit theta OS=Homo sapiens OX=9606 GN=CCT8 |
| 25 | 126.13 | 19 | 18 | 30819 | 710 | >sp\|P19338\|NUCL_HUMAN Nucleolin OS=Homo sapiens OX=9606 GN=NCL PE=1 SV=3 |
| 26 | 124.78 | 10 | 10 | 199 | 553 | >sp\|P25705\|ATPA_HUMAN ATP synthase subunit alpha; mitochondrial OS=Homo sapiens OX=9606 GN=ATP5F1A PE=1 SV=1 |
| 27 | 124.46 | 20 | 17 | 14875 | 854 | >sp\|P07900-2\|HS90A_HUMAN Isoform 2 of Heat shock protein HSP 90-alpha OS=Homo sapiens OX=9606 GN=HSP90AA1 |
| 28 | 123.89 | 20 | 18 | 5883 | 573 | >sp\|P10809\|CH60_HUMAN 60 kDa heat shock protein; mitochondrial OS=Homo sapiens OX=9606 GN=HSPD1 PE=1 SV=2 |
| 29 | 122.43 | 15 | 13 | 13137 | 334 | >sp\|P07195\|LDHB_HUMAN L-lactate dehydrogenase B chain OS=Homo sapiens OX=9606 GN=LDHB PE=1 SV=2 |
| 30 | 122.31 | 17 | 15 | 34358 | 911 | >sp\|O43707\|ACTN4_HUMAN Alpha-actinin-4 OS=Homo sapiens OX=9606 GN=ACTN4 PE=1 SV=2 |
| 31 | 119.98 | 11 | 10 | 2722 | 529 | >sp\|P06576\|ATPB_HUMAN ATP synthase subunit beta; mitochondrial OS=Homo sapiens OX=9606 GN=ATP5F1B PE=1 SV=3 |
| 32 | 116.43 | 17 | 14 | 11476 | 817 | >sp\|Q92598-3\|HS105_HUMAN Isoform 3 of Heat shock protein 105 kDa OS=Homo sapiens OX=9606 GN=HSPH1 |
| 33 | 115.97 | 24 | 20 | 12573 | 691 | >sp\|P52272-2\|HNRPM_HUMAN Isoform 2 of Heterogeneous nuclear ribonucleoprotein M OS=Homo sapiens OX=9606 GN=HNRNPM |
| 34 | 115.76 | 20 | 18 | 7399 | 4646 | >sp\|Q14204\|DYHC1_HUMAN Cytoplasmic dynein 1 heavy chain 1 OS=Homo sapiens OX=9606 GN=DYNC1H1 PE=1 SV=5 |
| 35 | 115.37 | 15 | 14 | 7556 | 662 | >sp\|O00571\|DDX3X_HUMAN ATP-dependent RNA helicase DDX3X OS=Homo sapiens OX=9606 GN=DDX3X PE=1 SV=3 |
| 36 | 115.30 | 12 | 11 | 3770 | 255 | >sp\|P62258\|1433E_HUMAN 14-3-3 protein epsilon OS=Homo sapiens OX=9606 GN=YWHAE PE=1 SV=1 |
| 37 | 114.42 | 15 | 13 | 11112 | 711 | >sp\|Q92945\|FUBP2_HUMAN Far upstream element-binding protein 2 OS=Homo sapiens OX=9606 GN=KHSRP PE=1 SV=4 |
| 38 | 109.91 | 17 | 16 | 35928 | 590 | >sp\|P31948-2\|STIP1_HUMAN Isoform 2 of Stress-induced-phosphoprotein 1 OS=Homo sapiens OX=9606 GN=STIP1 |
| 39 | 105.21 | 11 | 10 | 12288 | 293 | >sp\|P07910-2\|HNRPC_HUMAN Isoform C1 of Heterogeneous nuclear ribonucleoproteins C1/C2 OS=Homo sapiens OX=9606 GN=HNRNPC |
| 40 | 105.19 | 15 | 13 | 38968 | 448 | >sp\|P48643-2\|TCPE_HUMAN Isoform 2 of T-complex protein 1 subunit epsilon OS=Homo sapiens OX=9606 GN=CCT5 |
| 41 | 104.73 | 15 | 10 | 14006 | 103 | >sp\|P62805\|H4_HUMAN Histone H4 OS=Homo sapiens OX=9606 GN=HIST1H4A PE=1 SV=2 |
| 42 | 104.51 | 11 | 10 | 11581 | 898 | >sp\|Q12906-7\|ILF3_HUMAN Isoform 7 of Interleukin enhancer-binding factor 3 OS=Homo sapiens OX=9606 GN=ILF3 |

| | | | | | | |
|---|---|---|---|---|---|---|
| 43 | 102.97 | 14 | 12 | 16301 | 406 | >sp\|P60842\|IF4A1_HUMAN Eukaryotic initiation factor 4A-I OS=Homo sapiens OX=9606 GN=EIF4A1 PE=1 SV=1 |
| 44 | 102.72 | 15 | 14 | 16431 | 430 | >sp\|P05783\|K1C18_HUMAN Keratin; type I cytoskeletal 18 OS=Homo sapiens OX=9606 GN=KRT18 PE=1 SV=2 |
| 45 | 100.85 | 11 | 10 | 23683 | 2452 | >sp\|Q13813-3\|SPTN1_HUMAN Isoform 3 of Spectrin alpha chain; non-erythrocytic 1 OS=Homo sapiens OX=9606 GN=SPTAN1 |
| 46 | 100.64 | 12 | 11 | 14295 | 679 | >sp\|P38646\|GRP75_HUMAN Stress-70 protein; mitochondrial OS=Homo sapiens OX=9606 GN=HSPA9 PE=1 SV=2 |
| 47 | 100.00 | 11 | 11 | 3913 | 364 | >sp\|P04075\|ALDOA_HUMAN Fructose-bisphosphate aldolase A OS=Homo sapiens OX=9606 GN=ALDOA PE=1 SV=2 |
| 48 | 99.89 | 10 | 10 | 37650 | 962 | >sp\|Q15029-3\|U5S1_HUMAN Isoform 3 of 116 kDa U5 small nuclear ribonucleoprotein component OS=Homo sapiens OX=9606 GN=EFTUD2 |
| 49 | 99.21 | 18 | 16 | 17290 | 1382 | >sp\|Q14152\|EIF3A_HUMAN Eukaryotic translation initiation factor 3 subunit A OS=Homo sapiens OX=9606 GN=EIF3A PE=1 SV=1 |
| 50 | 98.36 | 16 | 16 | 42235 | 4128 | >sp\|P78527\|PRKDC_HUMAN DNA-dependent protein kinase catalytic subunit OS=Homo sapiens OX=9606 GN=PRKDC PE=1 SV=3 |
| 51 | 96.36 | 15 | 12 | 28584 | 505 | >sp\|P30101\|PDIA3_HUMAN Protein disulfide-isomerase A3 OS=Homo sapiens OX=9606 GN=PDIA3 PE=1 SV=4 |
| 52 | 96.36 | 13 | 10 | 40876 | 249 | >sp\|P60174-1\|TPIS_HUMAN Isoform 2 of Triosephosphate isomerase OS=Homo sapiens OX=9606 GN=TPI1 |
| 53 | 95.73 | 14 | 13 | 37885 | 1018 | >sp\|P22314-2\|UBA1_HUMAN Isoform 2 of Ubiquitin-like modifier-activating enzyme 1 OS=Homo sapiens OX=9606 GN=UBA1 |
| 54 | 94.09 | 14 | 13 | 18698 | 382 | >sp\|Q15233-2\|NONO_HUMAN Isoform 2 of Non-POU domain-containing octamer-binding protein OS=Homo sapiens OX=9606 GN=NONO |
| 55 | 93.83 | 14 | 12 | 29243 | 199 | >sp\|Q06830\|PRDX1_HUMAN Peroxiredoxin-1 OS=Homo sapiens OX=9606 GN=PRDX1 PE=1 SV=1 |
| 56 | 90.78 | 11 | 9 | 34464 | 245 | >sp\|P63104\|1433Z_HUMAN 14-3-3 protein zeta/delta OS=Homo sapiens OX=9606 GN=YWHAZ PE=1 SV=1 |
| 57 | 90.32 | 11 | 11 | 38884 | 835 | >sp\|Q13263\|TIF1B_HUMAN Transcription intermediary factor 1-beta OS=Homo sapiens OX=9606 GN=TRIM28 PE=1 SV=5 |
| 58 | 90.29 | 13 | 12 | 14882 | 840 | >sp\|P34932\|HSP74_HUMAN Heat shock 70 kDa protein 4 OS=Homo sapiens OX=9606 GN=HSPA4 PE=1 SV=4 |
| 59 | 89.93 | 17 | 15 | 27434 | 417 | >sp\|P00558\|PGK1_HUMAN Phosphoglycerate kinase 1 OS=Homo sapiens OX=9606 GN=PGK1 PE=1 SV=3 |
| 60 | 88.72 | 12 | 11 | 16442 | 511 | >sp\|P05787-2\|K2C8_HUMAN Isoform 2 of Keratin; type II cytoskeletal 8 OS=Homo sapiens OX=9606 GN=KRT8 |
| 61 | 88.63 | 18 | 14 | 7610 | 1270 | >sp\|Q08211\|DHX9_HUMAN ATP-dependent RNA helicase A OS=Homo sapiens OX=9606 GN=DHX9 PE=1 SV=4 |
| 62 | 86.99 | 18 | 17 | 32061 | 577 | >sp\|P26038\|MOES_HUMAN Moesin OS=Homo sapiens OX=9606 GN=MSN PE=1 SV=3 |
| 63 | 86.89 | 12 | 9 | 12431 | 588 | >sp\|O60506-2\|HNRPQ_HUMAN Isoform 2 of Heterogeneous nuclear ribonucleoprotein Q OS=Homo sapiens OX=9606 GN=SYNCRIP |
| 64 | 85.89 | 12 | 12 | 4219 | 935 | >sp\|P11586\|C1TC_HUMAN C-1-tetrahydrofolate synthase; cytoplasmic OS=Homo sapiens OX=9606 GN=MTHFD1 PE=1 SV=3 |
| 65 | 85.89 | 9 | 9 | 663 | 992 | >sp\|P05023-3\|AT1A1_HUMAN Isoform 3 of Sodium/potassium-transporting ATPase subunit alpha-1 OS=Homo sapiens OX=9606 GN=ATP1A1 |
| 66 | 85.06 | 11 | 9 | 13423 | 531 | >sp\|P14618-2\|KPYM_HUMAN Isoform M1 of Pyruvate kinase PKM OS=Homo sapiens OX=9606 GN=PKM |
| 67 | 85.05 | 12 | 11 | 22202 | 353 | >sp\|P22626\|ROA2_HUMAN Heterogeneous nuclear ribonucleoproteins A2/B1 OS=Homo sapiens OX=9606 GN=HNRNPA2B1 PE=1 SV=2 |
| 68 | 85.00 | 10 | 8 | 17245 | 353 | >sp\|Q15717-2\|ELAV1_HUMAN Isoform 2 of ELAV-like protein 1 OS=Homo sapiens OX=9606 GN=ELAVL1 |
| 69 | 84.25 | 15 | 11 | 8068 | 535 | >sp\|P17844-2\|DDX5_HUMAN Isoform 2 of Probable ATP-dependent RNA helicase DDX5 OS=Homo sapiens OX=9606 GN=DDX5 |
| 70 | 82.82 | 9 | 8 | 28314 | 645 | >sp\|P13667\|PDIA4_HUMAN Protein disulfide-isomerase A4 OS=Homo sapiens OX=9606 GN=PDIA4 PE=1 SV=2 |
| 71 | 81.55 | 9 | 8 | 19342 | 463 | >sp\|Q9Y230\|RUVB2_HUMAN RuvB-like 2 OS=Homo sapiens OX=9606 GN=RUVBL2 PE=1 SV=3 |
| 72 | 81.48 | 8 | 8 | 22201 | 267 | >sp\|P09651-3\|ROA1_HUMAN Isoform 2 of Heterogeneous nuclear ribonucleoprotein A1 OS=Homo sapiens OX=9606 GN=HNRNPA1 |
| 73 | 81.02 | 11 | 10 | 38249 | 499 | >sp\|Q99832-3\|TCPH_HUMAN Isoform 3 of T-complex protein 1 subunit eta OS=Homo sapiens OX=9606 GN=CCT7 |
| 74 | 80.93 | 7 | 7 | 27427 | 254 | >sp\|P18669\|PGAM1_HUMAN Phosphoglycerate mutase 1 OS=Homo sapiens OX=9606 GN=PGAM1 PE=1 SV=2 |
| 75 | 79.81 | 11 | 10 | 13288 | 361 | >sp\|P00338-3\|LDHA_HUMAN Isoform 3 of L-lactate dehydrogenase A chain OS=Homo sapiens OX=9606 GN=LDHA |
| 76 | 79.00 | 12 | 11 | 41398 | 1066 | >sp\|P18206-2\|VINC_HUMAN Isoform 1 of Vinculin OS=Homo sapiens OX=9606 GN=VCL |
| 77 | 78.91 | 14 | 10 | 34825 | 416 | >sp\|Q71U36-2\|TBA1A_HUMAN Isoform 2 of Tubulin alpha-1A chain OS=Homo sapiens OX=9606 GN=TUBA1A |
| 78 | 78.14 | 13 | 10 | 12512 | 440 | >sp\|P61978-3\|HNRPK_HUMAN Isoform 3 of Heterogeneous nuclear ribonucleoprotein K OS=Homo sapiens OX=9606 GN=HNRNPK |
| 79 | 77.69 | 12 | 12 | 41146 | 2541 | >sp\|Q9Y490\|TLN1_HUMAN Talin-1 OS=Homo sapiens OX=9606 GN=TLN1 PE=1 SV=3 |
| 80 | 77.17 | 9 | 8 | 20532 | 707 | >sp\|P23246\|SFPQ_HUMAN Splicing factor; proline- and glutamine-rich OS=Homo sapiens OX=9606 GN=SFPQ PE=1 SV=2 |
| 81 | 76.87 | 14 | 11 | 28052 | 393 | >sp\|Q8NC51-3\|PAIRB_HUMAN Isoform 3 of Plasminogen activator inhibitor 1 RNA-binding protein OS=Homo sapiens OX=9606 GN=SERBP1 |
| 82 | 76.66 | 16 | 13 | 9793 | 966 | >sp\|Q14697-2\|GANAB_HUMAN Isoform 2 of Neutral alpha-glucosidase AB OS=Homo sapiens OX=9606 GN=GANAB |
| 83 | 76.42 | 8 | 6 | 19742 | 432 | >sp\|P23526\|SAHH_HUMAN Adenosylhomocysteinase OS=Homo sapiens OX=9606 GN=AHCY PE=1 SV=4 |
| 84 | 76.20 | 7 | 4 | 22777 | 110 | >sp\|P06454-2\|PTMA_HUMAN Isoform 2 of Prothymosin alpha OS=Homo sapiens OX=9606 GN=PTMA |
| 85 | 75.90 | 10 | 10 | 16289 | 381 | >sp\|P12277\|KCRB_HUMAN Creatine kinase B-type OS=Homo sapiens OX=9606 GN=CKB PE=1 SV=1 |
| 86 | 74.52 | 7 | 7 | 22096 | 295 | >sp\|P08865\|RSSA_HUMAN 40S ribosomal protein SA OS=Homo sapiens OX=9606 GN=RPSA PE=1 SV=4 |
| 87 | 73.79 | 9 | 7 | 38832 | 556 | >sp\|P17987\|TCPA_HUMAN T-complex protein 1 subunit alpha OS=Homo sapiens OX=9606 GN=TCP1 PE=1 SV=1 |
| 88 | 72.62 | 12 | 11 | 7519 | 462 | >sp\|Q5VTE0\|EF1A3_HUMAN Putative elongation factor 1-alpha-like 3 OS=Homo sapiens OX=9606 GN=EEF1A1P5 PE=5 SV=1 |
| 89 | 72.37 | 14 | 12 | 27271 | 585 | >sp\|P13797-3\|PLST_HUMAN Isoform 3 of Plastin-3 OS=Homo sapiens OX=9606 GN=PLS3 |
| 90 | 71.15 | 7 | 6 | 27451 | 187 | >sp\|P30086\|PEBP1_HUMAN Phosphatidylethanolamine-binding protein 1 OS=Homo sapiens OX=9606 GN=PEBP1 PE=1 SV=3 |
| 91 | 70.43 | 8 | 7 | 12429 | 287 | >sp\|Q14103-4\|HNRPD_HUMAN Isoform 4 of Heterogeneous nuclear ribonucleoprotein D0 OS=Homo sapiens OX=9606 GN=HNRNPD |
| 92 | 69.85 | 10 | 10 | 12860 | 694 | >sp\|P42166\|LAP2A_HUMAN Lamina-associated polypeptide 2; isoform alpha OS=Homo sapiens OX=9606 GN=TMPO PE=1 SV=2 |
| 93 | 69.33 | 13 | 12 | 21886 | 264 | >sp\|P61247\|RS3A_HUMAN 40S ribosomal protein S3a OS=Homo sapiens OX=9606 GN=RPS3A PE=1 SV=2 |
| 94 | 69.13 | 8 | 7 | 28888 | 919 | >sp\|P55786\|PSA_HUMAN Puromycin-sensitive aminopeptidase OS=Homo sapiens OX=9606 GN=NPEPPS PE=1 SV=2 |
| 95 | 69.00 | 12 | 11 | 18997 | 391 | >sp\|P38159\|RBMX_HUMAN RNA-binding motif protein; X chromosome OS=Homo sapiens OX=9606 GN=RBMX PE=1 SV=3 |
| 96 | 68.98 | 12 | 10 | 5624 | 1639 | >sp\|Q00610-2\|CLH1_HUMAN Isoform 2 of Clathrin heavy chain 1 OS=Homo sapiens OX=9606 GN=CLTC |
| 97 | 68.49 | 8 | 8 | 19134 | 211 | >sp\|P26373\|RL13_HUMAN 60S ribosomal protein L13 OS=Homo sapiens OX=9606 GN=RPL13 PE=1 SV=4 |
| 98 | 68.20 | 8 | 7 | 42044 | 317 | >sp\|P63244\|RACK1_HUMAN Receptor of activated protein C kinase 1 OS=Homo sapiens OX=9606 GN=RACK1 PE=1 SV=3 |
| 99 | 68.09 | 11 | 9 | 9850 | 525 | >sp\|P14314-2\|GLU2B_HUMAN Isoform 2 of Glucosidase 2 subunit beta OS=Homo sapiens OX=9606 GN=PRKCSH |
| 100 | 68.01 | 12 | 11 | 21381 | 248 | >sp\|P18124\|RL7_HUMAN 60S ribosomal protein L7 OS=Homo sapiens OX=9606 GN=RPL7 PE=1 SV=1 |

| | | | | | | |
|---|---|---|---|---|---|---|
| 101 | 67.98 | 7 | 7 | 21436 | 255 | >sp\|P05388-2\|RLA0_HUMAN Isoform 2 of 60S acidic ribosomal protein P0 OS=Homo sapiens OX=9606 GN=RPLP0 |
| 102 | 67.93 | 6 | 6 | 37210 | 324 | >sp\|P67809\|YBOX1_HUMAN Nuclease-sensitive element-binding protein 1 OS=Homo sapiens OX=9606 GN=YBX1 PE=1 SV=3 |
| 103 | 67.30 | 6 | 6 | 37757 | 223 | >sp\|P09936\|UCHL1_HUMAN Ubiquitin carboxyl-terminal hydrolase isozyme L1 OS=Homo sapiens OX=9606 GN=UCHL1 PE=1 SV=2 |
| 104 | 67.24 | 7 | 7 | 7218 | 257 | >sp\|P29692-3\|EF1D_HUMAN Isoform 3 of Elongation factor 1-delta OS=Homo sapiens OX=9606 GN=EEF1D |
| 105 | 66.06 | 7 | 7 | 42264 | 908 | >sp\|Q13200\|PSMD2_HUMAN 26S proteasome non-ATPase regulatory subunit 2 OS=Homo sapiens OX=9606 GN=PSMD2 PE=1 SV=3 |
| 106 | 65.73 | 8 | 7 | 9596 | 483 | >sp\|P34897-3\|GLYM_HUMAN Isoform 3 of Serine hydroxymethyltransferase; mitochondrial OS=Homo sapiens OX=9606 GN=SHMT2 |
| 107 | 64.79 | 8 | 7 | 29443 | 271 | >sp\|Q13162\|PRDX4_HUMAN Peroxiredoxin-4 OS=Homo sapiens OX=9606 GN=PRDX4 PE=1 SV=1 |
| 108 | 64.00 | 9 | 7 | 21380 | 297 | >sp\|P46777\|RL5_HUMAN 60S ribosomal protein L5 OS=Homo sapiens OX=9606 GN=RPL5 PE=1 SV=3 |
| 109 | 63.86 | 8 | 7 | 41436 | 283 | >sp\|P21796\|VDAC1_HUMAN Voltage-dependent anion-selective channel protein 1 OS=Homo sapiens OX=9606 GN=VDAC1 PE=1 SV=2 |
| 110 | 63.61 | 11 | 10 | 22188 | 145 | >sp\|P39019\|RS19_HUMAN 40S ribosomal protein S19 OS=Homo sapiens OX=9606 GN=RPS19 PE=1 SV=2 |
| 111 | 63.17 | 11 | 10 | 41256 | 609 | >sp\|P12956\|XRCC6_HUMAN X-ray repair cross-complementing protein 6 OS=Homo sapiens OX=9606 GN=XRCC6 PE=1 SV=2 |
| 112 | 62.57 | 8 | 7 | 29326 | 165 | >sp\|P62937\|PPIA_HUMAN Peptidyl-prolyl cis-trans isomerase A OS=Homo sapiens OX=9606 GN=PPIA PE=1 SV=2 |
| 113 | 62.48 | 12 | 11 | 30785 | 636 | >sp\|P11940\|PABP1_HUMAN Polyadenylate-binding protein 1 OS=Homo sapiens OX=9606 GN=PABPC1 PE=1 SV=2 |
| 114 | 62.21 | 7 | 6 | 26393 | 614 | >sp\|P02545-6\|LMNA_HUMAN Isoform 6 of Prelamin-A/C OS=Homo sapiens OX=9606 GN=LMNA |
| 115 | 62.04 | 7 | 6 | 19303 | 293 | >sp\|P15880\|RS2_HUMAN 40S ribosomal protein S2 OS=Homo sapiens OX=9606 GN=RPS2 PE=1 SV=2 |
| 116 | 61.92 | 8 | 8 | 40575 | 651 | >sp\|Q12931-2\|TRAP1_HUMAN Isoform 2 of Heat shock protein 75 kDa; mitochondrial OS=Homo sapiens OX=9606 GN=TRAP1 |
| 117 | 61.81 | 8 | 7 | 29283 | 398 | >sp\|P62195-2\|PRS8_HUMAN Isoform 2 of 26S proteasome regulatory subunit 8 OS=Homo sapiens OX=9606 GN=PSMC5 |
| 118 | 61.28 | 8 | 7 | 7279 | 443 | >sp\|Q13838-2\|DX39B_HUMAN Isoform 2 of Spliceosome RNA helicase DDX39B OS=Homo sapiens OX=9606 GN=DDX39B |
| 119 | 60.73 | 10 | 10 | 29644 | 1394 | >sp\|P42704\|LPPRC_HUMAN Leucine-rich PPR motif-containing protein; mitochondrial OS=Homo sapiens OX=9606 GN=LRPPRC PE=1 SV=3 |
| 120 | 60.28 | 9 | 9 | 21434 | 427 | >sp\|P36578\|RL4_HUMAN 60S ribosomal protein L4 OS=Homo sapiens OX=9606 GN=RPL4 PE=1 SV=5 |
| 121 | 60.24 | 10 | 8 | 5190 | 417 | >sp\|P27797\|CALR_HUMAN Calreticulin OS=Homo sapiens OX=9606 GN=CALR PE=1 SV=1 |
| 122 | 60.19 | 9 | 8 | 38497 | 531 | >sp\|P40227\|TCPZ_HUMAN T-complex protein 1 subunit zeta OS=Homo sapiens OX=9606 GN=CCT6A PE=1 SV=3 |
| 123 | 59.93 | 9 | 8 | 36277 | 513 | >sp\|Q14247-3\|SRC8_HUMAN Isoform 3 of Src substrate cortactin OS=Homo sapiens OX=9606 GN=CTTN |
| 124 | 59.04 | 12 | 8 | 12654 | 335 | >sp\|P04406\|G3P_HUMAN Glyceraldehyde-3-phosphate dehydrogenase OS=Homo sapiens OX=9606 GN=GAPDH PE=1 SV=3 |
| 125 | 58.49 | 7 | 7 | 19252 | 135 | >sp\|P08708\|RS17_HUMAN 40S ribosomal protein S17 OS=Homo sapiens OX=9606 GN=RPS17 PE=1 SV=2 |
| 126 | 58.41 | 9 | 8 | 15545 | 569 | >sp\|P06744-2\|G6PI_HUMAN Isoform 2 of Glucose-6-phosphate isomerase OS=Homo sapiens OX=9606 GN=GPI |
| 127 | 58.11 | 8 | 8 | 3563 | 430 | >sp\|P00505\|AATM_HUMAN Aspartate aminotransferase; mitochondrial OS=Homo sapiens OX=9606 GN=GOT2 PE=1 SV=3 |
| 128 | 57.80 | 13 | 10 | 30780 | 394 | >sp\|Q9UQ80\|PA2G4_HUMAN Proliferation-associated protein 2G4 OS=Homo sapiens OX=9606 GN=PA2G4 PE=1 SV=3 |
| 129 | 57.58 | 8 | 8 | 7432 | 487 | >sp\|P26641-2\|EF1G_HUMAN Isoform 2 of Elongation factor 1-gamma OS=Homo sapiens OX=9606 GN=EEF1G |
| 130 | 57.53 | 4 | 4 | 27443 | 272 | >sp\|P35232\|PHB_HUMAN Prohibitin OS=Homo sapiens OX=9606 GN=PHB PE=1 SV=1 |
| 131 | 57.27 | 7 | 7 | 24592 | 357 | >sp\|P07355-2\|ANXA2_HUMAN Isoform 2 of Annexin A2 OS=Homo sapiens OX=9606 GN=ANXA2 |
| 132 | 57.01 | 7 | 7 | 34806 | 220 | >sp\|P37802-2\|TAGL2_HUMAN Isoform 2 of Transgelin-2 OS=Homo sapiens OX=9606 GN=TAGLN2 |
| 133 | 56.18 | 6 | 6 | 21332 | 257 | >sp\|P62917\|RL8_HUMAN 60S ribosomal protein L8 OS=Homo sapiens OX=9606 GN=RPL8 PE=1 SV=2 |
| 134 | 56.11 | 6 | 6 | 20411 | 533 | >sp\|O43175\|SERA_HUMAN D-3-phosphoglycerate dehydrogenase OS=Homo sapiens OX=9606 GN=PHGDH PE=1 SV=4 |
| 135 | 56.09 | 8 | 8 | 11111 | 653 | >sp\|Q96AE4-2\|FUBP1_HUMAN Isoform 2 of Far upstream element-binding protein 1 OS=Homo sapiens OX=9606 GN=FUBP1 |
| 136 | 55.84 | 7 | 7 | 4017 | 346 | >sp\|P04083\|ANXA1_HUMAN Annexin A1 OS=Homo sapiens OX=9606 GN=ANXA1 PE=1 SV=2 |
| 137 | 55.35 | 5 | 5 | 4124 | 1062 | >sp\|Q86VP6-2\|CAND1_HUMAN Isoform 2 of Cullin-associated NEDD8-dissociated protein 1 OS=Homo sapiens OX=9606 GN=CAND1 |
| 138 | 55.28 | 9 | 9 | 19840 | 1304 | >sp\|O75533\|SF3B1_HUMAN Splicing factor 3B subunit 1 OS=Homo sapiens OX=9606 GN=SF3B1 PE=1 SV=3 |
| 139 | 54.21 | 7 | 7 | 26541 | 620 | >sp\|Q03252\|LMNB2_HUMAN Lamin-B2 OS=Homo sapiens OX=9606 GN=LMNB2 PE=1 SV=4 |
| 140 | 53.59 | 6 | 6 | 500 | 504 | >sp\|Q9BZZ5-2\|API5_HUMAN Isoform 2 of Apoptosis inhibitor 5 OS=Homo sapiens OX=9606 GN=API5 |
| 141 | 53.57 | 7 | 7 | 40872 | 765 | >sp\|P11387\|TOP1_HUMAN DNA topoisomerase 1 OS=Homo sapiens OX=9606 GN=TOP1 PE=1 SV=2 |
| 142 | 53.48 | 5 | 4 | 12190 | 221 | >sp\|P16402\|H13_HUMAN Histone H1.3 OS=Homo sapiens OX=9606 GN=HIST1H1D PE=1 SV=2 |
| 143 | 53.46 | 6 | 5 | 13900 | 210 | >sp\|P09211\|GSTP1_HUMAN Glutathione S-transferase P OS=Homo sapiens OX=9606 GN=GSTP1 PE=1 SV=2 |
| 144 | 53.33 | 7 | 7 | 28589 | 488 | >sp\|Q15084-5\|PDIA6_HUMAN Isoform 5 of Protein disulfide-isomerase A6 OS=Homo sapiens OX=9606 GN=PDIA6 |
| 145 | 53.21 | 8 | 7 | 12624 | 415 | >sp\|P52597\|HNRPF_HUMAN Heterogeneous nuclear ribonucleoprotein F OS=Homo sapiens OX=9606 GN=HNRNPF PE=1 SV=3 |
| 146 | 53.01 | 9 | 9 | 15619 | 459 | >sp\|Q02790\|FKBP4_HUMAN Peptidyl-prolyl cis-trans isomerase FKBP4 OS=Homo sapiens OX=9606 GN=FKBP4 PE=1 SV=3 |
| 147 | 52.97 | 7 | 6 | 42249 | 433 | >sp\|P35998\|PRS7_HUMAN 26S proteasome regulatory subunit 7 OS=Homo sapiens OX=9606 GN=PSMC2 PE=1 SV=3 |
| 148 | 52.74 | 5 | 5 | 35309 | 427 | >sp\|P24752\|THIL_HUMAN Acetyl-CoA acetyltransferase; mitochondrial OS=Homo sapiens OX=9606 GN=ACAT1 PE=1 SV=1 |
| 149 | 52.48 | 10 | 5 | 36287 | 248 | >sp\|Q07955\|SRSF1_HUMAN Serine/arginine-rich splicing factor 1 OS=Homo sapiens OX=9606 GN=SRSF1 PE=1 SV=2 |
| 150 | 52.40 | 10 | 9 | 626 | 586 | >sp\|Q9NVI7-2\|ATD3A_HUMAN Isoform 2 of ATPase family AAA domain-containing protein 3A OS=Homo sapiens OX=9606 GN=ATAD3A |
| 151 | 51.92 | 7 | 7 | 1462 | 2039 | >sp\|Q14008-3\|CKAP5_HUMAN Isoform 3 of Cytoskeleton-associated protein 5 OS=Homo sapiens OX=9606 GN=CKAP5 |
| 152 | 51.81 | 10 | 8 | 11200 | 466 | >sp\|Q13283\|G3BP1_HUMAN Ras GTPase-activating protein-binding protein 1 OS=Homo sapiens OX=9606 GN=G3BP1 PE=1 SV=1 |
| 153 | 51.21 | 6 | 6 | 17443 | 516 | >sp\|Q9Y262-2\|EIF3L_HUMAN Isoform 2 of Eukaryotic translation initiation factor 3 subunit L OS=Homo sapiens OX=9606 GN=EIF3L |
| 154 | 51.18 | 10 | 7 | 13614 | 963 | >sp\|P33176\|KINH_HUMAN Kinesin-1 heavy chain OS=Homo sapiens OX=9606 GN=KIF5B PE=1 SV=1 |
| 155 | 51.02 | 6 | 6 | 21802 | 188 | >sp\|Q07020\|RL18_HUMAN 60S ribosomal protein L18 OS=Homo sapiens OX=9606 GN=RPL18 PE=1 SV=2 |
| 156 | 50.82 | 6 | 6 | 4409 | 277 | >sp\|P16152\|CBR1_HUMAN Carbonyl reductase [NADPH] 1 OS=Homo sapiens OX=9606 GN=CBR1 PE=1 SV=3 |
| 157 | 50.65 | 6 | 6 | 21936 | 505 | >sp\|Q9Y3I0\|RTCB_HUMAN tRNA-splicing ligase RtcB homolog OS=Homo sapiens OX=9606 GN=RTCB PE=1 SV=1 |

| | | | | | | |
|---|---|---|---|---|---|---|
| 158 | 50.17 | 8 | 4 | 1226 | 166 | >sp\|P23528\|COF1_HUMAN Cofilin-1 OS=Homo sapiens OX=9606 GN=CFL1 PE=1 SV=3 |
| 159 | 50.14 | 9 | 7 | 16626 | 289 | >sp\|Q15181\|IPYR_HUMAN Inorganic pyrophosphatase OS=Homo sapiens OX=9606 GN=PPA1 PE=1 SV=2 |
| 160 | 50.05 | 9 | 9 | 4021 | 641 | >sp\|P08133-2\|ANXA6_HUMAN Isoform 2 of Annexin A6 OS=Homo sapiens OX=9606 GN=ANXA6 |
| 161 | 49.94 | 9 | 9 | 31097 | 331 | >sp\|Q9Y266\|NUDC_HUMAN Nuclear migration protein nudC OS=Homo sapiens OX=9606 GN=NUDC PE=1 SV=1 |
| 162 | 49.92 | 4 | 4 | 39025 | 193 | >sp\|Q99426-2\|TBCB_HUMAN Isoform 2 of Tubulin-folding cofactor B OS=Homo sapiens OX=9606 GN=TBCB |
| 163 | 49.78 | 6 | 5 | 26655 | 1985 | >sp\|P35580-5\|MYH10_HUMAN Isoform 5 of Myosin-10 OS=Homo sapiens OX=9606 GN=MYH10 |
| 164 | 49.61 | 7 | 5 | 6433 | 102 | >sp\|P61604\|CH10_HUMAN 10 kDa heat shock protein; mitochondrial OS=Homo sapiens OX=9606 GN=HSPE1 PE=1 SV=2 |
| 165 | 48.86 | 8 | 8 | 34873 | 1262 | >sp\|P41252\|SYIC_HUMAN Isoleucine--tRNA ligase; cytoplasmic OS=Homo sapiens OX=9606 GN=IARS PE=1 SV=2 |
| 166 | 48.76 | 6 | 6 | 34468 | 589 | >sp\|P30153\|2AAA_HUMAN Serine/threonine-protein phosphatase 2A 65 kDa regulatory subunit A alpha isoform OS=Homo sapiens OX=9606 |
| 167 | 48.57 | 6 | 6 | 28625 | 299 | >sp\|Q99623\|PHB2_HUMAN Prohibitin-2 OS=Homo sapiens OX=9606 GN=PHB2 PE=1 SV=2 |
| 168 | 48.57 | 6 | 6 | 4739 | 475 | >sp\|Q01518\|CAP1_HUMAN Adenylyl cyclase-associated protein 1 OS=Homo sapiens OX=9606 GN=CAP1 PE=1 SV=5 |
| 169 | 47.41 | 11 | 10 | 34627 | 1512 | >sp\|P07814\|SYEP_HUMAN Bifunctional glutamate/proline--tRNA ligase OS=Homo sapiens OX=9606 GN=EPRS PE=1 SV=5 |
| 170 | 47.40 | 6 | 6 | 26001 | 338 | >sp\|P40926\|MDHM_HUMAN Malate dehydrogenase; mitochondrial OS=Homo sapiens OX=9606 GN=MDH2 PE=1 SV=3 |
| 171 | 47.34 | 7 | 6 | 14631 | 577 | >sp\|Q9NZI8\|IF2B1_HUMAN Insulin-like growth factor 2 mRNA-binding protein 1 OS=Homo sapiens OX=9606 GN=IGF2BP1 PE=1 SV=2 |
| 172 | 47.32 | 5 | 5 | 36192 | 221 | >sp\|Q13242\|SRSF9_HUMAN Serine/arginine-rich splicing factor 9 OS=Homo sapiens OX=9606 GN=SRSF9 PE=1 SV=1 |
| 173 | 47.09 | 8 | 6 | 14930 | 184 | >sp\|P63241-2\|IF5A1_HUMAN Isoform 2 of Eukaryotic translation initiation factor 5A-1 OS=Homo sapiens OX=9606 GN=EIF5A |
| 174 | 47.05 | 9 | 8 | 21287 | 156 | >sp\|P62750\|RL23A_HUMAN 60S ribosomal protein L23a OS=Homo sapiens OX=9606 GN=RPL23A PE=1 SV=1 |
| 175 | 46.86 | 7 | 7 | 22920 | 1197 | >sp\|O95347\|SMC2_HUMAN Structural maintenance of chromosomes protein 2 OS=Homo sapiens OX=9606 GN=SMC2 PE=1 SV=2 |
| 176 | 46.84 | 5 | 4 | 35782 | 756 | >sp\|P26639-2\|SYTC_HUMAN Isoform 2 of Threonine--tRNA ligase; cytoplasmic OS=Homo sapiens OX=9606 GN=TARS |
| 177 | 46.77 | 7 | 7 | 35064 | 509 | >sp\|P50991-2\|TCPD_HUMAN Isoform 2 of T-complex protein 1 subunit delta OS=Homo sapiens OX=9606 GN=CCT4 |
| 178 | 46.58 | 7 | 6 | 14985 | 529 | >sp\|P52292\|IMA1_HUMAN Importin subunit alpha-1 OS=Homo sapiens OX=9606 GN=KPNA2 PE=1 SV=1 |
| 179 | 46.33 | 7 | 6 | 11386 | 126 | >sp\|Q99880\|H2B1L_HUMAN Histone H2B type 1-L OS=Homo sapiens OX=9606 GN=HIST1H2BL PE=1 SV=3 |
| 180 | 46.27 | 5 | 4 | 12514 | 636 | >sp\|O43390-2\|HNRPR_HUMAN Isoform 2 of Heterogeneous nuclear ribonucleoprotein R OS=Homo sapiens OX=9606 GN=HNRNPR |
| 181 | 46.22 | 8 | 8 | 8085 | 2871 | >sp\|P15924\|DESP_HUMAN Desmoplakin OS=Homo sapiens OX=9606 GN=DSP PE=1 SV=3 |
| 182 | 46.10 | 6 | 6 | 1175 | 283 | >sp\|Q15417-3\|CNN3_HUMAN Isoform 3 of Calponin-3 OS=Homo sapiens OX=9606 GN=CNN3 |
| 183 | 45.49 | 5 | 5 | 10934 | 873 | >sp\|P55884-2\|EIF3B_HUMAN Isoform 2 of Eukaryotic translation initiation factor 3 subunit B OS=Homo sapiens OX=9606 GN=EIF3B |
| 184 | 45.16 | 9 | 7 | 31862 | 267 | >sp\|P22392-2\|NDKB_HUMAN Isoform 3 of Nucleoside diphosphate kinase B OS=Homo sapiens OX=9606 GN=NME2 |
| 185 | 45.01 | 8 | 8 | 22334 | 592 | >sp\|P31939\|PUR9_HUMAN Bifunctional purine biosynthesis protein PURH OS=Homo sapiens OX=9606 GN=ATIC PE=1 SV=3 |
| 186 | 45.00 | 7 | 7 | 32551 | 732 | >sp\|Q08J23-2\|NSUN2_HUMAN Isoform 2 of tRNA (cytosine(34)-C(5))-methyltransferase OS=Homo sapiens OX=9606 GN=NSUN2 |
| 187 | 44.89 | 5 | 5 | 19880 | 456 | >sp\|Q9Y265\|RUVB1_HUMAN RuvB-like 1 OS=Homo sapiens OX=9606 GN=RUVBL1 PE=1 SV=1 |
| 188 | 44.61 | 5 | 5 | 26244 | 711 | >sp\|P49959-3\|MRE11_HUMAN Isoform 3 of Double-strand break repair protein MRE11 OS=Homo sapiens OX=9606 GN=MRE11 |
| 189 | 44.32 | 10 | 10 | 11605 | 876 | >sp\|Q14974\|IMB1_HUMAN Importin subunit beta-1 OS=Homo sapiens OX=9606 GN=KPNB1 PE=1 SV=2 |
| 190 | 44.13 | 5 | 5 | 10792 | 600 | >sp\|Q01844-6\|EWS_HUMAN Isoform 6 of RNA-binding protein EWS OS=Homo sapiens OX=9606 GN=EWSR1 |
| 191 | 44.05 | 7 | 6 | 11247 | 452 | >sp\|P49411\|EFTU_HUMAN Elongation factor Tu; mitochondrial OS=Homo sapiens OX=9606 GN=TUFM PE=1 SV=2 |
| 192 | 43.77 | 8 | 7 | 38916 | 337 | >sp\|P37837\|TALDO_HUMAN Transaldolase OS=Homo sapiens OX=9606 GN=TALDO1 PE=1 SV=2 |
| 193 | 43.00 | 6 | 6 | 10904 | 290 | >sp\|P30084\|ECHM_HUMAN Enoyl-CoA hydratase; mitochondrial OS=Homo sapiens OX=9606 GN=ECHS1 PE=1 SV=4 |
| 194 | 42.88 | 5 | 5 | 2781 | 213 | >sp\|P48047\|ATPO_HUMAN ATP synthase subunit O; mitochondrial OS=Homo sapiens OX=9606 GN=ATP5O PE=1 SV=1 |
| 195 | 42.72 | 6 | 6 | 30919 | 439 | >sp\|P04181\|OAT_HUMAN Ornithine aminotransferase; mitochondrial OS=Homo sapiens OX=9606 GN=OAT PE=1 SV=1 |
| 196 | 42.62 | 9 | 9 | 19307 | 152 | >sp\|P62269\|RS18_HUMAN 40S ribosomal protein S18 OS=Homo sapiens OX=9606 GN=RPS18 PE=1 SV=3 |
| 197 | 42.51 | 5 | 4 | 21316 | 148 | >sp\|P46776\|RL27A_HUMAN 60S ribosomal protein L27a OS=Homo sapiens OX=9606 GN=RPL27A PE=1 SV=2 |
| 198 | 42.49 | 5 | 4 | 24904 | 249 | >sp\|P39687\|AN32A_HUMAN Acidic leucine-rich nuclear phosphoprotein 32 family member A OS=Homo sapiens OX=9606 GN=ANP32A PE=1 SV= |
| 199 | 42.47 | 3 | 3 | 18182 | 925 | >sp\|E9PAV3-2\|NACAM_HUMAN Isoform skNAC-2 of Nascent polypeptide-associated complex subunit alpha; muscle-specific form OS=Homo |
| 200 | 42.46 | 5 | 5 | 21314 | 196 | >sp\|P84098\|RL19_HUMAN 60S ribosomal protein L19 OS=Homo sapiens OX=9606 GN=RPL19 PE=1 SV=1 |
| 201 | 42.14 | 3 | 3 | 16304 | 572 | >sp\|P23588-2\|IF4B_HUMAN Isoform 2 of Eukaryotic translation initiation factor 4B OS=Homo sapiens OX=9606 GN=EIF4B |
| 202 | 41.61 | 4 | 4 | 25995 | 543 | >sp\|P33993-3\|MCM7_HUMAN Isoform 3 of DNA replication licensing factor MCM7 OS=Homo sapiens OX=9606 GN=MCM7 |
| 203 | 41.47 | 7 | 7 | 6405 | 297 | >sp\|P06493\|CDK1_HUMAN Cyclin-dependent kinase 1 OS=Homo sapiens OX=9606 GN=CDK1 PE=1 SV=3 |
| 204 | 40.64 | 4 | 4 | 16997 | 284 | >sp\|P13804-2\|ETFA_HUMAN Isoform 2 of Electron transfer flavoprotein subunit alpha; mitochondrial OS=Homo sapiens OX=9606 GN=ETF |
| 205 | 40.59 | 3 | 3 | 32839 | 396 | >sp\|Q9NTK5\|OLA1_HUMAN Obg-like ATPase 1 OS=Homo sapiens OX=9606 GN=OLA1 PE=1 SV=2 |
| 206 | 40.53 | 8 | 7 | 21386 | 288 | >sp\|Q02878\|RL6_HUMAN 60S ribosomal protein L6 OS=Homo sapiens OX=9606 GN=RPL6 PE=1 SV=3 |
| 207 | 40.47 | 4 | 4 | 36772 | 732 | >sp\|P13010\|XRCC5_HUMAN X-ray repair cross-complementing protein 5 OS=Homo sapiens OX=9606 GN=XRCC5 PE=1 SV=3 |
| 208 | 40.45 | 4 | 3 | 34830 | 445 | >sp\|Q9BVA1\|TBB2B_HUMAN Tubulin beta-2B chain OS=Homo sapiens OX=9606 GN=TUBB2B PE=1 SV=1 |
| 209 | 40.35 | 7 | 7 | 34612 | 298 | >sp\|P05141\|ADT2_HUMAN ADP/ATP translocase 2 OS=Homo sapiens OX=9606 GN=SLC25A5 PE=1 SV=7 |
| 210 | 40.32 | 7 | 6 | 27891 | 508 | >sp\|P07237\|PDIA1_HUMAN Protein disulfide-isomerase OS=Homo sapiens OX=9606 GN=P4HB PE=1 SV=3 |
| 211 | 40.15 | 3 | 3 | 18702 | 386 | >sp\|Q99733-2\|NP1L4_HUMAN Isoform 2 of Nucleosome assembly protein 1-like 4 OS=Homo sapiens OX=9606 GN=NAP1L4 |
| 212 | 40.14 | 7 | 6 | 11534 | 315 | >sp\|P05198\|IF2A_HUMAN Eukaryotic translation initiation factor 2 subunit 1 OS=Homo sapiens OX=9606 GN=EIF2S1 PE=1 SV=3 |

| | | | | | | |
|---|---|---|---|---|---|---|
| 213 | 40.07 | 5 | 5 | 15783 | 572 | >sp\|Q96I24\|FUBP3_HUMAN Far upstream element-binding protein 3 OS=Homo sapiens OX=9606 GN=FUBP3 PE=1 SV=2 |
| 214 | 39.98 | 11 | 11 | 29263 | 2335 | >sp\|Q6P2Q9\|PRP8_HUMAN Pre-mRNA-processing-splicing factor 8 OS=Homo sapiens OX=9606 GN=PRPF8 PE=1 SV=2 |
| 215 | 39.97 | 6 | 6 | 36094 | 363 | >sp\|Q9Y3F4-2\|STRAP_HUMAN Isoform 2 of Serine-threonine kinase receptor-associated protein OS=Homo sapiens OX=9606 GN=STRAP |
| 216 | 39.87 | 4 | 4 | 34462 | 246 | >sp\|P31946\|1433B_HUMAN 14-3-3 protein beta/alpha OS=Homo sapiens OX=9606 GN=YWHAB PE=1 SV=3 |
| 217 | 39.84 | 4 | 4 | 32423 | 594 | >sp\|O00567\|NOP56_HUMAN Nucleolar protein 56 OS=Homo sapiens OX=9606 GN=NOP56 PE=1 SV=4 |
| 218 | 39.64 | 5 | 5 | 20210 | 324 | >sp\|Q9Y617-2\|SERC_HUMAN Isoform 2 of Phosphoserine aminotransferase OS=Homo sapiens OX=9606 GN=PSAT1 |
| 219 | 39.41 | 11 | 7 | 41253 | 915 | >sp\|P55060-4\|XPO2_HUMAN Isoform 4 of Exportin-2 OS=Homo sapiens OX=9606 GN=CSE1L |
| 220 | 39.31 | 6 | 6 | 36164 | 2155 | >sp\|Q01082-3\|SPTB2_HUMAN Isoform 2 of Spectrin beta chain; non-erythrocytic 1 OS=Homo sapiens OX=9606 GN=SPTBN1 |
| 221 | 39.28 | 4 | 4 | 21694 | 305 | >sp\|Q13151\|ROA0_HUMAN Heterogeneous nuclear ribonucleoprotein A0 OS=Homo sapiens OX=9606 GN=HNRNPA0 PE=1 SV=1 |
| 222 | 39.15 | 6 | 4 | 19310 | 249 | >sp\|P62753\|RS6_HUMAN 40S ribosomal protein S6 OS=Homo sapiens OX=9606 GN=RPS6 PE=1 SV=1 |
| 223 | 39.10 | 4 | 4 | 35745 | 992 | >sp\|P49588-2\|SYAC_HUMAN Isoform 2 of Alanine--tRNA ligase; cytoplasmic OS=Homo sapiens OX=9606 GN=AARS |
| 224 | 38.93 | 4 | 4 | 28847 | 158 | >sp\|P24666\|PPAC_HUMAN Low molecular weight phosphotyrosine protein phosphatase OS=Homo sapiens OX=9606 GN=ACP1 PE=1 SV=3 |
| 225 | 38.85 | 5 | 5 | 26028 | 757 | >sp\|Q16891-4\|MIC60_HUMAN Isoform 4 of MICOS complex subunit MIC60 OS=Homo sapiens OX=9606 GN=IMMT |
| 226 | 38.82 | 4 | 3 | 19305 | 208 | >sp\|P62241\|RS8_HUMAN 40S ribosomal protein S8 OS=Homo sapiens OX=9606 GN=RPS8 PE=1 SV=2 |
| 227 | 38.76 | 6 | 6 | 19239 | 165 | >sp\|P46783\|RS10_HUMAN 40S ribosomal protein S10 OS=Homo sapiens OX=9606 GN=RPS10 PE=1 SV=1 |
| 228 | 38.61 | 3 | 3 | 7704 | 341 | >sp\|P35659-2\|DEK_HUMAN Isoform 2 of Protein DEK OS=Homo sapiens OX=9606 GN=DEK |
| 229 | 38.56 | 3 | 3 | 20681 | 305 | >sp\|P40938-2\|RFC3_HUMAN Isoform 2 of Replication factor C subunit 3 OS=Homo sapiens OX=9606 GN=RFC3 |
| 230 | 38.48 | 5 | 5 | 10545 | 533 | >sp\|O15371-3\|EIF3D_HUMAN Isoform 3 of Eukaryotic translation initiation factor 3 subunit D OS=Homo sapiens OX=9606 GN=EIF3D |
| 231 | 37.98 | 5 | 5 | 25642 | 591 | >sp\|P17812\|PYRG1_HUMAN CTP synthase 1 OS=Homo sapiens OX=9606 GN=CTPS1 PE=1 SV=2 |
| 232 | 37.95 | 6 | 5 | 38830 | 1412 | >sp\|Q13428-8\|TCOF_HUMAN Isoform 8 of Treacle protein OS=Homo sapiens OX=9606 GN=TCOF1 |
| 233 | 37.65 | 6 | 5 | 4954 | 694 | >sp\|Q14444-2\|CAPR1_HUMAN Isoform 2 of Caprin-1 OS=Homo sapiens OX=9606 GN=CAPRIN1 |
| 234 | 37.56 | 5 | 5 | 29289 | 248 | >sp\|O14818\|PSA7_HUMAN Proteasome subunit alpha type-7 OS=Homo sapiens OX=9606 GN=PSMA7 PE=1 SV=1 |
| 235 | 37.29 | 6 | 6 | 19311 | 194 | >sp\|P46781\|RS9_HUMAN 40S ribosomal protein S9 OS=Homo sapiens OX=9606 GN=RPS9 PE=1 SV=3 |
| 236 | 37.16 | 4 | 4 | 22648 | 265 | >sp\|P61289-3\|PSME3_HUMAN Isoform 3 of Proteasome activator complex subunit 3 OS=Homo sapiens OX=9606 GN=PSME3 |
| 237 | 37.11 | 7 | 7 | 10547 | 914 | >sp\|B5ME19\|EIFCL_HUMAN Eukaryotic translation initiation factor 3 subunit C-like protein OS=Homo sapiens OX=9606 GN=EIF3CL PE=3 |
| 238 | 36.89 | 3 | 3 | 38613 | 444 | >sp\|P07437\|TBB5_HUMAN Tubulin beta chain OS=Homo sapiens OX=9606 GN=TUBB PE=1 SV=2 |
| 239 | 36.75 | 6 | 5 | 34655 | 588 | >sp\|P54136-2\|SYRC_HUMAN Isoform Monomeric of Arginine--tRNA ligase; cytoplasmic OS=Homo sapiens OX=9606 GN=RARS |
| 240 | 36.57 | 6 | 5 | 32958 | 268 | >sp\|Q15691\|MARE1_HUMAN Microtubule-associated protein RP/EB family member 1 OS=Homo sapiens OX=9606 GN=MAPRE1 PE=1 SV=3 |
| 241 | 36.57 | 4 | 4 | 22857 | 418 | >sp\|P50454\|SERPH_HUMAN Serpin H1 OS=Homo sapiens OX=9606 GN=SERPINH1 PE=1 SV=2 |
| 242 | 36.27 | 7 | 5 | 13299 | 1329 | >sp\|Q86UP2-4\|KTN1_HUMAN Isoform 4 of Kinectin OS=Homo sapiens OX=9606 GN=KTN1 |
| 243 | 36.21 | 6 | 6 | 33358 | 853 | >sp\|P25205-2\|MCM3_HUMAN Isoform 2 of DNA replication licensing factor MCM3 OS=Homo sapiens OX=9606 GN=MCM3 |
| 244 | 36.14 | 5 | 5 | 19265 | 607 | >sp\|P04843\|RPN1_HUMAN Dolichyl-diphosphooligosaccharide--protein glycosyltransferase subunit 1 OS=Homo sapiens OX=9606 GN=RPN1 |
| 245 | 36.06 | 6 | 5 | 6909 | 438 | >sp\|Q99615-2\|DNJC7_HUMAN Isoform 2 of DnaJ homolog subfamily C member 7 OS=Homo sapiens OX=9606 GN=DNAJC7 |
| 246 | 35.97 | 4 | 4 | 28433 | 366 | >sp\|Q15366-2\|PCBP2_HUMAN Isoform 2 of Poly(rC)-binding protein 2 OS=Homo sapiens OX=9606 GN=PCBP2 |
| 247 | 35.86 | 5 | 3 | 28453 | 125 | >sp\|O14737\|PDCD5_HUMAN Programmed cell death protein 5 OS=Homo sapiens OX=9606 GN=PDCD5 PE=1 SV=3 |
| 248 | 35.74 | 8 | 8 | 22232 | 263 | >sp\|P62701\|RS4X_HUMAN 40S ribosomal protein S4; X isoform OS=Homo sapiens OX=9606 GN=RPS4X PE=1 SV=2 |
| 249 | 35.57 | 6 | 6 | 13122 | 408 | >sp\|P05455\|LA_HUMAN Lupus La protein OS=Homo sapiens OX=9606 GN=SSB PE=1 SV=2 |
| 250 | 35.50 | 5 | 5 | 9133 | 185 | >sp\|Q9HB71-3\|CYBP_HUMAN Isoform 3 of Calcyclin-binding protein OS=Homo sapiens OX=9606 GN=CACYBP |
| 251 | 35.32 | 7 | 6 | 18777 | 294 | >sp\|P06748\|NPM_HUMAN Nucleophosmin OS=Homo sapiens OX=9606 GN=NPM1 PE=1 SV=2 |
| 252 | 35.27 | 4 | 4 | 2841 | 269 | >sp\|P35613-2\|BASI_HUMAN Isoform 2 of Basigin OS=Homo sapiens OX=9606 GN=BSG |
| 253 | 35.21 | 4 | 4 | 11102 | 493 | >sp\|Q16658\|FSCN1_HUMAN Fascin OS=Homo sapiens OX=9606 GN=FSCN1 PE=1 SV=3 |
| 254 | 35.11 | 6 | 4 | 12365 | 130 | >sp\|Q6FI13\|H2A2A_HUMAN Histone H2A type 2-A OS=Homo sapiens OX=9606 GN=HIST2H2AA3 PE=1 SV=3 |
| 255 | 35.01 | 5 | 4 | 12561 | 346 | >sp\|P31942\|HNRH3_HUMAN Heterogeneous nuclear ribonucleoprotein H3 OS=Homo sapiens OX=9606 GN=HNRNPH3 PE=1 SV=2 |
| 256 | 34.98 | 7 | 7 | 21306 | 204 | >sp\|P61313\|RL15_HUMAN 60S ribosomal protein L15 OS=Homo sapiens OX=9606 GN=RPL15 PE=1 SV=2 |
| 257 | 34.98 | 4 | 4 | 3384 | 134 | >sp\|P07741-2\|APT_HUMAN Isoform 2 of Adenine phosphoribosyltransferase OS=Homo sapiens OX=9606 GN=APRT |
| 258 | 34.76 | 5 | 5 | 29328 | 216 | >sp\|P23284\|PPIB_HUMAN Peptidyl-prolyl cis-trans isomerase B OS=Homo sapiens OX=9606 GN=PPIB PE=1 SV=2 |
| 259 | 34.61 | 5 | 3 | 11014 | 224 | >sp\|Q00688\|FKBP3_HUMAN Peptidyl-prolyl cis-trans isomerase FKBP3 OS=Homo sapiens OX=9606 GN=FKBP3 PE=1 SV=1 |
| 260 | 34.47 | 4 | 3 | 19133 | 214 | >sp\|P27635\|RL10_HUMAN 60S ribosomal protein L10 OS=Homo sapiens OX=9606 GN=RPL10 PE=1 SV=4 |
| 261 | 34.39 | 5 | 4 | 4295 | 627 | >sp\|P27824-2\|CALX_HUMAN Isoform 2 of Calnexin OS=Homo sapiens OX=9606 GN=CANX |
| 262 | 34.37 | 5 | 4 | 22726 | 432 | >sp\|P22234-2\|PUR6_HUMAN Isoform 2 of Multifunctional protein ADE2 OS=Homo sapiens OX=9606 GN=PAICS |
| 263 | 34.34 | 6 | 6 | 19251 | 151 | >sp\|P62277\|RS13_HUMAN 40S ribosomal protein S13 OS=Homo sapiens OX=9606 GN=RPS13 PE=1 SV=2 |
| 264 | 34.26 | 6 | 5 | 21378 | 165 | >sp\|P30050\|RL12_HUMAN 60S ribosomal protein L12 OS=Homo sapiens OX=9606 GN=RPL12 PE=1 SV=1 |
| 265 | 34.11 | 5 | 3 | 21526 | 192 | >sp\|P32969\|RL9_HUMAN 60S ribosomal protein L9 OS=Homo sapiens OX=9606 GN=RPL9 PE=1 SV=1 |
| 266 | 34.09 | 5 | 5 | 6960 | 715 | >sp\|Q9NR30-2\|DDX21_HUMAN Isoform 2 of Nucleolar RNA helicase 2 OS=Homo sapiens OX=9606 GN=DDX21 |
| 267 | 34.07 | 6 | 6 | 39068 | 172 | >sp\|P13693\|TCTP_HUMAN Translationally-controlled tumor protein OS=Homo sapiens OX=9606 GN=TPT1 PE=1 SV=1 |
| 268 | 33.85 | 6 | 5 | 22345 | 216 | >sp\|P62826\|RAN_HUMAN GTP-binding nuclear protein Ran OS=Homo sapiens OX=9606 GN=RAN PE=1 SV=3 |
| 269 | 33.77 | 7 | 6 | 10546 | 357 | >sp\|O00303\|EIF3F_HUMAN Eukaryotic translation initiation factor 3 subunit F OS=Homo sapiens OX=9606 GN=EIF3F PE=1 SV=1 |
| 270 | 33.68 | 3 | 3 | 7107 | 695 | >sp\|Q16643-3\|DREB_HUMAN Isoform 3 of Drebrin OS=Homo sapiens OX=9606 GN=DBN1 |
| 271 | 33.61 | 4 | 4 | 15590 | 400 | >sp\|P50395-2\|GDIB_HUMAN Isoform 2 of Rab GDP dissociation inhibitor beta OS=Homo sapiens OX=9606 GN=GDI2 |
| 272 | 33.47 | 4 | 4 | 2541 | 303 | >sp\|P46109\|CRKL_HUMAN Crk-like protein OS=Homo sapiens OX=9606 GN=CRKL PE=1 SV=1 |
| 273 | 33.43 | 6 | 6 | 17292 | 320 | >sp\|O75821\|EIF3G_HUMAN Eukaryotic translation initiation factor 3 subunit G OS=Homo sapiens OX=9606 GN=EIF3G PE=1 SV=2 |

| | | | | | | |
|---|---|---|---|---|---|---|
| 274 | 33.19 | 7 | 6 | 236 | 320 | >sp\|P08758\|ANXA5_HUMAN Annexin A5 OS=Homo sapiens OX=9606 GN=ANXA5 PE=1 SV=2 |
| 275 | 33.08 | 5 | 5 | 32152 | 845 | >sp\|P46087-4\|NOP2_HUMAN Isoform 4 of Probable 28S rRNA (cytosine(4447)-C(5))-methyltransferase OS=Homo sapiens OX=9606 GN=NOP2 |
| 276 | 32.78 | 7 | 6 | 9361 | 282 | >sp\|P10768\|ESTD_HUMAN S-formylglutathione hydrolase OS=Homo sapiens OX=9606 GN=ESD PE=1 SV=2 |
| 277 | 32.77 | 4 | 4 | 7514 | 148 | >sp\|O60869\|EDF1_HUMAN Endothelial differentiation-related factor 1 OS=Homo sapiens OX=9606 GN=EDF1 PE=1 SV=1 |
| 278 | 32.74 | 5 | 5 | 21288 | 140 | >sp\|P62829\|RL23_HUMAN 60S ribosomal protein L23 OS=Homo sapiens OX=9606 GN=RPL23 PE=1 SV=1 |
| 279 | 32.59 | 6 | 6 | 6110 | 923 | >sp\|Q8N163-2\|CCAR2_HUMAN Isoform 2 of Cell cycle and apoptosis regulator protein 2 OS=Homo sapiens OX=9606 GN=CCAR2 |
| 280 | 32.52 | 6 | 6 | 15591 | 204 | >sp\|P52565\|GDIR1_HUMAN Rho GDP-dissociation inhibitor 1 OS=Homo sapiens OX=9606 GN=ARHGDIA PE=1 SV=3 |
| 281 | 32.51 | 4 | 4 | 19125 | 193 | >sp\|P08134\|RHOC_HUMAN Rho-related GTP-binding protein RhoC OS=Homo sapiens OX=9606 GN=RHOC PE=1 SV=1 |
| 282 | 32.45 | 4 | 4 | 41837 | 368 | >sp\|P78406\|RAE1L_HUMAN mRNA export factor OS=Homo sapiens OX=9606 GN=RAE1 PE=1 SV=1 |
| 283 | 32.42 | 4 | 3 | 16075 | 628 | >sp\|Q07866-6\|KLC1_HUMAN Isoform N of Kinesin light chain 1 OS=Homo sapiens OX=9606 GN=KLC1 |
| 284 | 32.29 | 5 | 5 | 14669 | 4365 | >sp\|Q7Z6Z7-3\|HUWE1_HUMAN Isoform 3 of E3 ubiquitin-protein ligase HUWE1 OS=Homo sapiens OX=9606 GN=HUWE1 |
| 285 | 32.17 | 4 | 4 | 21312 | 228 | >sp\|P18621-3\|RL17_HUMAN Isoform 3 of 60S ribosomal protein L17 OS=Homo sapiens OX=9606 GN=RPL17 |
| 286 | 32.04 | 5 | 4 | 18951 | 587 | >sp\|P46060\|RAGP1_HUMAN Ran GTPase-activating protein 1 OS=Homo sapiens OX=9606 GN=RANGAP1 PE=1 SV=1 |
| 287 | 32.01 | 4 | 4 | 9191 | 369 | >sp\|P50502\|F10A1_HUMAN Hsc70-interacting protein OS=Homo sapiens OX=9606 GN=ST13 PE=1 SV=2 |
| 288 | 31.79 | 9 | 9 | 21309 | 203 | >sp\|P40429\|RL13A_HUMAN 60S ribosomal protein L13a OS=Homo sapiens OX=9606 GN=RPL13A PE=1 SV=2 |
| 289 | 31.77 | 5 | 3 | 33005 | 352 | >sp\|P40925-3\|MDHC_HUMAN Isoform 3 of Malate dehydrogenase; cytoplasmic OS=Homo sapiens OX=9606 GN=MDH1 |
| 290 | 31.66 | 5 | 4 | 32688 | 372 | >sp\|Q9UNZ2-5\|NSF1C_HUMAN Isoform 3 of NSFL1 cofactor p47 OS=Homo sapiens OX=9606 GN=NSFL1C |
| 291 | 31.56 | 8 | 8 | 7600 | 795 | >sp\|O43143\|DHX15_HUMAN Pre-mRNA-splicing factor ATP-dependent RNA helicase DHX15 OS=Homo sapiens OX=9606 GN=DHX15 PE=1 SV=2 |
| 292 | 31.21 | 3 | 3 | 33566 | 583 | >sp\|Q8N1G4\|LRC47_HUMAN Leucine-rich repeat-containing protein 47 OS=Homo sapiens OX=9606 GN=LRRC47 PE=1 SV=1 |
| 293 | 31.08 | 4 | 3 | 21879 | 217 | >sp\|P62906\|RL10A_HUMAN 60S ribosomal protein L10a OS=Homo sapiens OX=9606 GN=RPL10A PE=1 SV=2 |
| 294 | 31.07 | 4 | 4 | 12359 | 594 | >sp\|P49915-2\|GUAA_HUMAN Isoform 2 of GMP synthase [glutamine-hydrolyzing] OS=Homo sapiens OX=9606 GN=GMPS |
| 295 | 31.06 | 5 | 5 | 19764 | 210 | >sp\|P82979\|SARNP_HUMAN SAP domain-containing ribonucleoprotein OS=Homo sapiens OX=9606 GN=SARNP PE=1 SV=3 |
| 296 | 31.03 | 4 | 4 | 16835 | 2896 | >sp\|P46013-2\|KI67_HUMAN Isoform Short of Proliferation marker protein Ki-67 OS=Homo sapiens OX=9606 GN=MKI67 |
| 297 | 30.89 | 5 | 4 | 20444 | 313 | >sp\|O43765\|SGTA_HUMAN Small glutamine-rich tetratricopeptide repeat-containing protein alpha OS=Homo sapiens OX=9606 GN=SGTA PE |
| 298 | 30.87 | 4 | 4 | 41065 | 229 | >sp\|O43399-7\|TPD54_HUMAN Isoform 7 of Tumor protein D54 OS=Homo sapiens OX=9606 GN=TPD52L2 |
| 299 | 30.86 | 3 | 3 | 34203 | 246 | >sp\|Q04917\|1433F_HUMAN 14-3-3 protein eta OS=Homo sapiens OX=9606 GN=YWHAH PE=1 SV=4 |
| 300 | 30.81 | 4 | 4 | 34719 | 1176 | >sp\|Q9P2J5\|SYLC_HUMAN Leucine--tRNA ligase; cytoplasmic OS=Homo sapiens OX=9606 GN=LARS PE=1 SV=2 |
| 301 | 30.68 | 4 | 4 | 2630 | 327 | >sp\|Q9UJU6-5\|DBNL_HUMAN Isoform 5 of Drebrin-like protein OS=Homo sapiens OX=9606 GN=DBNL |
| 302 | 30.57 | 6 | 4 | 21474 | 266 | >sp\|P62424\|RL7A_HUMAN 60S ribosomal protein L7a OS=Homo sapiens OX=9606 GN=RPL7A PE=1 SV=2 |
| 303 | 30.49 | 4 | 4 | 25053 | 1074 | >sp\|Q96KR1\|ZFR_HUMAN Zinc finger RNA-binding protein OS=Homo sapiens OX=9606 GN=ZFR PE=1 SV=2 |
| 304 | 30.43 | 10 | 9 | 23083 | 1233 | >sp\|Q14683\|SMC1A_HUMAN Structural maintenance of chromosomes protein 1A OS=Homo sapiens OX=9606 GN=SMC1A PE=1 SV=2 |
| 305 | 30.43 | 7 | 7 | 11626 | 1115 | >sp\|O00410-3\|IPO5_HUMAN Isoform 3 of Importin-5 OS=Homo sapiens OX=9606 GN=IPO5 |
| 306 | 30.35 | 9 | 9 | 15628 | 2633 | >sp\|O75369-8\|FLNB_HUMAN Isoform 8 of Filamin-B OS=Homo sapiens OX=9606 GN=FLNB |
| 307 | 30.12 | 5 | 4 | 21432 | 105 | >sp\|Q9Y3U8\|RL36_HUMAN 60S ribosomal protein L36 OS=Homo sapiens OX=9606 GN=RPL36 PE=1 SV=3 |
| 308 | 30.11 | 3 | 3 | 4314 | 412 | >sp\|P07339\|CATD_HUMAN Cathepsin D OS=Homo sapiens OX=9606 GN=CTSD PE=1 SV=1 |
| 309 | 30.02 | 4 | 3 | 38242 | 108 | >sp\|O75347\|TBCA_HUMAN Tubulin-specific chaperone A OS=Homo sapiens OX=9606 GN=TBCA PE=1 SV=3 |
| 310 | 29.91 | 4 | 4 | 29011 | 378 | >sp\|Q9UNM6-2\|PSD13_HUMAN Isoform 2 of 26S proteasome non-ATPase regulatory subunit 13 OS=Homo sapiens OX=9606 GN=PSMD13 |
| 311 | 29.53 | 2 | 2 | 21801 | 215 | >sp\|P50914\|RL14_HUMAN 60S ribosomal protein L14 OS=Homo sapiens OX=9606 GN=RPL14 PE=1 SV=4 |
| 312 | 29.49 | 3 | 3 | 37673 | 483 | >sp\|P54578-3\|UBP14_HUMAN Isoform 3 of Ubiquitin carboxyl-terminal hydrolase 14 OS=Homo sapiens OX=9606 GN=USP14 |
| 313 | 29.48 | 4 | 3 | 38831 | 127 | >sp\|P53999\|TCP4_HUMAN Activated RNA polymerase II transcriptional coactivator p15 OS=Homo sapiens OX=9606 GN=SUB1 PE=1 SV=3 |
| 314 | 29.47 | 6 | 5 | 19304 | 204 | >sp\|P46782\|RS5_HUMAN 40S ribosomal protein S5 OS=Homo sapiens OX=9606 GN=RPS5 PE=1 SV=4 |
| 315 | 29.23 | 4 | 4 | 19615 | 895 | >sp\|Q13435\|SF3B2_HUMAN Splicing factor 3B subunit 2 OS=Homo sapiens OX=9606 GN=SF3B2 PE=1 SV=2 |
| 316 | 29.17 | 5 | 4 | 20644 | 280 | >sp\|Q15293-2\|RCN1_HUMAN Isoform 2 of Reticulocalbin-1 OS=Homo sapiens OX=9606 GN=RCN1 |
| 317 | 29.08 | 3 | 3 | 9631 | 467 | >sp\|P07954-2\|FUMH_HUMAN Isoform Cytoplasmic of Fumarate hydratase; mitochondrial OS=Homo sapiens OX=9606 GN=FH |
| 318 | 29.06 | 3 | 3 | 22727 | 234 | >sp\|P25787\|PSA2_HUMAN Proteasome subunit alpha type-2 OS=Homo sapiens OX=9606 GN=PSMA2 PE=1 SV=2 |
| 319 | 29.05 | 3 | 3 | 29308 | 330 | >sp\|P62136\|PP1A_HUMAN Serine/threonine-protein phosphatase PP1-alpha catalytic subunit OS=Homo sapiens OX=9606 GN=PPP1CA PE=1 S |
| 320 | 28.95 | 4 | 4 | 11406 | 2079 | >sp\|P51610-4\|HCFC1_HUMAN Isoform 4 of Host cell factor 1 OS=Homo sapiens OX=9606 GN=HCFC1 |
| 321 | 28.88 | 7 | 7 | 1436 | 466 | >sp\|O75390\|CISY_HUMAN Citrate synthase; mitochondrial OS=Homo sapiens OX=9606 GN=CS PE=1 SV=2 |
| 322 | 28.76 | 10 | 9 | 26929 | 4684 | >sp\|Q15149\|PLEC_HUMAN Plectin OS=Homo sapiens OX=9606 GN=PLEC PE=1 SV=3 |
| 323 | 28.51 | 5 | 5 | 23403 | 1288 | >sp\|Q9NTJ3\|SMC4_HUMAN Structural maintenance of chromosomes protein 4 OS=Homo sapiens OX=9606 GN=SMC4 PE=1 SV=2 |
| 324 | 28.50 | 9 | 7 | 12446 | 215 | >sp\|P09429\|HMGB1_HUMAN High mobility group protein B1 OS=Homo sapiens OX=9606 GN=HMGB1 PE=1 SV=3 |
| 325 | 28.41 | 5 | 5 | 16999 | 346 | >sp\|P38117-2\|ETFB_HUMAN Isoform 2 of Electron transfer flavoprotein subunit beta OS=Homo sapiens OX=9606 GN=ETFB |
| 326 | 28.40 | 6 | 5 | 16376 | 514 | >sp\|P12268\|IMDH2_HUMAN Inosine-5'-monophosphate dehydrogenase 2 OS=Homo sapiens OX=9606 GN=IMPDH2 PE=1 SV=2 |
| 327 | 28.38 | 9 | 7 | 40982 | 2363 | >sp\|P12270\|TPR_HUMAN Nucleoprotein TPR OS=Homo sapiens OX=9606 GN=TPR PE=1 SV=3 |
| 328 | 28.23 | 6 | 6 | 21290 | 136 | >sp\|P61353\|RL27_HUMAN 60S ribosomal protein L27 OS=Homo sapiens OX=9606 GN=RPL27 PE=1 SV=2 |
| 329 | 28.02 | 3 | 3 | 22906 | 107 | >sp\|Q9GZT3-2\|SLIRP_HUMAN Isoform 2 of SRA stem-loop-interacting RNA-binding protein; mitochondrial OS=Homo sapiens OX=9606 GN=S |
| 330 | 27.95 | 5 | 5 | 24714 | 475 | >sp\|P49419-4\|AL7A1_HUMAN Isoform 4 of Alpha-aminoadipic semialdehyde dehydrogenase OS=Homo sapiens OX=9606 GN=ALDH7A1 |

| | | | | | | |
|---|---|---|---|---|---|---|
| 331 | 27.85 | 2 | 2 | 27422 | 154 | >sp\|Q9UHV9\|PFD2_HUMAN Prefoldin subunit 2 OS=Homo sapiens OX=9606 GN=PFDN2 PE=1 SV=1 |
| 332 | 27.83 | 4 | 4 | 8033 | 819 | >sp\|Q86XP3-2\|DDX42_HUMAN Isoform 2 of ATP-dependent RNA helicase DDX42 OS=Homo sapiens OX=9606 GN=DDX42 |
| 333 | 27.77 | 4 | 4 | 42254 | 248 | >sp\|P25788-2\|PSA3_HUMAN Isoform 2 of Proteasome subunit alpha type-3 OS=Homo sapiens OX=9606 GN=PSMA3 |
| 334 | 27.76 | 2 | 2 | 21300 | 128 | >sp\|P35268\|RL22_HUMAN 60S ribosomal protein L22 OS=Homo sapiens OX=9606 GN=RPL22 PE=1 SV=2 |
| 335 | 27.58 | 3 | 3 | 22581 | 409 | >sp\|P54727\|RD23B_HUMAN UV excision repair protein RAD23 homolog B OS=Homo sapiens OX=9606 GN=RAD23B PE=1 SV=1 |
| 336 | 27.49 | 3 | 3 | 13707 | 261 | >sp\|Q99714\|HCD2_HUMAN 3-hydroxyacyl-CoA dehydrogenase type-2 OS=Homo sapiens OX=9606 GN=HSD17B10 PE=1 SV=3 |
| 337 | 27.39 | 5 | 5 | 28812 | 224 | >sp\|P30041\|PRDX6_HUMAN Peroxiredoxin-6 OS=Homo sapiens OX=9606 GN=PRDX6 PE=1 SV=3 |
| 338 | 27.34 | 3 | 3 | 2845 | 173 | >sp\|P80723-2\|BASP1_HUMAN Isoform 2 of Brain acid soluble protein 1 OS=Homo sapiens OX=9606 GN=BASP1 |
| 339 | 27.28 | 4 | 4 | 2503 | 169 | >sp\|P13073\|COX41_HUMAN Cytochrome c oxidase subunit 4 isoform 1; mitochondrial OS=Homo sapiens OX=9606 GN=COX4I1 PE=1 SV=1 |
| 340 | 27.10 | 4 | 4 | 41927 | 480 | >sp\|P31930\|QCR1_HUMAN Cytochrome b-c1 complex subunit 1; mitochondrial OS=Homo sapiens OX=9606 GN=UQCRC1 PE=1 SV=3 |
| 341 | 26.99 | 5 | 4 | 38851 | 139 | >sp\|Q15185-4\|TEBP_HUMAN Isoform 4 of Prostaglandin E synthase 3 OS=Homo sapiens OX=9606 GN=PTGES3 |
| 342 | 26.91 | 4 | 4 | 8150 | 420 | >sp\|O60832-2\|DKC1_HUMAN Isoform 3 of H/ACA ribonucleoprotein complex subunit 4 OS=Homo sapiens OX=9606 GN=DKC1 |
| 343 | 26.84 | 5 | 5 | 16432 | 400 | >sp\|P08727\|K1C19_HUMAN Keratin; type I cytoskeletal 19 OS=Homo sapiens OX=9606 GN=KRT19 PE=1 SV=4 |
| 344 | 26.83 | 5 | 5 | 37646 | 2136 | >sp\|O75643\|U520_HUMAN U5 small nuclear ribonucleoprotein 200 kDa helicase OS=Homo sapiens OX=9606 GN=SNRNP200 PE=1 SV=2 |
| 345 | 26.81 | 4 | 3 | 24762 | 220 | >sp\|Q9BTT0-3\|AN32E_HUMAN Isoform 3 of Acidic leucine-rich nuclear phosphoprotein 32 family member E OS=Homo sapiens OX=9606 GN= |
| 346 | 26.58 | 7 | 6 | 22230 | 259 | >sp\|P23396-2\|RS3_HUMAN Isoform 2 of 40S ribosomal protein S3 OS=Homo sapiens OX=9606 GN=RPS3 |
| 347 | 26.48 | 7 | 6 | 712 | 303 | >sp\|Q9UNE7\|CHIP_HUMAN E3 ubiquitin-protein ligase CHIP OS=Homo sapiens OX=9606 GN=STUB1 PE=1 SV=2 |
| 348 | 26.31 | 6 | 4 | 41727 | 283 | >sp\|P45880-2\|VDAC2_HUMAN Isoform 2 of Voltage-dependent anion-selective channel protein 2 OS=Homo sapiens OX=9606 GN=VDAC2 |
| 349 | 26.31 | 3 | 3 | 34664 | 366 | >sp\|P52788\|SPSY_HUMAN Spermine synthase OS=Homo sapiens OX=9606 GN=SMS PE=1 SV=2 |
| 350 | 26.30 | 3 | 3 | 13269 | 432 | >sp\|O95232\|LC7L3_HUMAN Luc7-like protein 3 OS=Homo sapiens OX=9606 GN=LUC7L3 PE=1 SV=2 |
| 351 | 26.22 | 5 | 5 | 34564 | 376 | >sp\|P61163\|ACTZ_HUMAN Alpha-centractin OS=Homo sapiens OX=9606 GN=ACTR1A PE=1 SV=1 |
| 352 | 26.13 | 2 | 2 | 12196 | 181 | >sp\|Q13442\|HAP28_HUMAN 28 kDa heat- and acid-stable phosphoprotein OS=Homo sapiens OX=9606 GN=PDAP1 PE=1 SV=1 |
| 353 | 25.90 | 3 | 2 | 21364 | 115 | >sp\|P05387\|RLA2_HUMAN 60S acidic ribosomal protein P2 OS=Homo sapiens OX=9606 GN=RPLP2 PE=1 SV=1 |
| 354 | 25.87 | 2 | 2 | 2165 | 352 | >sp\|Q9BT78-2\|CSN4_HUMAN Isoform 2 of COP9 signalosome complex subunit 4 OS=Homo sapiens OX=9606 GN=COPS4 |
| 355 | 25.84 | 3 | 3 | 31975 | 239 | >sp\|Q9UKD2\|MRT4_HUMAN mRNA turnover protein 4 homolog OS=Homo sapiens OX=9606 GN=MRTO4 PE=1 SV=2 |
| 356 | 25.81 | 5 | 5 | 30993 | 793 | >sp\|P54886-2\|P5CS_HUMAN Isoform Short of Delta-1-pyrroline-5-carboxylate synthase OS=Homo sapiens OX=9606 GN=ALDH18A1 |
| 357 | 25.76 | 4 | 3 | 8893 | 478 | >sp\|Q16630-3\|CPSF6_HUMAN Isoform 3 of Cleavage and polyadenylation specificity factor subunit 6 OS=Homo sapiens OX=9606 GN=CPSF |
| 358 | 25.75 | 6 | 6 | 36188 | 2752 | >sp\|Q9UQ35\|SRRM2_HUMAN Serine/arginine repetitive matrix protein 2 OS=Homo sapiens OX=9606 GN=SRRM2 PE=1 SV=2 |
| 359 | 25.57 | 2 | 2 | 36776 | 58 | >sp\|Q96IX5\|USMG5_HUMAN Up-regulated during skeletal muscle growth protein 5 OS=Homo sapiens OX=9606 GN=USMG5 PE=1 SV=1 |
| 360 | 25.39 | 2 | 2 | 20127 | 321 | >sp\|O14828-2\|SCAM3_HUMAN Isoform 2 of Secretory carrier-associated membrane protein 3 OS=Homo sapiens OX=9606 GN=SCAMP3 |
| 361 | 25.33 | 3 | 3 | 15080 | 504 | >sp\|Q9NPH2-3\|INO1_HUMAN Isoform 3 of Inositol-3-phosphate synthase 1 OS=Homo sapiens OX=9606 GN=ISYNA1 |
| 362 | 25.31 | 2 | 2 | 14105 | 1293 | >sp\|Q6Y7W6-5\|GGYF2_HUMAN Isoform 4 of GRB10-interacting GYF protein 2 OS=Homo sapiens OX=9606 GN=GIGYF2 |
| 363 | 25.30 | 3 | 3 | 3129 | 313 | >sp\|P51572-2\|BAP31_HUMAN Isoform 2 of B-cell receptor-associated protein 31 OS=Homo sapiens OX=9606 GN=BCAP31 |
| 364 | 25.21 | 4 | 4 | 24897 | 488 | >sp\|P28838-2\|AMPL_HUMAN Isoform 2 of Cytosol aminopeptidase OS=Homo sapiens OX=9606 GN=LAP3 |
| 365 | 25.18 | 3 | 3 | 40291 | 243 | >sp\|Q9BVC6\|TM109_HUMAN Transmembrane protein 109 OS=Homo sapiens OX=9606 GN=TMEM109 PE=1 SV=1 |
| 366 | 25.11 | 2 | 2 | 3069 | 298 | >sp\|P36542\|ATPG_HUMAN ATP synthase subunit gamma; mitochondrial OS=Homo sapiens OX=9606 GN=ATP5F1C PE=1 SV=1 |
| 367 | 25.06 | 4 | 4 | 41389 | 617 | >sp\|P38606\|VATA_HUMAN V-type proton ATPase catalytic subunit A OS=Homo sapiens OX=9606 GN=ATP6V1A PE=1 SV=2 |
| 368 | 24.96 | 5 | 4 | 13927 | 371 | >sp\|O75367-3\|H2AY_HUMAN Isoform 3 of Core histone macro-H2A.1 OS=Homo sapiens OX=9606 GN=H2AFY |
| 369 | 24.94 | 5 | 5 | 37032 | 1235 | >sp\|Q00341-2\|VIGLN_HUMAN Isoform 2 of Vigilin OS=Homo sapiens OX=9606 GN=HDLBP |
| 370 | 24.90 | 2 | 2 | 21697 | 332 | >sp\|Q99729\|ROAA_HUMAN Heterogeneous nuclear ribonucleoprotein A/B OS=Homo sapiens OX=9606 GN=HNRNPAB PE=1 SV=2 |
| 371 | 24.80 | 4 | 3 | 22922 | 1217 | >sp\|Q9UQE7\|SMC3_HUMAN Structural maintenance of chromosomes protein 3 OS=Homo sapiens OX=9606 GN=SMC3 PE=1 SV=2 |
| 372 | 24.54 | 4 | 4 | 10165 | 533 | >sp\|Q8N8S7-3\|ENAH_HUMAN Isoform 3 of Protein enabled homolog OS=Homo sapiens OX=9606 GN=ENAH |
| 373 | 24.52 | 3 | 3 | 12637 | 218 | >sp\|P00492\|HPRT_HUMAN Hypoxanthine-guanine phosphoribosyltransferase OS=Homo sapiens OX=9606 GN=HPRT1 PE=1 SV=2 |
| 374 | 24.46 | 4 | 4 | 12626 | 456 | >sp\|P14866-2\|HNRPL_HUMAN Isoform 2 of Heterogeneous nuclear ribonucleoprotein L OS=Homo sapiens OX=9606 GN=HNRNPL |
| 375 | 24.44 | 2 | 2 | 5553 | 202 | >sp\|Q9Y3Y2-4\|CHTOP_HUMAN Isoform 3 of Chromatin target of PRMT1 protein OS=Homo sapiens OX=9606 GN=CHTOP |
| 376 | 24.44 | 3 | 3 | 22228 | 59 | >sp\|P62861\|RS30_HUMAN 40S ribosomal protein S30 OS=Homo sapiens OX=9606 GN=FAU PE=1 SV=1 |
| 377 | 24.43 | 3 | 3 | 21356 | 135 | >sp\|P62910\|RL32_HUMAN 60S ribosomal protein L32 OS=Homo sapiens OX=9606 GN=RPL32 PE=1 SV=2 |
| 378 | 24.38 | 3 | 3 | 12592 | 205 | >sp\|P04792\|HSPB1_HUMAN Heat shock protein beta-1 OS=Homo sapiens OX=9606 GN=HSPB1 PE=1 SV=2 |
| 379 | 24.38 | 3 | 3 | 29061 | 255 | >sp\|P67775-2\|PP2AA_HUMAN Isoform 2 of Serine/threonine-protein phosphatase 2A catalytic subunit alpha isoform OS=Homo sapiens O |
| 380 | 24.37 | 2 | 2 | 11533 | 144 | >sp\|P47813\|IF1AX_HUMAN Eukaryotic translation initiation factor 1A; X-chromosomal OS=Homo sapiens OX=9606 GN=EIF1AX PE=1 SV=2 |
| 381 | 24.30 | 5 | 4 | 29256 | 140 | >sp\|P07737\|PROF1_HUMAN Profilin-1 OS=Homo sapiens OX=9606 GN=PFN1 PE=1 SV=2 |
| 382 | 24.24 | 2 | 2 | 15022 | 237 | >sp\|P54105\|ICLN_HUMAN Methylosome subunit pICln OS=Homo sapiens OX=9606 GN=CLNS1A PE=1 SV=1 |
| 383 | 24.18 | 2 | 2 | 21623 | 282 | >sp\|Q9Y3B9\|RRP15_HUMAN RRP15-like protein OS=Homo sapiens OX=9606 GN=RRP15 PE=1 SV=2 |
| 384 | 24.08 | 5 | 4 | 3616 | 1091 | >sp\|P53396-2\|ACLY_HUMAN Isoform 2 of ATP-citrate synthase OS=Homo sapiens OX=9606 GN=ACLY |

| | | | | | | |
|---|---|---|---|---|---|---|
| 385 | 24.06 | 4 | 3 | 19565 | 616 | >sp\|P31040-2\|SDHA_HUMAN Isoform 2 of Succinate dehydrogenase [ubiquinone] flavoprotein subunit; mitochondrial OS=Homo sapiens O |
| 386 | 24.02 | 5 | 5 | 34764 | 528 | >sp\|P54577\|SYYC_HUMAN Tyrosine--tRNA ligase; cytoplasmic OS=Homo sapiens OX=9606 GN=YARS PE=1 SV=4 |
| 387 | 23.99 | 4 | 4 | 33834 | 587 | >sp\|P09960-4\|LKHA4_HUMAN Isoform 4 of Leukotriene A-4 hydrolase OS=Homo sapiens OX=9606 GN=LTA4H |
| 388 | 23.98 | 2 | 2 | 1459 | 602 | >sp\|Q07065\|CKAP4_HUMAN Cytoskeleton-associated protein 4 OS=Homo sapiens OX=9606 GN=CKAP4 PE=1 SV=2 |
| 389 | 23.97 | 3 | 3 | 26884 | 228 | >sp\|P22061-2\|PIMT_HUMAN Isoform 2 of Protein-L-isoaspartate(D-aspartate) O-methyltransferase OS=Homo sapiens OX=9606 GN=PCMT1 |
| 390 | 23.96 | 3 | 3 | 10935 | 325 | >sp\|Q13347\|EIF3I_HUMAN Eukaryotic translation initiation factor 3 subunit I OS=Homo sapiens OX=9606 GN=EIF3I PE=1 SV=1 |
| 391 | 23.89 | 5 | 5 | 18198 | 169 | >sp\|Q04760-2\|LGUL_HUMAN Isoform 2 of Lactoylglutathione lyase OS=Homo sapiens OX=9606 GN=GLO1 |
| 392 | 23.73 | 5 | 5 | 21329 | 403 | >sp\|P39023\|RL3_HUMAN 60S ribosomal protein L3 OS=Homo sapiens OX=9606 GN=RPL3 PE=1 SV=2 |
| 393 | 23.73 | 3 | 3 | 16994 | 1114 | >sp\|Q9BSJ8-2\|ESYT1_HUMAN Isoform 2 of Extended synaptotagmin-1 OS=Homo sapiens OX=9606 GN=ESYT1 |
| 394 | 23.71 | 3 | 3 | 36556 | 796 | >sp\|Q96QK1\|VPS35_HUMAN Vacuolar protein sorting-associated protein 35 OS=Homo sapiens OX=9606 GN=VPS35 PE=1 SV=2 |
| 395 | 23.53 | 5 | 5 | 21921 | 125 | >sp\|P62851\|RS25_HUMAN 40S ribosomal protein S25 OS=Homo sapiens OX=9606 GN=RPS25 PE=1 SV=1 |
| 396 | 23.37 | 4 | 4 | 16422 | 218 | >sp\|Q9H910-3\|JUPI2_HUMAN Isoform 3 of Jupiter microtubule associated homolog 2 OS=Homo sapiens OX=9606 GN=JPT2 |
| 397 | 23.33 | 4 | 4 | 37907 | 1104 | >sp\|Q14157-5\|UBP2L_HUMAN Isoform 5 of Ubiquitin-associated protein 2-like OS=Homo sapiens OX=9606 GN=UBAP2L |
| 398 | 23.04 | 4 | 4 | 22391 | 364 | >sp\|Q9BWF3\|RBM4_HUMAN RNA-binding protein 4 OS=Homo sapiens OX=9606 GN=RBM4 PE=1 SV=1 |
| 399 | 23.03 | 2 | 2 | 6451 | 303 | >sp\|P11802\|CDK4_HUMAN Cyclin-dependent kinase 4 OS=Homo sapiens OX=9606 GN=CDK4 PE=1 SV=2 |
| 400 | 23.01 | 7 | 6 | 14646 | 1606 | >sp\|Q04637-9\|IF4G1_HUMAN Isoform 9 of Eukaryotic translation initiation factor 4 gamma 1 OS=Homo sapiens OX=9606 GN=EIF4G1 |
| 401 | 22.98 | 3 | 3 | 19293 | 326 | >sp\|Q9NQG5\|RPR1B_HUMAN Regulation of nuclear pre-mRNA domain-containing protein 1B OS=Homo sapiens OX=9606 GN=RPRD1B PE=1 SV=1 |
| 402 | 22.98 | 4 | 4 | 42093 | 456 | >sp\|P30520\|PURA2_HUMAN Adenylosuccinate synthetase isozyme 2 OS=Homo sapiens OX=9606 GN=ADSS PE=1 SV=3 |
| 403 | 22.95 | 4 | 4 | 41620 | 152 | >sp\|P61088\|UBE2N_HUMAN Ubiquitin-conjugating enzyme E2 N OS=Homo sapiens OX=9606 GN=UBE2N PE=1 SV=1 |
| 404 | 22.84 | 3 | 3 | 10940 | 112 | >sp\|Q15369\|ELOC_HUMAN Elongin-C OS=Homo sapiens OX=9606 GN=ELOC PE=1 SV=1 |
| 405 | 22.81 | 2 | 2 | 4272 | 559 | >sp\|Q13112\|CAF1B_HUMAN Chromatin assembly factor 1 subunit B OS=Homo sapiens OX=9606 GN=CHAF1B PE=1 SV=1 |
| 406 | 22.76 | 4 | 3 | 18064 | 368 | >sp\|P55209-2\|NP1L1_HUMAN Isoform 2 of Nucleosome assembly protein 1-like 1 OS=Homo sapiens OX=9606 GN=NAP1L1 |
| 407 | 22.64 | 4 | 4 | 40218 | 188 | >sp\|P62995-3\|TRA2B_HUMAN Isoform 3 of Transformer-2 protein homolog beta OS=Homo sapiens OX=9606 GN=TRA2B |
| 408 | 22.60 | 2 | 2 | 28430 | 233 | >sp\|Q9BVG4\|PBDC1_HUMAN Protein PBDC1 OS=Homo sapiens OX=9606 GN=PBDC1 PE=1 SV=1 |
| 409 | 22.51 | 2 | 1 | 35209 | 142 | >sp\|Q9NS69\|TOM22_HUMAN Mitochondrial import receptor subunit TOM22 homolog OS=Homo sapiens OX=9606 GN=TOMM22 PE=1 SV=3 |
| 410 | 22.51 | 4 | 4 | 19306 | 132 | >sp\|P25398\|RS12_HUMAN 40S ribosomal protein S12 OS=Homo sapiens OX=9606 GN=RPS12 PE=1 SV=3 |
| 411 | 22.48 | 4 | 4 | 19878 | 166 | >sp\|P08621-3\|RU17_HUMAN Isoform 3 of U1 small nuclear ribonucleoprotein 70 kDa OS=Homo sapiens OX=9606 GN=SNRNP70 |
| 412 | 22.48 | 4 | 3 | 21382 | 123 | >sp\|P42766\|RL35_HUMAN 60S ribosomal protein L35 OS=Homo sapiens OX=9606 GN=RPL35 PE=1 SV=2 |
| 413 | 22.22 | 2 | 2 | 22643 | 377 | >sp\|P55036\|PSMD4_HUMAN 26S proteasome non-ATPase regulatory subunit 4 OS=Homo sapiens OX=9606 GN=PSMD4 PE=1 SV=1 |
| 414 | 22.14 | 3 | 3 | 21377 | 177 | >sp\|P62913-2\|RL11_HUMAN Isoform 2 of 60S ribosomal protein L11 OS=Homo sapiens OX=9606 GN=RPL11 |
| 415 | 22.14 | 7 | 7 | 21289 | 145 | >sp\|P61254\|RL26_HUMAN 60S ribosomal protein L26 OS=Homo sapiens OX=9606 GN=RPL26 PE=1 SV=1 |
| 416 | 22.10 | 6 | 6 | 13911 | 977 | >sp\|P78347-4\|GTF2I_HUMAN Isoform 4 of General transcription factor II-I OS=Homo sapiens OX=9606 GN=GTF2I |
| 417 | 22.09 | 2 | 2 | 7924 | 467 | >sp\|Q9UJW0-3\|DCTN4_HUMAN Isoform 3 of Dynactin subunit 4 OS=Homo sapiens OX=9606 GN=DCTN4 |
| 418 | 21.95 | 2 | 2 | 32646 | 416 | >sp\|P30419-2\|NMT1_HUMAN Isoform Short of Glycylpeptide N-tetradecanoyltransferase 1 OS=Homo sapiens OX=9606 GN=NMT1 |
| 419 | 21.95 | 2 | 2 | 34204 | 247 | >sp\|P61981\|1433G_HUMAN 14-3-3 protein gamma OS=Homo sapiens OX=9606 GN=YWHAG PE=1 SV=2 |
| 420 | 21.93 | 3 | 3 | 23654 | 154 | >sp\|P00441\|SODC_HUMAN Superoxide dismutase [Cu-Zn] OS=Homo sapiens OX=9606 GN=SOD1 PE=1 SV=2 |
| 421 | 21.86 | 3 | 3 | 20527 | 673 | >sp\|Q15637-5\|SF01_HUMAN Isoform 5 of Splicing factor 1 OS=Homo sapiens OX=9606 GN=SF1 |
| 422 | 21.83 | 5 | 4 | 12737 | 224 | >sp\|P54819-5\|KAD2_HUMAN Isoform 5 of Adenylate kinase 2; mitochondrial OS=Homo sapiens OX=9606 GN=AK2 |
| 423 | 21.83 | 3 | 3 | 1424 | 313 | >sp\|Q9UHD1-2\|CHRD1_HUMAN Isoform 2 of Cysteine and histidine-rich domain-containing protein 1 OS=Homo sapiens OX=9606 GN=CHORDC |
| 424 | 21.75 | 2 | 2 | 34390 | 245 | >sp\|P27348\|1433T_HUMAN 14-3-3 protein theta OS=Homo sapiens OX=9606 GN=YWHAQ PE=1 SV=1 |
| 425 | 21.75 | 4 | 3 | 16013 | 740 | >sp\|P51812\|KS6A3_HUMAN Ribosomal protein S6 kinase alpha-3 OS=Homo sapiens OX=9606 GN=RPS6KA3 PE=1 SV=1 |
| 426 | 21.61 | 2 | 2 | 32763 | 440 | >sp\|Q9UKX7-2\|NUP50_HUMAN Isoform 2 of Nuclear pore complex protein Nup50 OS=Homo sapiens OX=9606 GN=NUP50 |
| 427 | 21.55 | 5 | 4 | 9255 | 321 | >sp\|P22087\|FBRL_HUMAN rRNA 2'-O-methyltransferase fibrillarin OS=Homo sapiens OX=9606 GN=FBL PE=1 SV=2 |
| 428 | 21.50 | 2 | 2 | 37977 | 158 | >sp\|P63279\|UBC9_HUMAN SUMO-conjugating enzyme UBC9 OS=Homo sapiens OX=9606 GN=UBE2I PE=1 SV=1 |
| 429 | 21.40 | 4 | 3 | 20876 | 452 | >sp\|P18754-2\|RCC1_HUMAN Isoform 2 of Regulator of chromosome condensation OS=Homo sapiens OX=9606 GN=RCC1 |
| 430 | 21.37 | 2 | 2 | 42050 | 593 | >sp\|Q06124-2\|PTN11_HUMAN Isoform 2 of Tyrosine-protein phosphatase non-receptor type 11 OS=Homo sapiens OX=9606 GN=PTPN11 |
| 431 | 21.30 | 2 | 2 | 21949 | 159 | >sp\|P09234\|RU1C_HUMAN U1 small nuclear ribonucleoprotein C OS=Homo sapiens OX=9606 GN=SNRPC PE=1 SV=1 |
| 432 | 21.24 | 2 | 2 | 6839 | 191 | >sp\|P60953\|CDC42_HUMAN Cell division control protein 42 homolog OS=Homo sapiens OX=9606 GN=CDC42 PE=1 SV=2 |
| 433 | 21.23 | 2 | 2 | 22461 | 239 | >sp\|P28072\|PSB6_HUMAN Proteasome subunit beta type-6 OS=Homo sapiens OX=9606 GN=PSMB6 PE=1 SV=4 |
| 434 | 21.23 | 4 | 4 | 36131 | 108 | >sp\|P62316-2\|SMD2_HUMAN Isoform 2 of Small nuclear ribonucleoprotein Sm D2 OS=Homo sapiens OX=9606 GN=SNRPD2 |
| 435 | 21.18 | 3 | 3 | 21880 | 83 | >sp\|P63220\|RS21_HUMAN 40S ribosomal protein S21 OS=Homo sapiens OX=9606 GN=RPS21 PE=1 SV=1 |
| 436 | 21.18 | 2 | 2 | 21699 | 285 | >sp\|Q99729-3\|ROAA_HUMAN Isoform 3 of Heterogeneous nuclear ribonucleoprotein A/B OS=Homo sapiens OX=9606 GN=HNRNPAB |
| 437 | 21.11 | 3 | 3 | 23973 | 402 | >sp\|O60749-2\|SNX2_HUMAN Isoform 2 of Sorting nexin-2 OS=Homo sapiens OX=9606 GN=SNX2 |
| 438 | 20.99 | 3 | 3 | 12280 | 449 | >sp\|P31943\|HNRH1_HUMAN Heterogeneous nuclear ribonucleoprotein H OS=Homo sapiens OX=9606 GN=HNRNPH1 PE=1 SV=4 |
| 439 | 20.95 | 2 | 2 | 21696 | 356 | >sp\|P51991-2\|ROA3_HUMAN Isoform 2 of Heterogeneous nuclear ribonucleoprotein A3 OS=Homo sapiens OX=9606 GN=HNRNPA3 |

| | | | | | | |
|---|---|---|---|---|---|---|
| 440 | 20.94 | 2 | 2 | 26853 | 125 | >sp\|Q9NRX4\|PHP14_HUMAN 14 kDa phosphohistidine phosphatase OS=Homo sapiens OX=9606 GN=PHPT1 PE=1 SV=1 |
| 441 | 20.87 | 6 | 6 | 21038 | 3224 | >sp\|P49792\|RBP2_HUMAN E3 SUMO-protein ligase RanBP2 OS=Homo sapiens OX=9606 GN=RANBP2 PE=1 SV=2 |
| 442 | 20.86 | 4 | 4 | 23623 | 910 | >sp\|Q7KZF4\|SND1_HUMAN Staphylococcal nuclease domain-containing protein 1 OS=Homo sapiens OX=9606 GN=SND1 PE=1 SV=1 |
| 443 | 20.78 | 2 | 2 | 35086 | 257 | >sp\|Q86V81\|THOC4_HUMAN THO complex subunit 4 OS=Homo sapiens OX=9606 GN=ALYREF PE=1 SV=3 |
| 444 | 20.78 | 2 | 2 | 15951 | 404 | >sp\|Q07666-3\|KHDR1_HUMAN Isoform 3 of KH domain-containing; RNA-binding; signal transduction-associated protein 1 OS=Homo sapie |
| 445 | 20.78 | 3 | 3 | 22729 | 241 | >sp\|P20618\|PSB1_HUMAN Proteasome subunit beta type-1 OS=Homo sapiens OX=9606 GN=PSMB1 PE=1 SV=2 |
| 446 | 20.67 | 3 | 3 | 24230 | 930 | >sp\|P12814-4\|ACTN1_HUMAN Isoform 4 of Alpha-actinin-1 OS=Homo sapiens OX=9606 GN=ACTN1 |
| 447 | 20.61 | 4 | 4 | 31506 | 151 | >sp\|P60660-2\|MYL6_HUMAN Isoform Smooth muscle of Myosin light polypeptide 6 OS=Homo sapiens OX=9606 GN=MYL6 |
| 448 | 20.53 | 2 | 2 | 35210 | 309 | >sp\|Q15785\|TOM34_HUMAN Mitochondrial import receptor subunit TOM34 OS=Homo sapiens OX=9606 GN=TOMM34 PE=1 SV=2 |
| 449 | 20.47 | 3 | 2 | 21531 | 198 | >sp\|P52815\|RM12_HUMAN 39S ribosomal protein L12; mitochondrial OS=Homo sapiens OX=9606 GN=MRPL12 PE=1 SV=2 |
| 450 | 20.41 | 3 | 3 | 16466 | 325 | >sp\|Q96CX2\|KCD12_HUMAN BTB/POZ domain-containing protein KCTD12 OS=Homo sapiens OX=9606 GN=KCTD12 PE=1 SV=1 |
| 451 | 20.31 | 4 | 3 | 41426 | 294 | >sp\|Q9P0L0-2\|VAPA_HUMAN Isoform 2 of Vesicle-associated membrane protein-associated protein A OS=Homo sapiens OX=9606 GN=VAPA |
| 452 | 20.23 | 4 | 4 | 20408 | 371 | >sp\|Q15019-3\|SEPT2_HUMAN Isoform 3 of Septin-2 OS=Homo sapiens OX=9606 GN=SEPT2 |
| 453 | 20.14 | 2 | 2 | 27077 | 195 | >sp\|O00264\|PGRC1_HUMAN Membrane-associated progesterone receptor component 1 OS=Homo sapiens OX=9606 GN=PGRMC1 PE=1 SV=3 |
| 454 | 20.12 | 7 | 7 | 25810 | 1332 | >sp\|Q9BQG0-2\|MBB1A_HUMAN Isoform 2 of Myb-binding protein 1A OS=Homo sapiens OX=9606 GN=MYBBP1A |
| 455 | 20.12 | 3 | 3 | 20838 | 508 | >sp\|Q14498-3\|RBM39_HUMAN Isoform 3 of RNA-binding protein 39 OS=Homo sapiens OX=9606 GN=RBM39 |
| 456 | 20.08 | 3 | 3 | 10959 | 254 | >sp\|P50402\|EMD_HUMAN Emerin OS=Homo sapiens OX=9606 GN=EMD PE=1 SV=1 |
| 457 | 20.05 | 4 | 4 | 37287 | 546 | >sp\|P40222\|TXLNA_HUMAN Alpha-taxilin OS=Homo sapiens OX=9606 GN=TXLNA PE=1 SV=3 |
| 458 | 19.95 | 3 | 3 | 31489 | 866 | >sp\|Q9BXJ9\|NAA15_HUMAN N-alpha-acetyltransferase 15; NatA auxiliary subunit OS=Homo sapiens OX=9606 GN=NAA15 PE=1 SV=1 |
| 459 | 19.94 | 6 | 2 | 37667 | 229 | >sp\|P0CG47\|UBB_HUMAN Polyubiquitin-B OS=Homo sapiens OX=9606 GN=UBB PE=1 SV=1 |
| 460 | 19.91 | 4 | 2 | 19309 | 142 | >sp\|P60866-2\|RS20_HUMAN Isoform 2 of 40S ribosomal protein S20 OS=Homo sapiens OX=9606 GN=RPS20 |
| 461 | 19.91 | 3 | 3 | 26503 | 489 | >sp\|O75439\|MPPB_HUMAN Mitochondrial-processing peptidase subunit beta OS=Homo sapiens OX=9606 GN=PMPCB PE=1 SV=2 |
| 462 | 19.86 | 2 | 2 | 26486 | 221 | >sp\|Q7L9L4-2\|MOB1B_HUMAN Isoform 2 of MOB kinase activator 1B OS=Homo sapiens OX=9606 GN=MOB1B |
| 463 | 19.84 | 3 | 3 | 22227 | 151 | >sp\|P62263\|RS14_HUMAN 40S ribosomal protein S14 OS=Homo sapiens OX=9606 GN=RPS14 PE=1 SV=3 |
| 464 | 19.70 | 4 | 3 | 9589 | 335 | >sp\|O76003\|GLRX3_HUMAN Glutaredoxin-3 OS=Homo sapiens OX=9606 GN=GLRX3 PE=1 SV=2 |
| 465 | 19.69 | 4 | 4 | 19839 | 501 | >sp\|Q12874\|SF3A3_HUMAN Splicing factor 3A subunit 3 OS=Homo sapiens OX=9606 GN=SF3A3 PE=1 SV=1 |
| 466 | 19.69 | 5 | 4 | 17694 | 586 | >sp\|P15311\|EZRI_HUMAN Ezrin OS=Homo sapiens OX=9606 GN=EZR PE=1 SV=4 |
| 467 | 19.65 | 2 | 2 | 29405 | 317 | >sp\|Q15435-2\|PP1R7_HUMAN Isoform 2 of Protein phosphatase 1 regulatory subunit 7 OS=Homo sapiens OX=9606 GN=PPP1R7 |
| 468 | 19.56 | 6 | 4 | 20186 | 439 | >sp\|Q9NVA2-2\|SEP11_HUMAN Isoform 2 of Septin-11 OS=Homo sapiens OX=9606 GN=SEPT11 |
| 469 | 19.56 | 4 | 4 | 1892 | 527 | >sp\|Q9ULV4-3\|COR1C_HUMAN Isoform 3 of Coronin-1C OS=Homo sapiens OX=9606 GN=CORO1C |
| 470 | 19.53 | 3 | 3 | 27196 | 717 | >sp\|Q9H307\|PININ_HUMAN Pinin OS=Homo sapiens OX=9606 GN=PNN PE=1 SV=5 |
| 471 | 19.51 | 4 | 4 | 22539 | 423 | >sp\|O00231-2\|PSD11_HUMAN Isoform 2 of 26S proteasome non-ATPase regulatory subunit 11 OS=Homo sapiens OX=9606 GN=PSMD11 |
| 472 | 19.45 | 3 | 3 | 17440 | 445 | >sp\|P60228\|EIF3E_HUMAN Eukaryotic translation initiation factor 3 subunit E OS=Homo sapiens OX=9606 GN=EIF3E PE=1 SV=1 |
| 473 | 19.28 | 4 | 4 | 37749 | 1086 | >sp\|Q93009-3\|UBP7_HUMAN Isoform 3 of Ubiquitin carboxyl-terminal hydrolase 7 OS=Homo sapiens OX=9606 GN=USP7 |
| 474 | 19.22 | 3 | 3 | 12699 | 181 | >sp\|Q9UK76-2\|JUPI1_HUMAN Isoform 2 of Jupiter microtubule associated homolog 1 OS=Homo sapiens OX=9606 GN=JPT1 |
| 475 | 19.22 | 3 | 3 | 23951 | 513 | >sp\|Q2TAY7\|SMU1_HUMAN WD40 repeat-containing protein SMU1 OS=Homo sapiens OX=9606 GN=SMU1 PE=1 SV=2 |
| 476 | 19.20 | 3 | 3 | 11218 | 164 | >sp\|P33316-2\|DUT_HUMAN Isoform 2 of Deoxyuridine 5'-triphosphate nucleotidohydrolase; mitochondrial OS=Homo sapiens OX=9606 GN= |
| 477 | 19.18 | 2 | 2 | 7017 | 738 | >sp\|O00429-8\|DNM1L_HUMAN Isoform 8 of Dynamin-1-like protein OS=Homo sapiens OX=9606 GN=DNM1L |
| 478 | 19.15 | 2 | 2 | 13831 | 483 | >sp\|P49840\|GSK3A_HUMAN Glycogen synthase kinase-3 alpha OS=Homo sapiens OX=9606 GN=GSK3A PE=1 SV=2 |
| 479 | 19.12 | 2 | 2 | 31739 | 81 | >sp\|Q9GZZ1-2\|NAA50_HUMAN Isoform 2 of N-alpha-acetyltransferase 50 OS=Homo sapiens OX=9606 GN=NAA50 |
| 480 | 19.12 | 2 | 2 | 11677 | 390 | >sp\|Q12905\|ILF2_HUMAN Interleukin enhancer-binding factor 2 OS=Homo sapiens OX=9606 GN=ILF2 PE=1 SV=2 |
| 481 | 19.11 | 3 | 2 | 41614 | 471 | >sp\|P26368-2\|U2AF2_HUMAN Isoform 2 of Splicing factor U2AF 65 kDa subunit OS=Homo sapiens OX=9606 GN=U2AF2 |
| 482 | 19.04 | 3 | 2 | 2886 | 869 | >sp\|Q9NYF8-3\|BCLF1_HUMAN Isoform 3 of Bcl-2-associated transcription factor 1 OS=Homo sapiens OX=9606 GN=BCLAF1 |
| 483 | 19.02 | 3 | 3 | 21301 | 157 | >sp\|P83731\|RL24_HUMAN 60S ribosomal protein L24 OS=Homo sapiens OX=9606 GN=RPL24 PE=1 SV=1 |
| 484 | 18.98 | 2 | 2 | 20243 | 97 | >sp\|P60903\|S10AA_HUMAN Protein S100-A10 OS=Homo sapiens OX=9606 GN=S100A10 PE=1 SV=2 |
| 485 | 18.93 | 2 | 2 | 34835 | 502 | >sp\|O95793-3\|STAU1_HUMAN Isoform 3 of Double-stranded RNA-binding protein Staufen homolog 1 OS=Homo sapiens OX=9606 GN=STAU1 |
| 486 | 18.84 | 2 | 2 | 34157 | 425 | >sp\|P11310-2\|ACADM_HUMAN Isoform 2 of Medium-chain specific acyl-CoA dehydrogenase; mitochondrial OS=Homo sapiens OX=9606 GN=AC |
| 487 | 18.70 | 2 | 2 | 34826 | 451 | >sp\|P68363\|TBA1B_HUMAN Tubulin alpha-1B chain OS=Homo sapiens OX=9606 GN=TUBA1B PE=1 SV=1 |
| 488 | 18.38 | 3 | 3 | 30638 | 271 | >sp\|Q96FW1\|OTUB1_HUMAN Ubiquitin thioesterase OTUB1 OS=Homo sapiens OX=9606 GN=OTUB1 PE=1 SV=2 |
| 489 | 18.32 | 4 | 3 | 4210 | 326 | >sp\|O43684-2\|BUB3_HUMAN Isoform 2 of Mitotic checkpoint protein BUB3 OS=Homo sapiens OX=9606 GN=BUB3 |
| 490 | 18.31 | 3 | 3 | 28549 | 261 | >sp\|P12004\|PCNA_HUMAN Proliferating cell nuclear antigen OS=Homo sapiens OX=9606 GN=PCNA PE=1 SV=1 |
| 491 | 18.25 | 3 | 3 | 18062 | 301 | >sp\|Q9Y314\|NOSIP_HUMAN Nitric oxide synthase-interacting protein OS=Homo sapiens OX=9606 GN=NOSIP PE=1 SV=1 |
| 492 | 18.24 | 2 | 2 | 7147 | 306 | >sp\|Q99848\|EBP2_HUMAN Probable rRNA-processing protein EBP2 OS=Homo sapiens OX=9606 GN=EBNA1BP2 PE=1 SV=2 |
| 493 | 18.13 | 3 | 2 | 15027 | 366 | >sp\|P50213\|IDH3A_HUMAN Isocitrate dehydrogenase [NAD] subunit alpha; mitochondrial OS=Homo sapiens OX=9606 GN=IDH3A PE=1 SV=1 |
| 494 | 18.11 | 7 | 4 | 15672 | 525 | >sp\|P35637-2\|FUS_HUMAN Isoform Short of RNA-binding protein FUS OS=Homo sapiens OX=9606 GN=FUS |
| 495 | 18.08 | 3 | 3 | 21885 | 69 | >sp\|P62857\|RS28_HUMAN 40S ribosomal protein S28 OS=Homo sapiens OX=9606 GN=RPS28 PE=1 SV=1 |

| | | | | | | |
|---|---|---|---|---|---|---|
| 496 | 18.06 | 2 | 2 | 32284 | 128 | >sp\|P55769\|NH2L1_HUMAN NHP2-like protein 1 OS=Homo sapiens OX=9606 GN=SNU13 PE=1 SV=3 |
| 497 | 17.99 | 2 | 2 | 6904 | 331 | >sp\|P31689-2\|DNJA1_HUMAN Isoform 2 of DnaJ homolog subfamily A member 1 OS=Homo sapiens OX=9606 GN=DNAJA1 |
| 498 | 17.94 | 2 | 2 | 21113 | 1118 | >sp\|Q92900-2\|RENT1_HUMAN Isoform 2 of Regulator of nonsense transcripts 1 OS=Homo sapiens OX=9606 GN=UPF1 |
| 499 | 17.89 | 5 | 4 | 37663 | 170 | >sp\|Q13404-7\|UB2V1_HUMAN Isoform 5 of Ubiquitin-conjugating enzyme E2 variant 1 OS=Homo sapiens OX=9606 GN=UBE2V1 |
| 500 | 17.89 | 4 | 4 | 440 | 1015 | >sp\|P16615-4\|AT2A2_HUMAN Isoform 4 of Sarcoplasmic/endoplasmic reticulum calcium ATPase 2 OS=Homo sapiens OX=9606 GN=ATP2A2 |
| 501 | 17.85 | 5 | 5 | 20530 | 793 | >sp\|Q15459\|SF3A1_HUMAN Splicing factor 3A subunit 1 OS=Homo sapiens OX=9606 GN=SF3A1 PE=1 SV=1 |
| 502 | 17.79 | 2 | 2 | 25668 | 201 | >sp\|Q9H0U4\|RAB1B_HUMAN Ras-related protein Rab-1B OS=Homo sapiens OX=9606 GN=RAB1B PE=1 SV=1 |

#### Score distribution

827 peptide matches above score 32, 1106 (17 reverse) matches above 20
Top score for reverse: 46, top score for reverse with > 9 aa's: 24
Charge distribution: z=+1: 0, z=+2: 795, z=+3: 32, z=+4: 0

#### Mass measurement errors

**Precursors**

*Before recalibration:*
Median precursor m/z error (observed minus true): -0.0004 Da, -0.7 ppm
Median precursor accuracy (absolute value of error): 0.0004 Da, 0.8 ppm
Precursors measured too high: 64 Too low: 649

*After recalibration:*
Median precursor m/z error (observed minus true): 0.0000 Da, 0.0 ppm
Median precursor accuracy (absolute value of error): 0.0002 Da, 0.3 ppm
Precursors measured too high: 358 Too low: 351

Off-by-one errors (nominal mass is +1 isotope): 11.1% (88/795)

**Fragments**

*Before recalibration:*
Median fragment m/z error (observed minus true): 0.0930 Da, 141.0 ppm
Median fragment accuracy (absolute value of error): 0.0935 Da, 142.6 ppm
Fragments (within ±0.5 Da) measured too high: 4176 Too low: 142

*After recalibration:*
Median fragment m/z error (observed minus true): 0.0176 Da, 27.3 ppm
Median fragment accuracy (absolute value of error): 0.0323 Da, 50.1 ppm
Fragments (within ±0.5 Da) measured too high: 2574 Too low: 1744

#### Cysteine

Cysteine set to a fixed modification of +57
Completeness: 86.4% (51/59) peptides C[+57], 11.9% (7/59) C[+0], 1.7% (1/59) C[+71] (from gel)
Carbamidomethylation artifacts (H,K,N-terminus[+57/+114]): 0.0% (0/1072) of peptides
DTT artifacts (C[+209]): 5.6% (64/1136) of peptides
Disulfide bridge (unmodified_C[-2]): 33.3% (1/3) of peptides containing at least two C's

#### Nonspecific cleavage

Cleavage sites (C-side): RK
Missed cleavage: 31.8% (269/847) of semitryptic peptides contain an internal K or R not followed by P
Semitryptic peptides (% of tryptic and semitryptic): 5.4% (46/847) ragged-N, 3.7% (31/847) ragged-C
Nontryptic peptides (% of all peptides): 0.0% (0/841)

#### Oxidation

Oxidized methionine (M[+16]): 15.5% (67/433) of peptides containing M[+0] or M[+16]
Side chain loss from oxidized methionine (M[-48]): 0.3% (1/367) of peptides containing M[+0] or M[-48]
Doubly oxidized methionine (M[+32]): 0.7% (2/304) of peptides containing one M[+32] or M[+0]
Oxidized histidine and tryptophan (H[+16], W[+16]): 0.8% (2/255) (2 H, 0 W) of peptides containing H or W
Doubly oxidized tryptophan (W[+32]): 0.0% (0/82) of peptides containing W but not M
Triply oxidized cysteine (C[+48]): 1.1% (1/88) of peptides containing C

#### Chemical modifications

Deamidated asparagine or glutamine: 11.3% (109/966) (82 N, 27 Q) of peptides containing N and/or Q
Amidated aspartic or glutamic acid: 4.7% (57/1212) of peptides containing D and/or E
Pyro-glu N-terminus (Q[-17], E[-18], C[+57][-17]): 10.6% (11/104) (10 -17, 1 -18) of peptides with N-terminal Q, E, or camC
Sodiation: 0.0% (6/1354) of all peptides (4 on E and D)
Carbamylation (N-terminus[+43], R[+43], K[+43]): 1.0% (14/1366) (5 N-term) of peptides
Carbamylated methionine (M[+43]): 1.0% (4/408) of peptides containing M
Formaldehyde (N-terminus[+12], W[+12]): 0.0% (0/1354) of peptides
Acetaldehyde (N-terminus[+26], H[+26], K[+26]): 0.3% (4/1354) (4 N-term) of peptides
N-terminal methylation/dimethylation (N-terminus[+14/+28]): 0.6% (8/1354) (8 +14, 0 +28) of peptides
Peptide (not protein) N-terminal acetylation (N-terminus[+42]): 0.0% (0/1354) of peptides
Cysteine propionamide (C[+71]): 0.0% (0/78) of peptides containing C
Methyl ester (E[+14]): 0.9% (7/819) of peptides containing E
Formylation (S[+28], T[+28]): 0.3% (2/746) of peptides containing S or T

#### Posttranslational modifications

Hydroxyproline: 3.0% (15/497) of peptides containing P
Phosphorylation: 0.1% (1/870) (0 S, 0 T, 1 Y) of peptides containing S, T, or Y
Beta-elimination (S[-18], T[-18]): 0.0% (0/744) peptides containing S or T
Dimethylation (K[+28], R[+28]): 0.9% (11/1228) (11 K, 0 R) of peptides containing K or R
Methylation (K[+14], H[+14], N[+14], R[+14]): 1.1% (14/1281) (1 K) of peptides containing K, H, N, or R

Acetylation (or guanidination or trimethylation) (K[+42]): 0.7% (6/801) of peptides containing K
Protein N-terminal acetylation: 58.3% (7/12) of N-terminal peptides

#### Computational information

Spectrum file: D:\Luci\Moggridge et al data\ch_03Fe...p_1.raw_Preview_181130_173155\objs\ch_03Fe...-100p_1.mgf
Protein database: D:\Luci\PD Databases\Schulz_CP_Homo_sapiens_proteomeUP000005640reveiwed_Download_20180420_proteins_20303.fasta
Presets: Cysteine 57, Lysine 0, Arginine 0, N-terminus 0, C-terminus 0
CID fragmentation
Digestion after RK
First pass is fully tryptic digestion
No wildcard search
Read in 22398 spectrum/charge combinations: z=+1 0, z=+2 14410, z=+3 6857, z=+4 1131
Precursor mass from 756 to 3001 Da
Read 42313 proteins
Scored 354349926 candidate/spectrum/charge combinations

Preview version: v2.13.17
Output folder: D:\Luci\Moggridge et al data\ch_03Fe...p_1.raw_Preview_181130_173155

Click to open **parameters** used for this run.

Note: The Details page reports the rates of modification on eligible peptides, whereas the Summary page reports potential gains in the total number of identifications. Denominators in the percentages may also vary from search to search due to "second-order" effects such as multiply modified peptides and corrections for hits to decoys.

**Summary** **Detail**

www.proteinmetrics.com

PROTEIN METRICS

**Preview™ by PROTEIN METRICS Inc.**

Summary Detail

#### Table of Contents (ch_03Feb2018_SP3-Solvents_acetone-ph8-100p_1.raw) (Schulz_CP_Homo_sapiens_proteomeUP000005640reveiwed_Download_20180420_proteins_20303.fasta)


1. Representative proteins
2. Score distribution
3. Mass measurement errors
4. Cysteine
5. Nonspecific cleavage
6. Oxidation
7. Chemical modifications
8. Posttranslational modifications
9. Computation information


#### Representative proteins

Right click (or option-click) and save link: **peptide and protein identifications** (in csv format). Or, access the **folder**.

| Rank | ProtScore | # Spectra ID'd | # Unique Peptides | Database Sequence # | # aa's | Protein Name |
|---|---|---|---|---|---|---|
| 1 | 282.55 | 44 | 32 | 14875 | 854 | >sp\|P07900-2\|HS90A_HUMAN Isoform 2 of Heat shock protein HSP 90-alpha OS=Homo sapiens OX=9606 GN=HSP90AA1 |
| 2 | 253.52 | 40 | 34 | 10117 | 858 | >sp\|P13639\|EF2_HUMAN Elongation factor 2 OS=Homo sapiens OX=9606 GN=EEF2 PE=1 SV=4 |
| 3 | 251.53 | 33 | 26 | 12650 | 641 | >sp\|P0DMV9\|HS71B_HUMAN Heat shock 70 kDa protein 1B OS=Homo sapiens OX=9606 GN=HSPA1B PE=1 SV=1 |
| 4 | 250.04 | 30 | 27 | 29794 | 1960 | >sp\|P35579\|MYH9_HUMAN Myosin-9 OS=Homo sapiens OX=9606 GN=MYH9 PE=1 SV=4 |
| 5 | 218.69 | 27 | 24 | 41440 | 466 | >sp\|P08670\|VIME_HUMAN Vimentin OS=Homo sapiens OX=9606 GN=VIM PE=1 SV=4 |
| 6 | 199.00 | 28 | 25 | 17426 | 2639 | >sp\|P21333-2\|FLNA_HUMAN Isoform 2 of Filamin-A OS=Homo sapiens OX=9606 GN=FLNA |
| 7 | 198.50 | 23 | 20 | 11486 | 646 | >sp\|P11142\|HSP7C_HUMAN Heat shock cognate 71 kDa protein OS=Homo sapiens OX=9606 GN=HSPA8 PE=1 SV=1 |
| 8 | 186.80 | 20 | 20 | 27671 | 2101 | >sp\|Q14980-2\|NUMA1_HUMAN Isoform 2 of Nuclear mitotic apparatus protein 1 OS=Homo sapiens OX=9606 GN=NUMA1 |
| 9 | 179.78 | 22 | 19 | 5194 | 654 | >sp\|P11021\|BIP_HUMAN Endoplasmic reticulum chaperone BiP OS=Homo sapiens OX=9606 GN=HSPA5 PE=1 SV=2 |
| 10 | 172.68 | 22 | 18 | 10608 | 434 | >sp\|P06733\|ENOA_HUMAN Alpha-enolase OS=Homo sapiens OX=9606 GN=ENO1 PE=1 SV=2 |
| 11 | 170.32 | 22 | 20 | 38833 | 548 | >sp\|P50990\|TCPQ_HUMAN T-complex protein 1 subunit theta OS=Homo sapiens OX=9606 GN=CCT8 PE=1 SV=4 |
| 12 | 169.25 | 17 | 16 | 26540 | 586 | >sp\|P20700\|LMNB1_HUMAN Lamin-B1 OS=Homo sapiens OX=9606 GN=LMNB1 PE=1 SV=2 |
| 13 | 168.51 | 21 | 20 | 38994 | 806 | >sp\|P55072\|TERA_HUMAN Transitional endoplasmic reticulum ATPase OS=Homo sapiens OX=9606 GN=VCP PE=1 SV=4 |
| 14 | 165.64 | 23 | 19 | 34832 | 445 | >sp\|P68371\|TBB4B_HUMAN Tubulin beta-4B chain OS=Homo sapiens OX=9606 GN=TUBB4B PE=1 SV=1 |
| 15 | 163.57 | 28 | 28 | 3268 | 5890 | >sp\|Q09666\|AHNK_HUMAN Neuroblast differentiation-associated protein AHNAK OS=Homo sapiens OX=9606 GN=AHNAK PE=1 SV=2 |
| 16 | 163.24 | 18 | 18 | 35062 | 488 | >sp\|P78371-2\|TCPB_HUMAN Isoform 2 of T-complex protein 1 subunit beta OS=Homo sapiens OX=9606 GN=CCT2 |
| 17 | 154.22 | 19 | 18 | 27856 | 1014 | >sp\|P09874\|PARP1_HUMAN Poly [ADP-ribose] polymerase 1 OS=Homo sapiens OX=9606 GN=PARP1 PE=1 SV=4 |
| 18 | 149.16 | 20 | 19 | 5883 | 573 | >sp\|P10809\|CH60_HUMAN 60 kDa heat shock protein; mitochondrial OS=Homo sapiens OX=9606 GN=HSPD1 PE=1 SV=2 |
| 19 | 148.02 | 24 | 23 | 35928 | 590 | >sp\|P31948-2\|STIP1_HUMAN Isoform 2 of Stress-induced-phosphoprotein 1 OS=Homo sapiens OX=9606 GN=STIP1 |
| 20 | 147.50 | 23 | 21 | 12573 | 691 | >sp\|P52272-2\|HNRPM_HUMAN Isoform 2 of Heterogeneous nuclear ribonucleoprotein M OS=Homo sapiens OX=9606 GN=HNRNPM |
| 21 | 146.41 | 21 | 20 | 17366 | 2511 | >sp\|P49327\|FAS_HUMAN Fatty acid synthase OS=Homo sapiens OX=9606 GN=FASN PE=1 SV=3 |
| 22 | 145.90 | 17 | 15 | 7556 | 662 | >sp\|O00571\|DDX3X_HUMAN ATP-dependent RNA helicase DDX3X OS=Homo sapiens OX=9606 GN=DDX3X PE=1 SV=3 |
| 23 | 143.05 | 15 | 14 | 39061 | 545 | >sp\|P49368\|TCPG_HUMAN T-complex protein 1 subunit gamma OS=Homo sapiens OX=9606 GN=CCT3 PE=1 SV=4 |
| 24 | 141.44 | 19 | 17 | 34358 | 911 | >sp\|O43707\|ACTN4_HUMAN Alpha-actinin-4 OS=Homo sapiens OX=9606 GN=ACTN4 PE=1 SV=2 |
| 25 | 138.26 | 26 | 16 | 34144 | 375 | >sp\|P60709\|ACTB_HUMAN Actin; cytoplasmic 1 OS=Homo sapiens OX=9606 GN=ACTB PE=1 SV=1 |
| 26 | 136.06 | 21 | 19 | 17521 | 803 | >sp\|P14625\|ENPL_HUMAN Endoplasmin OS=Homo sapiens OX=9606 GN=HSP90B1 PE=1 SV=1 |
| 27 | 135.95 | 15 | 14 | 13137 | 334 | >sp\|P07195\|LDHB_HUMAN L-lactate dehydrogenase B chain OS=Homo sapiens OX=9606 GN=LDHB PE=1 SV=2 |
| 28 | 134.58 | 12 | 12 | 199 | 553 | >sp\|P25705\|ATPA_HUMAN ATP synthase subunit alpha; mitochondrial OS=Homo sapiens OX=9606 GN=ATP5F1A PE=1 SV=1 |
| 29 | 134.16 | 15 | 13 | 40876 | 249 | >sp\|P60174-1\|TPIS_HUMAN Isoform 2 of Triosephosphate isomerase OS=Homo sapiens OX=9606 GN=TPI1 |
| 30 | 128.94 | 21 | 20 | 42235 | 4128 | >sp\|P78527\|PRKDC_HUMAN DNA-dependent protein kinase catalytic subunit OS=Homo sapiens OX=9606 GN=PRKDC PE=1 SV=3 |
| 31 | 126.51 | 19 | 16 | 27269 | 630 | >sp\|P13797\|PLST_HUMAN Plastin-3 OS=Homo sapiens OX=9606 GN=PLS3 PE=1 SV=4 |
| 32 | 126.11 | 16 | 11 | 14006 | 103 | >sp\|P62805\|H4_HUMAN Histone H4 OS=Homo sapiens OX=9606 GN=HIST1H4A PE=1 SV=2 |
| 33 | 125.25 | 16 | 13 | 12288 | 293 | >sp\|P07910-2\|HNRPC_HUMAN Isoform C1 of Heterogeneous nuclear ribonucleoproteins C1/C2 OS=Homo sapiens OX=9606 GN=HNRNPC |
| 34 | 124.57 | 16 | 15 | 12627 | 825 | >sp\|Q00839\|HNRPU_HUMAN Heterogeneous nuclear ribonucleoprotein U OS=Homo sapiens OX=9606 GN=HNRNPU PE=1 SV=6 |
| 35 | 123.01 | 18 | 16 | 8068 | 535 | >sp\|P17844-2\|DDX5_HUMAN Isoform 2 of Probable ATP-dependent RNA helicase DDX5 OS=Homo sapiens OX=9606 GN=DDX5 |
| 36 | 122.95 | 16 | 15 | 11112 | 711 | >sp\|Q92945\|FUBP2_HUMAN Far upstream element-binding protein 2 OS=Homo sapiens OX=9606 GN=KHSRP PE=1 SV=4 |
| 37 | 121.50 | 19 | 15 | 14876 | 724 | >sp\|P08238\|HS90B_HUMAN Heat shock protein HSP 90-beta OS=Homo sapiens OX=9606 GN=HSP90AB1 PE=1 SV=4 |
| 38 | 117.29 | 18 | 18 | 7399 | 4646 | >sp\|Q14204\|DYHC1_HUMAN Cytoplasmic dynein 1 heavy chain 1 OS=Homo sapiens OX=9606 GN=DYNC1H1 PE=1 SV=5 |
| 39 | 113.39 | 16 | 12 | 14295 | 679 | >sp\|P38646\|GRP75_HUMAN Stress-70 protein; mitochondrial OS=Homo sapiens OX=9606 GN=HSPA9 PE=1 SV=2 |
| 40 | 111.90 | 19 | 16 | 37885 | 1018 | >sp\|P22314-2\|UBA1_HUMAN Isoform 2 of Ubiquitin-like modifier-activating enzyme 1 OS=Homo sapiens OX=9606 GN=UBA1 |
| 41 | 110.85 | 16 | 13 | 4219 | 935 | >sp\|P11586\|C1TC_HUMAN C-1-tetrahydrofolate synthase; cytoplasmic OS=Homo sapiens OX=9606 GN=MTHFD1 PE=1 SV=3 |

| | | | | | | |
|---|---|---|---|---|---|---|
| 42 | 110.81 | 9 | 9 | 2722 | 529 | >sp\|P06576\|ATPB_HUMAN ATP synthase subunit beta; mitochondrial OS=Homo sapiens OX=9606 GN=ATP5F1B PE=1 SV=3 |
| 43 | 109.35 | 14 | 13 | 16442 | 511 | >sp\|P05787-2\|K2C8_HUMAN Isoform 2 of Keratin; type II cytoskeletal 8 OS=Homo sapiens OX=9606 GN=KRT8 |
| 44 | 107.05 | 15 | 13 | 16301 | 406 | >sp\|P60842\|IF4A1_HUMAN Eukaryotic initiation factor 4A-I OS=Homo sapiens OX=9606 GN=EIF4A1 PE=1 SV=1 |
| 45 | 106.75 | 15 | 13 | 11581 | 898 | >sp\|Q12906-7\|ILF3_HUMAN Isoform 7 of Interleukin enhancer-binding factor 3 OS=Homo sapiens OX=9606 GN=ILF3 |
| 46 | 106.70 | 20 | 18 | 17290 | 1382 | >sp\|Q14152\|EIF3A_HUMAN Eukaryotic translation initiation factor 3 subunit A OS=Homo sapiens OX=9606 GN=EIF3A PE=1 SV=1 |
| 47 | 106.37 | 16 | 14 | 13423 | 531 | >sp\|P14618-2\|KPYM_HUMAN Isoform M1 of Pyruvate kinase PKM OS=Homo sapiens OX=9606 GN=PKM |
| 48 | 106.31 | 17 | 13 | 28052 | 393 | >sp\|Q8NC51-3\|PAIRB_HUMAN Isoform 3 of Plasminogen activator inhibitor 1 RNA-binding protein OS=Homo sapiens OX=9606 GN=SERBP1 |
| 49 | 103.87 | 16 | 13 | 5624 | 1639 | >sp\|Q00610-2\|CLH1_HUMAN Isoform 2 of Clathrin heavy chain 1 OS=Homo sapiens OX=9606 GN=CLTC |
| 50 | 103.42 | 12 | 12 | 37650 | 962 | >sp\|Q15029-3\|U5S1_HUMAN Isoform 3 of 116 kDa U5 small nuclear ribonucleoprotein component OS=Homo sapiens OX=9606 GN=EFTUD2 |
| 51 | 103.20 | 17 | 16 | 18698 | 382 | >sp\|Q15233-2\|NONO_HUMAN Isoform 2 of Non-POU domain-containing octamer-binding protein OS=Homo sapiens OX=9606 GN=NONO |
| 52 | 102.40 | 12 | 9 | 27427 | 254 | >sp\|P18669\|PGAM1_HUMAN Phosphoglycerate mutase 1 OS=Homo sapiens OX=9606 GN=PGAM1 PE=1 SV=2 |
| 53 | 101.20 | 13 | 8 | 40882 | 248 | >sp\|P06753-5\|TPM3_HUMAN Isoform 5 of Tropomyosin alpha-3 chain OS=Homo sapiens OX=9606 GN=TPM3 |
| 54 | 98.56 | 15 | 12 | 11476 | 817 | >sp\|Q92598-3\|HS105_HUMAN Isoform 3 of Heat shock protein 105 kDa OS=Homo sapiens OX=9606 GN=HSPH1 |
| 55 | 96.59 | 12 | 12 | 38968 | 448 | >sp\|P48643-2\|TCPE_HUMAN Isoform 2 of T-complex protein 1 subunit epsilon OS=Homo sapiens OX=9606 GN=CCT5 |
| 56 | 95.77 | 15 | 13 | 29243 | 199 | >sp\|Q06830\|PRDX1_HUMAN Peroxiredoxin-1 OS=Homo sapiens OX=9606 GN=PRDX1 PE=1 SV=1 |
| 57 | 94.29 | 14 | 13 | 7610 | 1270 | >sp\|Q08211\|DHX9_HUMAN ATP-dependent RNA helicase A OS=Homo sapiens OX=9606 GN=DHX9 PE=1 SV=4 |
| 58 | 94.19 | 14 | 14 | 41146 | 2541 | >sp\|Q9Y490\|TLN1_HUMAN Talin-1 OS=Homo sapiens OX=9606 GN=TLN1 PE=1 SV=3 |
| 59 | 93.77 | 11 | 11 | 3770 | 255 | >sp\|P62258\|1433E_HUMAN 14-3-3 protein epsilon OS=Homo sapiens OX=9606 GN=YWHAE PE=1 SV=1 |
| 60 | 92.41 | 11 | 10 | 12512 | 440 | >sp\|P61978-3\|HNRPK_HUMAN Isoform 3 of Heterogeneous nuclear ribonucleoprotein K OS=Homo sapiens OX=9606 GN=HNRNPK |
| 61 | 91.66 | 14 | 13 | 41256 | 609 | >sp\|P12956\|XRCC6_HUMAN X-ray repair cross-complementing protein 6 OS=Homo sapiens OX=9606 GN=XRCC6 PE=1 SV=2 |
| 62 | 90.53 | 17 | 13 | 20532 | 707 | >sp\|P23246\|SFPQ_HUMAN Splicing factor; proline- and glutamine-rich OS=Homo sapiens OX=9606 GN=SFPQ PE=1 SV=2 |
| 63 | 90.32 | 19 | 15 | 32061 | 577 | >sp\|P26038\|MOES_HUMAN Moesin OS=Homo sapiens OX=9606 GN=MSN PE=1 SV=3 |
| 64 | 90.07 | 13 | 12 | 38884 | 835 | >sp\|Q13263\|TIF1B_HUMAN Transcription intermediary factor 1-beta OS=Homo sapiens OX=9606 GN=TRIM28 PE=1 SV=5 |
| 65 | 90.05 | 15 | 15 | 30819 | 710 | >sp\|P19338\|NUCL_HUMAN Nucleolin OS=Homo sapiens OX=9606 GN=NCL PE=1 SV=3 |
| 66 | 87.33 | 16 | 13 | 28584 | 505 | >sp\|P30101\|PDIA3_HUMAN Protein disulfide-isomerase A3 OS=Homo sapiens OX=9606 GN=PDIA3 PE=1 SV=4 |
| 67 | 86.74 | 13 | 12 | 23083 | 1233 | >sp\|Q14683\|SMC1A_HUMAN Structural maintenance of chromosomes protein 1A OS=Homo sapiens OX=9606 GN=SMC1A PE=1 SV=2 |
| 68 | 86.49 | 14 | 14 | 29644 | 1394 | >sp\|P42704\|LPPRC_HUMAN Leucine-rich PPR motif-containing protein; mitochondrial OS=Homo sapiens OX=9606 GN=LRPPRC PE=1 SV=3 |
| 69 | 85.92 | 16 | 14 | 30785 | 636 | >sp\|P11940\|PABP1_HUMAN Polyadenylate-binding protein 1 OS=Homo sapiens OX=9606 GN=PABPC1 PE=1 SV=2 |
| 70 | 85.31 | 10 | 9 | 13614 | 963 | >sp\|P33176\|KINH_HUMAN Kinesin-1 heavy chain OS=Homo sapiens OX=9606 GN=KIF5B PE=1 SV=1 |
| 71 | 85.16 | 13 | 12 | 23683 | 2452 | >sp\|Q13813-3\|SPTN1_HUMAN Isoform 3 of Spectrin alpha chain; non-erythrocytic 1 OS=Homo sapiens OX=9606 GN=SPTAN1 |
| 72 | 84.50 | 11 | 11 | 14882 | 840 | >sp\|P34932\|HSP74_HUMAN Heat shock 70 kDa protein 4 OS=Homo sapiens OX=9606 GN=HSPA4 PE=1 SV=4 |
| 73 | 83.40 | 12 | 12 | 22200 | 320 | >sp\|P09651-2\|ROA1_HUMAN Isoform A1-A of Heterogeneous nuclear ribonucleoprotein A1 OS=Homo sapiens OX=9606 GN=HNRNPA1 |
| 74 | 83.08 | 12 | 11 | 38497 | 531 | >sp\|P40227\|TCPZ_HUMAN T-complex protein 1 subunit zeta OS=Homo sapiens OX=9606 GN=CCT6A PE=1 SV=3 |
| 75 | 82.84 | 9 | 8 | 34464 | 245 | >sp\|P63104\|1433Z_HUMAN 14-3-3 protein zeta/delta OS=Homo sapiens OX=9606 GN=YWHAZ PE=1 SV=1 |
| 76 | 81.04 | 15 | 11 | 34826 | 451 | >sp\|P68363\|TBA1B_HUMAN Tubulin alpha-1B chain OS=Homo sapiens OX=9606 GN=TUBA1B PE=1 SV=1 |
| 77 | 80.84 | 9 | 7 | 38832 | 556 | >sp\|P17987\|TCPA_HUMAN T-complex protein 1 subunit alpha OS=Homo sapiens OX=9606 GN=TCP1 PE=1 SV=1 |
| 78 | 80.02 | 12 | 10 | 12655 | 293 | >sp\|P04406-2\|G3P_HUMAN Isoform 2 of Glyceraldehyde-3-phosphate dehydrogenase OS=Homo sapiens OX=9606 GN=GAPDH |
| 79 | 79.59 | 8 | 8 | 42044 | 317 | >sp\|P63244\|RACK1_HUMAN Receptor of activated protein C kinase 1 OS=Homo sapiens OX=9606 GN=RACK1 PE=1 SV=3 |
| 80 | 79.55 | 16 | 14 | 9793 | 966 | >sp\|Q14697-2\|GANAB_HUMAN Isoform 2 of Neutral alpha-glucosidase AB OS=Homo sapiens OX=9606 GN=GANAB |
| 81 | 79.42 | 12 | 10 | 3913 | 364 | >sp\|P04075\|ALDOA_HUMAN Fructose-bisphosphate aldolase A OS=Homo sapiens OX=9606 GN=ALDOA PE=1 SV=2 |
| 82 | 79.16 | 10 | 9 | 16289 | 381 | >sp\|P12277\|KCRB_HUMAN Creatine kinase B-type OS=Homo sapiens OX=9606 GN=CKB PE=1 SV=1 |
| 83 | 78.58 | 8 | 7 | 22096 | 295 | >sp\|P08865\|RSSA_HUMAN 40S ribosomal protein SA OS=Homo sapiens OX=9606 GN=RPSA PE=1 SV=4 |
| 84 | 77.59 | 9 | 8 | 41436 | 283 | >sp\|P21796\|VDAC1_HUMAN Voltage-dependent anion-selective channel protein 1 OS=Homo sapiens OX=9606 GN=VDAC1 PE=1 SV=2 |
| 85 | 76.94 | 10 | 10 | 663 | 992 | >sp\|P05023-3\|AT1A1_HUMAN Isoform 3 of Sodium/potassium-transporting ATPase subunit alpha-1 OS=Homo sapiens OX=9606 GN=ATP1A1 |
| 86 | 76.91 | 9 | 8 | 41398 | 1066 | >sp\|P18206-2\|VINC_HUMAN Isoform 1 of Vinculin OS=Homo sapiens OX=9606 GN=VCL |
| 87 | 76.77 | 9 | 8 | 12428 | 306 | >sp\|Q14103-3\|HNRPD_HUMAN Isoform 3 of Heterogeneous nuclear ribonucleoprotein D0 OS=Homo sapiens OX=9606 GN=HNRNPD |
| 88 | 76.69 | 15 | 12 | 21434 | 427 | >sp\|P36578\|RL4_HUMAN 60S ribosomal protein L4 OS=Homo sapiens OX=9606 GN=RPL4 PE=1 SV=5 |
| 89 | 76.36 | 8 | 8 | 19342 | 463 | >sp\|Q9Y230\|RUVB2_HUMAN RuvB-like 2 OS=Homo sapiens OX=9606 GN=RUVBL2 PE=1 SV=3 |
| 90 | 76.08 | 9 | 8 | 7218 | 257 | >sp\|P29692-3\|EF1D_HUMAN Isoform 3 of Elongation factor 1-delta OS=Homo sapiens OX=9606 GN=EEF1D |
| 91 | 75.35 | 13 | 11 | 12431 | 588 | >sp\|O60506-2\|HNRPQ_HUMAN Isoform 2 of Heterogeneous nuclear ribonucleoprotein Q OS=Homo sapiens OX=9606 GN=SYNCRIP |
| 92 | 75.17 | 10 | 10 | 42264 | 908 | >sp\|Q13200\|PSMD2_HUMAN 26S proteasome non-ATPase regulatory subunit 2 OS=Homo sapiens OX=9606 GN=PSMD2 PE=1 SV=3 |
| 93 | 74.62 | 8 | 8 | 19742 | 432 | >sp\|P23526\|SAHH_HUMAN Adenosylhomocysteinase OS=Homo sapiens OX=9606 GN=AHCY PE=1 SV=4 |
| 94 | 74.47 | 9 | 8 | 20411 | 533 | >sp\|O43175\|SERA_HUMAN D-3-phosphoglycerate dehydrogenase OS=Homo sapiens OX=9606 GN=PHGDH PE=1 SV=4 |
| 95 | 74.34 | 7 | 7 | 6958 | 652 | >sp\|Q92841-3\|DDX17_HUMAN Isoform 4 of Probable ATP-dependent RNA helicase DDX17 OS=Homo sapiens OX=9606 GN=DDX17 |
| 96 | 74.24 | 10 | 10 | 16431 | 430 | >sp\|P05783\|K1C18_HUMAN Keratin; type I cytoskeletal 18 OS=Homo sapiens OX=9606 GN=KRT18 PE=1 SV=2 |
| 97 | 72.65 | 8 | 8 | 38249 | 499 | >sp\|Q99832-3\|TCPH_HUMAN Isoform 3 of T-complex protein 1 subunit eta OS=Homo sapiens OX=9606 GN=CCT7 |
| 98 | 72.20 | 6 | 4 | 22777 | 110 | >sp\|P06454-2\|PTMA_HUMAN Isoform 2 of Prothymosin alpha OS=Homo sapiens OX=9606 GN=PTMA |
| 99 | 71.29 | 12 | 11 | 31097 | 331 | >sp\|Q9Y266\|NUDC_HUMAN Nuclear migration protein nudC OS=Homo sapiens OX=9606 GN=NUDC PE=1 SV=1 |

| | | | | | | |
|---|---|---|---|---|---|---|
| 100 | 70.70 | 8 | 8 | 11111 | 653 | >sp\|Q96AE4-2\|FUBP1_HUMAN Isoform 2 of Far upstream element-binding protein 1 OS=Homo sapiens OX=9606 GN=FUBP1 |
| 101 | 70.54 | 15 | 12 | 15619 | 459 | >sp\|Q02790\|FKBP4_HUMAN Peptidyl-prolyl cis-trans isomerase FKBP4 OS=Homo sapiens OX=9606 GN=FKBP4 PE=1 SV=3 |
| 102 | 70.47 | 10 | 7 | 37757 | 223 | >sp\|P09936\|UCHL1_HUMAN Ubiquitin carboxyl-terminal hydrolase isozyme L1 OS=Homo sapiens OX=9606 GN=UCHL1 PE=1 SV=2 |
| 103 | 70.34 | 8 | 7 | 29443 | 271 | >sp\|Q13162\|PRDX4_HUMAN Peroxiredoxin-4 OS=Homo sapiens OX=9606 GN=PRDX4 PE=1 SV=1 |
| 104 | 70.08 | 9 | 9 | 22202 | 353 | >sp\|P22626\|ROA2_HUMAN Heterogeneous nuclear ribonucleoproteins A2/B1 OS=Homo sapiens OX=9606 GN=HNRNPA2B1 PE=1 SV=2 |
| 105 | 70.00 | 8 | 6 | 1226 | 166 | >sp\|P23528\|COF1_HUMAN Cofilin-1 OS=Homo sapiens OX=9606 GN=CFL1 PE=1 SV=3 |
| 106 | 69.98 | 9 | 7 | 29326 | 165 | >sp\|P62937\|PPIA_HUMAN Peptidyl-prolyl cis-trans isomerase A OS=Homo sapiens OX=9606 GN=PPIA PE=1 SV=2 |
| 107 | 69.60 | 9 | 8 | 4123 | 1230 | >sp\|Q86VP6\|CAND1_HUMAN Cullin-associated NEDD8-dissociated protein 1 OS=Homo sapiens OX=9606 GN=CAND1 PE=1 SV=2 |
| 108 | 69.04 | 9 | 8 | 28314 | 645 | >sp\|P13667\|PDIA4_HUMAN Protein disulfide-isomerase A4 OS=Homo sapiens OX=9606 GN=PDIA4 PE=1 SV=2 |
| 109 | 68.82 | 9 | 7 | 6433 | 102 | >sp\|P61604\|CH10_HUMAN 10 kDa heat shock protein; mitochondrial OS=Homo sapiens OX=9606 GN=HSPE1 PE=1 SV=2 |
| 110 | 68.65 | 12 | 12 | 28888 | 919 | >sp\|P55786\|PSA_HUMAN Puromycin-sensitive aminopeptidase OS=Homo sapiens OX=9606 GN=NPEPPS PE=1 SV=2 |
| 111 | 67.83 | 12 | 9 | 7519 | 462 | >sp\|Q5VTE0\|EF1A3_HUMAN Putative elongation factor 1-alpha-like 3 OS=Homo sapiens OX=9606 GN=EEF1A1P5 PE=5 SV=1 |
| 112 | 67.71 | 9 | 8 | 35782 | 756 | >sp\|P26639-2\|SYTC_HUMAN Isoform 2 of Threonine--tRNA ligase; cytoplasmic OS=Homo sapiens OX=9606 GN=TARS |
| 113 | 67.25 | 8 | 8 | 17245 | 353 | >sp\|Q15717-2\|ELAV1_HUMAN Isoform 2 of ELAV-like protein 1 OS=Homo sapiens OX=9606 GN=ELAVL1 |
| 114 | 67.00 | 7 | 7 | 19303 | 293 | >sp\|P15880\|RS2_HUMAN 40S ribosomal protein S2 OS=Homo sapiens OX=9606 GN=RPS2 PE=1 SV=2 |
| 115 | 66.13 | 9 | 9 | 11626 | 1115 | >sp\|O00410-3\|IPO5_HUMAN Isoform 3 of Importin-5 OS=Homo sapiens OX=9606 GN=IPO5 |
| 116 | 65.88 | 6 | 5 | 13900 | 210 | >sp\|P09211\|GSTP1_HUMAN Glutathione S-transferase P OS=Homo sapiens OX=9606 GN=GSTP1 PE=1 SV=2 |
| 117 | 65.65 | 7 | 7 | 28589 | 488 | >sp\|Q15084-5\|PDIA6_HUMAN Isoform 5 of Protein disulfide-isomerase A6 OS=Homo sapiens OX=9606 GN=PDIA6 |
| 118 | 65.31 | 10 | 10 | 22230 | 259 | >sp\|P23396-2\|RS3_HUMAN Isoform 2 of 40S ribosomal protein S3 OS=Homo sapiens OX=9606 GN=RPS3 |
| 119 | 64.94 | 8 | 8 | 20444 | 313 | >sp\|O43765\|SGTA_HUMAN Small glutamine-rich tetratricopeptide repeat-containing protein alpha OS=Homo sapiens OX=9606 GN=SGTA PE |
| 120 | 64.94 | 7 | 7 | 12734 | 232 | >sp\|P54819-2\|KAD2_HUMAN Isoform 2 of Adenylate kinase 2; mitochondrial OS=Homo sapiens OX=9606 GN=AK2 |
| 121 | 64.34 | 6 | 5 | 27451 | 187 | >sp\|P30086\|PEBP1_HUMAN Phosphatidylethanolamine-binding protein 1 OS=Homo sapiens OX=9606 GN=PEBP1 PE=1 SV=3 |
| 122 | 64.22 | 10 | 10 | 40982 | 2363 | >sp\|P12270\|TPR_HUMAN Nucleoprotein TPR OS=Homo sapiens OX=9606 GN=TPR PE=1 SV=3 |
| 123 | 63.97 | 10 | 10 | 40575 | 651 | >sp\|Q12931-2\|TRAP1_HUMAN Isoform 2 of Heat shock protein 75 kDa; mitochondrial OS=Homo sapiens OX=9606 GN=TRAP1 |
| 124 | 63.64 | 8 | 7 | 7279 | 443 | >sp\|Q13838-2\|DX39B_HUMAN Isoform 2 of Spliceosome RNA helicase DDX39B OS=Homo sapiens OX=9606 GN=DDX39B |
| 125 | 63.42 | 6 | 6 | 26541 | 620 | >sp\|Q03252\|LMNB2_HUMAN Lamin-B2 OS=Homo sapiens OX=9606 GN=LMNB2 PE=1 SV=4 |
| 126 | 63.39 | 8 | 8 | 14631 | 577 | >sp\|Q9NZI8\|IF2B1_HUMAN Insulin-like growth factor 2 mRNA-binding protein 1 OS=Homo sapiens OX=9606 GN=IGF2BP1 PE=1 SV=2 |
| 127 | 63.24 | 9 | 8 | 34468 | 589 | >sp\|P30153\|2AAA_HUMAN Serine/threonine-protein phosphatase 2A 65 kDa regulatory subunit A alpha isoform OS=Homo sapiens OX=9606 |
| 128 | 63.07 | 9 | 8 | 28625 | 299 | >sp\|Q99623\|PHB2_HUMAN Prohibitin-2 OS=Homo sapiens OX=9606 GN=PHB2 PE=1 SV=2 |
| 129 | 61.86 | 12 | 10 | 27434 | 417 | >sp\|P00558\|PGK1_HUMAN Phosphoglycerate kinase 1 OS=Homo sapiens OX=9606 GN=PGK1 PE=1 SV=3 |
| 130 | 60.92 | 6 | 6 | 12513 | 633 | >sp\|O43390\|HNRPR_HUMAN Heterogeneous nuclear ribonucleoprotein R OS=Homo sapiens OX=9606 GN=HNRNPR PE=1 SV=1 |
| 131 | 60.83 | 12 | 12 | 21886 | 264 | >sp\|P61247\|RS3A_HUMAN 40S ribosomal protein S3a OS=Homo sapiens OX=9606 GN=RPS3A PE=1 SV=2 |
| 132 | 60.34 | 9 | 8 | 14646 | 1606 | >sp\|Q04637-9\|IF4G1_HUMAN Isoform 9 of Eukaryotic translation initiation factor 4 gamma 1 OS=Homo sapiens OX=9606 GN=EIF4G1 |
| 133 | 60.05 | 6 | 5 | 3563 | 430 | >sp\|P00505\|AATM_HUMAN Aspartate aminotransferase; mitochondrial OS=Homo sapiens OX=9606 GN=GOT2 PE=1 SV=3 |
| 134 | 59.81 | 11 | 10 | 15628 | 2633 | >sp\|O75369-8\|FLNB_HUMAN Isoform 8 of Filamin-B OS=Homo sapiens OX=9606 GN=FLNB |
| 135 | 59.65 | 6 | 6 | 10934 | 873 | >sp\|P55884-2\|EIF3B_HUMAN Isoform 2 of Eukaryotic translation initiation factor 3 subunit B OS=Homo sapiens OX=9606 GN=EIF3B |
| 136 | 59.46 | 15 | 12 | 30781 | 340 | >sp\|Q9UQ80-2\|PA2G4_HUMAN Isoform 2 of Proliferation-associated protein 2G4 OS=Homo sapiens OX=9606 GN=PA2G4 |
| 137 | 59.22 | 6 | 6 | 27443 | 272 | >sp\|P35232\|PHB_HUMAN Prohibitin OS=Homo sapiens OX=9606 GN=PHB PE=1 SV=1 |
| 138 | 59.06 | 6 | 6 | 12860 | 694 | >sp\|P42166\|LAP2A_HUMAN Lamina-associated polypeptide 2; isoform alpha OS=Homo sapiens OX=9606 GN=TMPO PE=1 SV=2 |
| 139 | 57.70 | 10 | 9 | 16626 | 289 | >sp\|Q15181\|IPYR_HUMAN Inorganic pyrophosphatase OS=Homo sapiens OX=9606 GN=PPA1 PE=1 SV=2 |
| 140 | 57.47 | 7 | 6 | 6960 | 715 | >sp\|Q9NR30-2\|DDX21_HUMAN Isoform 2 of Nucleolar RNA helicase 2 OS=Homo sapiens OX=9606 GN=DDX21 |
| 141 | 57.45 | 10 | 9 | 5190 | 417 | >sp\|P27797\|CALR_HUMAN Calreticulin OS=Homo sapiens OX=9606 GN=CALR PE=1 SV=1 |
| 142 | 57.26 | 9 | 9 | 37210 | 324 | >sp\|P67809\|YBOX1_HUMAN Nuclease-sensitive element-binding protein 1 OS=Homo sapiens OX=9606 GN=YBX1 PE=1 SV=3 |
| 143 | 57.01 | 7 | 6 | 24904 | 249 | >sp\|P39687\|AN32A_HUMAN Acidic leucine-rich nuclear phosphoprotein 32 family member A OS=Homo sapiens OX=9606 GN=ANP32A PE=1 SV= |
| 144 | 56.53 | 11 | 8 | 38242 | 108 | >sp\|O75347\|TBCA_HUMAN Tubulin-specific chaperone A OS=Homo sapiens OX=9606 GN=TBCA PE=1 SV=3 |
| 145 | 56.26 | 8 | 8 | 15101 | 1657 | >sp\|P46940\|IQGA1_HUMAN Ras GTPase-activating-like protein IQGAP1 OS=Homo sapiens OX=9606 GN=IQGAP1 PE=1 SV=1 |
| 146 | 56.25 | 10 | 10 | 34627 | 1512 | >sp\|P07814\|SYEP_HUMAN Bifunctional glutamate/proline--tRNA ligase OS=Homo sapiens OX=9606 GN=EPRS PE=1 SV=5 |
| 147 | 56.23 | 10 | 9 | 22920 | 1197 | >sp\|O95347\|SMC2_HUMAN Structural maintenance of chromosomes protein 2 OS=Homo sapiens OX=9606 GN=SMC2 PE=1 SV=2 |
| 148 | 56.17 | 7 | 7 | 22539 | 423 | >sp\|O00231-2\|PSD11_HUMAN Isoform 2 of 26S proteasome non-ATPase regulatory subunit 11 OS=Homo sapiens OX=9606 GN=PSMD11 |
| 149 | 55.81 | 6 | 6 | 26001 | 338 | >sp\|P40926\|MDHM_HUMAN Malate dehydrogenase; mitochondrial OS=Homo sapiens OX=9606 GN=MDH2 PE=1 SV=3 |
| 150 | 55.74 | 8 | 6 | 15672 | 525 | >sp\|P35637-2\|FUS_HUMAN Isoform Short of RNA-binding protein FUS OS=Homo sapiens OX=9606 GN=FUS |
| 151 | 55.57 | 8 | 8 | 34806 | 220 | >sp\|P37802-2\|TAGL2_HUMAN Isoform 2 of Transgelin-2 OS=Homo sapiens OX=9606 GN=TAGLN2 |
| 152 | 55.51 | 7 | 7 | 24592 | 357 | >sp\|P07355-2\|ANXA2_HUMAN Isoform 2 of Annexin A2 OS=Homo sapiens OX=9606 GN=ANXA2 |
| 153 | 55.04 | 6 | 6 | 9255 | 321 | >sp\|P22087\|FBRL_HUMAN rRNA 2'-O-methyltransferase fibrillarin OS=Homo sapiens OX=9606 GN=FBL PE=1 SV=2 |

| | | | | | | |
|---|---|---|---|---|---|---|
| 154 | 54.92 | 5 | 5 | 35309 | 427 | >sp\|P24752\|THIL_HUMAN Acetyl-CoA acetyltransferase; mitochondrial OS=Homo sapiens OX=9606 GN=ACAT1 PE=1 SV=1 |
| 155 | 54.89 | 10 | 10 | 9596 | 483 | >sp\|P34897-3\|GLYM_HUMAN Isoform 3 of Serine hydroxymethyltransferase; mitochondrial OS=Homo sapiens OX=9606 GN=SHMT2 |
| 156 | 54.52 | 6 | 6 | 26393 | 614 | >sp\|P02545-6\|LMNA_HUMAN Isoform 6 of Prelamin-A/C OS=Homo sapiens OX=9606 GN=LMNA |
| 157 | 53.85 | 4 | 4 | 16304 | 572 | >sp\|P23588-2\|IF4B_HUMAN Isoform 2 of Eukaryotic translation initiation factor 4B OS=Homo sapiens OX=9606 GN=EIF4B |
| 158 | 53.44 | 9 | 8 | 20530 | 793 | >sp\|Q15459\|SF3A1_HUMAN Splicing factor 3A subunit 1 OS=Homo sapiens OX=9606 GN=SF3A1 PE=1 SV=1 |
| 159 | 52.99 | 7 | 6 | 25642 | 591 | >sp\|P17812\|PYRG1_HUMAN CTP synthase 1 OS=Homo sapiens OX=9606 GN=CTPS1 PE=1 SV=2 |
| 160 | 52.97 | 8 | 6 | 4295 | 627 | >sp\|P27824-2\|CALX_HUMAN Isoform 2 of Calnexin OS=Homo sapiens OX=9606 GN=CANX |
| 161 | 52.78 | 4 | 4 | 18702 | 386 | >sp\|Q99733-2\|NP1L4_HUMAN Isoform 2 of Nucleosome assembly protein 1-like 4 OS=Homo sapiens OX=9606 GN=NAP1L4 |
| 162 | 52.48 | 8 | 7 | 13288 | 361 | >sp\|P00338-3\|LDHA_HUMAN Isoform 3 of L-lactate dehydrogenase A chain OS=Homo sapiens OX=9606 GN=LDHA |
| 163 | 52.39 | 5 | 5 | 34655 | 588 | >sp\|P54136-2\|SYRC_HUMAN Isoform Monomeric of Arginine--tRNA ligase; cytoplasmic OS=Homo sapiens OX=9606 GN=RARS |
| 164 | 52.05 | 5 | 5 | 19134 | 211 | >sp\|P26373\|RL13_HUMAN 60S ribosomal protein L13 OS=Homo sapiens OX=9606 GN=RPL13 PE=1 SV=4 |
| 165 | 51.86 | 7 | 7 | 500 | 504 | >sp\|Q9BZZ5-2\|API5_HUMAN Isoform 2 of Apoptosis inhibitor 5 OS=Homo sapiens OX=9606 GN=API5 |
| 166 | 51.41 | 7 | 6 | 36094 | 363 | >sp\|Q9Y3F4-2\|STRAP_HUMAN Isoform 2 of Serine-threonine kinase receptor-associated protein OS=Homo sapiens OX=9606 GN=STRAP |
| 167 | 51.32 | 7 | 7 | 9850 | 525 | >sp\|P14314-2\|GLU2B_HUMAN Isoform 2 of Glucosidase 2 subunit beta OS=Homo sapiens OX=9606 GN=PRKCSH |
| 168 | 51.26 | 10 | 10 | 22334 | 592 | >sp\|P31939\|PUR9_HUMAN Bifunctional purine biosynthesis protein PURH OS=Homo sapiens OX=9606 GN=ATIC PE=1 SV=3 |
| 169 | 50.94 | 7 | 7 | 12280 | 449 | >sp\|P31943\|HNRH1_HUMAN Heterogeneous nuclear ribonucleoprotein H OS=Homo sapiens OX=9606 GN=HNRNPH1 PE=1 SV=4 |
| 170 | 50.65 | 7 | 7 | 32152 | 845 | >sp\|P46087-4\|NOP2_HUMAN Isoform 4 of Probable 28S rRNA (cytosine(4447)-C(5))-methyltransferase OS=Homo sapiens OX=9606 GN=NOP2 |
| 171 | 50.64 | 9 | 9 | 36275 | 550 | >sp\|Q14247\|SRC8_HUMAN Src substrate cortactin OS=Homo sapiens OX=9606 GN=CTTN PE=1 SV=2 |
| 172 | 50.61 | 6 | 5 | 11014 | 224 | >sp\|Q00688\|FKBP3_HUMAN Peptidyl-prolyl cis-trans isomerase FKBP3 OS=Homo sapiens OX=9606 GN=FKBP3 PE=1 SV=1 |
| 173 | 50.53 | 8 | 8 | 22232 | 263 | >sp\|P62701\|RS4X_HUMAN 40S ribosomal protein S4; X isoform OS=Homo sapiens OX=9606 GN=RPS4X PE=1 SV=2 |
| 174 | 50.47 | 7 | 6 | 4739 | 475 | >sp\|Q01518\|CAP1_HUMAN Adenylyl cyclase-associated protein 1 OS=Homo sapiens OX=9606 GN=CAP1 PE=1 SV=5 |
| 175 | 50.37 | 7 | 6 | 29256 | 140 | >sp\|P07737\|PROF1_HUMAN Profilin-1 OS=Homo sapiens OX=9606 GN=PFN1 PE=1 SV=2 |
| 176 | 50.30 | 5 | 4 | 41068 | 248 | >sp\|P67936\|TPM4_HUMAN Tropomyosin alpha-4 chain OS=Homo sapiens OX=9606 GN=TPM4 PE=1 SV=3 |
| 177 | 50.17 | 5 | 5 | 10792 | 600 | >sp\|Q01844-6\|EWS_HUMAN Isoform 6 of RNA-binding protein EWS OS=Homo sapiens OX=9606 GN=EWSR1 |
| 178 | 49.84 | 6 | 6 | 21436 | 255 | >sp\|P05388-2\|RLA0_HUMAN Isoform 2 of 60S acidic ribosomal protein P0 OS=Homo sapiens OX=9606 GN=RPLP0 |
| 179 | 49.78 | 6 | 5 | 34462 | 246 | >sp\|P31946\|1433B_HUMAN 14-3-3 protein beta/alpha OS=Homo sapiens OX=9606 GN=YWHAB PE=1 SV=3 |
| 180 | 49.76 | 8 | 7 | 18778 | 265 | >sp\|P06748-2\|NPM_HUMAN Isoform 2 of Nucleophosmin OS=Homo sapiens OX=9606 GN=NPM1 |
| 181 | 49.66 | 4 | 4 | 12196 | 181 | >sp\|Q13442\|HAP28_HUMAN 28 kDa heat- and acid-stable phosphoprotein OS=Homo sapiens OX=9606 GN=PDAP1 PE=1 SV=1 |
| 182 | 49.63 | 11 | 11 | 19307 | 152 | >sp\|P62269\|RS18_HUMAN 40S ribosomal protein S18 OS=Homo sapiens OX=9606 GN=RPS18 PE=1 SV=3 |
| 183 | 49.56 | 13 | 12 | 626 | 586 | >sp\|Q9NVI7-2\|ATD3A_HUMAN Isoform 2 of ATPase family AAA domain-containing protein 3A OS=Homo sapiens OX=9606 GN=ATAD3A |
| 184 | 49.48 | 11 | 10 | 11247 | 452 | >sp\|P49411\|EFTU_HUMAN Elongation factor Tu; mitochondrial OS=Homo sapiens OX=9606 GN=TUFM PE=1 SV=2 |
| 185 | 49.15 | 8 | 7 | 36556 | 796 | >sp\|Q96QK1\|VPS35_HUMAN Vacuolar protein sorting-associated protein 35 OS=Homo sapiens OX=9606 GN=VPS35 PE=1 SV=2 |
| 186 | 48.92 | 6 | 6 | 21332 | 257 | >sp\|P62917\|RL8_HUMAN 60S ribosomal protein L8 OS=Homo sapiens OX=9606 GN=RPL8 PE=1 SV=2 |
| 187 | 48.85 | 11 | 9 | 18998 | 378 | >sp\|P38159-2\|RBMX_HUMAN Isoform 2 of RNA-binding motif protein; X chromosome OS=Homo sapiens OX=9606 GN=RBMX |
| 188 | 48.55 | 7 | 7 | 1462 | 2039 | >sp\|Q14008-3\|CKAP5_HUMAN Isoform 3 of Cytoskeleton-associated protein 5 OS=Homo sapiens OX=9606 GN=CKAP5 |
| 189 | 48.40 | 10 | 8 | 16376 | 514 | >sp\|P12268\|IMDH2_HUMAN Inosine-5'-monophosphate dehydrogenase 2 OS=Homo sapiens OX=9606 GN=IMPDH2 PE=1 SV=2 |
| 190 | 48.39 | 5 | 5 | 22726 | 432 | >sp\|P22234-2\|PUR6_HUMAN Isoform 2 of Multifunctional protein ADE2 OS=Homo sapiens OX=9606 GN=PAICS |
| 191 | 48.10 | 9 | 7 | 30993 | 793 | >sp\|P54886-2\|P5CS_HUMAN Isoform Short of Delta-1-pyrroline-5-carboxylate synthase OS=Homo sapiens OX=9606 GN=ALDH18A1 |
| 192 | 47.80 | 7 | 5 | 21380 | 297 | >sp\|P46777\|RL5_HUMAN 60S ribosomal protein L5 OS=Homo sapiens OX=9606 GN=RPL5 PE=1 SV=3 |
| 193 | 47.74 | 6 | 5 | 12190 | 221 | >sp\|P16402\|H13_HUMAN Histone H1.3 OS=Homo sapiens OX=9606 GN=HIST1H1D PE=1 SV=2 |
| 194 | 47.33 | 5 | 5 | 42257 | 246 | >sp\|P60900\|PSA6_HUMAN Proteasome subunit alpha type-6 OS=Homo sapiens OX=9606 GN=PSMA6 PE=1 SV=1 |
| 195 | 47.32 | 6 | 6 | 37907 | 1104 | >sp\|Q14157-5\|UBP2L_HUMAN Isoform 5 of Ubiquitin-associated protein 2-like OS=Homo sapiens OX=9606 GN=UBAP2L |
| 196 | 47.31 | 7 | 7 | 34764 | 528 | >sp\|P54577\|SYYC_HUMAN Tyrosine--tRNA ligase; cytoplasmic OS=Homo sapiens OX=9606 GN=YARS PE=1 SV=4 |
| 197 | 47.01 | 11 | 6 | 12365 | 130 | >sp\|Q6FI13\|H2A2A_HUMAN Histone H2A type 2-A OS=Homo sapiens OX=9606 GN=HIST2H2AA3 PE=1 SV=3 |
| 198 | 46.84 | 10 | 9 | 15545 | 569 | >sp\|P06744-2\|G6PI_HUMAN Isoform 2 of Glucose-6-phosphate isomerase OS=Homo sapiens OX=9606 GN=GPI |
| 199 | 46.75 | 8 | 7 | 10904 | 290 | >sp\|P30084\|ECHM_HUMAN Enoyl-CoA hydratase; mitochondrial OS=Homo sapiens OX=9606 GN=ECHS1 PE=1 SV=4 |
| 200 | 46.61 | 9 | 9 | 13911 | 977 | >sp\|P78347-4\|GTF2I_HUMAN Isoform 4 of General transcription factor II-I OS=Homo sapiens OX=9606 GN=GTF2I |
| 201 | 46.45 | 6 | 5 | 36772 | 732 | >sp\|P13010\|XRCC5_HUMAN X-ray repair cross-complementing protein 5 OS=Homo sapiens OX=9606 GN=XRCC5 PE=1 SV=3 |
| 202 | 46.41 | 7 | 7 | 11102 | 493 | >sp\|Q16658\|FSCN1_HUMAN Fascin OS=Homo sapiens OX=9606 GN=FSCN1 PE=1 SV=3 |
| 203 | 46.38 | 9 | 7 | 22345 | 216 | >sp\|P62826\|RAN_HUMAN GTP-binding nuclear protein Ran OS=Homo sapiens OX=9606 GN=RAN PE=1 SV=3 |
| 204 | 46.37 | 5 | 5 | 9191 | 369 | >sp\|P50502\|F10A1_HUMAN Hsc70-interacting protein OS=Homo sapiens OX=9606 GN=ST13 PE=1 SV=2 |
| 205 | 46.29 | 6 | 5 | 36192 | 221 | >sp\|Q13242\|SRSF9_HUMAN Serine/arginine-rich splicing factor 9 OS=Homo sapiens OX=9606 GN=SRSF9 PE=1 SV=1 |
| 206 | 46.28 | 4 | 4 | 28433 | 366 | >sp\|Q15366-2\|PCBP2_HUMAN Isoform 2 of Poly(rC)-binding protein 2 OS=Homo sapiens OX=9606 GN=PCBP2 |
| 207 | 46.15 | 10 | 7 | 31862 | 267 | >sp\|P22392-2\|NDKB_HUMAN Isoform 3 of Nucleoside diphosphate kinase B OS=Homo sapiens OX=9606 GN=NME2 |
| 208 | 46.15 | 5 | 4 | 35685 | 548 | >sp\|O43776\|SYNC_HUMAN Asparagine--tRNA ligase; cytoplasmic OS=Homo sapiens OX=9606 GN=NARS PE=1 SV=1 |
| 209 | 45.91 | 8 | 7 | 42249 | 433 | >sp\|P35998\|PRS7_HUMAN 26S proteasome regulatory subunit 7 OS=Homo sapiens OX=9606 GN=PSMC2 PE=1 SV=3 |
| 210 | 45.87 | 9 | 9 | 4021 | 641 | >sp\|P08133-2\|ANXA6_HUMAN Isoform 2 of Annexin A6 OS=Homo sapiens OX=9606 GN=ANXA6 |

| | | | | | | |
|---|---|---|---|---|---|---|
| 211 | 45.83 | 7 | 6 | 21936 | 505 | >sp\|Q9Y3I0\|RTCB_HUMAN tRNA-splicing ligase RtcB homolog OS=Homo sapiens OX=9606 GN=RTCB PE=1 SV=1 |
| 212 | 45.72 | 3 | 3 | 18182 | 925 | >sp\|E9PAV3-2\|NACAM_HUMAN Isoform skNAC-2 of Nascent polypeptide-associated complex subunit alpha; muscle-specific form OS=Homo |
| 213 | 45.71 | 10 | 8 | 7432 | 487 | >sp\|P26641-2\|EF1G_HUMAN Isoform 2 of Elongation factor 1-gamma OS=Homo sapiens OX=9606 GN=EEF1G |
| 214 | 45.67 | 9 | 8 | 42301 | 1338 | >sp\|O15067\|PUR4_HUMAN Phosphoribosylformylglycinamidine synthase OS=Homo sapiens OX=9606 GN=PFAS PE=1 SV=4 |
| 215 | 45.51 | 4 | 4 | 34390 | 245 | >sp\|P27348\|1433T_HUMAN 14-3-3 protein theta OS=Homo sapiens OX=9606 GN=YWHAQ PE=1 SV=1 |
| 216 | 45.45 | 7 | 5 | 38831 | 127 | >sp\|P53999\|TCP4_HUMAN Activated RNA polymerase II transcriptional coactivator p15 OS=Homo sapiens OX=9606 GN=SUB1 PE=1 SV=3 |
| 217 | 45.40 | 6 | 5 | 20643 | 331 | >sp\|Q15293\|RCN1_HUMAN Reticulocalbin-1 OS=Homo sapiens OX=9606 GN=RCN1 PE=1 SV=1 |
| 218 | 45.38 | 6 | 6 | 1175 | 283 | >sp\|Q15417-3\|CNN3_HUMAN Isoform 3 of Calponin-3 OS=Homo sapiens OX=9606 GN=CNN3 |
| 219 | 45.33 | 9 | 8 | 21381 | 248 | >sp\|P18124\|RL7_HUMAN 60S ribosomal protein L7 OS=Homo sapiens OX=9606 GN=RPL7 PE=1 SV=1 |
| 220 | 45.31 | 7 | 6 | 22453 | 439 | >sp\|P17980\|PRS6A_HUMAN 26S proteasome regulatory subunit 6A OS=Homo sapiens OX=9606 GN=PSMC3 PE=1 SV=3 |
| 221 | 45.05 | 5 | 5 | 14720 | 414 | >sp\|O75874\|IDHC_HUMAN Isocitrate dehydrogenase [NADP] cytoplasmic OS=Homo sapiens OX=9606 GN=IDH1 PE=1 SV=2 |
| 222 | 44.91 | 4 | 4 | 19305 | 208 | >sp\|P62241\|RS8_HUMAN 40S ribosomal protein S8 OS=Homo sapiens OX=9606 GN=RPS8 PE=1 SV=2 |
| 223 | 44.49 | 5 | 5 | 15591 | 204 | >sp\|P52565\|GDIR1_HUMAN Rho GDP-dissociation inhibitor 1 OS=Homo sapiens OX=9606 GN=ARHGDIA PE=1 SV=3 |
| 224 | 44.32 | 6 | 6 | 23207 | 290 | >sp\|Q01105\|SET_HUMAN Protein SET OS=Homo sapiens OX=9606 GN=SET PE=1 SV=3 |
| 225 | 44.22 | 5 | 5 | 15783 | 572 | >sp\|Q96I24\|FUBP3_HUMAN Far upstream element-binding protein 3 OS=Homo sapiens OX=9606 GN=FUBP3 PE=1 SV=2 |
| 226 | 43.98 | 5 | 4 | 18951 | 587 | >sp\|P46060\|RAGP1_HUMAN Ran GTPase-activating protein 1 OS=Homo sapiens OX=9606 GN=RANGAP1 PE=1 SV=1 |
| 227 | 43.87 | 7 | 7 | 14985 | 529 | >sp\|P52292\|IMA1_HUMAN Importin subunit alpha-1 OS=Homo sapiens OX=9606 GN=KPNA2 PE=1 SV=1 |
| 228 | 43.81 | 6 | 6 | 29328 | 216 | >sp\|P23284\|PPIB_HUMAN Peptidyl-prolyl cis-trans isomerase B OS=Homo sapiens OX=9606 GN=PPIB PE=1 SV=2 |
| 229 | 43.42 | 7 | 5 | 36287 | 248 | >sp\|Q07955\|SRSF1_HUMAN Serine/arginine-rich splicing factor 1 OS=Homo sapiens OX=9606 GN=SRSF1 PE=1 SV=2 |
| 230 | 43.39 | 6 | 5 | 8893 | 478 | >sp\|Q16630-3\|CPSF6_HUMAN Isoform 3 of Cleavage and polyadenylation specificity factor subunit 6 OS=Homo sapiens OX=9606 GN=CPSF |
| 231 | 43.16 | 12 | 9 | 41253 | 915 | >sp\|P55060-4\|XPO2_HUMAN Isoform 4 of Exportin-2 OS=Homo sapiens OX=9606 GN=CSE1L |
| 232 | 42.99 | 4 | 4 | 29290 | 178 | >sp\|O14818-2\|PSA7_HUMAN Isoform 2 of Proteasome subunit alpha type-7 OS=Homo sapiens OX=9606 GN=PSMA7 |
| 233 | 42.88 | 4 | 4 | 4409 | 277 | >sp\|P16152\|CBR1_HUMAN Carbonyl reductase [NADPH] 1 OS=Homo sapiens OX=9606 GN=CBR1 PE=1 SV=3 |
| 234 | 42.57 | 6 | 5 | 19252 | 135 | >sp\|P08708\|RS17_HUMAN 40S ribosomal protein S17 OS=Homo sapiens OX=9606 GN=RPS17 PE=1 SV=2 |
| 235 | 42.47 | 11 | 11 | 33358 | 853 | >sp\|P25205-2\|MCM3_HUMAN Isoform 2 of DNA replication licensing factor MCM3 OS=Homo sapiens OX=9606 GN=MCM3 |
| 236 | 42.42 | 4 | 4 | 21364 | 115 | >sp\|P05387\|RLA2_HUMAN 60S acidic ribosomal protein P2 OS=Homo sapiens OX=9606 GN=RPLP2 PE=1 SV=1 |
| 237 | 42.15 | 6 | 6 | 2886 | 869 | >sp\|Q9NYF8-3\|BCLF1_HUMAN Isoform 3 of Bcl-2-associated transcription factor 1 OS=Homo sapiens OX=9606 GN=BCLAF1 |
| 238 | 42.09 | 3 | 3 | 34830 | 445 | >sp\|Q9BVA1\|TBB2B_HUMAN Tubulin beta-2B chain OS=Homo sapiens OX=9606 GN=TUBB2B PE=1 SV=1 |
| 239 | 42.01 | 8 | 6 | 11386 | 126 | >sp\|Q99880\|H2B1L_HUMAN Histone H2B type 1-L OS=Homo sapiens OX=9606 GN=HIST1H2BL PE=1 SV=3 |
| 240 | 41.75 | 6 | 6 | 21802 | 188 | >sp\|Q07020\|RL18_HUMAN 60S ribosomal protein L18 OS=Homo sapiens OX=9606 GN=RPL18 PE=1 SV=2 |
| 241 | 41.58 | 6 | 6 | 21474 | 266 | >sp\|P62424\|RL7A_HUMAN 60S ribosomal protein L7a OS=Homo sapiens OX=9606 GN=RPL7A PE=1 SV=2 |
| 242 | 41.54 | 6 | 6 | 21314 | 196 | >sp\|P84098\|RL19_HUMAN 60S ribosomal protein L19 OS=Homo sapiens OX=9606 GN=RPL19 PE=1 SV=1 |
| 243 | 41.50 | 7 | 6 | 11200 | 466 | >sp\|Q13283\|G3BP1_HUMAN Ras GTPase-activating protein-binding protein 1 OS=Homo sapiens OX=9606 GN=G3BP1 PE=1 SV=1 |
| 244 | 41.26 | 5 | 4 | 21316 | 148 | >sp\|P46776\|RL27A_HUMAN 60S ribosomal protein L27a OS=Homo sapiens OX=9606 GN=RPL27A PE=1 SV=2 |
| 245 | 40.75 | 5 | 5 | 13269 | 432 | >sp\|O95232\|LC7L3_HUMAN Luc7-like protein 3 OS=Homo sapiens OX=9606 GN=LUC7L3 PE=1 SV=2 |
| 246 | 40.68 | 5 | 5 | 33005 | 352 | >sp\|P40925-3\|MDHC_HUMAN Isoform 3 of Malate dehydrogenase; cytoplasmic OS=Homo sapiens OX=9606 GN=MDH1 |
| 247 | 40.60 | 3 | 3 | 26853 | 125 | >sp\|Q9NRX4\|PHP14_HUMAN 14 kDa phosphohistidine phosphatase OS=Homo sapiens OX=9606 GN=PHPT1 PE=1 SV=1 |
| 248 | 40.49 | 5 | 4 | 21526 | 192 | >sp\|P32969\|RL9_HUMAN 60S ribosomal protein L9 OS=Homo sapiens OX=9606 GN=RPL9 PE=1 SV=1 |
| 249 | 40.45 | 6 | 6 | 38851 | 139 | >sp\|Q15185-4\|TEBP_HUMAN Isoform 4 of Prostaglandin E synthase 3 OS=Homo sapiens OX=9606 GN=PTGES3 |
| 250 | 40.44 | 7 | 7 | 20623 | 669 | >sp\|Q96PK6\|RBM14_HUMAN RNA-binding protein 14 OS=Homo sapiens OX=9606 GN=RBM14 PE=1 SV=2 |
| 251 | 40.29 | 7 | 6 | 7017 | 738 | >sp\|O00429-8\|DNM1L_HUMAN Isoform 8 of Dynamin-1-like protein OS=Homo sapiens OX=9606 GN=DNM1L |
| 252 | 40.15 | 10 | 8 | 35063 | 539 | >sp\|P50991\|TCPD_HUMAN T-complex protein 1 subunit delta OS=Homo sapiens OX=9606 GN=CCT4 PE=1 SV=4 |
| 253 | 40.12 | 7 | 7 | 19840 | 1304 | >sp\|O75533\|SF3B1_HUMAN Splicing factor 3B subunit 1 OS=Homo sapiens OX=9606 GN=SF3B1 PE=1 SV=3 |
| 254 | 39.85 | 6 | 6 | 21287 | 156 | >sp\|P62750\|RL23A_HUMAN 60S ribosomal protein L23a OS=Homo sapiens OX=9606 GN=RPL23A PE=1 SV=1 |
| 255 | 39.66 | 4 | 3 | 39025 | 193 | >sp\|Q99426-2\|TBCB_HUMAN Isoform 2 of Tubulin-folding cofactor B OS=Homo sapiens OX=9606 GN=TBCB |
| 256 | 39.66 | 8 | 7 | 34835 | 502 | >sp\|O95793-3\|STAU1_HUMAN Isoform 3 of Double-stranded RNA-binding protein Staufen homolog 1 OS=Homo sapiens OX=9606 GN=STAU1 |
| 257 | 39.59 | 3 | 3 | 28453 | 125 | >sp\|O14737\|PDCD5_HUMAN Programmed cell death protein 5 OS=Homo sapiens OX=9606 GN=PDCD5 PE=1 SV=3 |
| 258 | 39.35 | 8 | 8 | 10546 | 357 | >sp\|O00303\|EIF3F_HUMAN Eukaryotic translation initiation factor 3 subunit F OS=Homo sapiens OX=9606 GN=EIF3F PE=1 SV=1 |
| 259 | 39.32 | 4 | 4 | 21948 | 244 | >sp\|Q9Y224\|RTRAF_HUMAN RNA transcription; translation and transport factor protein OS=Homo sapiens OX=9606 GN=RTRAF PE=1 SV=1 |
| 260 | 39.32 | 8 | 8 | 23403 | 1288 | >sp\|Q9NTJ3\|SMC4_HUMAN Structural maintenance of chromosomes protein 4 OS=Homo sapiens OX=9606 GN=SMC4 PE=1 SV=2 |
| 261 | 38.91 | 7 | 6 | 27891 | 508 | >sp\|P07237\|PDIA1_HUMAN Protein disulfide-isomerase OS=Homo sapiens OX=9606 GN=P4HB PE=1 SV=3 |
| 262 | 38.85 | 5 | 5 | 19310 | 249 | >sp\|P62753\|RS6_HUMAN 40S ribosomal protein S6 OS=Homo sapiens OX=9606 GN=RPS6 PE=1 SV=1 |
| 263 | 38.73 | 6 | 6 | 20536 | 322 | >sp\|Q9H9B4\|SFXN1_HUMAN Sideroflexin-1 OS=Homo sapiens OX=9606 GN=SFXN1 PE=1 SV=4 |
| 264 | 38.72 | 8 | 8 | 13122 | 408 | >sp\|P05455\|LA_HUMAN Lupus La protein OS=Homo sapiens OX=9606 GN=SSB PE=1 SV=2 |
| 265 | 38.63 | 3 | 3 | 5435 | 378 | >sp\|Q16543\|CDC37_HUMAN Hsp90 co-chaperone Cdc37 OS=Homo sapiens OX=9606 GN=CDC37 PE=1 SV=1 |
| 266 | 38.45 | 5 | 5 | 30919 | 439 | >sp\|P04181\|OAT_HUMAN Ornithine aminotransferase; mitochondrial OS=Homo sapiens OX=9606 GN=OAT PE=1 SV=1 |
| 267 | 38.36 | 6 | 5 | 19133 | 214 | >sp\|P27635\|RL10_HUMAN 60S ribosomal protein L10 OS=Homo sapiens OX=9606 GN=RPL10 PE=1 SV=4 |
| 268 | 38.35 | 4 | 4 | 7891 | 367 | >sp\|Q9Y295\|DRG1_HUMAN Developmentally-regulated GTP-binding protein 1 OS=Homo sapiens OX=9606 GN=DRG1 PE=1 SV=1 |
| 269 | 38.26 | 4 | 4 | 3129 | 313 | >sp\|P51572-2\|BAP31_HUMAN Isoform 2 of B-cell receptor-associated protein 31 OS=Homo sapiens OX=9606 GN=BCAP31 |

| | | | | | | |
|---|---|---|---|---|---|---|
| 270 | 38.21 | 4 | 4 | 13707 | 261 | >sp\|Q99714\|HCD2_HUMAN 3-hydroxyacyl-CoA dehydrogenase type-2 OS=Homo sapiens OX=9606 GN=HSD17B10 PE=1 SV=3 |
| 271 | 38.08 | 7 | 5 | 6908 | 494 | >sp\|Q99615\|DNJC7_HUMAN DnaJ homolog subfamily C member 7 OS=Homo sapiens OX=9606 GN=DNAJC7 PE=1 SV=2 |
| 272 | 38.00 | 3 | 3 | 28847 | 158 | >sp\|P24666\|PPAC_HUMAN Low molecular weight phosphotyrosine protein phosphatase OS=Homo sapiens OX=9606 GN=ACP1 PE=1 SV=3 |
| 273 | 37.99 | 7 | 6 | 38916 | 337 | >sp\|P37837\|TALDO_HUMAN Transaldolase OS=Homo sapiens OX=9606 GN=TALDO1 PE=1 SV=2 |
| 274 | 37.94 | 5 | 5 | 25995 | 543 | >sp\|P33993-3\|MCM7_HUMAN Isoform 3 of DNA replication licensing factor MCM7 OS=Homo sapiens OX=9606 GN=MCM7 |
| 275 | 37.63 | 6 | 5 | 26244 | 711 | >sp\|P49959-3\|MRE11_HUMAN Isoform 3 of Double-strand break repair protein MRE11 OS=Homo sapiens OX=9606 GN=MRE11 |
| 276 | 37.61 | 7 | 6 | 3616 | 1091 | >sp\|P53396-2\|ACLY_HUMAN Isoform 2 of ATP-citrate synthase OS=Homo sapiens OX=9606 GN=ACLY |
| 277 | 37.42 | 5 | 5 | 4017 | 346 | >sp\|P04083\|ANXA1_HUMAN Annexin A1 OS=Homo sapiens OX=9606 GN=ANXA1 PE=1 SV=2 |
| 278 | 37.41 | 5 | 5 | 21113 | 1118 | >sp\|Q92900-2\|RENT1_HUMAN Isoform 2 of Regulator of nonsense transcripts 1 OS=Homo sapiens OX=9606 GN=UPF1 |
| 279 | 37.41 | 5 | 5 | 24897 | 488 | >sp\|P28838-2\|AMPL_HUMAN Isoform 2 of Cytosol aminopeptidase OS=Homo sapiens OX=9606 GN=LAP3 |
| 280 | 37.33 | 6 | 6 | 34612 | 298 | >sp\|P05141\|ADT2_HUMAN ADP/ATP translocase 2 OS=Homo sapiens OX=9606 GN=SLC25A5 PE=1 SV=7 |
| 281 | 37.33 | 5 | 4 | 21879 | 217 | >sp\|P62906\|RL10A_HUMAN 60S ribosomal protein L10a OS=Homo sapiens OX=9606 GN=RPL10A PE=1 SV=2 |
| 282 | 37.25 | 4 | 4 | 25053 | 1074 | >sp\|Q96KR1\|ZFR_HUMAN Zinc finger RNA-binding protein OS=Homo sapiens OX=9606 GN=ZFR PE=1 SV=2 |
| 283 | 37.00 | 7 | 7 | 38830 | 1412 | >sp\|Q13428-8\|TCOF_HUMAN Isoform 8 of Treacle protein OS=Homo sapiens OX=9606 GN=TCOF1 |
| 284 | 36.86 | 6 | 6 | 19251 | 151 | >sp\|P62277\|RS13_HUMAN 40S ribosomal protein S13 OS=Homo sapiens OX=9606 GN=RPS13 PE=1 SV=2 |
| 285 | 36.66 | 6 | 6 | 17292 | 320 | >sp\|O75821\|EIF3G_HUMAN Eukaryotic translation initiation factor 3 subunit G OS=Homo sapiens OX=9606 GN=EIF3G PE=1 SV=2 |
| 286 | 36.62 | 4 | 4 | 34203 | 246 | >sp\|Q04917\|1433F_HUMAN 14-3-3 protein eta OS=Homo sapiens OX=9606 GN=YWHAH PE=1 SV=4 |
| 287 | 36.49 | 9 | 8 | 23623 | 910 | >sp\|Q7KZF4\|SND1_HUMAN Staphylococcal nuclease domain-containing protein 1 OS=Homo sapiens OX=9606 GN=SND1 PE=1 SV=1 |
| 288 | 36.39 | 9 | 8 | 7600 | 795 | >sp\|O43143\|DHX15_HUMAN Pre-mRNA-splicing factor ATP-dependent RNA helicase DHX15 OS=Homo sapiens OX=9606 GN=DHX15 PE=1 SV=2 |
| 289 | 36.24 | 5 | 4 | 26503 | 489 | >sp\|O75439\|MPPB_HUMAN Mitochondrial-processing peptidase subunit beta OS=Homo sapiens OX=9606 GN=PMPCB PE=1 SV=2 |
| 290 | 36.14 | 5 | 5 | 41390 | 584 | >sp\|P38606-2\|VATA_HUMAN Isoform 2 of V-type proton ATPase catalytic subunit A OS=Homo sapiens OX=9606 GN=ATP6V1A |
| 291 | 36.04 | 5 | 4 | 32551 | 732 | >sp\|Q08J23-2\|NSUN2_HUMAN Isoform 2 of tRNA (cytosine(34)-C(5))-methyltransferase OS=Homo sapiens OX=9606 GN=NSUN2 |
| 292 | 36.01 | 8 | 8 | 22188 | 145 | >sp\|P39019\|RS19_HUMAN 40S ribosomal protein S19 OS=Homo sapiens OX=9606 GN=RPS19 PE=1 SV=2 |
| 293 | 35.98 | 4 | 4 | 33831 | 611 | >sp\|P09960\|LKHA4_HUMAN Leukotriene A-4 hydrolase OS=Homo sapiens OX=9606 GN=LTA4H PE=1 SV=2 |
| 294 | 35.76 | 4 | 4 | 3321 | 180 | >sp\|P18085\|ARF4_HUMAN ADP-ribosylation factor 4 OS=Homo sapiens OX=9606 GN=ARF4 PE=1 SV=3 |
| 295 | 35.76 | 9 | 7 | 4742 | 272 | >sp\|P47756-2\|CAPZB_HUMAN Isoform 2 of F-actin-capping protein subunit beta OS=Homo sapiens OX=9606 GN=CAPZB |
| 296 | 35.73 | 3 | 3 | 28590 | 329 | >sp\|O00151\|PDLI1_HUMAN PDZ and LIM domain protein 1 OS=Homo sapiens OX=9606 GN=PDLIM1 PE=1 SV=4 |
| 297 | 35.68 | 5 | 5 | 42031 | 207 | >sp\|P51149\|RAB7A_HUMAN Ras-related protein Rab-7a OS=Homo sapiens OX=9606 GN=RAB7A PE=1 SV=1 |
| 298 | 35.68 | 5 | 4 | 41727 | 283 | >sp\|P45880-2\|VDAC2_HUMAN Isoform 2 of Voltage-dependent anion-selective channel protein 2 OS=Homo sapiens OX=9606 GN=VDAC2 |
| 299 | 35.67 | 5 | 4 | 34519 | 470 | >sp\|P52209-2\|6PGD_HUMAN Isoform 2 of 6-phosphogluconate dehydrogenase; decarboxylating OS=Homo sapiens OX=9606 GN=PGD |
| 300 | 35.55 | 7 | 5 | 21288 | 140 | >sp\|P62829\|RL23_HUMAN 60S ribosomal protein L23 OS=Homo sapiens OX=9606 GN=RPL23 PE=1 SV=1 |
| 301 | 35.53 | 8 | 8 | 21309 | 203 | >sp\|P40429\|RL13A_HUMAN 60S ribosomal protein L13a OS=Homo sapiens OX=9606 GN=RPL13A PE=1 SV=2 |
| 302 | 35.37 | 5 | 5 | 16422 | 218 | >sp\|Q9H910-3\|JUPI2_HUMAN Isoform 3 of Jupiter microtubule associated homolog 2 OS=Homo sapiens OX=9606 GN=JPT2 |
| 303 | 35.28 | 4 | 4 | 16834 | 3256 | >sp\|P46013\|KI67_HUMAN Proliferation marker protein Ki-67 OS=Homo sapiens OX=9606 GN=MKI67 PE=1 SV=2 |
| 304 | 34.94 | 5 | 4 | 23973 | 402 | >sp\|O60749-2\|SNX2_HUMAN Isoform 2 of Sorting nexin-2 OS=Homo sapiens OX=9606 GN=SNX2 |
| 305 | 34.89 | 7 | 6 | 17443 | 516 | >sp\|Q9Y262-2\|EIF3L_HUMAN Isoform 2 of Eukaryotic translation initiation factor 3 subunit L OS=Homo sapiens OX=9606 GN=EIF3L |
| 306 | 34.78 | 5 | 5 | 10935 | 325 | >sp\|Q13347\|EIF3I_HUMAN Eukaryotic translation initiation factor 3 subunit I OS=Homo sapiens OX=9606 GN=EIF3I PE=1 SV=1 |
| 307 | 34.73 | 5 | 4 | 18934 | 249 | >sp\|P51148-2\|RAB5C_HUMAN Isoform 2 of Ras-related protein Rab-5C OS=Homo sapiens OX=9606 GN=RAB5C |
| 308 | 34.71 | 5 | 5 | 12561 | 346 | >sp\|P31942\|HNRH3_HUMAN Heterogeneous nuclear ribonucleoprotein H3 OS=Homo sapiens OX=9606 GN=HNRNPH3 PE=1 SV=2 |
| 309 | 34.65 | 6 | 5 | 8044 | 622 | >sp\|Q92499-2\|DDX1_HUMAN Isoform 2 of ATP-dependent RNA helicase DDX1 OS=Homo sapiens OX=9606 GN=DDX1 |
| 310 | 34.38 | 9 | 8 | 19311 | 194 | >sp\|P46781\|RS9_HUMAN 40S ribosomal protein S9 OS=Homo sapiens OX=9606 GN=RPS9 PE=1 SV=3 |
| 311 | 34.34 | 5 | 5 | 20209 | 370 | >sp\|Q9Y617\|SERC_HUMAN Phosphoserine aminotransferase OS=Homo sapiens OX=9606 GN=PSAT1 PE=1 SV=2 |
| 312 | 34.19 | 3 | 3 | 38613 | 444 | >sp\|P07437\|TBB5_HUMAN Tubulin beta chain OS=Homo sapiens OX=9606 GN=TUBB PE=1 SV=2 |
| 313 | 34.06 | 4 | 4 | 10165 | 533 | >sp\|Q8N8S7-3\|ENAH_HUMAN Isoform 3 of Protein enabled homolog OS=Homo sapiens OX=9606 GN=ENAH |
| 314 | 34.03 | 5 | 4 | 23408 | 120 | >sp\|P62318-2\|SMD3_HUMAN Isoform 2 of Small nuclear ribonucleoprotein Sm D3 OS=Homo sapiens OX=9606 GN=SNRPD3 |
| 315 | 34.01 | 6 | 5 | 12625 | 589 | >sp\|P14866\|HNRPL_HUMAN Heterogeneous nuclear ribonucleoprotein L OS=Homo sapiens OX=9606 GN=HNRNPL PE=1 SV=2 |
| 316 | 34.00 | 6 | 6 | 7558 | 1031 | >sp\|Q7L014\|DDX46_HUMAN Probable ATP-dependent RNA helicase DDX46 OS=Homo sapiens OX=9606 GN=DDX46 PE=1 SV=2 |
| 317 | 33.70 | 3 | 3 | 20681 | 305 | >sp\|P40938-2\|RFC3_HUMAN Isoform 2 of Replication factor C subunit 3 OS=Homo sapiens OX=9606 GN=RFC3 |
| 318 | 33.66 | 5 | 5 | 22581 | 409 | >sp\|P54727\|RD23B_HUMAN UV excision repair protein RAD23 homolog B OS=Homo sapiens OX=9606 GN=RAD23B PE=1 SV=1 |
| 319 | 33.61 | 6 | 5 | 22391 | 364 | >sp\|Q9BWF3\|RBM4_HUMAN RNA-binding protein 4 OS=Homo sapiens OX=9606 GN=RBM4 PE=1 SV=1 |
| 320 | 33.58 | 4 | 4 | 21885 | 69 | >sp\|P62857\|RS28_HUMAN 40S ribosomal protein S28 OS=Homo sapiens OX=9606 GN=RPS28 PE=1 SV=1 |
| 321 | 33.49 | 8 | 6 | 16999 | 346 | >sp\|P38117-2\|ETFB_HUMAN Isoform 2 of Electron transfer flavoprotein subunit beta OS=Homo sapiens OX=9606 GN=ETFB |
| 322 | 33.38 | 6 | 6 | 8033 | 819 | >sp\|Q86XP3-2\|DDX42_HUMAN Isoform 2 of ATP-dependent RNA helicase DDX42 OS=Homo sapiens OX=9606 GN=DDX42 |
| 323 | 33.21 | 6 | 5 | 19292 | 146 | >sp\|P62249\|RS16_HUMAN 40S ribosomal protein S16 OS=Homo sapiens OX=9606 GN=RPS16 PE=1 SV=2 |
| 324 | 32.78 | 6 | 6 | 21386 | 288 | >sp\|Q02878\|RL6_HUMAN 60S ribosomal protein L6 OS=Homo sapiens OX=9606 GN=RPL6 PE=1 SV=3 |

| | | | | | | |
|---|---|---|---|---|---|---|
| 325 | 32.69 | 4 | 4 | 2781 | 213 | >sp\|P48047\|ATPO_HUMAN ATP synthase subunit O; mitochondrial OS=Homo sapiens OX=9606 GN=ATP5O PE=1 SV=1 |
| 326 | 32.64 | 3 | 3 | 2845 | 173 | >sp\|P80723-2\|BASP1_HUMAN Isoform 2 of Brain acid soluble protein 1 OS=Homo sapiens OX=9606 GN=BASP1 |
| 327 | 32.52 | 4 | 4 | 9589 | 335 | >sp\|O76003\|GLRX3_HUMAN Glutaredoxin-3 OS=Homo sapiens OX=9606 GN=GLRX3 PE=1 SV=2 |
| 328 | 32.44 | 3 | 3 | 27422 | 154 | >sp\|Q9UHV9\|PFD2_HUMAN Prefoldin subunit 2 OS=Homo sapiens OX=9606 GN=PFDN2 PE=1 SV=1 |
| 329 | 32.29 | 5 | 4 | 41614 | 471 | >sp\|P26368-2\|U2AF2_HUMAN Isoform 2 of Splicing factor U2AF 65 kDa subunit OS=Homo sapiens OX=9606 GN=U2AF2 |
| 330 | 32.05 | 7 | 5 | 14930 | 184 | >sp\|P63241-2\|IF5A1_HUMAN Isoform 2 of Eukaryotic translation initiation factor 5A-1 OS=Homo sapiens OX=9606 GN=EIF5A |
| 331 | 31.96 | 5 | 5 | 11406 | 2079 | >sp\|P51610-4\|HCFC1_HUMAN Isoform 4 of Host cell factor 1 OS=Homo sapiens OX=9606 GN=HCFC1 |
| 332 | 31.84 | 3 | 3 | 18056 | 529 | >sp\|Q9Y2X3\|NOP58_HUMAN Nucleolar protein 58 OS=Homo sapiens OX=9606 GN=NOP58 PE=1 SV=1 |
| 333 | 31.83 | 3 | 3 | 13299 | 1329 | >sp\|Q86UP2-4\|KTN1_HUMAN Isoform 4 of Kinectin OS=Homo sapiens OX=9606 GN=KTN1 |
| 334 | 31.61 | 8 | 5 | 9361 | 282 | >sp\|P10768\|ESTD_HUMAN S-formylglutathione hydrolase OS=Homo sapiens OX=9606 GN=ESD PE=1 SV=2 |
| 335 | 31.39 | 7 | 7 | 28812 | 224 | >sp\|P30041\|PRDX6_HUMAN Peroxiredoxin-6 OS=Homo sapiens OX=9606 GN=PRDX6 PE=1 SV=3 |
| 336 | 31.36 | 3 | 3 | 7555 | 670 | >sp\|Q9NVP1\|DDX18_HUMAN ATP-dependent RNA helicase DDX18 OS=Homo sapiens OX=9606 GN=DDX18 PE=1 SV=2 |
| 337 | 31.33 | 4 | 4 | 38686 | 426 | >sp\|Q9BWD1-2\|THIC_HUMAN Isoform 2 of Acetyl-CoA acetyltransferase; cytosolic OS=Homo sapiens OX=9606 GN=ACAT2 |
| 338 | 30.99 | 5 | 5 | 17478 | 261 | >sp\|P30040\|ERP29_HUMAN Endoplasmic reticulum resident protein 29 OS=Homo sapiens OX=9606 GN=ERP29 PE=1 SV=4 |
| 339 | 30.94 | 7 | 7 | 8085 | 2871 | >sp\|P15924\|DESP_HUMAN Desmoplakin OS=Homo sapiens OX=9606 GN=DSP PE=1 SV=3 |
| 340 | 30.87 | 6 | 5 | 19265 | 607 | >sp\|P04843\|RPN1_HUMAN Dolichyl-diphosphooligosaccharide--protein glycosyltransferase subunit 1 OS=Homo sapiens OX=9606 GN=RPN1 |
| 341 | 30.80 | 4 | 4 | 19880 | 456 | >sp\|Q9Y265\|RUVB1_HUMAN RuvB-like 1 OS=Homo sapiens OX=9606 GN=RUVBL1 PE=1 SV=1 |
| 342 | 30.78 | 5 | 4 | 4954 | 694 | >sp\|Q14444-2\|CAPR1_HUMAN Isoform 2 of Caprin-1 OS=Homo sapiens OX=9606 GN=CAPRIN1 |
| 343 | 30.62 | 4 | 4 | 7703 | 375 | >sp\|P35659\|DEK_HUMAN Protein DEK OS=Homo sapiens OX=9606 GN=DEK PE=1 SV=1 |
| 344 | 30.61 | 3 | 3 | 13871 | 240 | >sp\|P51858\|HDGF_HUMAN Hepatoma-derived growth factor OS=Homo sapiens OX=9606 GN=HDGF PE=1 SV=1 |
| 345 | 30.51 | 4 | 4 | 37663 | 170 | >sp\|Q13404-7\|UB2V1_HUMAN Isoform 5 of Ubiquitin-conjugating enzyme E2 variant 1 OS=Homo sapiens OX=9606 GN=UBE2V1 |
| 346 | 30.33 | 3 | 3 | 42174 | 269 | >sp\|P25786-2\|PSA1_HUMAN Isoform Long of Proteasome subunit alpha type-1 OS=Homo sapiens OX=9606 GN=PSMA1 |
| 347 | 30.26 | 7 | 6 | 10547 | 914 | >sp\|B5ME19\|EIFCL_HUMAN Eukaryotic translation initiation factor 3 subunit C-like protein OS=Homo sapiens OX=9606 GN=EIF3CL PE=3 |
| 348 | 30.16 | 5 | 5 | 29283 | 398 | >sp\|P62195-2\|PRS8_HUMAN Isoform 2 of 26S proteasome regulatory subunit 8 OS=Homo sapiens OX=9606 GN=PSMC5 |
| 349 | 30.09 | 4 | 3 | 29308 | 330 | >sp\|P62136\|PP1A_HUMAN Serine/threonine-protein phosphatase PP1-alpha catalytic subunit OS=Homo sapiens OX=9606 GN=PPP1CA PE=1 S |
| 350 | 30.07 | 6 | 6 | 36164 | 2155 | >sp\|Q01082-3\|SPTB2_HUMAN Isoform 2 of Spectrin beta chain; non-erythrocytic 1 OS=Homo sapiens OX=9606 GN=SPTBN1 |
| 351 | 30.02 | 7 | 7 | 21378 | 165 | >sp\|P30050\|RL12_HUMAN 60S ribosomal protein L12 OS=Homo sapiens OX=9606 GN=RPL12 PE=1 SV=1 |
| 352 | 29.91 | 3 | 3 | 27996 | 873 | >sp\|Q8WUM4-2\|PDC6I_HUMAN Isoform 2 of Programmed cell death 6-interacting protein OS=Homo sapiens OX=9606 GN=PDCD6IP |
| 353 | 29.63 | 4 | 4 | 573 | 399 | >sp\|P61160-2\|ARP2_HUMAN Isoform 2 of Actin-related protein 2 OS=Homo sapiens OX=9606 GN=ACTR2 |
| 354 | 29.61 | 4 | 3 | 21697 | 332 | >sp\|Q99729\|ROAA_HUMAN Heterogeneous nuclear ribonucleoprotein A/B OS=Homo sapiens OX=9606 GN=HNRNPAB PE=1 SV=2 |
| 355 | 29.56 | 3 | 3 | 25870 | 395 | >sp\|P31153\|METK2_HUMAN S-adenosylmethionine synthase isoform type-2 OS=Homo sapiens OX=9606 GN=MAT2A PE=1 SV=1 |
| 356 | 29.49 | 5 | 4 | 12503 | 209 | >sp\|P26583\|HMGB2_HUMAN High mobility group protein B2 OS=Homo sapiens OX=9606 GN=HMGB2 PE=1 SV=2 |
| 357 | 29.46 | 2 | 2 | 21300 | 128 | >sp\|P35268\|RL22_HUMAN 60S ribosomal protein L22 OS=Homo sapiens OX=9606 GN=RPL22 PE=1 SV=2 |
| 358 | 29.24 | 4 | 3 | 10545 | 533 | >sp\|O15371-3\|EIF3D_HUMAN Isoform 3 of Eukaryotic translation initiation factor 3 subunit D OS=Homo sapiens OX=9606 GN=EIF3D |
| 359 | 29.24 | 5 | 4 | 19361 | 255 | >sp\|P09661\|RU2A_HUMAN U2 small nuclear ribonucleoprotein A' OS=Homo sapiens OX=9606 GN=SNRPA1 PE=1 SV=2 |
| 360 | 29.17 | 5 | 5 | 34873 | 1262 | >sp\|P41252\|SYIC_HUMAN Isoleucine--tRNA ligase; cytoplasmic OS=Homo sapiens OX=9606 GN=IARS PE=1 SV=2 |
| 361 | 29.02 | 6 | 6 | 31489 | 866 | >sp\|Q9BXJ9\|NAA15_HUMAN N-alpha-acetyltransferase 15; NatA auxiliary subunit OS=Homo sapiens OX=9606 GN=NAA15 PE=1 SV=1 |
| 362 | 29.00 | 3 | 3 | 37672 | 459 | >sp\|P54578-2\|UBP14_HUMAN Isoform 2 of Ubiquitin carboxyl-terminal hydrolase 14 OS=Homo sapiens OX=9606 GN=USP14 |
| 363 | 28.94 | 5 | 4 | 20186 | 439 | >sp\|Q9NVA2-2\|SEP11_HUMAN Isoform 2 of Septin-11 OS=Homo sapiens OX=9606 GN=SEPT11 |
| 364 | 28.88 | 3 | 3 | 22906 | 107 | >sp\|Q9GZT3-2\|SLIRP_HUMAN Isoform 2 of SRA stem-loop-interacting RNA-binding protein; mitochondrial OS=Homo sapiens OX=9606 GN=S |
| 365 | 28.88 | 4 | 4 | 16075 | 628 | >sp\|Q07866-6\|KLC1_HUMAN Isoform N of Kinesin light chain 1 OS=Homo sapiens OX=9606 GN=KLC1 |
| 366 | 28.82 | 3 | 3 | 32839 | 396 | >sp\|Q9NTK5\|OLA1_HUMAN Obg-like ATPase 1 OS=Homo sapiens OX=9606 GN=OLA1 PE=1 SV=2 |
| 367 | 28.68 | 4 | 4 | 34664 | 366 | >sp\|P52788\|SPSY_HUMAN Spermine synthase OS=Homo sapiens OX=9606 GN=SMS PE=1 SV=2 |
| 368 | 28.65 | 3 | 3 | 3384 | 134 | >sp\|P07741-2\|APT_HUMAN Isoform 2 of Adenine phosphoribosyltransferase OS=Homo sapiens OX=9606 GN=APRT |
| 369 | 28.57 | 3 | 3 | 7215 | 225 | >sp\|P24534\|EF1B_HUMAN Elongation factor 1-beta OS=Homo sapiens OX=9606 GN=EEF1B2 PE=1 SV=3 |
| 370 | 28.49 | 3 | 3 | 34204 | 247 | >sp\|P61981\|1433G_HUMAN 14-3-3 protein gamma OS=Homo sapiens OX=9606 GN=YWHAG PE=1 SV=2 |
| 371 | 28.42 | 4 | 4 | 21356 | 135 | >sp\|P62910\|RL32_HUMAN 60S ribosomal protein L32 OS=Homo sapiens OX=9606 GN=RPL32 PE=1 SV=2 |
| 372 | 28.35 | 3 | 3 | 32243 | 334 | >sp\|P00387-3\|NB5R3_HUMAN Isoform 3 of NADH-cytochrome b5 reductase 3 OS=Homo sapiens OX=9606 GN=CYB5R3 |
| 373 | 28.28 | 3 | 3 | 22948 | 225 | >sp\|O75934\|SPF27_HUMAN Pre-mRNA-splicing factor SPF27 OS=Homo sapiens OX=9606 GN=BCAS2 PE=1 SV=1 |
| 374 | 28.28 | 5 | 4 | 20408 | 371 | >sp\|Q15019-3\|SEPT2_HUMAN Isoform 3 of Septin-2 OS=Homo sapiens OX=9606 GN=SEPT2 |
| 375 | 28.28 | 9 | 7 | 40872 | 765 | >sp\|P11387\|TOP1_HUMAN DNA topoisomerase 1 OS=Homo sapiens OX=9606 GN=TOP1 PE=1 SV=2 |
| 376 | 28.19 | 4 | 4 | 2503 | 169 | >sp\|P13073\|COX41_HUMAN Cytochrome c oxidase subunit 4 isoform 1; mitochondrial OS=Homo sapiens OX=9606 GN=COX4I1 PE=1 SV=1 |
| 377 | 28.18 | 4 | 2 | 1459 | 602 | >sp\|Q07065\|CKAP4_HUMAN Cytoskeleton-associated protein 4 OS=Homo sapiens OX=9606 GN=CKAP4 PE=1 SV=2 |
| 378 | 28.10 | 5 | 5 | 6405 | 297 | >sp\|P06493\|CDK1_HUMAN Cyclin-dependent kinase 1 OS=Homo sapiens OX=9606 GN=CDK1 PE=1 SV=3 |
| 379 | 28.07 | 4 | 4 | 37032 | 1235 | >sp\|Q00341-2\|VIGLN_HUMAN Isoform 2 of Vigilin OS=Homo sapiens OX=9606 GN=HDLBP |
| 380 | 27.98 | 5 | 5 | 19764 | 210 | >sp\|P82979\|SARNP_HUMAN SAP domain-containing ribonucleoprotein OS=Homo sapiens OX=9606 GN=SARNP PE=1 SV=3 |
| 381 | 27.65 | 3 | 3 | 204 | 353 | >sp\|Q99873-3\|ANM1_HUMAN Isoform 3 of Protein arginine N-methyltransferase 1 OS=Homo sapiens OX=9606 GN=PRMT1 |

| | | | | | | |
|---|---|---|---|---|---|---|
| 382 | 27.64 | 4 | 4 | 31975 | 239 | >sp\|Q9UKD2\|MRT4_HUMAN mRNA turnover protein 4 homolog OS=Homo sapiens OX=9606 GN=MRTO4 PE=1 SV=2 |
| 383 | 27.63 | 9 | 8 | 26929 | 4684 | >sp\|Q15149\|PLEC_HUMAN Plectin OS=Homo sapiens OX=9606 GN=PLEC PE=1 SV=3 |
| 384 | 27.58 | 4 | 4 | 31331 | 819 | >sp\|Q8N1F7\|NUP93_HUMAN Nuclear pore complex protein Nup93 OS=Homo sapiens OX=9606 GN=NUP93 PE=1 SV=2 |
| 385 | 27.58 | 5 | 4 | 7147 | 306 | >sp\|Q99848\|EBP2_HUMAN Probable rRNA-processing protein EBP2 OS=Homo sapiens OX=9606 GN=EBNA1BP2 PE=1 SV=2 |
| 386 | 27.49 | 3 | 3 | 41057 | 233 | >sp\|P55327-7\|TPD52_HUMAN Isoform 7 of Tumor protein D52 OS=Homo sapiens OX=9606 GN=TPD52 |
| 387 | 27.25 | 3 | 3 | 36193 | 148 | >sp\|Q04837\|SSBP_HUMAN Single-stranded DNA-binding protein; mitochondrial OS=Homo sapiens OX=9606 GN=SSBP1 PE=1 SV=1 |
| 388 | 27.20 | 5 | 4 | 29945 | 1135 | >sp\|P27816-6\|MAP4_HUMAN Isoform 6 of Microtubule-associated protein 4 OS=Homo sapiens OX=9606 GN=MAP4 |
| 389 | 27.19 | 4 | 4 | 26028 | 757 | >sp\|Q16891-4\|MIC60_HUMAN Isoform 4 of MICOS complex subunit MIC60 OS=Homo sapiens OX=9606 GN=IMMT |
| 390 | 27.16 | 3 | 3 | 26884 | 228 | >sp\|P22061-2\|PIMT_HUMAN Isoform 2 of Protein-L-isoaspartate(D-aspartate) O-methyltransferase OS=Homo sapiens OX=9606 GN=PCMT1 |
| 391 | 27.12 | 5 | 4 | 29060 | 309 | >sp\|P67775\|PP2AA_HUMAN Serine/threonine-protein phosphatase 2A catalytic subunit alpha isoform OS=Homo sapiens OX=9606 GN=PPP2C |
| 392 | 27.10 | 8 | 6 | 31826 | 724 | >sp\|P49321-4\|NASP_HUMAN Isoform 4 of Nuclear autoantigenic sperm protein OS=Homo sapiens OX=9606 GN=NASP |
| 393 | 27.06 | 7 | 7 | 25835 | 821 | >sp\|Q14566\|MCM6_HUMAN DNA replication licensing factor MCM6 OS=Homo sapiens OX=9606 GN=MCM6 PE=1 SV=1 |
| 394 | 26.96 | 4 | 3 | 22857 | 418 | >sp\|P50454\|SERPH_HUMAN Serpin H1 OS=Homo sapiens OX=9606 GN=SERPINH1 PE=1 SV=2 |
| 395 | 26.93 | 3 | 3 | 37154 | 489 | >sp\|P12081-4\|SYHC_HUMAN Isoform 4 of Histidine--tRNA ligase; cytoplasmic OS=Homo sapiens OX=9606 GN=HARS |
| 396 | 26.90 | 4 | 4 | 38667 | 246 | >sp\|Q00059\|TFAM_HUMAN Transcription factor A; mitochondrial OS=Homo sapiens OX=9606 GN=TFAM PE=1 SV=1 |
| 397 | 26.86 | 9 | 8 | 236 | 320 | >sp\|P08758\|ANXA5_HUMAN Annexin A5 OS=Homo sapiens OX=9606 GN=ANXA5 PE=1 SV=2 |
| 398 | 26.81 | 3 | 3 | 20410 | 436 | >sp\|Q16181-2\|SEPT7_HUMAN Isoform 2 of Septin-7 OS=Homo sapiens OX=9606 GN=SEPT7 |
| 399 | 26.79 | 5 | 5 | 27898 | 392 | >sp\|Q9BY77-2\|PDIP3_HUMAN Isoform 2 of Polymerase delta-interacting protein 3 OS=Homo sapiens OX=9606 GN=POLDIP3 |
| 400 | 26.75 | 4 | 4 | 21312 | 228 | >sp\|P18621-3\|RL17_HUMAN Isoform 3 of 60S ribosomal protein L17 OS=Homo sapiens OX=9606 GN=RPL17 |
| 401 | 26.75 | 3 | 3 | 2631 | 382 | >sp\|Q9UJU6-6\|DBNL_HUMAN Isoform 6 of Drebrin-like protein OS=Homo sapiens OX=9606 GN=DBNL |
| 402 | 26.67 | 5 | 5 | 1436 | 466 | >sp\|O75390\|CISY_HUMAN Citrate synthase; mitochondrial OS=Homo sapiens OX=9606 GN=CS PE=1 SV=2 |
| 403 | 26.66 | 3 | 2 | 21195 | 461 | >sp\|P13489\|RINI_HUMAN Ribonuclease inhibitor OS=Homo sapiens OX=9606 GN=RNH1 PE=1 SV=2 |
| 404 | 26.63 | 6 | 5 | 34717 | 589 | >sp\|Q9NSD9\|SYFB_HUMAN Phenylalanine--tRNA ligase beta subunit OS=Homo sapiens OX=9606 GN=FARSB PE=1 SV=3 |
| 405 | 26.56 | 3 | 3 | 38867 | 689 | >sp\|P52888\|THOP1_HUMAN Thimet oligopeptidase OS=Homo sapiens OX=9606 GN=THOP1 PE=1 SV=2 |
| 406 | 26.52 | 4 | 4 | 7595 | 718 | >sp\|P51659-3\|DHB4_HUMAN Isoform 3 of Peroxisomal multifunctional enzyme type 2 OS=Homo sapiens OX=9606 GN=HSD17B4 |
| 407 | 26.40 | 4 | 4 | 34564 | 376 | >sp\|P61163\|ACTZ_HUMAN Alpha-centractin OS=Homo sapiens OX=9606 GN=ACTR1A PE=1 SV=1 |
| 408 | 26.36 | 5 | 5 | 17694 | 586 | >sp\|P15311\|EZRI_HUMAN Ezrin OS=Homo sapiens OX=9606 GN=EZR PE=1 SV=4 |
| 409 | 26.30 | 2 | 2 | 24186 | 336 | >sp\|Q12904-2\|AIMP1_HUMAN Isoform 2 of Aminoacyl tRNA synthase complex-interacting multifunctional protein 1 OS=Homo sapiens OX= |
| 410 | 26.26 | 6 | 5 | 21329 | 403 | >sp\|P39023\|RL3_HUMAN 60S ribosomal protein L3 OS=Homo sapiens OX=9606 GN=RPL3 PE=1 SV=2 |
| 411 | 26.18 | 3 | 3 | 19844 | 399 | >sp\|Q15393-3\|SF3B3_HUMAN Isoform 3 of Splicing factor 3B subunit 3 OS=Homo sapiens OX=9606 GN=SF3B3 |
| 412 | 26.14 | 5 | 5 | 30638 | 271 | >sp\|Q96FW1\|OTUB1_HUMAN Ubiquitin thioesterase OTUB1 OS=Homo sapiens OX=9606 GN=OTUB1 PE=1 SV=2 |
| 413 | 26.01 | 3 | 3 | 37746 | 858 | >sp\|P45974\|UBP5_HUMAN Ubiquitin carboxyl-terminal hydrolase 5 OS=Homo sapiens OX=9606 GN=USP5 PE=1 SV=2 |
| 414 | 25.98 | 5 | 4 | 11889 | 1038 | >sp\|O95373\|IPO7_HUMAN Importin-7 OS=Homo sapiens OX=9606 GN=IPO7 PE=1 SV=1 |
| 415 | 25.97 | 5 | 4 | 13927 | 371 | >sp\|O75367-3\|H2AY_HUMAN Isoform 3 of Core histone macro-H2A.1 OS=Homo sapiens OX=9606 GN=H2AFY |
| 416 | 25.90 | 5 | 5 | 32688 | 372 | >sp\|Q9UNZ2-5\|NSF1C_HUMAN Isoform 3 of NSFL1 cofactor p47 OS=Homo sapiens OX=9606 GN=NSFL1C |
| 417 | 25.79 | 3 | 3 | 7514 | 148 | >sp\|O60869\|EDF1_HUMAN Endothelial differentiation-related factor 1 OS=Homo sapiens OX=9606 GN=EDF1 PE=1 SV=1 |
| 418 | 25.75 | 2 | 2 | 32646 | 416 | >sp\|P30419-2\|NMT1_HUMAN Isoform Short of Glycylpeptide N-tetradecanoyltransferase 1 OS=Homo sapiens OX=9606 GN=NMT1 |
| 419 | 25.72 | 4 | 4 | 7549 | 403 | >sp\|Q13561-3\|DCTN2_HUMAN Isoform 3 of Dynactin subunit 2 OS=Homo sapiens OX=9606 GN=DCTN2 |
| 420 | 25.69 | 3 | 3 | 24230 | 930 | >sp\|P12814-4\|ACTN1_HUMAN Isoform 4 of Alpha-actinin-1 OS=Homo sapiens OX=9606 GN=ACTN1 |
| 421 | 25.52 | 6 | 5 | 26658 | 1945 | >sp\|P35749-3\|MYH11_HUMAN Isoform 3 of Myosin-11 OS=Homo sapiens OX=9606 GN=MYH11 |
| 422 | 25.47 | 5 | 5 | 21290 | 136 | >sp\|P61353\|RL27_HUMAN 60S ribosomal protein L27 OS=Homo sapiens OX=9606 GN=RPL27 PE=1 SV=2 |
| 423 | 25.41 | 3 | 2 | 24762 | 220 | >sp\|Q9BTT0-3\|AN32E_HUMAN Isoform 3 of Acidic leucine-rich nuclear phosphoprotein 32 family member E OS=Homo sapiens OX=9606 GN= |
| 424 | 25.38 | 2 | 2 | 27195 | 133 | >sp\|Q9Y237-3\|PIN4_HUMAN Isoform 3 of Peptidyl-prolyl cis-trans isomerase NIMA-interacting 4 OS=Homo sapiens OX=9606 GN=PIN4 |
| 425 | 25.37 | 6 | 4 | 14832 | 411 | >sp\|P38919\|IF4A3_HUMAN Eukaryotic initiation factor 4A-III OS=Homo sapiens OX=9606 GN=EIF4A3 PE=1 SV=4 |
| 426 | 25.31 | 4 | 3 | 19615 | 895 | >sp\|Q13435\|SF3B2_HUMAN Splicing factor 3B subunit 2 OS=Homo sapiens OX=9606 GN=SF3B2 PE=1 SV=2 |
| 427 | 25.24 | 5 | 4 | 42254 | 248 | >sp\|P25788-2\|PSA3_HUMAN Isoform 2 of Proteasome subunit alpha type-3 OS=Homo sapiens OX=9606 GN=PSMA3 |
| 428 | 25.23 | 4 | 4 | 28819 | 901 | >sp\|O94906-2\|PRP6_HUMAN Isoform 2 of Pre-mRNA-processing factor 6 OS=Homo sapiens OX=9606 GN=PRPF6 |
| 429 | 25.16 | 4 | 4 | 15590 | 400 | >sp\|P50395-2\|GDIB_HUMAN Isoform 2 of Rab GDP dissociation inhibitor beta OS=Homo sapiens OX=9606 GN=GDI2 |
| 430 | 25.13 | 2 | 2 | 20127 | 321 | >sp\|O14828-2\|SCAM3_HUMAN Isoform 2 of Secretory carrier-associated membrane protein 3 OS=Homo sapiens OX=9606 GN=SCAMP3 |
| 431 | 25.05 | 3 | 3 | 19293 | 326 | >sp\|Q9NQG5\|RPR1B_HUMAN Regulation of nuclear pre-mRNA domain-containing protein 1B OS=Homo sapiens OX=9606 GN=RPRD1B PE=1 SV=1 |
| 432 | 24.98 | 2 | 2 | 6451 | 303 | >sp\|P11802\|CDK4_HUMAN Cyclin-dependent kinase 4 OS=Homo sapiens OX=9606 GN=CDK4 PE=1 SV=2 |
| 433 | 24.86 | 3 | 3 | 31475 | 943 | >sp\|O14974-5\|MYPT1_HUMAN Isoform 5 of Protein phosphatase 1 regulatory subunit 12A OS=Homo sapiens OX=9606 GN=PPP1R12A |
| 434 | 24.85 | 3 | 3 | 32423 | 594 | >sp\|O00567\|NOP56_HUMAN Nucleolar protein 56 OS=Homo sapiens OX=9606 GN=NOP56 PE=1 SV=4 |
| 435 | 24.83 | 4 | 4 | 4314 | 412 | >sp\|P07339\|CATD_HUMAN Cathepsin D OS=Homo sapiens OX=9606 GN=CTSD PE=1 SV=1 |
| 436 | 24.82 | 4 | 3 | 7107 | 695 | >sp\|Q16643-3\|DREB_HUMAN Isoform 3 of Drebrin OS=Homo sapiens OX=9606 GN=DBN1 |
| 437 | 24.80 | 4 | 4 | 34719 | 1176 | >sp\|Q9P2J5\|SYLC_HUMAN Leucine--tRNA ligase; cytoplasmic OS=Homo sapiens OX=9606 GN=LARS PE=1 SV=2 |
| 438 | 24.76 | 3 | 3 | 16997 | 284 | >sp\|P13804-2\|ETFA_HUMAN Isoform 2 of Electron transfer flavoprotein subunit alpha; mitochondrial OS=Homo sapiens OX=9606 GN=ETF |
| 439 | 24.53 | 2 | 2 | 15951 | 404 | >sp\|Q07666-3\|KHDR1_HUMAN Isoform 3 of KH domain-containing; RNA-binding; signal transduction-associated protein 1 OS=Homo sapie |

| | | | | | | |
|---|---|---|---|---|---|---|
| 440 | 24.48 | 2 | 2 | 36776 | 58 | >sp\|Q96IX5\|USMG5_HUMAN Up-regulated during skeletal muscle growth protein 5 OS=Homo sapiens OX=9606 GN=USMG5 PE=1 SV=1 |
| 441 | 24.41 | 7 | 6 | 11605 | 876 | >sp\|Q14974\|IMB1_HUMAN Importin subunit beta-1 OS=Homo sapiens OX=9606 GN=KPNB1 PE=1 SV=2 |
| 442 | 24.38 | 2 | 2 | 29405 | 317 | >sp\|Q15435-2\|PP1R7_HUMAN Isoform 2 of Protein phosphatase 1 regulatory subunit 7 OS=Homo sapiens OX=9606 GN=PPP1R7 |
| 443 | 24.32 | 5 | 4 | 19239 | 165 | >sp\|P46783\|RS10_HUMAN 40S ribosomal protein S10 OS=Homo sapiens OX=9606 GN=RPS10 PE=1 SV=1 |
| 444 | 24.32 | 4 | 4 | 9133 | 185 | >sp\|Q9HB71-3\|CYBP_HUMAN Isoform 3 of Calcyclin-binding protein OS=Homo sapiens OX=9606 GN=CACYBP |
| 445 | 24.22 | 7 | 6 | 712 | 303 | >sp\|Q9UNE7\|CHIP_HUMAN E3 ubiquitin-protein ligase CHIP OS=Homo sapiens OX=9606 GN=STUB1 PE=1 SV=2 |
| 446 | 24.14 | 3 | 3 | 11218 | 164 | >sp\|P33316-2\|DUT_HUMAN Isoform 2 of Deoxyuridine 5'-triphosphate nucleotidohydrolase; mitochondrial OS=Homo sapiens OX=9606 GN= |
| 447 | 24.14 | 4 | 4 | 21044 | 1173 | >sp\|Q92878-3\|RAD50_HUMAN Isoform 3 of DNA repair protein RAD50 OS=Homo sapiens OX=9606 GN=RAD50 |
| 448 | 24.11 | 5 | 4 | 36666 | 1071 | >sp\|O14980\|XPO1_HUMAN Exportin-1 OS=Homo sapiens OX=9606 GN=XPO1 PE=1 SV=1 |
| 449 | 24.09 | 3 | 3 | 20206 | 567 | >sp\|Q9UHD8-7\|SEPT9_HUMAN Isoform 7 of Septin-9 OS=Homo sapiens OX=9606 GN=SEPT9 |
| 450 | 24.03 | 2 | 2 | 11381 | 90 | >sp\|P52926-6\|HMGA2_HUMAN Isoform 6 of High mobility group protein HMGI-C OS=Homo sapiens OX=9606 GN=HMGA2 |
| 451 | 23.90 | 5 | 4 | 17440 | 445 | >sp\|P60228\|EIF3E_HUMAN Eukaryotic translation initiation factor 3 subunit E OS=Homo sapiens OX=9606 GN=EIF3E PE=1 SV=1 |
| 452 | 23.79 | 3 | 3 | 34734 | 900 | >sp\|P56192\|SYMC_HUMAN Methionine--tRNA ligase; cytoplasmic OS=Homo sapiens OX=9606 GN=MARS PE=1 SV=2 |
| 453 | 23.71 | 4 | 3 | 24831 | 288 | >sp\|O95433-2\|AHSA1_HUMAN Isoform 2 of Activator of 90 kDa heat shock protein ATPase homolog 1 OS=Homo sapiens OX=9606 GN=AHSA1 |
| 454 | 23.66 | 4 | 4 | 42169 | 440 | >sp\|P62191\|PRS4_HUMAN 26S proteasome regulatory subunit 4 OS=Homo sapiens OX=9606 GN=PSMC1 PE=1 SV=1 |
| 455 | 23.61 | 3 | 3 | 14493 | 472 | >sp\|P41091\|IF2G_HUMAN Eukaryotic translation initiation factor 2 subunit 3 OS=Homo sapiens OX=9606 GN=EIF2S3 PE=1 SV=3 |
| 456 | 23.57 | 3 | 2 | 14105 | 1293 | >sp\|Q6Y7W6-5\|GGYF2_HUMAN Isoform 4 of GRB10-interacting GYF protein 2 OS=Homo sapiens OX=9606 GN=GIGYF2 |
| 457 | 23.53 | 2 | 2 | 21694 | 305 | >sp\|Q13151\|ROA0_HUMAN Heterogeneous nuclear ribonucleoprotein A0 OS=Homo sapiens OX=9606 GN=HNRNPA0 PE=1 SV=1 |
| 458 | 23.51 | 2 | 2 | 21699 | 285 | >sp\|Q99729-3\|ROAA_HUMAN Isoform 3 of Heterogeneous nuclear ribonucleoprotein A/B OS=Homo sapiens OX=9606 GN=HNRNPAB |
| 459 | 23.44 | 4 | 4 | 7148 | 763 | >sp\|P40939\|ECHA_HUMAN Trifunctional enzyme subunit alpha; mitochondrial OS=Homo sapiens OX=9606 GN=HADHA PE=1 SV=2 |
| 460 | 23.32 | 5 | 5 | 14431 | 1220 | >sp\|O60841\|IF2P_HUMAN Eukaryotic translation initiation factor 5B OS=Homo sapiens OX=9606 GN=EIF5B PE=1 SV=4 |
| 461 | 23.20 | 2 | 2 | 18064 | 368 | >sp\|P55209-2\|NP1L1_HUMAN Isoform 2 of Nucleosome assembly protein 1-like 1 OS=Homo sapiens OX=9606 GN=NAP1L1 |
| 462 | 23.16 | 4 | 4 | 6110 | 923 | >sp\|Q8N163-2\|CCAR2_HUMAN Isoform 2 of Cell cycle and apoptosis regulator protein 2 OS=Homo sapiens OX=9606 GN=CCAR2 |
| 463 | 23.11 | 2 | 2 | 2841 | 269 | >sp\|P35613-2\|BASI_HUMAN Isoform 2 of Basigin OS=Homo sapiens OX=9606 GN=BSG |
| 464 | 23.10 | 3 | 3 | 16119 | 196 | >sp\|P30085\|KCY_HUMAN UMP-CMP kinase OS=Homo sapiens OX=9606 GN=CMPK1 PE=1 SV=3 |
| 465 | 23.08 | 5 | 5 | 21306 | 204 | >sp\|P61313\|RL15_HUMAN 60S ribosomal protein L15 OS=Homo sapiens OX=9606 GN=RPL15 PE=1 SV=2 |
| 466 | 22.97 | 3 | 3 | 3560 | 661 | >sp\|P08195-4\|4F2_HUMAN Isoform 4 of 4F2 cell-surface antigen heavy chain OS=Homo sapiens OX=9606 GN=SLC3A2 |
| 467 | 22.94 | 5 | 5 | 41617 | 200 | >sp\|P61086\|UBE2K_HUMAN Ubiquitin-conjugating enzyme E2 K OS=Homo sapiens OX=9606 GN=UBE2K PE=1 SV=3 |
| 468 | 22.89 | 4 | 4 | 6195 | 802 | >sp\|Q99459\|CDC5L_HUMAN Cell division cycle 5-like protein OS=Homo sapiens OX=9606 GN=CDC5L PE=1 SV=2 |
| 469 | 22.86 | 3 | 2 | 26972 | 562 | >sp\|P36871\|PGM1_HUMAN Phosphoglucomutase-1 OS=Homo sapiens OX=9606 GN=PGM1 PE=1 SV=3 |
| 470 | 22.67 | 5 | 5 | 21004 | 453 | >sp\|P22695\|QCR2_HUMAN Cytochrome b-c1 complex subunit 2; mitochondrial OS=Homo sapiens OX=9606 GN=UQCRC2 PE=1 SV=3 |
| 471 | 22.66 | 2 | 2 | 18299 | 227 | >sp\|Q9NX63\|MIC19_HUMAN MICOS complex subunit MIC19 OS=Homo sapiens OX=9606 GN=CHCHD3 PE=1 SV=1 |
| 472 | 22.61 | 5 | 4 | 35745 | 992 | >sp\|P49588-2\|SYAC_HUMAN Isoform 2 of Alanine--tRNA ligase; cytoplasmic OS=Homo sapiens OX=9606 GN=AARS |
| 473 | 22.59 | 3 | 3 | 39068 | 172 | >sp\|P13693\|TCTP_HUMAN Translationally-controlled tumor protein OS=Homo sapiens OX=9606 GN=TPT1 PE=1 SV=1 |
| 474 | 22.50 | 2 | 2 | 35007 | 446 | >sp\|Q9BUF5\|TBB6_HUMAN Tubulin beta-6 chain OS=Homo sapiens OX=9606 GN=TUBB6 PE=1 SV=1 |
| 475 | 22.44 | 2 | 2 | 28430 | 233 | >sp\|Q9BVG4\|PBDC1_HUMAN Protein PBDC1 OS=Homo sapiens OX=9606 GN=PBDC1 PE=1 SV=1 |
| 476 | 22.36 | 4 | 4 | 40089 | 342 | >sp\|P16989-3\|YBOX3_HUMAN Isoform 3 of Y-box-binding protein 3 OS=Homo sapiens OX=9606 GN=YBX3 |
| 477 | 22.29 | 2 | 2 | 30760 | 634 | >sp\|Q86YP4-3\|P66A_HUMAN Isoform 3 of Transcriptional repressor p66-alpha OS=Homo sapiens OX=9606 GN=GATAD2A |
| 478 | 22.24 | 2 | 2 | 24480 | 237 | >sp\|O95831-6\|AIFM1_HUMAN Isoform 6 of Apoptosis-inducing factor 1; mitochondrial OS=Homo sapiens OX=9606 GN=AIFM1 |
| 479 | 22.17 | 3 | 2 | 42050 | 593 | >sp\|Q06124-2\|PTN11_HUMAN Isoform 2 of Tyrosine-protein phosphatase non-receptor type 11 OS=Homo sapiens OX=9606 GN=PTPN11 |
| 480 | 21.95 | 5 | 5 | 36130 | 118 | >sp\|P62316\|SMD2_HUMAN Small nuclear ribonucleoprotein Sm D2 OS=Homo sapiens OX=9606 GN=SNRPD2 PE=1 SV=1 |
| 481 | 21.91 | 5 | 3 | 22227 | 151 | >sp\|P62263\|RS14_HUMAN 40S ribosomal protein S14 OS=Homo sapiens OX=9606 GN=RPS14 PE=1 SV=3 |
| 482 | 21.89 | 4 | 3 | 17314 | 637 | >sp\|P15170-3\|ERF3A_HUMAN Isoform 3 of Eukaryotic peptide chain release factor GTP-binding subunit ERF3A OS=Homo sapiens OX=9606 |
| 483 | 21.73 | 2 | 2 | 3069 | 298 | >sp\|P36542\|ATPG_HUMAN ATP synthase subunit gamma; mitochondrial OS=Homo sapiens OX=9606 GN=ATP5F1C PE=1 SV=1 |
| 484 | 21.73 | 2 | 2 | 22565 | 102 | >sp\|P20962\|PTMS_HUMAN Parathymosin OS=Homo sapiens OX=9606 GN=PTMS PE=1 SV=2 |
| 485 | 21.72 | 3 | 3 | 19757 | 198 | >sp\|Q9NR31\|SAR1A_HUMAN GTP-binding protein SAR1a OS=Homo sapiens OX=9606 GN=SAR1A PE=1 SV=1 |
| 486 | 21.66 | 2 | 2 | 36034 | 173 | >sp\|P51571\|SSRD_HUMAN Translocon-associated protein subunit delta OS=Homo sapiens OX=9606 GN=SSR4 PE=1 SV=1 |
| 487 | 21.64 | 3 | 3 | 33604 | 211 | >sp\|Q8WZA0-2\|LZIC_HUMAN Isoform 2 of Protein LZIC OS=Homo sapiens OX=9606 GN=LZIC |
| 488 | 21.63 | 4 | 4 | 19566 | 519 | >sp\|P31040-3\|SDHA_HUMAN Isoform 3 of Succinate dehydrogenase [ubiquinone] flavoprotein subunit; mitochondrial OS=Homo sapiens O |
| 489 | 21.63 | 4 | 4 | 29244 | 198 | >sp\|P32119\|PRDX2_HUMAN Peroxiredoxin-2 OS=Homo sapiens OX=9606 GN=PRDX2 PE=1 SV=5 |
| 490 | 21.60 | 5 | 5 | 16432 | 400 | >sp\|P08727\|K1C19_HUMAN Keratin; type I cytoskeletal 19 OS=Homo sapiens OX=9606 GN=KRT19 PE=1 SV=4 |
| 491 | 21.49 | 2 | 2 | 5553 | 202 | >sp\|Q9Y3Y2-4\|CHTOP_HUMAN Isoform 3 of Chromatin target of PRMT1 protein OS=Homo sapiens OX=9606 GN=CHTOP |
| 492 | 21.47 | 2 | 2 | 256 | 537 | >sp\|Q9UJX3-2\|APC7_HUMAN Isoform 2 of Anaphase-promoting complex subunit 7 OS=Homo sapiens OX=9606 GN=ANAPC7 |

<table>
<tr><td>493</td><td>21.46</td><td>4</td><td>3</td><td>10959</td><td>254</td><td>>sp|P50402|EMD_HUMAN Emerin OS=Homo sapiens OX=9606 GN=EMD PE=1 SV=1</td></tr>
<tr><td>494</td><td>21.40</td><td>6</td><td>6</td><td>12623</td><td>747</td><td>>sp|Q1KMD3|HNRL2_HUMAN Heterogeneous nuclear ribonucleoprotein U-like protein 2 OS=Homo sapiens OX=9606 GN=HNRNPUL2 PE=1 SV=1</td></tr>
<tr><td>495</td><td>21.31</td><td>2</td><td>2</td><td>22648</td><td>265</td><td>>sp|P61289-3|PSME3_HUMAN Isoform 3 of Proteasome activator complex subunit 3 OS=Homo sapiens OX=9606 GN=PSME3</td></tr>
<tr><td>496</td><td>21.31</td><td>5</td><td>4</td><td>19250</td><td>158</td><td>>sp|P62280|RS11_HUMAN 40S ribosomal protein S11 OS=Homo sapiens OX=9606 GN=RPS11 PE=1 SV=3</td></tr>
<tr><td>497</td><td>21.29</td><td>5</td><td>4</td><td>32087</td><td>1068</td><td>>sp|P52701-2|MSH6_HUMAN Isoform GTBP-alt of DNA mismatch repair protein Msh6 OS=Homo sapiens OX=9606 GN=MSH6</td></tr>
<tr><td>498</td><td>21.27</td><td>3</td><td>2</td><td>37977</td><td>158</td><td>>sp|P63279|UBC9_HUMAN SUMO-conjugating enzyme UBC9 OS=Homo sapiens OX=9606 GN=UBE2I PE=1 SV=1</td></tr>
<tr><td>499</td><td>21.26</td><td>3</td><td>3</td><td>42093</td><td>456</td><td>>sp|P30520|PURA2_HUMAN Adenylosuccinate synthetase isozyme 2 OS=Homo sapiens OX=9606 GN=ADSS PE=1 SV=3</td></tr>
<tr><td>500</td><td>21.23</td><td>6</td><td>6</td><td>41926</td><td>2225</td><td>>sp|P27708|PYR1_HUMAN CAD protein OS=Homo sapiens OX=9606 GN=CAD PE=1 SV=3</td></tr>
<tr><td>501</td><td>21.20</td><td>4</td><td>4</td><td>14497</td><td>869</td><td>>sp|P78344-2|IF4G2_HUMAN Isoform 2 of Eukaryotic translation initiation factor 4 gamma 2 OS=Homo sapiens OX=9606 GN=EIF4G2</td></tr>
<tr><td>502</td><td>21.18</td><td>5</td><td>4</td><td>32755</td><td>1332</td><td>>sp|O75694-2|NU155_HUMAN Isoform 2 of Nuclear pore complex protein Nup155 OS=Homo sapiens OX=9606 GN=NUP155</td></tr>
<tr><td>503</td><td>21.18</td><td>6</td><td>6</td><td>1072</td><td>75</td><td>>sp|P09669|COX6C_HUMAN Cytochrome c oxidase subunit 6C OS=Homo sapiens OX=9606 GN=COX6C PE=1 SV=2</td></tr>
<tr><td>504</td><td>21.11</td><td>2</td><td>2</td><td>1424</td><td>313</td><td>>sp|Q9UHD1-2|CHRD1_HUMAN Isoform 2 of Cysteine and histidine-rich domain-containing protein 1 OS=Homo sapiens OX=9606 GN=CHORDC</td></tr>
<tr><td>505</td><td>20.97</td><td>2</td><td>2</td><td>3835</td><td>387</td><td>>sp|O96019-2|ACL6A_HUMAN Isoform 2 of Actin-like protein 6A OS=Homo sapiens OX=9606 GN=ACTL6A</td></tr>
<tr><td>506</td><td>20.89</td><td>4</td><td>4</td><td>1892</td><td>527</td><td>>sp|Q9ULV4-3|COR1C_HUMAN Isoform 3 of Coronin-1C OS=Homo sapiens OX=9606 GN=CORO1C</td></tr>
<tr><td>507</td><td>20.76</td><td>4</td><td>4</td><td>37719</td><td>183</td><td>>sp|P61081|UBC12_HUMAN NEDD8-conjugating enzyme Ubc12 OS=Homo sapiens OX=9606 GN=UBE2M PE=1 SV=1</td></tr>
<tr><td>508</td><td>20.68</td><td>4</td><td>2</td><td>37667</td><td>229</td><td>>sp|P0CG47|UBB_HUMAN Polyubiquitin-B OS=Homo sapiens OX=9606 GN=UBB PE=1 SV=1</td></tr>
<tr><td>509</td><td>20.66</td><td>5</td><td>5</td><td>20019</td><td>765</td><td>>sp|Q15436|SC23A_HUMAN Protein transport protein Sec23A OS=Homo sapiens OX=9606 GN=SEC23A PE=1 SV=2</td></tr>
<tr><td>510</td><td>20.61</td><td>6</td><td>6</td><td>12446</td><td>215</td><td>>sp|P09429|HMGB1_HUMAN High mobility group protein B1 OS=Homo sapiens OX=9606 GN=HMGB1 PE=1 SV=3</td></tr>
<tr><td>511</td><td>20.59</td><td>3</td><td>2</td><td>12592</td><td>205</td><td>>sp|P04792|HSPB1_HUMAN Heat shock protein beta-1 OS=Homo sapiens OX=9606 GN=HSPB1 PE=1 SV=2</td></tr>
<tr><td>512</td><td>20.54</td><td>2</td><td>2</td><td>34157</td><td>425</td><td>>sp|P11310-2|ACADM_HUMAN Isoform 2 of Medium-chain specific acyl-CoA dehydrogenase; mitochondrial OS=Homo sapiens OX=9606 GN=AC</td></tr>
<tr><td>513</td><td>20.51</td><td>7</td><td>5</td><td>25975</td><td>847</td><td>>sp|P43243|MATR3_HUMAN Matrin-3 OS=Homo sapiens OX=9606 GN=MATR3 PE=1 SV=2</td></tr>
<tr><td>514</td><td>20.43</td><td>3</td><td>3</td><td>2611</td><td>483</td><td>>sp|P26196|DDX6_HUMAN Probable ATP-dependent RNA helicase DDX6 OS=Homo sapiens OX=9606 GN=DDX6 PE=1 SV=2</td></tr>
<tr><td>515</td><td>20.43</td><td>4</td><td>3</td><td>10744</td><td>104</td><td>>sp|P84090|ERH_HUMAN Enhancer of rudimentary homolog OS=Homo sapiens OX=9606 GN=ERH PE=1 SV=1</td></tr>
<tr><td>516</td><td>20.35</td><td>3</td><td>3</td><td>8977</td><td>299</td><td>>sp|Q6FI81-3|CPIN1_HUMAN Isoform 3 of Anamorsin OS=Homo sapiens OX=9606 GN=CIAPIN1</td></tr>
<tr><td>517</td><td>20.24</td><td>3</td><td>3</td><td>28386</td><td>697</td><td>>sp|Q32P28-4|P3H1_HUMAN Isoform 4 of Prolyl 3-hydroxylase 1 OS=Homo sapiens OX=9606 GN=P3H1</td></tr>
<tr><td>518</td><td>20.23</td><td>2</td><td>2</td><td>11415</td><td>458</td><td>>sp|Q92769-3|HDAC2_HUMAN Isoform 2 of Histone deacetylase 2 OS=Homo sapiens OX=9606 GN=HDAC2</td></tr>
<tr><td>519</td><td>20.22</td><td>5</td><td>5</td><td>21918</td><td>130</td><td>>sp|P62244|RS15A_HUMAN 40S ribosomal protein S15a OS=Homo sapiens OX=9606 GN=RPS15A PE=1 SV=2</td></tr>
<tr><td>520</td><td>20.22</td><td>3</td><td>3</td><td>35714</td><td>339</td><td>>sp|Q9P289-2|STK26_HUMAN Isoform 2 of Serine/threonine-protein kinase 26 OS=Homo sapiens OX=9606 GN=STK26</td></tr>
<tr><td>521</td><td>20.17</td><td>3</td><td>3</td><td>23951</td><td>513</td><td>>sp|Q2TAY7|SMU1_HUMAN WD40 repeat-containing protein SMU1 OS=Homo sapiens OX=9606 GN=SMU1 PE=1 SV=2</td></tr>
<tr><td>522</td><td>20.15</td><td>2</td><td>2</td><td>16217</td><td>223</td><td>>sp|P27144|KAD4_HUMAN Adenylate kinase 4; mitochondrial OS=Homo sapiens OX=9606 GN=AK4 PE=1 SV=1</td></tr>
<tr><td>523</td><td>20.10</td><td>7</td><td>6</td><td>6904</td><td>331</td><td>>sp|P31689-2|DNJA1_HUMAN Isoform 2 of DnaJ homolog subfamily A member 1 OS=Homo sapiens OX=9606 GN=DNAJA1</td></tr>
<tr><td>524</td><td>20.03</td><td>2</td><td>2</td><td>22228</td><td>59</td><td>>sp|P62861|RS30_HUMAN 40S ribosomal protein S30 OS=Homo sapiens OX=9606 GN=FAU PE=1 SV=1</td></tr>
<tr><td>525</td><td>19.99</td><td>5</td><td>4</td><td>21313</td><td>176</td><td>>sp|Q02543|RL18A_HUMAN 60S ribosomal protein L18a OS=Homo sapiens OX=9606 GN=RPL18A PE=1 SV=2</td></tr>
<tr><td>526</td><td>19.96</td><td>2</td><td>2</td><td>22734</td><td>333</td><td>>sp|O75475-2|PSIP1_HUMAN Isoform 2 of PC4 and SFRS1-interacting protein OS=Homo sapiens OX=9606 GN=PSIP1</td></tr>
<tr><td>527</td><td>19.84</td><td>4</td><td>4</td><td>33566</td><td>583</td><td>>sp|Q8N1G4|LRC47_HUMAN Leucine-rich repeat-containing protein 47 OS=Homo sapiens OX=9606 GN=LRRC47 PE=1 SV=1</td></tr>
<tr><td>528</td><td>19.71</td><td>4</td><td>4</td><td>37796</td><td>727</td><td>>sp|P17480-2|UBF1_HUMAN Isoform UBF2 of Nucleolar transcription factor 1 OS=Homo sapiens OX=9606 GN=UBTF</td></tr>
<tr><td>529</td><td>19.61</td><td>4</td><td>4</td><td>29415</td><td>546</td><td>>sp|O15355|PPM1G_HUMAN Protein phosphatase 1G OS=Homo sapiens OX=9606 GN=PPM1G PE=1 SV=1</td></tr>
<tr><td>530</td><td>19.54</td><td>3</td><td>3</td><td>17441</td><td>352</td><td>>sp|O15372|EIF3H_HUMAN Eukaryotic translation initiation factor 3 subunit H OS=Homo sapiens OX=9606 GN=EIF3H PE=1 SV=1</td></tr>
<tr><td>531</td><td>19.49</td><td>2</td><td>2</td><td>22461</td><td>239</td><td>>sp|P28072|PSB6_HUMAN Proteasome subunit beta type-6 OS=Homo sapiens OX=9606 GN=PSMB6 PE=1 SV=4</td></tr>
<tr><td>532</td><td>19.44</td><td>4</td><td>4</td><td>14503</td><td>228</td><td>>sp|Q15056-2|IF4H_HUMAN Isoform Short of Eukaryotic translation initiation factor 4H OS=Homo sapiens OX=9606 GN=EIF4H</td></tr>
<tr><td>533</td><td>19.41</td><td>5</td><td>5</td><td>25810</td><td>1332</td><td>>sp|Q9BQG0-2|MBB1A_HUMAN Isoform 2 of Myb-binding protein 1A OS=Homo sapiens OX=9606 GN=MYBBP1A</td></tr>
<tr><td>534</td><td>19.36</td><td>3</td><td>3</td><td>9631</td><td>467</td><td>>sp|P07954-2|FUMH_HUMAN Isoform Cytoplasmic of Fumarate hydratase; mitochondrial OS=Homo sapiens OX=9606 GN=FH</td></tr>
<tr><td>535</td><td>19.34</td><td>4</td><td>3</td><td>42160</td><td>504</td><td>>sp|Q9UMS4|PRP19_HUMAN Pre-mRNA-processing factor 19 OS=Homo sapiens OX=9606 GN=PRPF19 PE=1 SV=1</td></tr>
<tr><td>536</td><td>19.34</td><td>3</td><td>3</td><td>22415</td><td>335</td><td>>sp|Q14257-2|RCN2_HUMAN Isoform 2 of Reticulocalbin-2 OS=Homo sapiens OX=9606 GN=RCN2</td></tr>
<tr><td>537</td><td>19.30</td><td>4</td><td>4</td><td>19839</td><td>501</td><td>>sp|Q12874|SF3A3_HUMAN Splicing factor 3A subunit 3 OS=Homo sapiens OX=9606 GN=SF3A3 PE=1 SV=1</td></tr>
<tr><td>538</td><td>19.28</td><td>4</td><td>3</td><td>35216</td><td>1612</td><td>>sp|P11388-4|TOP2A_HUMAN Isoform 4 of DNA topoisomerase 2-alpha OS=Homo sapiens OX=9606 GN=TOP2A</td></tr>
<tr><td>539</td><td>19.28</td><td>2</td><td>2</td><td>41621</td><td>309</td><td>>sp|O14562|UBFD1_HUMAN Ubiquitin domain-containing protein UBFD1 OS=Homo sapiens OX=9606 GN=UBFD1 PE=1 SV=2</td></tr>
<tr><td>540</td><td>19.23</td><td>3</td><td>3</td><td>15080</td><td>504</td><td>>sp|Q9NPH2-3|INO1_HUMAN Isoform 3 of Inositol-3-phosphate synthase 1 OS=Homo sapiens OX=9606 GN=ISYNA1</td></tr>
<tr><td>541</td><td>19.23</td><td>2</td><td>2</td><td>32763</td><td>440</td><td>>sp|Q9UKX7-2|NUP50_HUMAN Isoform 2 of Nuclear pore complex protein Nup50 OS=Homo sapiens OX=9606 GN=NUP50</td></tr>
<tr><td>542</td><td>19.21</td><td>2</td><td>2</td><td>14994</td><td>566</td><td>>sp|P20839-7|IMDH1_HUMAN Isoform 7 of Inosine-5'-monophosphate dehydrogenase 1 OS=Homo sapiens OX=9606 GN=IMPDH1</td></tr>
<tr><td>543</td><td>19.21</td><td>3</td><td>2</td><td>21801</td><td>215</td><td>>sp|P50914|RL14_HUMAN 60S ribosomal protein L14 OS=Homo sapiens OX=9606 GN=RPL14 PE=1 SV=4</td></tr>
<tr><td>544</td><td>19.20</td><td>2</td><td>2</td><td>13126</td><td>392</td><td>>sp|Q9Y383|LC7L2_HUMAN Putative RNA-binding protein Luc7-like 2 OS=Homo sapiens OX=9606 GN=LUC7L2 PE=1 SV=2</td></tr>
<tr><td>545</td><td>19.18</td><td>3</td><td>3</td><td>21377</td><td>177</td><td>>sp|P62913-2|RL11_HUMAN Isoform 2 of 60S ribosomal protein L11 OS=Homo sapiens OX=9606 GN=RPL11</td></tr>
<tr><td>546</td><td>19.17</td><td>5</td><td>5</td><td>18198</td><td>169</td><td>>sp|Q04760-2|LGUL_HUMAN Isoform 2 of Lactoylglutathione lyase OS=Homo sapiens OX=9606 GN=GLO1</td></tr>
<tr><td>547</td><td>19.15</td><td>4</td><td>4</td><td>30538</td><td>660</td><td>>sp|Q13310-3|PABP4_HUMAN Isoform 3 of Polyadenylate-binding protein 4 OS=Homo sapiens OX=9606 GN=PABPC4</td></tr>
<tr><td>548</td><td>19.12</td><td>3</td><td>3</td><td>35798</td><td>298</td><td>>sp|Q13148-4|TADBP_HUMAN Isoform 2 of TAR DNA-binding protein 43 OS=Homo sapiens OX=9606 GN=TARDBP</td></tr>
<tr><td>549</td><td>19.09</td><td>3</td><td>3</td><td>26655</td><td>1985</td><td>>sp|P35580-5|MYH10_HUMAN Isoform 5 of Myosin-10 OS=Homo sapiens OX=9606 GN=MYH10</td></tr>
<tr><td>550</td><td>19.09</td><td>2</td><td>2</td><td>21531</td><td>198</td><td>>sp|P52815|RM12_HUMAN 39S ribosomal protein L12; mitochondrial OS=Homo sapiens OX=9606 GN=MRPL12 PE=1 SV=2</td></tr>
</table>

| | | | | | | |
|---|---|---|---|---|---|---|
| 551 | 19.02 | 3 | 2 | 11677 | 390 | >sp\|Q12905\|ILF2_HUMAN Interleukin enhancer-binding factor 2 OS=Homo sapiens OX=9606 GN=ILF2 PE=1 SV=2 |
| 552 | 19.00 | 3 | 2 | 22727 | 234 | >sp\|P25787\|PSA2_HUMAN Proteasome subunit alpha type-2 OS=Homo sapiens OX=9606 GN=PSMA2 PE=1 SV=2 |
| 553 | 18.98 | 2 | 2 | 1260 | 391 | >sp\|Q96KP4-2\|CNDP2_HUMAN Isoform 2 of Cytosolic non-specific dipeptidase OS=Homo sapiens OX=9606 GN=CNDP2 |
| 554 | 18.98 | 3 | 3 | 7851 | 412 | >sp\|O60884\|DNJA2_HUMAN DnaJ homolog subfamily A member 2 OS=Homo sapiens OX=9606 GN=DNAJA2 PE=1 SV=1 |
| 555 | 18.94 | 2 | 2 | 1346 | 423 | >sp\|P48444-2\|COPD_HUMAN Isoform 2 of Coatomer subunit delta OS=Homo sapiens OX=9606 GN=ARCN1 |
| 556 | 18.87 | 3 | 3 | 40854 | 311 | >sp\|P53007\|TXTP_HUMAN Tricarboxylate transport protein; mitochondrial OS=Homo sapiens OX=9606 GN=SLC25A1 PE=1 SV=2 |
| 557 | 18.84 | 2 | 2 | 40218 | 188 | >sp\|P62995-3\|TRA2B_HUMAN Isoform 3 of Transformer-2 protein homolog beta OS=Homo sapiens OX=9606 GN=TRA2B |
| 558 | 18.83 | 3 | 3 | 28549 | 261 | >sp\|P12004\|PCNA_HUMAN Proliferating cell nuclear antigen OS=Homo sapiens OX=9606 GN=PCNA PE=1 SV=1 |
| 559 | 18.82 | 3 | 3 | 35210 | 309 | >sp\|Q15785\|TOM34_HUMAN Mitochondrial import receptor subunit TOM34 OS=Homo sapiens OX=9606 GN=TOMM34 PE=1 SV=2 |
| 560 | 18.81 | 3 | 3 | 24192 | 330 | >sp\|O00170\|AIP_HUMAN AH receptor-interacting protein OS=Homo sapiens OX=9606 GN=AIP PE=1 SV=2 |
| 561 | 18.78 | 3 | 3 | 437 | 1042 | >sp\|P16615\|AT2A2_HUMAN Sarcoplasmic/endoplasmic reticulum calcium ATPase 2 OS=Homo sapiens OX=9606 GN=ATP2A2 PE=1 SV=1 |
| 562 | 18.75 | 4 | 4 | 37749 | 1086 | >sp\|Q93009-3\|UBP7_HUMAN Isoform 3 of Ubiquitin carboxyl-terminal hydrolase 7 OS=Homo sapiens OX=9606 GN=USP7 |
| 563 | 18.61 | 4 | 4 | 29006 | 261 | >sp\|P25789\|PSA4_HUMAN Proteasome subunit alpha type-4 OS=Homo sapiens OX=9606 GN=PSMA4 PE=1 SV=1 |
| 564 | 18.53 | 2 | 2 | 2924 | 333 | >sp\|Q92843-2\|B2CL2_HUMAN Isoform 3 of Bcl-2-like protein 2 OS=Homo sapiens OX=9606 GN=BCL2L2 |
| 565 | 18.37 | 2 | 2 | 40291 | 243 | >sp\|Q9BVC6\|TM109_HUMAN Transmembrane protein 109 OS=Homo sapiens OX=9606 GN=TMEM109 PE=1 SV=1 |
| 566 | 18.36 | 3 | 3 | 41223 | 1204 | >sp\|Q9HAV4\|XPO5_HUMAN Exportin-5 OS=Homo sapiens OX=9606 GN=XPO5 PE=1 SV=1 |
| 567 | 18.33 | 2 | 2 | 34935 | 71 | >sp\|P61956-2\|SUMO2_HUMAN Isoform 2 of Small ubiquitin-related modifier 2 OS=Homo sapiens OX=9606 GN=SUMO2 |
| 568 | 18.31 | 2 | 2 | 11533 | 144 | >sp\|P47813\|IF1AX_HUMAN Eukaryotic translation initiation factor 1A; X-chromosomal OS=Homo sapiens OX=9606 GN=EIF1AX PE=1 SV=2 |
| 569 | 18.23 | 4 | 4 | 40868 | 608 | >sp\|O94826\|TOM70_HUMAN Mitochondrial import receptor subunit TOM70 OS=Homo sapiens OX=9606 GN=TOMM70 PE=1 SV=1 |
| 570 | 18.10 | 2 | 2 | 21742 | 166 | >sp\|Q13405\|RM49_HUMAN 39S ribosomal protein L49; mitochondrial OS=Homo sapiens OX=9606 GN=MRPL49 PE=1 SV=1 |
| 571 | 18.08 | 5 | 5 | 1068 | 1233 | >sp\|P53621-2\|COPA_HUMAN Isoform 2 of Coatomer subunit alpha OS=Homo sapiens OX=9606 GN=COPA |
| 572 | 18.07 | 3 | 3 | 21880 | 83 | >sp\|P63220\|RS21_HUMAN 40S ribosomal protein S21 OS=Homo sapiens OX=9606 GN=RPS21 PE=1 SV=1 |
| 573 | 18.02 | 2 | 2 | 10940 | 112 | >sp\|Q15369\|ELOC_HUMAN Elongin-C OS=Homo sapiens OX=9606 GN=ELOC PE=1 SV=1 |
| 574 | 18.01 | 3 | 2 | 16994 | 1114 | >sp\|Q9BSJ8-2\|ESYT1_HUMAN Isoform 2 of Extended synaptotagmin-1 OS=Homo sapiens OX=9606 GN=ESYT1 |
| 575 | 17.99 | 3 | 3 | 39613 | 572 | >sp\|Q15942\|ZYX_HUMAN Zyxin OS=Homo sapiens OX=9606 GN=ZYX PE=1 SV=1 |
| 576 | 17.97 | 4 | 3 | 18062 | 301 | >sp\|Q9Y314\|NOSIP_HUMAN Nitric oxide synthase-interacting protein OS=Homo sapiens OX=9606 GN=NOSIP PE=1 SV=1 |
| 577 | 17.96 | 2 | 2 | 28810 | 170 | >sp\|P30044-3\|PRDX5_HUMAN Isoform 3 of Peroxiredoxin-5; mitochondrial OS=Homo sapiens OX=9606 GN=PRDX5 |
| 578 | 17.96 | 3 | 2 | 5363 | 223 | >sp\|Q96CT7\|CC124_HUMAN Coiled-coil domain-containing protein 124 OS=Homo sapiens OX=9606 GN=CCDC124 PE=1 SV=1 |
| 579 | 17.94 | 2 | 2 | 23654 | 154 | >sp\|P00441\|SODC_HUMAN Superoxide dismutase [Cu-Zn] OS=Homo sapiens OX=9606 GN=SOD1 PE=1 SV=2 |
| 580 | 17.92 | 2 | 2 | 12448 | 90 | >sp\|P05204\|HMGN2_HUMAN Non-histone chromosomal protein HMG-17 OS=Homo sapiens OX=9606 GN=HMGN2 PE=1 SV=3 |
| 581 | 17.90 | 3 | 3 | 16509 | 1127 | >sp\|O95239-2\|KIF4A_HUMAN Isoform 2 of Chromosome-associated kinesin KIF4A OS=Homo sapiens OX=9606 GN=KIF4A |
| 582 | 17.82 | 3 | 2 | 13509 | 323 | >sp\|Q14847-2\|LASP1_HUMAN Isoform 2 of LIM and SH3 domain protein 1 OS=Homo sapiens OX=9606 GN=LASP1 |
| 583 | 17.76 | 3 | 3 | 36278 | 271 | >sp\|Q9Y5M8\|SRPRB_HUMAN Signal recognition particle receptor subunit beta OS=Homo sapiens OX=9606 GN=SRPRB PE=1 SV=3 |
| 584 | 17.67 | 2 | 2 | 6648 | 167 | >sp\|P42771-4\|CDN2A_HUMAN Isoform 5 of Cyclin-dependent kinase inhibitor 2A OS=Homo sapiens OX=9606 GN=CDKN2A |
| 585 | 17.67 | 4 | 4 | 21432 | 105 | >sp\|Q9Y3U8\|RL36_HUMAN 60S ribosomal protein L36 OS=Homo sapiens OX=9606 GN=RPL36 PE=1 SV=3 |
| 586 | 17.61 | 3 | 2 | 17452 | 161 | >sp\|Q15370-2\|ELOB_HUMAN Isoform 2 of Elongin-B OS=Homo sapiens OX=9606 GN=ELOB |
| 587 | 17.60 | 5 | 5 | 36188 | 2752 | >sp\|Q9UQ35\|SRRM2_HUMAN Serine/arginine repetitive matrix protein 2 OS=Homo sapiens OX=9606 GN=SRRM2 PE=1 SV=2 |
| 588 | 17.51 | 2 | 2 | 22189 | 143 | >sp\|P62266\|RS23_HUMAN 40S ribosomal protein S23 OS=Homo sapiens OX=9606 GN=RPS23 PE=1 SV=3 |
| 589 | 17.50 | 3 | 3 | 19553 | 520 | >sp\|P55809\|SCOT1_HUMAN Succinyl-CoA:3-ketoacid coenzyme A transferase 1; mitochondrial OS=Homo sapiens OX=9606 GN=OXCT1 PE=1 SV |
| 590 | 17.47 | 1 | 1 | 35209 | 142 | >sp\|Q9NS69\|TOM22_HUMAN Mitochondrial import receptor subunit TOM22 homolog OS=Homo sapiens OX=9606 GN=TOMM22 PE=1 SV=3 |
| 591 | 17.45 | 3 | 3 | 31397 | 303 | >sp\|Q9Y6C9\|MTCH2_HUMAN Mitochondrial carrier homolog 2 OS=Homo sapiens OX=9606 GN=MTCH2 PE=1 SV=1 |
| 592 | 17.41 | 3 | 3 | 19270 | 977 | >sp\|Q9P2E9-3\|RRBP1_HUMAN Isoform 2 of Ribosome-binding protein 1 OS=Homo sapiens OX=9606 GN=RRBP1 |
| 593 | 17.40 | 2 | 2 | 10135 | 560 | >sp\|Q9BY44-3\|EIF2A_HUMAN Isoform 3 of Eukaryotic translation initiation factor 2A OS=Homo sapiens OX=9606 GN=EIF2A |
| 594 | 17.39 | 2 | 2 | 22918 | 183 | >sp\|O00193\|SMAP_HUMAN Small acidic protein OS=Homo sapiens OX=9606 GN=SMAP PE=1 SV=1 |
| 595 | 17.33 | 3 | 2 | 19301 | 56 | >sp\|P62273\|RS29_HUMAN 40S ribosomal protein S29 OS=Homo sapiens OX=9606 GN=RPS29 PE=1 SV=2 |
| 596 | 17.29 | 7 | 5 | 22922 | 1217 | >sp\|Q9UQE7\|SMC3_HUMAN Structural maintenance of chromosomes protein 3 OS=Homo sapiens OX=9606 GN=SMC3 PE=1 SV=2 |

#### Score distribution

890 peptide matches above score 32, 1188 (21 reverse) matches above 20
Top score for reverse: 45, top score for reverse with > 9 aa's: 23
Charge distribution: z=+1: 0, z=+2: 859, z=+3: 31, z=+4: 0

#### Mass measurement errors

**Precursors**

*Before recalibration:*
Median precursor m/z error (observed minus true): -0.0004 Da, -0.8 ppm
Median precursor accuracy (absolute value of error): 0.0004 Da, 0.8 ppm
Precursors measured too high: 69 Too low: 709

*After recalibration:*

Median precursor m/z error (observed minus true): 0.0000 Da, 0.0 ppm
Median precursor accuracy (absolute value of error): 0.0002 Da, 0.3 ppm
Precursors measured too high: 408 Too low: 375

Off-by-one errors (nominal mass is +1 isotope): 9.9% (86/867)

**Fragments**

*Before recalibration:*
Median fragment m/z error (observed minus true): 0.0909 Da, 140.6 ppm
Median fragment accuracy (absolute value of error): 0.0912 Da, 142.4 ppm
Fragments (within ±0.5 Da) measured too high: 4485 Too low: 172

*After recalibration:*
Median fragment m/z error (observed minus true): 0.0190 Da, 31.1 ppm
Median fragment accuracy (absolute value of error): 0.0324 Da, 51.7 ppm
Fragments (within ±0.5 Da) measured too high: 2812 Too low: 1844

#### Cysteine

Cysteine set to a fixed modification of +57
Completeness: 76.2% (48/63) peptides C[+57], 22.2% (14/63) C[+0], 1.6% (1/63) C[+71] (from gel)
Carbamidomethylation artifacts (H,K,N-terminus[+57/+114]): 0.0% (0/979) of peptides
DTT artifacts (C[+209]): 5.0% (51/1030) of peptides
Disulfide bridge (unmodified_C[-2]): 0.0% (0/2) of peptides containing at least two C's

#### Nonspecific cleavage

Cleavage sites (C-side): RK
Missed cleavage: 24.7% (197/797) of semitryptic peptides contain an internal K or R not followed by P
Semitryptic peptides (% of tryptic and semitryptic): 2.8% (22/797) ragged-N, 0.8% (6/797) ragged-C
Nontryptic peptides (% of all peptides): 0.1% (1/801)

#### Oxidation

Oxidized methionine (M[+16]): 19.7% (69/351) of peptides containing M[+0] or M[+16]
Side chain loss from oxidized methionine (M[-48]): 0.0% (0/281) of peptides containing M[+0] or M[-48]
Doubly oxidized methionine (M[+32]): 0.0% (0/226) of peptides containing one M[+32] or M[+0]
Oxidized histidine and tryptophan (H[+16], W[+16]): 2.1% (4/194) (2 H, 2 W) of peptides containing H or W
Doubly oxidized tryptophan (W[+32]): 0.0% (0/66) of peptides containing W but not M
Triply oxidized cysteine (C[+48]): 0.0% (0/56) of peptides containing C

#### Chemical modifications

Deamidated asparagine or glutamine: 9.9% (87/877) (84 N, 3 Q) of peptides containing N and/or Q
Amidated aspartic or glutamic acid: 0.6% (6/952) of peptides containing D and/or E
Pyro-glu N-terminus (Q[-17], E[-18], C[+57][-17]): 12.5% (10/80) (9 -17, 1 -18) of peptides with N-terminal Q, E, or camC
Sodiation: 0.0% (2/1124) of all peptides (0 on E and D)
Carbamylation (N-terminus[+43], R[+43], K[+43]): 0.4% (4/1128) (0 N-term) of peptides
Carbamylated methionine (M[+43]): 0.0% (0/351) of peptides containing M
Formaldehyde (N-terminus[+12], W[+12]): 0.0% (0/1124) of peptides
Acetaldehyde (N-terminus[+26], H[+26], K[+26]): 0.1% (1/1124) (1 N-term) of peptides
N-terminal methylation/dimethylation (N-terminus[+14/+28]): 0.4% (5/1124) (5 +14, 0 +28) of peptides
Peptide (not protein) N-terminal acetylation (N-terminus[+42]): 0.0% (0/1124) of peptides
Cysteine propionamide (C[+71]): 1.8% (1/56) of peptides containing C
Methyl ester (E[+14]): 0.6% (4/667) of peptides containing E
Formylation (S[+28], T[+28]): 0.0% (0/632) of peptides containing S or T

#### Posttranslational modifications

Hydroxyproline: 1.9% (8/418) of peptides containing P
Phosphorylation: 0.0% (0/746) (0 S, 0 T, 0 Y) of peptides containing S, T, or Y
Beta-elimination (S[-18], T[-18]): 0.0% (0/634) peptides containing S or T
Dimethylation (K[+28], R[+28]): 1.4% (15/1101) (13 K, 2 R) of peptides containing K or R
Methylation (K[+14], H[+14], N[+14], R[+14]): 0.8% (9/1104) (1 K) of peptides containing K, H, N, or R
Acetylation (or guanidination or trimethylation) (K[+42]): 1.4% (10/697) of peptides containing K
Protein N-terminal acetylation: 62.5% (5/8) of N-terminal peptides

#### Computational information

Spectrum file: D:\Luci\Moggridge et al data\ch_03Fe...p_1.raw_Preview_181130_173552\objs\ch_03Fe...-100p_1.mgf
Protein database: D:\Luci\PD Databases\Schulz_CP_Homo_sapiens_proteomeUP000005640reveiwed_Download_20180420_proteins_20303.fasta
Presets: Cysteine 57, Lysine 0, Arginine 0, N-terminus 0, C-terminus 0
CID fragmentation
Digestion after RK
First pass is fully tryptic digestion
No wildcard search
Read in 22629 spectrum/charge combinations: z=+1 0, z=+2 15754, z=+3 6053, z=+4 822
Precursor mass from 758 to 2999 Da
Read 42313 proteins
Scored 359262231 candidate/spectrum/charge combinations

Preview version: v2.13.17
Output folder: D:\Luci\Moggridge et al data\ch_03Fe...p_1.raw_Preview_181130_173552

Click to open **parameters** used for this run.

Note: The Details page reports the rates of modification on eligible peptides, whereas the Summary page reports potential gains in the total number of identifications. Denominators in the percentages may also vary from search to search due to "second-order" effects such as multiply modified peptides and corrections for hits to decoys.

Summary | Detail

www.proteinmetrics.com

**Preview™ by PROTEIN METRICS Inc.**

Summary | Detail

#### Table of Contents (ch_06Feb2018_SP3-Binding-ethanol-100p_mix-yes_rinse-yes_1.raw) (Schulz_CP_Homo_sapiens_proteomeUP000005640reveiwed_Download_20180420_proteins_20303.fasta)

1. Representative proteins
2. Score distribution
3. Mass measurement errors
4. Cysteine
5. Nonspecific cleavage
6. Oxidation
7. Chemical modifications
8. Posttranslational modifications
9. Computation information

#### Representative proteins

Right click (or option-click) and save link: **peptide and protein identifications** (in csv format). Or, access the **folder**.

| Rank | ProtScore | # Spectra ID'd | # Unique Peptides | Database Sequence # | # aa's | Protein Name |
|---|---|---|---|---|---|---|
| 1 | 284.29 | 30 | 29 | 29794 | 1960 | >sp\|P35579\|MYH9_HUMAN Myosin-9 OS=Homo sapiens OX=9606 GN=MYH9 PE=1 SV=4 |
| 2 | 251.68 | 36 | 29 | 10117 | 858 | >sp\|P13639\|EF2_HUMAN Elongation factor 2 OS=Homo sapiens OX=9606 GN=EEF2 PE=1 SV=4 |
| 3 | 225.91 | 27 | 25 | 27671 | 2101 | >sp\|Q14980-2\|NUMA1_HUMAN Isoform 2 of Nuclear mitotic apparatus protein 1 OS=Homo sapiens OX=9606 GN=NUMA1 |
| 4 | 223.62 | 34 | 25 | 14875 | 854 | >sp\|P07900-2\|HS90A_HUMAN Isoform 2 of Heat shock protein HSP 90-alpha OS=Homo sapiens OX=9606 GN=HSP90AA1 |
| 5 | 222.18 | 36 | 25 | 12650 | 641 | >sp\|P0DMV9\|HS71B_HUMAN Heat shock 70 kDa protein 1B OS=Homo sapiens OX=9606 GN=HSPA1B PE=1 SV=1 |
| 6 | 197.01 | 20 | 19 | 38994 | 806 | >sp\|P55072\|TERA_HUMAN Transitional endoplasmic reticulum ATPase OS=Homo sapiens OX=9606 GN=VCP PE=1 SV=4 |
| 7 | 183.36 | 24 | 22 | 17426 | 2639 | >sp\|P21333-2\|FLNA_HUMAN Isoform 2 of Filamin-A OS=Homo sapiens OX=9606 GN=FLNA |
| 8 | 183.34 | 21 | 16 | 11486 | 646 | >sp\|P11142\|HSP7C_HUMAN Heat shock cognate 71 kDa protein OS=Homo sapiens OX=9606 GN=HSPA8 PE=1 SV=1 |
| 9 | 172.12 | 21 | 21 | 12573 | 691 | >sp\|P52272-2\|HNRPM_HUMAN Isoform 2 of Heterogeneous nuclear ribonucleoprotein M OS=Homo sapiens OX=9606 GN=HNRNPM |
| 10 | 159.63 | 18 | 17 | 38833 | 548 | >sp\|P50990\|TCPQ_HUMAN T-complex protein 1 subunit theta OS=Homo sapiens OX=9606 GN=CCT8 PE=1 SV=4 |
| 11 | 159.14 | 20 | 16 | 26540 | 586 | >sp\|P20700\|LMNB1_HUMAN Lamin-B1 OS=Homo sapiens OX=9606 GN=LMNB1 PE=1 SV=2 |
| 12 | 152.70 | 22 | 22 | 17521 | 803 | >sp\|P14625\|ENPL_HUMAN Endoplasmin OS=Homo sapiens OX=9606 GN=HSP90B1 PE=1 SV=1 |
| 13 | 151.18 | 19 | 15 | 5194 | 654 | >sp\|P11021\|BIP_HUMAN Endoplasmic reticulum chaperone BiP OS=Homo sapiens OX=9606 GN=HSPA5 PE=1 SV=2 |
| 14 | 149.15 | 29 | 26 | 3268 | 5890 | >sp\|Q09666\|AHNK_HUMAN Neuroblast differentiation-associated protein AHNAK OS=Homo sapiens OX=9606 GN=AHNAK PE=1 SV=2 |
| 15 | 148.23 | 22 | 16 | 34832 | 445 | >sp\|P68371\|TBB4B_HUMAN Tubulin beta-4B chain OS=Homo sapiens OX=9606 GN=TUBB4B PE=1 SV=1 |
| 16 | 146.45 | 19 | 19 | 14295 | 679 | >sp\|P38646\|GRP75_HUMAN Stress-70 protein; mitochondrial OS=Homo sapiens OX=9606 GN=HSPA9 PE=1 SV=2 |
| 17 | 140.34 | 15 | 14 | 28314 | 645 | >sp\|P13667\|PDIA4_HUMAN Protein disulfide-isomerase A4 OS=Homo sapiens OX=9606 GN=PDIA4 PE=1 SV=2 |
| 18 | 139.14 | 20 | 14 | 34144 | 375 | >sp\|P60709\|ACTB_HUMAN Actin; cytoplasmic 1 OS=Homo sapiens OX=9606 GN=ACTB PE=1 SV=1 |
| 19 | 135.68 | 18 | 15 | 27856 | 1014 | >sp\|P09874\|PARP1_HUMAN Poly [ADP-ribose] polymerase 1 OS=Homo sapiens OX=9606 GN=PARP1 PE=1 SV=4 |
| 20 | 132.25 | 27 | 25 | 42235 | 4128 | >sp\|P78527\|PRKDC_HUMAN DNA-dependent protein kinase catalytic subunit OS=Homo sapiens OX=9606 GN=PRKDC PE=1 SV=3 |
| 21 | 131.99 | 22 | 18 | 10608 | 434 | >sp\|P06733\|ENOA_HUMAN Alpha-enolase OS=Homo sapiens OX=9606 GN=ENO1 PE=1 SV=2 |
| 22 | 130.96 | 16 | 13 | 16301 | 406 | >sp\|P60842\|IF4A1_HUMAN Eukaryotic initiation factor 4A-I OS=Homo sapiens OX=9606 GN=EIF4A1 PE=1 SV=1 |
| 23 | 130.81 | 19 | 17 | 41440 | 466 | >sp\|P08670\|VIME_HUMAN Vimentin OS=Homo sapiens OX=9606 GN=VIM PE=1 SV=4 |
| 24 | 130.57 | 16 | 13 | 28584 | 505 | >sp\|P30101\|PDIA3_HUMAN Protein disulfide-isomerase A3 OS=Homo sapiens OX=9606 GN=PDIA3 PE=1 SV=4 |
| 25 | 127.14 | 19 | 16 | 5883 | 573 | >sp\|P10809\|CH60_HUMAN 60 kDa heat shock protein; mitochondrial OS=Homo sapiens OX=9606 GN=HSPD1 PE=1 SV=2 |
| 26 | 120.99 | 26 | 21 | 17366 | 2511 | >sp\|P49327\|FAS_HUMAN Fatty acid synthase OS=Homo sapiens OX=9606 GN=FASN PE=1 SV=3 |
| 27 | 118.92 | 12 | 11 | 199 | 553 | >sp\|P25705\|ATPA_HUMAN ATP synthase subunit alpha; mitochondrial OS=Homo sapiens OX=9606 GN=ATP5F1A PE=1 SV=1 |
| 28 | 118.42 | 22 | 21 | 17290 | 1382 | >sp\|Q14152\|EIF3A_HUMAN Eukaryotic translation initiation factor 3 subunit A OS=Homo sapiens OX=9606 GN=EIF3A PE=1 SV=1 |
| 29 | 118.16 | 13 | 13 | 22202 | 353 | >sp\|P22626\|ROA2_HUMAN Heterogeneous nuclear ribonucleoproteins A2/B1 OS=Homo sapiens OX=9606 GN=HNRNPA2B1 PE=1 SV=2 |
| 30 | 116.47 | 18 | 17 | 35928 | 590 | >sp\|P31948-2\|STIP1_HUMAN Isoform 2 of Stress-induced-phosphoprotein 1 OS=Homo sapiens OX=9606 GN=STIP1 |
| 31 | 113.71 | 14 | 12 | 4219 | 935 | >sp\|P11586\|C1TC_HUMAN C-1-tetrahydrofolate synthase; cytoplasmic OS=Homo sapiens OX=9606 GN=MTHFD1 PE=1 SV=3 |
| 32 | 109.13 | 12 | 11 | 12627 | 825 | >sp\|Q00839\|HNRPU_HUMAN Heterogeneous nuclear ribonucleoprotein U OS=Homo sapiens OX=9606 GN=HNRNPU PE=1 SV=6 |
| 33 | 107.95 | 11 | 11 | 14876 | 724 | >sp\|P08238\|HS90B_HUMAN Heat shock protein HSP 90-beta OS=Homo sapiens OX=9606 GN=HSP90AB1 PE=1 SV=4 |
| 34 | 107.74 | 11 | 10 | 2722 | 529 | >sp\|P06576\|ATPB_HUMAN ATP synthase subunit beta; mitochondrial OS=Homo sapiens OX=9606 GN=ATP5F1B PE=1 SV=3 |
| 35 | 106.89 | 12 | 12 | 16442 | 511 | >sp\|P05787-2\|K2C8_HUMAN Isoform 2 of Keratin; type II cytoskeletal 8 OS=Homo sapiens OX=9606 GN=KRT8 |
| 36 | 106.84 | 12 | 11 | 14882 | 840 | >sp\|P34932\|HSP74_HUMAN Heat shock 70 kDa protein 4 OS=Homo sapiens OX=9606 GN=HSPA4 PE=1 SV=4 |
| 37 | 106.72 | 13 | 13 | 12431 | 588 | >sp\|O60506-2\|HNRPQ_HUMAN Isoform 2 of Heterogeneous nuclear ribonucleoprotein Q OS=Homo sapiens OX=9606 GN=SYNCRIP |
| 38 | 106.59 | 14 | 14 | 11581 | 898 | >sp\|Q12906-7\|ILF3_HUMAN Isoform 7 of Interleukin enhancer-binding factor 3 OS=Homo sapiens OX=9606 GN=ILF3 |
| 39 | 104.93 | 16 | 11 | 12512 | 440 | >sp\|P61978-3\|HNRPK_HUMAN Isoform 3 of Heterogeneous nuclear ribonucleoprotein K OS=Homo sapiens OX=9606 GN=HNRNPK |
| 40 | 104.59 | 14 | 13 | 34358 | 911 | >sp\|O43707\|ACTN4_HUMAN Alpha-actinin-4 OS=Homo sapiens OX=9606 GN=ACTN4 PE=1 SV=2 |
| 41 | 103.81 | 14 | 13 | 11476 | 817 | >sp\|Q92598-3\|HS105_HUMAN Isoform 3 of Heat shock protein 105 kDa OS=Homo sapiens OX=9606 GN=HSPH1 |
| 42 | 103.23 | 12 | 11 | 19742 | 432 | >sp\|P23526\|SAHH_HUMAN Adenosylhomocysteinase OS=Homo sapiens OX=9606 GN=AHCY PE=1 SV=4 |

| | | | | | | |
|---|---|---|---|---|---|---|
| 43 | 101.35 | 15 | 14 | 15628 | 2633 | >sp\|O75369-8\|FLNB_HUMAN Isoform 8 of Filamin-B OS=Homo sapiens OX=9606 GN=FLNB |
| 44 | 101.09 | 14 | 11 | 13137 | 334 | >sp\|P07195\|LDHB_HUMAN L-lactate dehydrogenase B chain OS=Homo sapiens OX=9606 GN=LDHB PE=1 SV=2 |
| 45 | 98.83 | 14 | 12 | 23683 | 2452 | >sp\|Q13813-3\|SPTN1_HUMAN Isoform 3 of Spectrin alpha chain; non-erythrocytic 1 OS=Homo sapiens OX=9606 GN=SPTAN1 |
| 46 | 98.78 | 14 | 13 | 8068 | 535 | >sp\|P17844-2\|DDX5_HUMAN Isoform 2 of Probable ATP-dependent RNA helicase DDX5 OS=Homo sapiens OX=9606 GN=DDX5 |
| 47 | 98.68 | 13 | 12 | 11112 | 711 | >sp\|Q92945\|FUBP2_HUMAN Far upstream element-binding protein 2 OS=Homo sapiens OX=9606 GN=KHSRP PE=1 SV=4 |
| 48 | 97.51 | 12 | 12 | 30785 | 636 | >sp\|P11940\|PABP1_HUMAN Polyadenylate-binding protein 1 OS=Homo sapiens OX=9606 GN=PABPC1 PE=1 SV=2 |
| 49 | 97.04 | 18 | 15 | 27269 | 630 | >sp\|P13797\|PLST_HUMAN Plastin-3 OS=Homo sapiens OX=9606 GN=PLS3 PE=1 SV=4 |
| 50 | 94.61 | 10 | 10 | 35062 | 488 | >sp\|P78371-2\|TCPB_HUMAN Isoform 2 of T-complex protein 1 subunit beta OS=Homo sapiens OX=9606 GN=CCT2 |
| 51 | 94.22 | 10 | 9 | 26393 | 614 | >sp\|P02545-6\|LMNA_HUMAN Isoform 6 of Prelamin-A/C OS=Homo sapiens OX=9606 GN=LMNA |
| 52 | 91.87 | 14 | 12 | 16431 | 430 | >sp\|P05783\|K1C18_HUMAN Keratin; type I cytoskeletal 18 OS=Homo sapiens OX=9606 GN=KRT18 PE=1 SV=2 |
| 53 | 91.74 | 11 | 10 | 22201 | 267 | >sp\|P09651-3\|ROA1_HUMAN Isoform 2 of Heterogeneous nuclear ribonucleoprotein A1 OS=Homo sapiens OX=9606 GN=HNRNPA1 |
| 54 | 90.82 | 14 | 11 | 7519 | 462 | >sp\|Q5VTE0\|EF1A3_HUMAN Putative elongation factor 1-alpha-like 3 OS=Homo sapiens OX=9606 GN=EEF1A1P5 PE=5 SV=1 |
| 55 | 90.65 | 14 | 11 | 34826 | 451 | >sp\|P68363\|TBA1B_HUMAN Tubulin alpha-1B chain OS=Homo sapiens OX=9606 GN=TUBA1B PE=1 SV=1 |
| 56 | 89.57 | 13 | 8 | 14006 | 103 | >sp\|P62805\|H4_HUMAN Histone H4 OS=Homo sapiens OX=9606 GN=HIST1H4A PE=1 SV=2 |
| 57 | 88.39 | 9 | 9 | 39061 | 545 | >sp\|P49368\|TCPG_HUMAN T-complex protein 1 subunit gamma OS=Homo sapiens OX=9606 GN=CCT3 PE=1 SV=4 |
| 58 | 87.79 | 12 | 10 | 20532 | 707 | >sp\|P23246\|SFPQ_HUMAN Splicing factor; proline- and glutamine-rich OS=Homo sapiens OX=9606 GN=SFPQ PE=1 SV=2 |
| 59 | 87.02 | 14 | 13 | 7610 | 1270 | >sp\|Q08211\|DHX9_HUMAN ATP-dependent RNA helicase A OS=Homo sapiens OX=9606 GN=DHX9 PE=1 SV=4 |
| 60 | 87.01 | 7 | 7 | 27427 | 254 | >sp\|P18669\|PGAM1_HUMAN Phosphoglycerate mutase 1 OS=Homo sapiens OX=9606 GN=PGAM1 PE=1 SV=2 |
| 61 | 86.94 | 11 | 9 | 38832 | 556 | >sp\|P17987\|TCPA_HUMAN T-complex protein 1 subunit alpha OS=Homo sapiens OX=9606 GN=TCP1 PE=1 SV=1 |
| 62 | 85.05 | 10 | 8 | 7218 | 257 | >sp\|P29692-3\|EF1D_HUMAN Isoform 3 of Elongation factor 1-delta OS=Homo sapiens OX=9606 GN=EEF1D |
| 63 | 84.55 | 14 | 11 | 3913 | 364 | >sp\|P04075\|ALDOA_HUMAN Fructose-bisphosphate aldolase A OS=Homo sapiens OX=9606 GN=ALDOA PE=1 SV=2 |
| 64 | 83.87 | 11 | 11 | 40982 | 2363 | >sp\|P12270\|TPR_HUMAN Nucleoprotein TPR OS=Homo sapiens OX=9606 GN=TPR PE=1 SV=3 |
| 65 | 82.99 | 13 | 13 | 23083 | 1233 | >sp\|Q14683\|SMC1A_HUMAN Structural maintenance of chromosomes protein 1A OS=Homo sapiens OX=9606 GN=SMC1A PE=1 SV=2 |
| 66 | 81.36 | 8 | 8 | 12655 | 293 | >sp\|P04406-2\|G3P_HUMAN Isoform 2 of Glyceraldehyde-3-phosphate dehydrogenase OS=Homo sapiens OX=9606 GN=GAPDH |
| 67 | 81.12 | 11 | 9 | 3770 | 255 | >sp\|P62258\|1433E_HUMAN 14-3-3 protein epsilon OS=Homo sapiens OX=9606 GN=YWHAE PE=1 SV=1 |
| 68 | 80.44 | 9 | 7 | 12428 | 306 | >sp\|Q14103-3\|HNRPD_HUMAN Isoform 3 of Heterogeneous nuclear ribonucleoprotein D0 OS=Homo sapiens OX=9606 GN=HNRNPD |
| 69 | 80.12 | 10 | 9 | 13614 | 963 | >sp\|P33176\|KINH_HUMAN Kinesin-1 heavy chain OS=Homo sapiens OX=9606 GN=KIF5B PE=1 SV=1 |
| 70 | 79.70 | 9 | 8 | 3563 | 430 | >sp\|P00505\|AATM_HUMAN Aspartate aminotransferase; mitochondrial OS=Homo sapiens OX=9606 GN=GOT2 PE=1 SV=3 |
| 71 | 79.65 | 9 | 9 | 4123 | 1230 | >sp\|Q86VP6\|CAND1_HUMAN Cullin-associated NEDD8-dissociated protein 1 OS=Homo sapiens OX=9606 GN=CAND1 PE=1 SV=2 |
| 72 | 79.42 | 16 | 15 | 7399 | 4646 | >sp\|Q14204\|DYHC1_HUMAN Cytoplasmic dynein 1 heavy chain 1 OS=Homo sapiens OX=9606 GN=DYNC1H1 PE=1 SV=5 |
| 73 | 79.27 | 17 | 11 | 15619 | 459 | >sp\|Q02790\|FKBP4_HUMAN Peptidyl-prolyl cis-trans isomerase FKBP4 OS=Homo sapiens OX=9606 GN=FKBP4 PE=1 SV=3 |
| 74 | 78.91 | 11 | 10 | 41146 | 2541 | >sp\|Q9Y490\|TLN1_HUMAN Talin-1 OS=Homo sapiens OX=9606 GN=TLN1 PE=1 SV=3 |
| 75 | 78.80 | 12 | 11 | 32551 | 732 | >sp\|Q08J23-2\|NSUN2_HUMAN Isoform 2 of tRNA (cytosine(34)-C(5))-methyltransferase OS=Homo sapiens OX=9606 GN=NSUN2 |
| 76 | 78.66 | 13 | 11 | 21434 | 427 | >sp\|P36578\|RL4_HUMAN 60S ribosomal protein L4 OS=Homo sapiens OX=9606 GN=RPL4 PE=1 SV=5 |
| 77 | 77.43 | 7 | 7 | 6958 | 652 | >sp\|Q92841-3\|DDX17_HUMAN Isoform 4 of Probable ATP-dependent RNA helicase DDX17 OS=Homo sapiens OX=9606 GN=DDX17 |
| 78 | 77.26 | 12 | 12 | 9793 | 966 | >sp\|Q14697-2\|GANAB_HUMAN Isoform 2 of Neutral alpha-glucosidase AB OS=Homo sapiens OX=9606 GN=GANAB |
| 79 | 77.03 | 9 | 9 | 34806 | 220 | >sp\|P37802-2\|TAGL2_HUMAN Isoform 2 of Transgelin-2 OS=Homo sapiens OX=9606 GN=TAGLN2 |
| 80 | 77.02 | 11 | 11 | 22920 | 1197 | >sp\|O95347\|SMC2_HUMAN Structural maintenance of chromosomes protein 2 OS=Homo sapiens OX=9606 GN=SMC2 PE=1 SV=2 |
| 81 | 76.78 | 8 | 8 | 41436 | 283 | >sp\|P21796\|VDAC1_HUMAN Voltage-dependent anion-selective channel protein 1 OS=Homo sapiens OX=9606 GN=VDAC1 PE=1 SV=2 |
| 82 | 76.74 | 12 | 11 | 13423 | 531 | >sp\|P14618-2\|KPYM_HUMAN Isoform M1 of Pyruvate kinase PKM OS=Homo sapiens OX=9606 GN=PKM |
| 83 | 76.54 | 7 | 7 | 40882 | 248 | >sp\|P06753-5\|TPM3_HUMAN Isoform 5 of Tropomyosin alpha-3 chain OS=Homo sapiens OX=9606 GN=TPM3 |
| 84 | 76.38 | 10 | 10 | 31097 | 331 | >sp\|Q9Y266\|NUDC_HUMAN Nuclear migration protein nudC OS=Homo sapiens OX=9606 GN=NUDC PE=1 SV=1 |
| 85 | 76.03 | 8 | 8 | 12280 | 449 | >sp\|P31943\|HNRH1_HUMAN Heterogeneous nuclear ribonucleoprotein H OS=Homo sapiens OX=9606 GN=HNRNPH1 PE=1 SV=4 |
| 86 | 75.93 | 12 | 11 | 17694 | 586 | >sp\|P15311\|EZRI_HUMAN Ezrin OS=Homo sapiens OX=9606 GN=EZR PE=1 SV=4 |
| 87 | 75.81 | 10 | 9 | 37210 | 324 | >sp\|P67809\|YBOX1_HUMAN Nuclease-sensitive element-binding protein 1 OS=Homo sapiens OX=9606 GN=YBX1 PE=1 SV=3 |
| 88 | 75.71 | 9 | 8 | 14832 | 411 | >sp\|P38919\|IF4A3_HUMAN Eukaryotic initiation factor 4A-III OS=Homo sapiens OX=9606 GN=EIF4A3 PE=1 SV=4 |
| 89 | 74.92 | 9 | 8 | 12288 | 293 | >sp\|P07910-2\|HNRPC_HUMAN Isoform C1 of Heterogeneous nuclear ribonucleoproteins C1/C2 OS=Homo sapiens OX=9606 GN=HNRNPC |
| 90 | 74.24 | 12 | 11 | 5624 | 1639 | >sp\|Q00610-2\|CLH1_HUMAN Isoform 2 of Clathrin heavy chain 1 OS=Homo sapiens OX=9606 GN=CLTC |
| 91 | 73.87 | 9 | 8 | 25995 | 543 | >sp\|P33993-3\|MCM7_HUMAN Isoform 3 of DNA replication licensing factor MCM7 OS=Homo sapiens OX=9606 GN=MCM7 |
| 92 | 73.57 | 11 | 8 | 40876 | 249 | >sp\|P60174-1\|TPIS_HUMAN Isoform 2 of Triosephosphate isomerase OS=Homo sapiens OX=9606 GN=TPI1 |
| 93 | 73.46 | 9 | 8 | 37650 | 962 | >sp\|Q15029-3\|U5S1_HUMAN Isoform 3 of 116 kDa U5 small nuclear ribonucleoprotein component OS=Homo sapiens OX=9606 GN=EFTUD2 |
| 94 | 73.18 | 11 | 11 | 38968 | 448 | >sp\|P48643-2\|TCPE_HUMAN Isoform 2 of T-complex protein 1 subunit epsilon OS=Homo sapiens OX=9606 GN=CCT5 |
| 95 | 71.98 | 8 | 6 | 27451 | 187 | >sp\|P30086\|PEBP1_HUMAN Phosphatidylethanolamine-binding protein 1 OS=Homo sapiens OX=9606 GN=PEBP1 PE=1 SV=3 |
| 96 | 71.46 | 11 | 10 | 40575 | 651 | >sp\|Q12931-2\|TRAP1_HUMAN Isoform 2 of Heat shock protein 75 kDa; mitochondrial OS=Homo sapiens OX=9606 GN=TRAP1 |
| 97 | 70.99 | 10 | 9 | 11247 | 452 | >sp\|P49411\|EFTU_HUMAN Elongation factor Tu; mitochondrial OS=Homo sapiens OX=9606 GN=TUFM PE=1 SV=2 |

| | | | | | | |
|---|---|---|---|---|---|---|
| 98 | 70.06 | 9 | 9 | 11110 | 644 | >sp\|Q96AE4\|FUBP1_HUMAN Far upstream element-binding protein 1 OS=Homo sapiens OX=9606 GN=FUBP1 PE=1 SV=3 |
| 99 | 69.98 | 10 | 9 | 42044 | 317 | >sp\|P63244\|RACK1_HUMAN Receptor of activated protein C kinase 1 OS=Homo sapiens OX=9606 GN=RACK1 PE=1 SV=3 |
| 100 | 69.91 | 14 | 12 | 21886 | 264 | >sp\|P61247\|RS3A_HUMAN 40S ribosomal protein S3a OS=Homo sapiens OX=9606 GN=RPS3A PE=1 SV=2 |
| 101 | 69.37 | 9 | 9 | 41256 | 609 | >sp\|P12956\|XRCC6_HUMAN X-ray repair cross-complementing protein 6 OS=Homo sapiens OX=9606 GN=XRCC6 PE=1 SV=2 |
| 102 | 68.69 | 9 | 8 | 28625 | 299 | >sp\|Q99623\|PHB2_HUMAN Prohibitin-2 OS=Homo sapiens OX=9606 GN=PHB2 PE=1 SV=2 |
| 103 | 68.16 | 7 | 7 | 21435 | 317 | >sp\|P05388\|RLA0_HUMAN 60S acidic ribosomal protein P0 OS=Homo sapiens OX=9606 GN=RPLP0 PE=1 SV=1 |
| 104 | 67.90 | 11 | 11 | 22334 | 592 | >sp\|P31939\|PUR9_HUMAN Bifunctional purine biosynthesis protein PURH OS=Homo sapiens OX=9606 GN=ATIC PE=1 SV=3 |
| 105 | 67.77 | 14 | 12 | 30819 | 710 | >sp\|P19338\|NUCL_HUMAN Nucleolin OS=Homo sapiens OX=9606 GN=NCL PE=1 SV=3 |
| 106 | 67.57 | 10 | 9 | 18997 | 391 | >sp\|P38159\|RBMX_HUMAN RNA-binding motif protein; X chromosome OS=Homo sapiens OX=9606 GN=RBMX PE=1 SV=3 |
| 107 | 65.46 | 8 | 8 | 16289 | 381 | >sp\|P12277\|KCRB_HUMAN Creatine kinase B-type OS=Homo sapiens OX=9606 GN=CKB PE=1 SV=1 |
| 108 | 65.16 | 5 | 4 | 18182 | 925 | >sp\|E9PAV3-2\|NACAM_HUMAN Isoform skNAC-2 of Nascent polypeptide-associated complex subunit alpha; muscle-specific form OS=Homo |
| 109 | 65.14 | 13 | 11 | 13288 | 361 | >sp\|P00338-3\|LDHA_HUMAN Isoform 3 of L-lactate dehydrogenase A chain OS=Homo sapiens OX=9606 GN=LDHA |
| 110 | 64.96 | 10 | 9 | 38884 | 835 | >sp\|Q13263\|TIF1B_HUMAN Transcription intermediary factor 1-beta OS=Homo sapiens OX=9606 GN=TRIM28 PE=1 SV=5 |
| 111 | 64.82 | 14 | 10 | 21936 | 505 | >sp\|Q9Y3I0\|RTCB_HUMAN tRNA-splicing ligase RtcB homolog OS=Homo sapiens OX=9606 GN=RTCB PE=1 SV=1 |
| 112 | 64.58 | 12 | 8 | 16376 | 514 | >sp\|P12268\|IMDH2_HUMAN Inosine-5'-monophosphate dehydrogenase 2 OS=Homo sapiens OX=9606 GN=IMPDH2 PE=1 SV=2 |
| 113 | 64.18 | 6 | 6 | 27443 | 272 | >sp\|P35232\|PHB_HUMAN Prohibitin OS=Homo sapiens OX=9606 GN=PHB PE=1 SV=1 |
| 114 | 64.10 | 9 | 8 | 42249 | 433 | >sp\|P35998\|PRS7_HUMAN 26S proteasome regulatory subunit 7 OS=Homo sapiens OX=9606 GN=PSMC2 PE=1 SV=3 |
| 115 | 63.78 | 12 | 10 | 38249 | 499 | >sp\|Q99832-3\|TCPH_HUMAN Isoform 3 of T-complex protein 1 subunit eta OS=Homo sapiens OX=9606 GN=CCT7 |
| 116 | 63.78 | 11 | 8 | 28052 | 393 | >sp\|Q8NC51-3\|PAIRB_HUMAN Isoform 3 of Plasminogen activator inhibitor 1 RNA-binding protein OS=Homo sapiens OX=9606 GN=SERBP1 |
| 117 | 63.52 | 9 | 9 | 7556 | 662 | >sp\|O00571\|DDX3X_HUMAN ATP-dependent RNA helicase DDX3X OS=Homo sapiens OX=9606 GN=DDX3X PE=1 SV=3 |
| 118 | 63.44 | 11 | 8 | 38497 | 531 | >sp\|P40227\|TCPZ_HUMAN T-complex protein 1 subunit zeta OS=Homo sapiens OX=9606 GN=CCT6A PE=1 SV=3 |
| 119 | 63.20 | 10 | 10 | 4021 | 641 | >sp\|P08133-2\|ANXA6_HUMAN Isoform 2 of Annexin A6 OS=Homo sapiens OX=9606 GN=ANXA6 |
| 120 | 62.50 | 8 | 7 | 26655 | 1985 | >sp\|P35580-5\|MYH10_HUMAN Isoform 5 of Myosin-10 OS=Homo sapiens OX=9606 GN=MYH10 |
| 121 | 62.32 | 5 | 5 | 7703 | 375 | >sp\|P35659\|DEK_HUMAN Protein DEK OS=Homo sapiens OX=9606 GN=DEK PE=1 SV=1 |
| 122 | 62.29 | 7 | 7 | 9850 | 525 | >sp\|P14314-2\|GLU2B_HUMAN Isoform 2 of Glucosidase 2 subunit beta OS=Homo sapiens OX=9606 GN=PRKCSH |
| 123 | 62.20 | 9 | 8 | 36556 | 796 | >sp\|Q96QK1\|VPS35_HUMAN Vacuolar protein sorting-associated protein 35 OS=Homo sapiens OX=9606 GN=VPS35 PE=1 SV=2 |
| 124 | 62.14 | 9 | 9 | 22232 | 263 | >sp\|P62701\|RS4X_HUMAN 40S ribosomal protein S4; X isoform OS=Homo sapiens OX=9606 GN=RPS4X PE=1 SV=2 |
| 125 | 61.95 | 7 | 7 | 6941 | 1140 | >sp\|Q16531\|DDB1_HUMAN DNA damage-binding protein 1 OS=Homo sapiens OX=9606 GN=DDB1 PE=1 SV=1 |
| 126 | 61.63 | 15 | 13 | 27434 | 417 | >sp\|P00558\|PGK1_HUMAN Phosphoglycerate kinase 1 OS=Homo sapiens OX=9606 GN=PGK1 PE=1 SV=3 |
| 127 | 61.30 | 8 | 6 | 24904 | 249 | >sp\|P39687\|AN32A_HUMAN Acidic leucine-rich nuclear phosphoprotein 32 family member A OS=Homo sapiens OX=9606 GN=ANP32A PE=1 SV= |
| 128 | 61.27 | 10 | 9 | 19303 | 293 | >sp\|P15880\|RS2_HUMAN 40S ribosomal protein S2 OS=Homo sapiens OX=9606 GN=RPS2 PE=1 SV=2 |
| 129 | 61.09 | 10 | 9 | 21038 | 3224 | >sp\|P49792\|RBP2_HUMAN E3 SUMO-protein ligase RanBP2 OS=Homo sapiens OX=9606 GN=RANBP2 PE=1 SV=2 |
| 130 | 60.96 | 8 | 6 | 19342 | 463 | >sp\|Q9Y230\|RUVB2_HUMAN RuvB-like 2 OS=Homo sapiens OX=9606 GN=RUVBL2 PE=1 SV=3 |
| 131 | 60.72 | 5 | 5 | 34464 | 245 | >sp\|P63104\|1433Z_HUMAN 14-3-3 protein zeta/delta OS=Homo sapiens OX=9606 GN=YWHAZ PE=1 SV=1 |
| 132 | 60.67 | 5 | 5 | 26002 | 296 | >sp\|P40926-2\|MDHM_HUMAN Isoform 2 of Malate dehydrogenase; mitochondrial OS=Homo sapiens OX=9606 GN=MDH2 |
| 133 | 60.63 | 10 | 9 | 38916 | 337 | >sp\|P37837\|TALDO_HUMAN Transaldolase OS=Homo sapiens OX=9606 GN=TALDO1 PE=1 SV=2 |
| 134 | 60.38 | 11 | 10 | 34627 | 1512 | >sp\|P07814\|SYEP_HUMAN Bifunctional glutamate/proline--tRNA ligase OS=Homo sapiens OX=9606 GN=EPRS PE=1 SV=5 |
| 135 | 59.91 | 8 | 7 | 35063 | 539 | >sp\|P50991\|TCPD_HUMAN T-complex protein 1 subunit delta OS=Homo sapiens OX=9606 GN=CCT4 PE=1 SV=4 |
| 136 | 59.71 | 7 | 7 | 12514 | 636 | >sp\|O43390-2\|HNRPR_HUMAN Isoform 2 of Heterogeneous nuclear ribonucleoprotein R OS=Homo sapiens OX=9606 GN=HNRNPR |
| 137 | 59.57 | 7 | 6 | 1459 | 602 | >sp\|Q07065\|CKAP4_HUMAN Cytoskeleton-associated protein 4 OS=Homo sapiens OX=9606 GN=CKAP4 PE=1 SV=2 |
| 138 | 59.45 | 6 | 4 | 22777 | 110 | >sp\|P06454-2\|PTMA_HUMAN Isoform 2 of Prothymosin alpha OS=Homo sapiens OX=9606 GN=PTMA |
| 139 | 59.32 | 10 | 8 | 23623 | 910 | >sp\|Q7KZF4\|SND1_HUMAN Staphylococcal nuclease domain-containing protein 1 OS=Homo sapiens OX=9606 GN=SND1 PE=1 SV=1 |
| 140 | 58.34 | 6 | 6 | 500 | 504 | >sp\|Q9BZZ5-2\|API5_HUMAN Isoform 2 of Apoptosis inhibitor 5 OS=Homo sapiens OX=9606 GN=API5 |
| 141 | 57.97 | 5 | 5 | 29283 | 398 | >sp\|P62195-2\|PRS8_HUMAN Isoform 2 of 26S proteasome regulatory subunit 8 OS=Homo sapiens OX=9606 GN=PSMC5 |
| 142 | 57.96 | 11 | 11 | 8085 | 2871 | >sp\|P15924\|DESP_HUMAN Desmoplakin OS=Homo sapiens OX=9606 GN=DSP PE=1 SV=3 |
| 143 | 57.89 | 9 | 9 | 15101 | 1657 | >sp\|P46940\|IQGA1_HUMAN Ras GTPase-activating-like protein IQGAP1 OS=Homo sapiens OX=9606 GN=IQGAP1 PE=1 SV=1 |
| 144 | 57.47 | 10 | 8 | 29243 | 199 | >sp\|Q06830\|PRDX1_HUMAN Peroxiredoxin-1 OS=Homo sapiens OX=9606 GN=PRDX1 PE=1 SV=1 |
| 145 | 56.86 | 7 | 7 | 663 | 992 | >sp\|P05023-3\|AT1A1_HUMAN Isoform 3 of Sodium/potassium-transporting ATPase subunit alpha-1 OS=Homo sapiens OX=9606 GN=ATP1A1 |
| 146 | 56.82 | 10 | 10 | 28888 | 919 | >sp\|P55786\|PSA_HUMAN Puromycin-sensitive aminopeptidase OS=Homo sapiens OX=9606 GN=NPEPPS PE=1 SV=2 |
| 147 | 56.65 | 8 | 7 | 4295 | 627 | >sp\|P27824-2\|CALX_HUMAN Isoform 2 of Calnexin OS=Homo sapiens OX=9606 GN=CANX |
| 148 | 56.51 | 8 | 8 | 34612 | 298 | >sp\|P05141\|ADT2_HUMAN ADP/ATP translocase 2 OS=Homo sapiens OX=9606 GN=SLC25A5 PE=1 SV=7 |
| 149 | 56.21 | 9 | 8 | 9596 | 483 | >sp\|P34897-3\|GLYM_HUMAN Isoform 3 of Serine hydroxymethyltransferase; mitochondrial OS=Homo sapiens OX=9606 GN=SHMT2 |
| 150 | 56.14 | 9 | 8 | 7432 | 487 | >sp\|P26641-2\|EF1G_HUMAN Isoform 2 of Elongation factor 1-gamma OS=Homo sapiens OX=9606 GN=EEF1G |
| 151 | 55.99 | 8 | 7 | 31825 | 790 | >sp\|P49321-3\|NASP_HUMAN Isoform 3 of Nuclear autoantigenic sperm protein OS=Homo sapiens OX=9606 GN=NASP |
| 152 | 55.99 | 10 | 8 | 3615 | 1101 | >sp\|P53396\|ACLY_HUMAN ATP-citrate synthase OS=Homo sapiens OX=9606 GN=ACLY PE=1 SV=3 |
| 153 | 55.82 | 7 | 7 | 41398 | 1066 | >sp\|P18206-2\|VINC_HUMAN Isoform 1 of Vinculin OS=Homo sapiens OX=9606 GN=VCL |
| 154 | 55.46 | 7 | 6 | 29326 | 165 | >sp\|P62937\|PPIA_HUMAN Peptidyl-prolyl cis-trans isomerase A OS=Homo sapiens OX=9606 GN=PPIA PE=1 SV=2 |

| | | | | | | |
|---|---|---|---|---|---|---|
| 155 | 53.64 | 7 | 7 | 35782 | 756 | >sp\|P26639-2\|SYTC_HUMAN Isoform 2 of Threonine--tRNA ligase; cytoplasmic OS=Homo sapiens OX=9606 GN=TARS |
| 156 | 53.63 | 10 | 9 | 18698 | 382 | >sp\|Q15233-2\|NONO_HUMAN Isoform 2 of Non-POU domain-containing octamer-binding protein OS=Homo sapiens OX=9606 GN=NONO |
| 157 | 53.38 | 8 | 7 | 31862 | 267 | >sp\|P22392-2\|NDKB_HUMAN Isoform 3 of Nucleoside diphosphate kinase B OS=Homo sapiens OX=9606 GN=NME2 |
| 158 | 53.35 | 7 | 6 | 29308 | 330 | >sp\|P62136\|PP1A_HUMAN Serine/threonine-protein phosphatase PP1-alpha catalytic subunit OS=Homo sapiens OX=9606 GN=PPP1CA PE=1 S |
| 159 | 53.26 | 7 | 5 | 13900 | 210 | >sp\|P09211\|GSTP1_HUMAN Glutathione S-transferase P OS=Homo sapiens OX=9606 GN=GSTP1 PE=1 SV=2 |
| 160 | 52.78 | 10 | 10 | 41253 | 915 | >sp\|P55060-4\|XPO2_HUMAN Isoform 4 of Exportin-2 OS=Homo sapiens OX=9606 GN=CSE1L |
| 161 | 52.68 | 5 | 5 | 34390 | 245 | >sp\|P27348\|1433T_HUMAN 14-3-3 protein theta OS=Homo sapiens OX=9606 GN=YWHAQ PE=1 SV=1 |
| 162 | 52.49 | 4 | 4 | 18702 | 386 | >sp\|Q99733-2\|NP1L4_HUMAN Isoform 2 of Nucleosome assembly protein 1-like 4 OS=Homo sapiens OX=9606 GN=NAP1L4 |
| 163 | 52.39 | 8 | 7 | 10547 | 914 | >sp\|B5ME19\|EIFCL_HUMAN Eukaryotic translation initiation factor 3 subunit C-like protein OS=Homo sapiens OX=9606 GN=EIF3CL PE=3 |
| 164 | 52.13 | 7 | 7 | 35375 | 955 | >sp\|Q9Y2W1\|TR150_HUMAN Thyroid hormone receptor-associated protein 3 OS=Homo sapiens OX=9606 GN=THRAP3 PE=1 SV=2 |
| 165 | 51.97 | 6 | 5 | 1226 | 166 | >sp\|P23528\|COF1_HUMAN Cofilin-1 OS=Homo sapiens OX=9606 GN=CFL1 PE=1 SV=3 |
| 166 | 51.84 | 11 | 11 | 36277 | 513 | >sp\|Q14247-3\|SRC8_HUMAN Isoform 3 of Src substrate cortactin OS=Homo sapiens OX=9606 GN=CTTN |
| 167 | 51.68 | 5 | 5 | 17245 | 353 | >sp\|Q15717-2\|ELAV1_HUMAN Isoform 2 of ELAV-like protein 1 OS=Homo sapiens OX=9606 GN=ELAVL1 |
| 168 | 51.64 | 7 | 7 | 21113 | 1118 | >sp\|Q92900-2\|RENT1_HUMAN Isoform 2 of Regulator of nonsense transcripts 1 OS=Homo sapiens OX=9606 GN=UPF1 |
| 169 | 50.94 | 8 | 8 | 19840 | 1304 | >sp\|O75533\|SF3B1_HUMAN Splicing factor 3B subunit 1 OS=Homo sapiens OX=9606 GN=SF3B1 PE=1 SV=3 |
| 170 | 50.91 | 10 | 8 | 21381 | 248 | >sp\|P18124\|RL7_HUMAN 60S ribosomal protein L7 OS=Homo sapiens OX=9606 GN=RPL7 PE=1 SV=1 |
| 171 | 50.81 | 7 | 7 | 13927 | 371 | >sp\|O75367-3\|H2AY_HUMAN Isoform 3 of Core histone macro-H2A.1 OS=Homo sapiens OX=9606 GN=H2AFY |
| 172 | 50.79 | 10 | 8 | 20530 | 793 | >sp\|Q15459\|SF3A1_HUMAN Splicing factor 3A subunit 1 OS=Homo sapiens OX=9606 GN=SF3A1 PE=1 SV=1 |
| 173 | 50.70 | 7 | 7 | 21474 | 266 | >sp\|P62424\|RL7A_HUMAN 60S ribosomal protein L7a OS=Homo sapiens OX=9606 GN=RPL7A PE=1 SV=2 |
| 174 | 50.44 | 4 | 4 | 29951 | 332 | >sp\|P29966\|MARCS_HUMAN Myristoylated alanine-rich C-kinase substrate OS=Homo sapiens OX=9606 GN=MARCKS PE=1 SV=4 |
| 175 | 50.06 | 7 | 7 | 34654 | 660 | >sp\|P54136\|SYRC_HUMAN Arginine--tRNA ligase; cytoplasmic OS=Homo sapiens OX=9606 GN=RARS PE=1 SV=2 |
| 176 | 49.98 | 6 | 5 | 42254 | 248 | >sp\|P25788-2\|PSA3_HUMAN Isoform 2 of Proteasome subunit alpha type-3 OS=Homo sapiens OX=9606 GN=PSMA3 |
| 177 | 49.82 | 6 | 6 | 38830 | 1412 | >sp\|Q13428-8\|TCOF_HUMAN Isoform 8 of Treacle protein OS=Homo sapiens OX=9606 GN=TCOF1 |
| 178 | 49.75 | 9 | 9 | 27891 | 508 | >sp\|P07237\|PDIA1_HUMAN Protein disulfide-isomerase OS=Homo sapiens OX=9606 GN=P4HB PE=1 SV=3 |
| 179 | 49.71 | 11 | 10 | 30780 | 394 | >sp\|Q9UQ80\|PA2G4_HUMAN Proliferation-associated protein 2G4 OS=Homo sapiens OX=9606 GN=PA2G4 PE=1 SV=3 |
| 180 | 49.45 | 5 | 5 | 10934 | 873 | >sp\|P55884-2\|EIF3B_HUMAN Isoform 2 of Eukaryotic translation initiation factor 3 subunit B OS=Homo sapiens OX=9606 GN=EIF3B |
| 181 | 49.43 | 10 | 8 | 1462 | 2039 | >sp\|Q14008-3\|CKAP5_HUMAN Isoform 3 of Cytoskeleton-associated protein 5 OS=Homo sapiens OX=9606 GN=CKAP5 |
| 182 | 48.96 | 5 | 5 | 16303 | 611 | >sp\|P23588\|IF4B_HUMAN Eukaryotic translation initiation factor 4B OS=Homo sapiens OX=9606 GN=EIF4B PE=1 SV=2 |
| 183 | 48.94 | 10 | 10 | 26929 | 4684 | >sp\|Q15149\|PLEC_HUMAN Plectin OS=Homo sapiens OX=9606 GN=PLEC PE=1 SV=3 |
| 184 | 48.77 | 7 | 6 | 18778 | 265 | >sp\|P06748-2\|NPM_HUMAN Isoform 2 of Nucleophosmin OS=Homo sapiens OX=9606 GN=NPM1 |
| 185 | 48.51 | 9 | 7 | 15672 | 525 | >sp\|P35637-2\|FUS_HUMAN Isoform Short of RNA-binding protein FUS OS=Homo sapiens OX=9606 GN=FUS |
| 186 | 48.27 | 7 | 7 | 33358 | 853 | >sp\|P25205-2\|MCM3_HUMAN Isoform 2 of DNA replication licensing factor MCM3 OS=Homo sapiens OX=9606 GN=MCM3 |
| 187 | 48.07 | 10 | 9 | 7600 | 795 | >sp\|O43143\|DHX15_HUMAN Pre-mRNA-splicing factor ATP-dependent RNA helicase DHX15 OS=Homo sapiens OX=9606 GN=DHX15 PE=1 SV=2 |
| 188 | 47.80 | 5 | 5 | 12561 | 346 | >sp\|P31942\|HNRH3_HUMAN Heterogeneous nuclear ribonucleoprotein H3 OS=Homo sapiens OX=9606 GN=HNRNPH3 PE=1 SV=2 |
| 189 | 47.43 | 7 | 5 | 14985 | 529 | >sp\|P52292\|IMA1_HUMAN Importin subunit alpha-1 OS=Homo sapiens OX=9606 GN=KPNA2 PE=1 SV=1 |
| 190 | 47.35 | 6 | 6 | 36094 | 363 | >sp\|Q9Y3F4-2\|STRAP_HUMAN Isoform 2 of Serine-threonine kinase receptor-associated protein OS=Homo sapiens OX=9606 GN=STRAP |
| 191 | 47.34 | 8 | 5 | 36287 | 248 | >sp\|Q07955\|SRSF1_HUMAN Serine/arginine-rich splicing factor 1 OS=Homo sapiens OX=9606 GN=SRSF1 PE=1 SV=2 |
| 192 | 47.11 | 10 | 10 | 37885 | 1018 | >sp\|P22314-2\|UBA1_HUMAN Isoform 2 of Ubiquitin-like modifier-activating enzyme 1 OS=Homo sapiens OX=9606 GN=UBA1 |
| 193 | 47.10 | 5 | 4 | 12190 | 221 | >sp\|P16402\|H13_HUMAN Histone H1.3 OS=Homo sapiens OX=9606 GN=HIST1H1D PE=1 SV=2 |
| 194 | 47.07 | 4 | 4 | 20411 | 533 | >sp\|O43175\|SERA_HUMAN D-3-phosphoglycerate dehydrogenase OS=Homo sapiens OX=9606 GN=PHGDH PE=1 SV=4 |
| 195 | 46.87 | 5 | 5 | 6960 | 715 | >sp\|Q9NR30-2\|DDX21_HUMAN Isoform 2 of Nucleolar RNA helicase 2 OS=Homo sapiens OX=9606 GN=DDX21 |
| 196 | 46.64 | 7 | 7 | 22188 | 145 | >sp\|P39019\|RS19_HUMAN 40S ribosomal protein S19 OS=Homo sapiens OX=9606 GN=RPS19 PE=1 SV=2 |
| 197 | 46.49 | 12 | 11 | 29644 | 1394 | >sp\|P42704\|LPPRC_HUMAN Leucine-rich PPR motif-containing protein; mitochondrial OS=Homo sapiens OX=9606 GN=LRPPRC PE=1 SV=3 |
| 198 | 46.42 | 5 | 4 | 28454 | 129 | >sp\|O14737-2\|PDCD5_HUMAN Isoform 2 of Programmed cell death protein 5 OS=Homo sapiens OX=9606 GN=PDCD5 |
| 199 | 46.09 | 4 | 4 | 19305 | 208 | >sp\|P62241\|RS8_HUMAN 40S ribosomal protein S8 OS=Homo sapiens OX=9606 GN=RPS8 PE=1 SV=2 |
| 200 | 45.94 | 4 | 4 | 32839 | 396 | >sp\|Q9NTK5\|OLA1_HUMAN Obg-like ATPase 1 OS=Homo sapiens OX=9606 GN=OLA1 PE=1 SV=2 |
| 201 | 45.91 | 5 | 4 | 22565 | 102 | >sp\|P20962\|PTMS_HUMAN Parathymosin OS=Homo sapiens OX=9606 GN=PTMS PE=1 SV=2 |
| 202 | 45.90 | 6 | 5 | 3320 | 144 | >sp\|P61204-2\|ARF3_HUMAN Isoform 2 of ADP-ribosylation factor 3 OS=Homo sapiens OX=9606 GN=ARF3 |
| 203 | 45.76 | 7 | 7 | 31489 | 866 | >sp\|Q9BXJ9\|NAA15_HUMAN N-alpha-acetyltransferase 15; NatA auxiliary subunit OS=Homo sapiens OX=9606 GN=NAA15 PE=1 SV=1 |
| 204 | 45.64 | 8 | 7 | 30993 | 793 | >sp\|P54886-2\|P5CS_HUMAN Isoform Short of Delta-1-pyrroline-5-carboxylate synthase OS=Homo sapiens OX=9606 GN=ALDH18A1 |
| 205 | 45.50 | 8 | 8 | 21386 | 288 | >sp\|Q02878\|RL6_HUMAN 60S ribosomal protein L6 OS=Homo sapiens OX=9606 GN=RPL6 PE=1 SV=3 |
| 206 | 44.71 | 8 | 8 | 15545 | 569 | >sp\|P06744-2\|G6PI_HUMAN Isoform 2 of Glucose-6-phosphate isomerase OS=Homo sapiens OX=9606 GN=GPI |
| 207 | 44.40 | 5 | 5 | 33737 | 863 | >sp\|P33991\|MCM4_HUMAN DNA replication licensing factor MCM4 OS=Homo sapiens OX=9606 GN=MCM4 PE=1 SV=5 |
| 208 | 44.09 | 5 | 5 | 21697 | 332 | >sp\|Q99729\|ROAA_HUMAN Heterogeneous nuclear ribonucleoprotein A/B OS=Homo sapiens OX=9606 GN=HNRNPAB PE=1 SV=2 |
| 209 | 43.94 | 10 | 10 | 22922 | 1217 | >sp\|Q9UQE7\|SMC3_HUMAN Structural maintenance of chromosomes protein 3 OS=Homo sapiens OX=9606 GN=SMC3 PE=1 SV=2 |

| | | | | | | |
|---|---|---|---|---|---|---|
| 210 | 43.82 | 6 | 6 | 37646 | 2136 | >sp\|O75643\|U520_HUMAN U5 small nuclear ribonucleoprotein 200 kDa helicase OS=Homo sapiens OX=9606 GN=SNRNP200 PE=1 SV=2 |
| 211 | 43.63 | 7 | 6 | 7279 | 443 | >sp\|Q13838-2\|DX39B_HUMAN Isoform 2 of Spliceosome RNA helicase DDX39B OS=Homo sapiens OX=9606 GN=DDX39B |
| 212 | 43.59 | 6 | 4 | 6908 | 494 | >sp\|Q99615\|DNJC7_HUMAN DnaJ homolog subfamily C member 7 OS=Homo sapiens OX=9606 GN=DNAJC7 PE=1 SV=2 |
| 213 | 43.41 | 4 | 4 | 11889 | 1038 | >sp\|O95373\|IPO7_HUMAN Importin-7 OS=Homo sapiens OX=9606 GN=IPO7 PE=1 SV=1 |
| 214 | 43.21 | 6 | 5 | 29443 | 271 | >sp\|Q13162\|PRDX4_HUMAN Peroxiredoxin-4 OS=Homo sapiens OX=9606 GN=PRDX4 PE=1 SV=1 |
| 215 | 43.14 | 5 | 5 | 11102 | 493 | >sp\|Q16658\|FSCN1_HUMAN Fascin OS=Homo sapiens OX=9606 GN=FSCN1 PE=1 SV=3 |
| 216 | 43.10 | 10 | 9 | 11605 | 876 | >sp\|Q14974\|IMB1_HUMAN Importin subunit beta-1 OS=Homo sapiens OX=9606 GN=KPNB1 PE=1 SV=2 |
| 217 | 43.00 | 7 | 6 | 38243 | 129 | >sp\|O75347-2\|TBCA_HUMAN Isoform 2 of Tubulin-specific chaperone A OS=Homo sapiens OX=9606 GN=TBCA |
| 218 | 42.66 | 4 | 3 | 42093 | 456 | >sp\|P30520\|PURA2_HUMAN Adenylosuccinate synthetase isozyme 2 OS=Homo sapiens OX=9606 GN=ADSS PE=1 SV=3 |
| 219 | 42.28 | 8 | 7 | 1068 | 1233 | >sp\|P53621-2\|COPA_HUMAN Isoform 2 of Coatomer subunit alpha OS=Homo sapiens OX=9606 GN=COPA |
| 220 | 42.26 | 5 | 5 | 21879 | 217 | >sp\|P62906\|RL10A_HUMAN 60S ribosomal protein L10a OS=Homo sapiens OX=9606 GN=RPL10A PE=1 SV=2 |
| 221 | 42.08 | 5 | 5 | 32755 | 1332 | >sp\|O75694-2\|NU155_HUMAN Isoform 2 of Nuclear pore complex protein Nup155 OS=Homo sapiens OX=9606 GN=NUP155 |
| 222 | 41.77 | 5 | 5 | 18951 | 587 | >sp\|P46060\|RAGP1_HUMAN Ran GTPase-activating protein 1 OS=Homo sapiens OX=9606 GN=RANGAP1 PE=1 SV=1 |
| 223 | 41.29 | 5 | 4 | 22857 | 418 | >sp\|P50454\|SERPH_HUMAN Serpin H1 OS=Homo sapiens OX=9606 GN=SERPINH1 PE=1 SV=2 |
| 224 | 41.17 | 3 | 3 | 26853 | 125 | >sp\|Q9NRX4\|PHP14_HUMAN 14 kDa phosphohistidine phosphatase OS=Homo sapiens OX=9606 GN=PHPT1 PE=1 SV=1 |
| 225 | 41.12 | 6 | 6 | 19310 | 249 | >sp\|P62753\|RS6_HUMAN 40S ribosomal protein S6 OS=Homo sapiens OX=9606 GN=RPS6 PE=1 SV=1 |
| 226 | 41.10 | 7 | 6 | 21332 | 257 | >sp\|P62917\|RL8_HUMAN 60S ribosomal protein L8 OS=Homo sapiens OX=9606 GN=RPL8 PE=1 SV=2 |
| 227 | 41.02 | 4 | 4 | 13269 | 432 | >sp\|O95232\|LC7L3_HUMAN Luc7-like protein 3 OS=Homo sapiens OX=9606 GN=LUC7L3 PE=1 SV=2 |
| 228 | 40.86 | 4 | 4 | 22726 | 432 | >sp\|P22234-2\|PUR6_HUMAN Isoform 2 of Multifunctional protein ADE2 OS=Homo sapiens OX=9606 GN=PAICS |
| 229 | 40.85 | 7 | 6 | 24592 | 357 | >sp\|P07355-2\|ANXA2_HUMAN Isoform 2 of Annexin A2 OS=Homo sapiens OX=9606 GN=ANXA2 |
| 230 | 40.82 | 8 | 8 | 19251 | 151 | >sp\|P62277\|RS13_HUMAN 40S ribosomal protein S13 OS=Homo sapiens OX=9606 GN=RPS13 PE=1 SV=2 |
| 231 | 40.65 | 7 | 7 | 11200 | 466 | >sp\|Q13283\|G3BP1_HUMAN Ras GTPase-activating protein-binding protein 1 OS=Homo sapiens OX=9606 GN=G3BP1 PE=1 SV=1 |
| 232 | 40.30 | 4 | 4 | 19842 | 1217 | >sp\|Q15393\|SF3B3_HUMAN Splicing factor 3B subunit 3 OS=Homo sapiens OX=9606 GN=SF3B3 PE=1 SV=4 |
| 233 | 40.25 | 6 | 6 | 24897 | 488 | >sp\|P28838-2\|AMPL_HUMAN Isoform 2 of Cytosol aminopeptidase OS=Homo sapiens OX=9606 GN=LAP3 |
| 234 | 40.20 | 4 | 4 | 39025 | 193 | >sp\|Q99426-2\|TBCB_HUMAN Isoform 2 of Tubulin-folding cofactor B OS=Homo sapiens OX=9606 GN=TBCB |
| 235 | 40.12 | 8 | 6 | 5190 | 417 | >sp\|P27797\|CALR_HUMAN Calreticulin OS=Homo sapiens OX=9606 GN=CALR PE=1 SV=1 |
| 236 | 39.42 | 5 | 5 | 29328 | 216 | >sp\|P23284\|PPIB_HUMAN Peptidyl-prolyl cis-trans isomerase B OS=Homo sapiens OX=9606 GN=PPIB PE=1 SV=2 |
| 237 | 39.22 | 8 | 8 | 36162 | 2364 | >sp\|Q01082\|SPTB2_HUMAN Spectrin beta chain; non-erythrocytic 1 OS=Homo sapiens OX=9606 GN=SPTBN1 PE=1 SV=2 |
| 238 | 39.09 | 6 | 5 | 13663 | 454 | >sp\|P42167\|LAP2B_HUMAN Lamina-associated polypeptide 2; isoforms beta/gamma OS=Homo sapiens OX=9606 GN=TMPO PE=1 SV=2 |
| 239 | 39.01 | 4 | 4 | 9191 | 369 | >sp\|P50502\|F10A1_HUMAN Hsc70-interacting protein OS=Homo sapiens OX=9606 GN=ST13 PE=1 SV=2 |
| 240 | 38.82 | 6 | 6 | 21803 | 159 | >sp\|Q07020-2\|RL18_HUMAN Isoform 2 of 60S ribosomal protein L18 OS=Homo sapiens OX=9606 GN=RPL18 |
| 241 | 38.80 | 6 | 6 | 42264 | 908 | >sp\|Q13200\|PSMD2_HUMAN 26S proteasome non-ATPase regulatory subunit 2 OS=Homo sapiens OX=9606 GN=PSMD2 PE=1 SV=3 |
| 242 | 38.70 | 4 | 3 | 19615 | 895 | >sp\|Q13435\|SF3B2_HUMAN Splicing factor 3B subunit 2 OS=Homo sapiens OX=9606 GN=SF3B2 PE=1 SV=2 |
| 243 | 38.62 | 5 | 5 | 24915 | 426 | >sp\|Q8WU90\|ZC3HF_HUMAN Zinc finger CCCH domain-containing protein 15 OS=Homo sapiens OX=9606 GN=ZC3H15 PE=1 SV=1 |
| 244 | 38.58 | 4 | 4 | 19252 | 135 | >sp\|P08708\|RS17_HUMAN 40S ribosomal protein S17 OS=Homo sapiens OX=9606 GN=RPS17 PE=1 SV=2 |
| 245 | 38.36 | 7 | 7 | 37154 | 489 | >sp\|P12081-4\|SYHC_HUMAN Isoform 4 of Histidine--tRNA ligase; cytoplasmic OS=Homo sapiens OX=9606 GN=HARS |
| 246 | 38.15 | 5 | 5 | 14631 | 577 | >sp\|Q9NZI8\|IF2B1_HUMAN Insulin-like growth factor 2 mRNA-binding protein 1 OS=Homo sapiens OX=9606 GN=IGF2BP1 PE=1 SV=2 |
| 247 | 38.11 | 6 | 6 | 25870 | 395 | >sp\|P31153\|METK2_HUMAN S-adenosylmethionine synthase isoform type-2 OS=Homo sapiens OX=9606 GN=MAT2A PE=1 SV=1 |
| 248 | 38.06 | 8 | 8 | 18965 | 390 | >sp\|Q09028-4\|RBBP4_HUMAN Isoform 4 of Histone-binding protein RBBP4 OS=Homo sapiens OX=9606 GN=RBBP4 |
| 249 | 37.88 | 6 | 6 | 21314 | 196 | >sp\|P84098\|RL19_HUMAN 60S ribosomal protein L19 OS=Homo sapiens OX=9606 GN=RPL19 PE=1 SV=1 |
| 250 | 37.76 | 4 | 4 | 573 | 399 | >sp\|P61160-2\|ARP2_HUMAN Isoform 2 of Actin-related protein 2 OS=Homo sapiens OX=9606 GN=ACTR2 |
| 251 | 37.69 | 5 | 5 | 1175 | 283 | >sp\|Q15417-3\|CNN3_HUMAN Isoform 3 of Calponin-3 OS=Homo sapiens OX=9606 GN=CNN3 |
| 252 | 37.61 | 8 | 6 | 4742 | 272 | >sp\|P47756-2\|CAPZB_HUMAN Isoform 2 of F-actin-capping protein subunit beta OS=Homo sapiens OX=9606 GN=CAPZB |
| 253 | 37.43 | 6 | 6 | 14930 | 184 | >sp\|P63241-2\|IF5A1_HUMAN Isoform 2 of Eukaryotic translation initiation factor 5A-1 OS=Homo sapiens OX=9606 GN=EIF5A |
| 254 | 37.41 | 6 | 6 | 29945 | 1135 | >sp\|P27816-6\|MAP4_HUMAN Isoform 6 of Microtubule-associated protein 4 OS=Homo sapiens OX=9606 GN=MAP4 |
| 255 | 37.39 | 9 | 8 | 626 | 586 | >sp\|Q9NVI7-2\|ATD3A_HUMAN Isoform 2 of ATPase family AAA domain-containing protein 3A OS=Homo sapiens OX=9606 GN=ATAD3A |
| 256 | 37.29 | 6 | 6 | 23656 | 2140 | >sp\|P18583-2\|SON_HUMAN Isoform A of Protein SON OS=Homo sapiens OX=9606 GN=SON |
| 257 | 37.01 | 6 | 6 | 235 | 472 | >sp\|P50995-2\|ANX11_HUMAN Isoform 2 of Annexin A11 OS=Homo sapiens OX=9606 GN=ANXA11 |
| 258 | 36.83 | 5 | 5 | 25834 | 904 | >sp\|P49736\|MCM2_HUMAN DNA replication licensing factor MCM2 OS=Homo sapiens OX=9606 GN=MCM2 PE=1 SV=4 |
| 259 | 36.78 | 3 | 3 | 21286 | 792 | >sp\|P23921\|RIR1_HUMAN Ribonucleoside-diphosphate reductase large subunit OS=Homo sapiens OX=9606 GN=RRM1 PE=1 SV=1 |
| 260 | 36.73 | 5 | 4 | 26541 | 620 | >sp\|Q03252\|LMNB2_HUMAN Lamin-B2 OS=Homo sapiens OX=9606 GN=LMNB2 PE=1 SV=4 |
| 261 | 36.66 | 8 | 7 | 16432 | 400 | >sp\|P08727\|K1C19_HUMAN Keratin; type I cytoskeletal 19 OS=Homo sapiens OX=9606 GN=KRT19 PE=1 SV=4 |
| 262 | 36.46 | 9 | 9 | 36772 | 732 | >sp\|P13010\|XRCC5_HUMAN X-ray repair cross-complementing protein 5 OS=Homo sapiens OX=9606 GN=XRCC5 PE=1 SV=3 |
| 263 | 35.95 | 3 | 3 | 33566 | 583 | >sp\|Q8N1G4\|LRC47_HUMAN Leucine-rich repeat-containing protein 47 OS=Homo sapiens OX=9606 GN=LRRC47 PE=1 SV=1 |
| 264 | 35.91 | 5 | 5 | 20209 | 370 | >sp\|Q9Y617\|SERC_HUMAN Phosphoserine aminotransferase OS=Homo sapiens OX=9606 GN=PSAT1 PE=1 SV=2 |
| 265 | 35.88 | 4 | 3 | 28847 | 158 | >sp\|P24666\|PPAC_HUMAN Low molecular weight phosphotyrosine protein phosphatase OS=Homo sapiens OX=9606 GN=ACP1 PE=1 SV=3 |
| 266 | 35.84 | 4 | 4 | 10792 | 600 | >sp\|Q01844-6\|EWS_HUMAN Isoform 6 of RNA-binding protein EWS OS=Homo sapiens OX=9606 GN=EWSR1 |
| 267 | 35.72 | 5 | 5 | 7017 | 738 | >sp\|O00429-8\|DNM1L_HUMAN Isoform 8 of Dynamin-1-like protein OS=Homo sapiens OX=9606 GN=DNM1L |

| | | | | | | |
|---|---|---|---|---|---|---|
| 268 | 35.70 | 3 | 3 | 36192 | 221 | >sp\|Q13242\|SRSF9_HUMAN Serine/arginine-rich splicing factor 9 OS=Homo sapiens OX=9606 GN=SRSF9 PE=1 SV=1 |
| 269 | 35.68 | 4 | 4 | 15950 | 418 | >sp\|Q07666-2\|KHDR1_HUMAN Isoform 2 of KH domain-containing; RNA-binding; signal transduction-associated protein 1 OS=Homo sapie |
| 270 | 35.48 | 5 | 5 | 4017 | 346 | >sp\|P04083\|ANXA1_HUMAN Annexin A1 OS=Homo sapiens OX=9606 GN=ANXA1 PE=1 SV=2 |
| 271 | 35.41 | 4 | 4 | 22345 | 216 | >sp\|P62826\|RAN_HUMAN GTP-binding nuclear protein Ran OS=Homo sapiens OX=9606 GN=RAN PE=1 SV=3 |
| 272 | 35.30 | 4 | 4 | 15783 | 572 | >sp\|Q96I24\|FUBP3_HUMAN Far upstream element-binding protein 3 OS=Homo sapiens OX=9606 GN=FUBP3 PE=1 SV=2 |
| 273 | 35.30 | 4 | 4 | 35547 | 1264 | >sp\|P26640\|SYVC_HUMAN Valine--tRNA ligase OS=Homo sapiens OX=9606 GN=VARS PE=1 SV=4 |
| 274 | 35.02 | 4 | 4 | 24830 | 338 | >sp\|O95433\|AHSA1_HUMAN Activator of 90 kDa heat shock protein ATPase homolog 1 OS=Homo sapiens OX=9606 GN=AHSA1 PE=1 SV=1 |
| 275 | 35.00 | 7 | 5 | 34873 | 1262 | >sp\|P41252\|SYIC_HUMAN Isoleucine--tRNA ligase; cytoplasmic OS=Homo sapiens OX=9606 GN=IARS PE=1 SV=2 |
| 276 | 34.76 | 5 | 5 | 6405 | 297 | >sp\|P06493\|CDK1_HUMAN Cyclin-dependent kinase 1 OS=Homo sapiens OX=9606 GN=CDK1 PE=1 SV=3 |
| 277 | 34.69 | 9 | 7 | 22229 | 243 | >sp\|P23396\|RS3_HUMAN 40S ribosomal protein S3 OS=Homo sapiens OX=9606 GN=RPS3 PE=1 SV=2 |
| 278 | 34.68 | 3 | 3 | 35309 | 427 | >sp\|P24752\|THIL_HUMAN Acetyl-CoA acetyltransferase; mitochondrial OS=Homo sapiens OX=9606 GN=ACAT1 PE=1 SV=1 |
| 279 | 34.64 | 3 | 3 | 28589 | 488 | >sp\|Q15084-5\|PDIA6_HUMAN Isoform 5 of Protein disulfide-isomerase A6 OS=Homo sapiens OX=9606 GN=PDIA6 |
| 280 | 34.61 | 7 | 5 | 12364 | 130 | >sp\|P0C0S8\|H2A1_HUMAN Histone H2A type 1 OS=Homo sapiens OX=9606 GN=HIST1H2AG PE=1 SV=2 |
| 281 | 34.48 | 7 | 7 | 17440 | 445 | >sp\|P60228\|EIF3E_HUMAN Eukaryotic translation initiation factor 3 subunit E OS=Homo sapiens OX=9606 GN=EIF3E PE=1 SV=1 |
| 282 | 34.44 | 4 | 3 | 11014 | 224 | >sp\|Q00688\|FKBP3_HUMAN Peptidyl-prolyl cis-trans isomerase FKBP3 OS=Homo sapiens OX=9606 GN=FKBP3 PE=1 SV=1 |
| 283 | 34.36 | 8 | 7 | 16626 | 289 | >sp\|Q15181\|IPYR_HUMAN Inorganic pyrophosphatase OS=Homo sapiens OX=9606 GN=PPA1 PE=1 SV=2 |
| 284 | 34.30 | 4 | 4 | 6433 | 102 | >sp\|P61604\|CH10_HUMAN 10 kDa heat shock protein; mitochondrial OS=Homo sapiens OX=9606 GN=HSPE1 PE=1 SV=2 |
| 285 | 34.23 | 7 | 6 | 11385 | 126 | >sp\|P06899\|H2B1J_HUMAN Histone H2B type 1-J OS=Homo sapiens OX=9606 GN=HIST1H2BJ PE=1 SV=3 |
| 286 | 34.21 | 7 | 6 | 10904 | 290 | >sp\|P30084\|ECHM_HUMAN Enoyl-CoA hydratase; mitochondrial OS=Homo sapiens OX=9606 GN=ECHS1 PE=1 SV=4 |
| 287 | 34.14 | 6 | 6 | 11626 | 1115 | >sp\|O00410-3\|IPO5_HUMAN Isoform 3 of Importin-5 OS=Homo sapiens OX=9606 GN=IPO5 |
| 288 | 33.86 | 6 | 4 | 18934 | 249 | >sp\|P51148-2\|RAB5C_HUMAN Isoform 2 of Ras-related protein Rab-5C OS=Homo sapiens OX=9606 GN=RAB5C |
| 289 | 33.80 | 5 | 5 | 9589 | 335 | >sp\|O76003\|GLRX3_HUMAN Glutaredoxin-3 OS=Homo sapiens OX=9606 GN=GLRX3 PE=1 SV=2 |
| 290 | 33.75 | 3 | 3 | 29289 | 248 | >sp\|O14818\|PSA7_HUMAN Proteasome subunit alpha type-7 OS=Homo sapiens OX=9606 GN=PSMA7 PE=1 SV=1 |
| 291 | 33.68 | 4 | 4 | 17671 | 536 | >sp\|Q16555-2\|DPYL2_HUMAN Isoform 2 of Dihydropyrimidinase-related protein 2 OS=Homo sapiens OX=9606 GN=DPYSL2 |
| 292 | 33.61 | 3 | 3 | 21195 | 461 | >sp\|P13489\|RINI_HUMAN Ribonuclease inhibitor OS=Homo sapiens OX=9606 GN=RNH1 PE=1 SV=2 |
| 293 | 33.61 | 7 | 7 | 25810 | 1332 | >sp\|Q9BQG0-2\|MBB1A_HUMAN Isoform 2 of Myb-binding protein 1A OS=Homo sapiens OX=9606 GN=MYBBP1A |
| 294 | 33.48 | 6 | 5 | 15310 | 739 | >sp\|P41250\|GARS_HUMAN Glycine--tRNA ligase OS=Homo sapiens OX=9606 GN=GARS PE=1 SV=3 |
| 295 | 33.47 | 6 | 6 | 25642 | 591 | >sp\|P17812\|PYRG1_HUMAN CTP synthase 1 OS=Homo sapiens OX=9606 GN=CTPS1 PE=1 SV=2 |
| 296 | 33.31 | 11 | 10 | 19307 | 152 | >sp\|P62269\|RS18_HUMAN 40S ribosomal protein S18 OS=Homo sapiens OX=9606 GN=RPS18 PE=1 SV=3 |
| 297 | 33.27 | 4 | 4 | 4740 | 474 | >sp\|Q01518-2\|CAP1_HUMAN Isoform 2 of Adenylyl cyclase-associated protein 1 OS=Homo sapiens OX=9606 GN=CAP1 |
| 298 | 33.19 | 6 | 6 | 21329 | 403 | >sp\|P39023\|RL3_HUMAN 60S ribosomal protein L3 OS=Homo sapiens OX=9606 GN=RPL3 PE=1 SV=2 |
| 299 | 33.10 | 5 | 5 | 21378 | 165 | >sp\|P30050\|RL12_HUMAN 60S ribosomal protein L12 OS=Homo sapiens OX=9606 GN=RPL12 PE=1 SV=1 |
| 300 | 33.08 | 4 | 4 | 38831 | 127 | >sp\|P53999\|TCP4_HUMAN Activated RNA polymerase II transcriptional coactivator p15 OS=Homo sapiens OX=9606 GN=SUB1 PE=1 SV=3 |
| 301 | 32.95 | 9 | 7 | 19311 | 194 | >sp\|P46781\|RS9_HUMAN 40S ribosomal protein S9 OS=Homo sapiens OX=9606 GN=RPS9 PE=1 SV=3 |
| 302 | 32.88 | 3 | 3 | 41065 | 229 | >sp\|O43399-7\|TPD54_HUMAN Isoform 7 of Tumor protein D54 OS=Homo sapiens OX=9606 GN=TPD52L2 |
| 303 | 32.87 | 3 | 3 | 38667 | 246 | >sp\|Q00059\|TFAM_HUMAN Transcription factor A; mitochondrial OS=Homo sapiens OX=9606 GN=TFAM PE=1 SV=1 |
| 304 | 32.69 | 3 | 3 | 22096 | 295 | >sp\|P08865\|RSSA_HUMAN 40S ribosomal protein SA OS=Homo sapiens OX=9606 GN=RPSA PE=1 SV=4 |
| 305 | 32.65 | 5 | 4 | 4409 | 277 | >sp\|P16152\|CBR1_HUMAN Carbonyl reductase [NADPH] 1 OS=Homo sapiens OX=9606 GN=CBR1 PE=1 SV=3 |
| 306 | 32.63 | 6 | 6 | 34835 | 502 | >sp\|O95793-3\|STAU1_HUMAN Isoform 3 of Double-stranded RNA-binding protein Staufen homolog 1 OS=Homo sapiens OX=9606 GN=STAU1 |
| 307 | 32.61 | 7 | 5 | 9361 | 282 | >sp\|P10768\|ESTD_HUMAN S-formylglutathione hydrolase OS=Homo sapiens OX=9606 GN=ESD PE=1 SV=2 |
| 308 | 32.61 | 3 | 3 | 33005 | 352 | >sp\|P40925-3\|MDHC_HUMAN Isoform 3 of Malate dehydrogenase; cytoplasmic OS=Homo sapiens OX=9606 GN=MDH1 |
| 309 | 32.61 | 5 | 4 | 16999 | 346 | >sp\|P38117-2\|ETFB_HUMAN Isoform 2 of Electron transfer flavoprotein subunit beta OS=Homo sapiens OX=9606 GN=ETFB |
| 310 | 32.54 | 6 | 6 | 21306 | 204 | >sp\|P61313\|RL15_HUMAN 60S ribosomal protein L15 OS=Homo sapiens OX=9606 GN=RPL15 PE=1 SV=2 |
| 311 | 32.53 | 4 | 4 | 2503 | 169 | >sp\|P13073\|COX41_HUMAN Cytochrome c oxidase subunit 4 isoform 1; mitochondrial OS=Homo sapiens OX=9606 GN=COX4I1 PE=1 SV=1 |
| 312 | 32.25 | 4 | 4 | 21004 | 453 | >sp\|P22695\|QCR2_HUMAN Cytochrome b-c1 complex subunit 2; mitochondrial OS=Homo sapiens OX=9606 GN=UQCRC2 PE=1 SV=3 |
| 313 | 32.19 | 4 | 4 | 17443 | 516 | >sp\|Q9Y262-2\|EIF3L_HUMAN Isoform 2 of Eukaryotic translation initiation factor 3 subunit L OS=Homo sapiens OX=9606 GN=EIF3L |
| 314 | 32.14 | 7 | 5 | 21380 | 297 | >sp\|P46777\|RL5_HUMAN 60S ribosomal protein L5 OS=Homo sapiens OX=9606 GN=RPL5 PE=1 SV=3 |
| 315 | 32.11 | 4 | 4 | 19292 | 146 | >sp\|P62249\|RS16_HUMAN 40S ribosomal protein S16 OS=Homo sapiens OX=9606 GN=RPS16 PE=1 SV=2 |
| 316 | 31.98 | 3 | 3 | 34462 | 246 | >sp\|P31946\|1433B_HUMAN 14-3-3 protein beta/alpha OS=Homo sapiens OX=9606 GN=YWHAB PE=1 SV=3 |
| 317 | 31.62 | 4 | 4 | 31975 | 239 | >sp\|Q9UKD2\|MRT4_HUMAN mRNA turnover protein 4 homolog OS=Homo sapiens OX=9606 GN=MRTO4 PE=1 SV=2 |
| 318 | 31.49 | 6 | 5 | 37757 | 223 | >sp\|P09936\|UCHL1_HUMAN Ubiquitin carboxyl-terminal hydrolase isozyme L1 OS=Homo sapiens OX=9606 GN=UCHL1 PE=1 SV=2 |
| 319 | 31.30 | 4 | 4 | 15591 | 204 | >sp\|P52565\|GDIR1_HUMAN Rho GDP-dissociation inhibitor 1 OS=Homo sapiens OX=9606 GN=ARHGDIA PE=1 SV=3 |
| 320 | 31.22 | 4 | 4 | 23487 | 1105 | >sp\|Q92922\|SMRC1_HUMAN SWI/SNF complex subunit SMARCC1 OS=Homo sapiens OX=9606 GN=SMARCC1 PE=1 SV=3 |
| 321 | 31.20 | 4 | 3 | 19565 | 616 | >sp\|P31040-2\|SDHA_HUMAN Isoform 2 of Succinate dehydrogenase [ubiquinone] flavoprotein subunit; mitochondrial OS=Homo sapiens O |
| 322 | 31.18 | 4 | 4 | 34468 | 589 | >sp\|P30153\|2AAA_HUMAN Serine/threonine-protein phosphatase 2A 65 kDa regulatory subunit A alpha isoform OS=Homo sapiens OX=9606 |
| 323 | 31.08 | 8 | 7 | 12285 | 756 | >sp\|Q9BUJ2-4\|HNRL1_HUMAN Isoform 4 of Heterogeneous nuclear ribonucleoprotein U-like protein 1 OS=Homo sapiens OX=9606 GN=HNRNP |
| 324 | 31.02 | 4 | 4 | 21356 | 135 | >sp\|P62910\|RL32_HUMAN 60S ribosomal protein L32 OS=Homo sapiens OX=9606 GN=RPL32 PE=1 SV=2 |

<table>
<tr><td>325</td><td>30.97</td><td>4</td><td>3</td><td>23408</td><td>120</td><td>>sp|P62318-2|SMD3_HUMAN Isoform 2 of Small nuclear ribonucleoprotein Sm D3 OS=Homo sapiens OX=9606 GN=SNRPD3</td></tr>
<tr><td>326</td><td>30.91</td><td>5</td><td>4</td><td>21357</td><td>70</td><td>>sp|P63173|RL38_HUMAN 60S ribosomal protein L38 OS=Homo sapiens OX=9606 GN=RPL38 PE=1 SV=2</td></tr>
<tr><td>327</td><td>30.89</td><td>5</td><td>3</td><td>13299</td><td>1329</td><td>>sp|Q86UP2-4|KTN1_HUMAN Isoform 4 of Kinectin OS=Homo sapiens OX=9606 GN=KTN1</td></tr>
<tr><td>328</td><td>30.81</td><td>4</td><td>4</td><td>31331</td><td>819</td><td>>sp|Q8N1F7|NUP93_HUMAN Nuclear pore complex protein Nup93 OS=Homo sapiens OX=9606 GN=NUP93 PE=1 SV=2</td></tr>
<tr><td>329</td><td>30.77</td><td>5</td><td>4</td><td>17478</td><td>261</td><td>>sp|P30040|ERP29_HUMAN Endoplasmic reticulum resident protein 29 OS=Homo sapiens OX=9606 GN=ERP29 PE=1 SV=4</td></tr>
<tr><td>330</td><td>30.71</td><td>3</td><td>3</td><td>14105</td><td>1293</td><td>>sp|Q6Y7W6-5|GGYF2_HUMAN Isoform 4 of GRB10-interacting GYF protein 2 OS=Homo sapiens OX=9606 GN=GIGYF2</td></tr>
<tr><td>331</td><td>30.63</td><td>5</td><td>5</td><td>8044</td><td>622</td><td>>sp|Q92499-2|DDX1_HUMAN Isoform 2 of ATP-dependent RNA helicase DDX1 OS=Homo sapiens OX=9606 GN=DDX1</td></tr>
<tr><td>332</td><td>30.62</td><td>6</td><td>5</td><td>32061</td><td>577</td><td>>sp|P26038|MOES_HUMAN Moesin OS=Homo sapiens OX=9606 GN=MSN PE=1 SV=3</td></tr>
<tr><td>333</td><td>30.51</td><td>3</td><td>3</td><td>36065</td><td>709</td><td>>sp|Q08945|SSRP1_HUMAN FACT complex subunit SSRP1 OS=Homo sapiens OX=9606 GN=SSRP1 PE=1 SV=1</td></tr>
<tr><td>334</td><td>30.29</td><td>5</td><td>5</td><td>22539</td><td>423</td><td>>sp|O00231-2|PSD11_HUMAN Isoform 2 of 26S proteasome non-ATPase regulatory subunit 11 OS=Homo sapiens OX=9606 GN=PSMD11</td></tr>
<tr><td>335</td><td>30.21</td><td>5</td><td>4</td><td>42031</td><td>207</td><td>>sp|P51149|RAB7A_HUMAN Ras-related protein Rab-7a OS=Homo sapiens OX=9606 GN=RAB7A PE=1 SV=1</td></tr>
<tr><td>336</td><td>30.19</td><td>3</td><td>3</td><td>2165</td><td>352</td><td>>sp|Q9BT78-2|CSN4_HUMAN Isoform 2 of COP9 signalosome complex subunit 4 OS=Homo sapiens OX=9606 GN=COPS4</td></tr>
<tr><td>337</td><td>30.07</td><td>5</td><td>4</td><td>14431</td><td>1220</td><td>>sp|O60841|IF2P_HUMAN Eukaryotic translation initiation factor 5B OS=Homo sapiens OX=9606 GN=EIF5B PE=1 SV=4</td></tr>
<tr><td>338</td><td>30.07</td><td>3</td><td>3</td><td>7555</td><td>670</td><td>>sp|Q9NVP1|DDX18_HUMAN ATP-dependent RNA helicase DDX18 OS=Homo sapiens OX=9606 GN=DDX18 PE=1 SV=2</td></tr>
<tr><td>339</td><td>30.03</td><td>4</td><td>3</td><td>2626</td><td>430</td><td>>sp|Q9UJU6|DBNL_HUMAN Drebrin-like protein OS=Homo sapiens OX=9606 GN=DBNL PE=1 SV=1</td></tr>
<tr><td>340</td><td>29.93</td><td>4</td><td>4</td><td>35086</td><td>257</td><td>>sp|Q86V81|THOC4_HUMAN THO complex subunit 4 OS=Homo sapiens OX=9606 GN=ALYREF PE=1 SV=3</td></tr>
<tr><td>341</td><td>29.89</td><td>4</td><td>4</td><td>17314</td><td>637</td><td>>sp|P15170-3|ERF3A_HUMAN Isoform 3 of Eukaryotic peptide chain release factor GTP-binding subunit ERF3A OS=Homo sapiens OX=9606</td></tr>
<tr><td>342</td><td>29.85</td><td>5</td><td>5</td><td>19877</td><td>428</td><td>>sp|P08621-2|RU17_HUMAN Isoform 2 of U1 small nuclear ribonucleoprotein 70 kDa OS=Homo sapiens OX=9606 GN=SNRNP70</td></tr>
<tr><td>343</td><td>29.57</td><td>5</td><td>5</td><td>41727</td><td>283</td><td>>sp|P45880-2|VDAC2_HUMAN Isoform 2 of Voltage-dependent anion-selective channel protein 2 OS=Homo sapiens OX=9606 GN=VDAC2</td></tr>
<tr><td>344</td><td>29.46</td><td>6</td><td>5</td><td>12626</td><td>456</td><td>>sp|P14866-2|HNRPL_HUMAN Isoform 2 of Heterogeneous nuclear ribonucleoprotein L OS=Homo sapiens OX=9606 GN=HNRNPL</td></tr>
<tr><td>345</td><td>29.43</td><td>3</td><td>3</td><td>26577</td><td>307</td><td>>sp|Q96AG4|LRC59_HUMAN Leucine-rich repeat-containing protein 59 OS=Homo sapiens OX=9606 GN=LRRC59 PE=1 SV=1</td></tr>
<tr><td>346</td><td>29.41</td><td>4</td><td>4</td><td>41617</td><td>200</td><td>>sp|P61086|UBE2K_HUMAN Ubiquitin-conjugating enzyme E2 K OS=Homo sapiens OX=9606 GN=UBE2K PE=1 SV=3</td></tr>
<tr><td>347</td><td>29.31</td><td>2</td><td>2</td><td>21300</td><td>128</td><td>>sp|P35268|RL22_HUMAN 60S ribosomal protein L22 OS=Homo sapiens OX=9606 GN=RPL22 PE=1 SV=2</td></tr>
<tr><td>348</td><td>29.26</td><td>6</td><td>6</td><td>19839</td><td>501</td><td>>sp|Q12874|SF3A3_HUMAN Splicing factor 3A subunit 3 OS=Homo sapiens OX=9606 GN=SF3A3 PE=1 SV=1</td></tr>
<tr><td>349</td><td>29.16</td><td>3</td><td>3</td><td>7595</td><td>718</td><td>>sp|P51659-3|DHB4_HUMAN Isoform 3 of Peroxisomal multifunctional enzyme type 2 OS=Homo sapiens OX=9606 GN=HSD17B4</td></tr>
<tr><td>350</td><td>29.11</td><td>4</td><td>4</td><td>11545</td><td>237</td><td>>sp|P06730-3|IF4E_HUMAN Isoform 3 of Eukaryotic translation initiation factor 4E OS=Homo sapiens OX=9606 GN=EIF4E</td></tr>
<tr><td>351</td><td>28.83</td><td>8</td><td>7</td><td>25975</td><td>847</td><td>>sp|P43243|MATR3_HUMAN Matrin-3 OS=Homo sapiens OX=9606 GN=MATR3 PE=1 SV=2</td></tr>
<tr><td>352</td><td>28.78</td><td>4</td><td>4</td><td>22730</td><td>263</td><td>>sp|P28074|PSB5_HUMAN Proteasome subunit beta type-5 OS=Homo sapiens OX=9606 GN=PSMB5 PE=1 SV=3</td></tr>
<tr><td>353</td><td>28.69</td><td>5</td><td>5</td><td>30538</td><td>660</td><td>>sp|Q13310-3|PABP4_HUMAN Isoform 3 of Polyadenylate-binding protein 4 OS=Homo sapiens OX=9606 GN=PABPC4</td></tr>
<tr><td>354</td><td>28.68</td><td>4</td><td>3</td><td>21694</td><td>305</td><td>>sp|Q13151|ROA0_HUMAN Heterogeneous nuclear ribonucleoprotein A0 OS=Homo sapiens OX=9606 GN=HNRNPA0 PE=1 SV=1</td></tr>
<tr><td>355</td><td>28.63</td><td>6</td><td>4</td><td>22391</td><td>364</td><td>>sp|Q9BWF3|RBM4_HUMAN RNA-binding protein 4 OS=Homo sapiens OX=9606 GN=RBM4 PE=1 SV=1</td></tr>
<tr><td>356</td><td>28.58</td><td>7</td><td>6</td><td>17292</td><td>320</td><td>>sp|O75821|EIF3G_HUMAN Eukaryotic translation initiation factor 3 subunit G OS=Homo sapiens OX=9606 GN=EIF3G PE=1 SV=2</td></tr>
<tr><td>357</td><td>28.46</td><td>4</td><td>4</td><td>29958</td><td>306</td><td>>sp|Q9NZL9-4|MAT2B_HUMAN Isoform 4 of Methionine adenosyltransferase 2 subunit beta OS=Homo sapiens OX=9606 GN=MAT2B</td></tr>
<tr><td>358</td><td>28.45</td><td>5</td><td>5</td><td>38867</td><td>689</td><td>>sp|P52888|THOP1_HUMAN Thimet oligopeptidase OS=Homo sapiens OX=9606 GN=THOP1 PE=1 SV=2</td></tr>
<tr><td>359</td><td>28.28</td><td>9</td><td>8</td><td>40872</td><td>765</td><td>>sp|P11387|TOP1_HUMAN DNA topoisomerase 1 OS=Homo sapiens OX=9606 GN=TOP1 PE=1 SV=2</td></tr>
<tr><td>360</td><td>28.27</td><td>4</td><td>4</td><td>25053</td><td>1074</td><td>>sp|Q96KR1|ZFR_HUMAN Zinc finger RNA-binding protein OS=Homo sapiens OX=9606 GN=ZFR PE=1 SV=2</td></tr>
<tr><td>361</td><td>28.18</td><td>3</td><td>3</td><td>41223</td><td>1204</td><td>>sp|Q9HAV4|XPO5_HUMAN Exportin-5 OS=Homo sapiens OX=9606 GN=XPO5 PE=1 SV=1</td></tr>
<tr><td>362</td><td>28.16</td><td>3</td><td>3</td><td>41620</td><td>152</td><td>>sp|P61088|UBE2N_HUMAN Ubiquitin-conjugating enzyme E2 N OS=Homo sapiens OX=9606 GN=UBE2N PE=1 SV=1</td></tr>
<tr><td>363</td><td>28.12</td><td>2</td><td>2</td><td>19175</td><td>268</td><td>>sp|Q15287-3|RNPS1_HUMAN Isoform 3 of RNA-binding protein with serine-rich domain 1 OS=Homo sapiens OX=9606 GN=RNPS1</td></tr>
<tr><td>364</td><td>28.11</td><td>6</td><td>5</td><td>34685</td><td>226</td><td>>sp|Q16629-4|SRSF7_HUMAN Isoform 4 of Serine/arginine-rich splicing factor 7 OS=Homo sapiens OX=9606 GN=SRSF7</td></tr>
<tr><td>365</td><td>28.08</td><td>3</td><td>3</td><td>15590</td><td>400</td><td>>sp|P50395-2|GDIB_HUMAN Isoform 2 of Rab GDP dissociation inhibitor beta OS=Homo sapiens OX=9606 GN=GDI2</td></tr>
<tr><td>366</td><td>27.94</td><td>4</td><td>4</td><td>19433</td><td>215</td><td>>sp|O75396|SC22B_HUMAN Vesicle-trafficking protein SEC22b OS=Homo sapiens OX=9606 GN=SEC22B PE=1 SV=4</td></tr>
<tr><td>367</td><td>27.85</td><td>7</td><td>6</td><td>22743</td><td>523</td><td>>sp|Q8WXF1|PSPC1_HUMAN Paraspeckle component 1 OS=Homo sapiens OX=9606 GN=PSPC1 PE=1 SV=1</td></tr>
<tr><td>368</td><td>27.81</td><td>4</td><td>4</td><td>28433</td><td>366</td><td>>sp|Q15366-2|PCBP2_HUMAN Isoform 2 of Poly(rC)-binding protein 2 OS=Homo sapiens OX=9606 GN=PCBP2</td></tr>
<tr><td>369</td><td>27.79</td><td>3</td><td>2</td><td>7538</td><td>534</td><td>>sp|Q9H4M9|EHD1_HUMAN EH domain-containing protein 1 OS=Homo sapiens OX=9606 GN=EHD1 PE=1 SV=2</td></tr>
<tr><td>370</td><td>27.75</td><td>3</td><td>2</td><td>22729</td><td>241</td><td>>sp|P20618|PSB1_HUMAN Proteasome subunit beta type-1 OS=Homo sapiens OX=9606 GN=PSMB1 PE=1 SV=2</td></tr>
<tr><td>371</td><td>27.72</td><td>5</td><td>5</td><td>37719</td><td>183</td><td>>sp|P61081|UBC12_HUMAN NEDD8-conjugating enzyme Ubc12 OS=Homo sapiens OX=9606 GN=UBE2M PE=1 SV=1</td></tr>
<tr><td>372</td><td>27.69</td><td>7</td><td>4</td><td>19239</td><td>165</td><td>>sp|P46783|RS10_HUMAN 40S ribosomal protein S10 OS=Homo sapiens OX=9606 GN=RPS10 PE=1 SV=1</td></tr>
<tr><td>373</td><td>27.64</td><td>7</td><td>6</td><td>14643</td><td>1403</td><td>>sp|Q04637-6|IF4G1_HUMAN Isoform E of Eukaryotic translation initiation factor 4 gamma 1 OS=Homo sapiens OX=9606 GN=EIF4G1</td></tr>
<tr><td>374</td><td>27.55</td><td>3</td><td>3</td><td>10935</td><td>325</td><td>>sp|Q13347|EIF3I_HUMAN Eukaryotic translation initiation factor 3 subunit I OS=Homo sapiens OX=9606 GN=EIF3I PE=1 SV=1</td></tr>
<tr><td>375</td><td>27.53</td><td>3</td><td>3</td><td>19133</td><td>214</td><td>>sp|P27635|RL10_HUMAN 60S ribosomal protein L10 OS=Homo sapiens OX=9606 GN=RPL10 PE=1 SV=4</td></tr>
<tr><td>376</td><td>27.52</td><td>4</td><td>4</td><td>23654</td><td>154</td><td>>sp|P00441|SODC_HUMAN Superoxide dismutase [Cu-Zn] OS=Homo sapiens OX=9606 GN=SOD1 PE=1 SV=2</td></tr>
<tr><td>377</td><td>27.40</td><td>4</td><td>3</td><td>21316</td><td>148</td><td>>sp|P46776|RL27A_HUMAN 60S ribosomal protein L27a OS=Homo sapiens OX=9606 GN=RPL27A PE=1 SV=2</td></tr>
<tr><td>378</td><td>27.20</td><td>7</td><td>5</td><td>19134</td><td>211</td><td>>sp|P26373|RL13_HUMAN 60S ribosomal protein L13 OS=Homo sapiens OX=9606 GN=RPL13 PE=1 SV=4</td></tr>
</table>

| | | | | | | |
|---|---|---|---|---|---|---|
| 379 | 27.16 | 3 | 2 | 12196 | 181 | >sp\|Q13442\|HAP28_HUMAN 28 kDa heat- and acid-stable phosphoprotein OS=Homo sapiens OX=9606 GN=PDAP1 PE=1 SV=1 |
| 380 | 27.11 | 3 | 3 | 3560 | 661 | >sp\|P08195-4\|4F2_HUMAN Isoform 4 of 4F2 cell-surface antigen heavy chain OS=Homo sapiens OX=9606 GN=SLC3A2 |
| 381 | 27.07 | 4 | 4 | 4113 | 700 | >sp\|P17655\|CAN2_HUMAN Calpain-2 catalytic subunit OS=Homo sapiens OX=9606 GN=CAPN2 PE=1 SV=6 |
| 382 | 27.06 | 5 | 5 | 42301 | 1338 | >sp\|O15067\|PUR4_HUMAN Phosphoribosylformylglycinamidine synthase OS=Homo sapiens OX=9606 GN=PFAS PE=1 SV=4 |
| 383 | 27.00 | 7 | 6 | 36666 | 1071 | >sp\|O14980\|XPO1_HUMAN Exportin-1 OS=Homo sapiens OX=9606 GN=XPO1 PE=1 SV=1 |
| 384 | 26.90 | 3 | 3 | 21526 | 192 | >sp\|P32969\|RL9_HUMAN 60S ribosomal protein L9 OS=Homo sapiens OX=9606 GN=RPL9 PE=1 SV=1 |
| 385 | 26.88 | 6 | 6 | 7454 | 1280 | >sp\|P26358-3\|DNMT1_HUMAN Isoform 3 of DNA (cytosine-5)-methyltransferase 1 OS=Homo sapiens OX=9606 GN=DNMT1 |
| 386 | 26.82 | 3 | 3 | 22648 | 265 | >sp\|P61289-3\|PSME3_HUMAN Isoform 3 of Proteasome activator complex subunit 3 OS=Homo sapiens OX=9606 GN=PSME3 |
| 387 | 26.80 | 3 | 3 | 10338 | 380 | >sp\|P39748\|FEN1_HUMAN Flap endonuclease 1 OS=Homo sapiens OX=9606 GN=FEN1 PE=1 SV=1 |
| 388 | 26.65 | 2 | 2 | 20681 | 305 | >sp\|P40938-2\|RFC3_HUMAN Isoform 2 of Replication factor C subunit 3 OS=Homo sapiens OX=9606 GN=RFC3 |
| 389 | 26.64 | 4 | 4 | 19881 | 386 | >sp\|Q9Y265-2\|RUVB1_HUMAN Isoform 2 of RuvB-like 1 OS=Homo sapiens OX=9606 GN=RUVBL1 |
| 390 | 26.58 | 5 | 5 | 21287 | 156 | >sp\|P62750\|RL23A_HUMAN 60S ribosomal protein L23a OS=Homo sapiens OX=9606 GN=RPL23A PE=1 SV=1 |
| 391 | 26.45 | 5 | 4 | 4416 | 183 | >sp\|Q13185\|CBX3_HUMAN Chromobox protein homolog 3 OS=Homo sapiens OX=9606 GN=CBX3 PE=1 SV=4 |
| 392 | 26.41 | 6 | 4 | 8893 | 478 | >sp\|Q16630-3\|CPSF6_HUMAN Isoform 3 of Cleavage and polyadenylation specificity factor subunit 6 OS=Homo sapiens OX=9606 GN=CPSF |
| 393 | 26.40 | 4 | 3 | 38613 | 444 | >sp\|P07437\|TBB5_HUMAN Tubulin beta chain OS=Homo sapiens OX=9606 GN=TUBB PE=1 SV=2 |
| 394 | 26.28 | 3 | 3 | 24230 | 930 | >sp\|P12814-4\|ACTN1_HUMAN Isoform 4 of Alpha-actinin-1 OS=Homo sapiens OX=9606 GN=ACTN1 |
| 395 | 26.19 | 5 | 4 | 6904 | 331 | >sp\|P31689-2\|DNJA1_HUMAN Isoform 2 of DnaJ homolog subfamily A member 1 OS=Homo sapiens OX=9606 GN=DNAJA1 |
| 396 | 26.17 | 4 | 4 | 22453 | 439 | >sp\|P17980\|PRS6A_HUMAN 26S proteasome regulatory subunit 6A OS=Homo sapiens OX=9606 GN=PSMC3 PE=1 SV=3 |
| 397 | 26.16 | 4 | 3 | 32958 | 268 | >sp\|Q15691\|MARE1_HUMAN Microtubule-associated protein RP/EB family member 1 OS=Homo sapiens OX=9606 GN=MAPRE1 PE=1 SV=3 |
| 398 | 26.12 | 3 | 3 | 21044 | 1173 | >sp\|Q92878-3\|RAD50_HUMAN Isoform 3 of DNA repair protein RAD50 OS=Homo sapiens OX=9606 GN=RAD50 |
| 399 | 26.11 | 4 | 4 | 17189 | 285 | >sp\|P42126-2\|ECI1_HUMAN Isoform 2 of Enoyl-CoA delta isomerase 1; mitochondrial OS=Homo sapiens OX=9606 GN=ECI1 |
| 400 | 26.10 | 8 | 8 | 20623 | 669 | >sp\|Q96PK6\|RBM14_HUMAN RNA-binding protein 14 OS=Homo sapiens OX=9606 GN=RBM14 PE=1 SV=2 |
| 401 | 26.08 | 5 | 5 | 9133 | 185 | >sp\|Q9HB71-3\|CYBP_HUMAN Isoform 3 of Calcyclin-binding protein OS=Homo sapiens OX=9606 GN=CACYBP |
| 402 | 25.93 | 9 | 9 | 29263 | 2335 | >sp\|Q6P2Q9\|PRP8_HUMAN Pre-mRNA-processing-splicing factor 8 OS=Homo sapiens OX=9606 GN=PRPF8 PE=1 SV=2 |
| 403 | 25.93 | 5 | 5 | 10546 | 357 | >sp\|O00303\|EIF3F_HUMAN Eukaryotic translation initiation factor 3 subunit F OS=Homo sapiens OX=9606 GN=EIF3F PE=1 SV=1 |
| 404 | 25.86 | 5 | 5 | 30638 | 271 | >sp\|Q96FW1\|OTUB1_HUMAN Ubiquitin thioesterase OTUB1 OS=Homo sapiens OX=9606 GN=OTUB1 PE=1 SV=2 |
| 405 | 25.74 | 2 | 2 | 28827 | 389 | >sp\|P62333\|PRS10_HUMAN 26S proteasome regulatory subunit 10B OS=Homo sapiens OX=9606 GN=PSMC6 PE=1 SV=1 |
| 406 | 25.68 | 5 | 3 | 19309 | 142 | >sp\|P60866-2\|RS20_HUMAN Isoform 2 of 40S ribosomal protein S20 OS=Homo sapiens OX=9606 GN=RPS20 |
| 407 | 25.65 | 4 | 3 | 204 | 353 | >sp\|Q99873-3\|ANM1_HUMAN Isoform 3 of Protein arginine N-methyltransferase 1 OS=Homo sapiens OX=9606 GN=PRMT1 |
| 408 | 25.55 | 5 | 4 | 7792 | 165 | >sp\|P60981\|DEST_HUMAN Destrin OS=Homo sapiens OX=9606 GN=DSTN PE=1 SV=3 |
| 409 | 25.53 | 3 | 3 | 41385 | 973 | >sp\|O60763-2\|USO1_HUMAN Isoform 2 of General vesicular transport factor p115 OS=Homo sapiens OX=9606 GN=USO1 |
| 410 | 25.48 | 3 | 3 | 33834 | 587 | >sp\|P09960-4\|LKHA4_HUMAN Isoform 4 of Leukotriene A-4 hydrolase OS=Homo sapiens OX=9606 GN=LTA4H |
| 411 | 25.42 | 3 | 3 | 32243 | 334 | >sp\|P00387-3\|NB5R3_HUMAN Isoform 3 of NADH-cytochrome b5 reductase 3 OS=Homo sapiens OX=9606 GN=CYB5R3 |
| 412 | 25.39 | 3 | 3 | 34830 | 445 | >sp\|Q9BVA1\|TBB2B_HUMAN Tubulin beta-2B chain OS=Homo sapiens OX=9606 GN=TUBB2B PE=1 SV=1 |
| 413 | 25.39 | 6 | 5 | 28812 | 224 | >sp\|P30041\|PRDX6_HUMAN Peroxiredoxin-6 OS=Homo sapiens OX=9606 GN=PRDX6 PE=1 SV=3 |
| 414 | 25.36 | 2 | 2 | 16997 | 284 | >sp\|P13804-2\|ETFA_HUMAN Isoform 2 of Electron transfer flavoprotein subunit alpha; mitochondrial OS=Homo sapiens OX=9606 GN=ETF |
| 415 | 25.10 | 7 | 6 | 24713 | 511 | >sp\|P49419-2\|AL7A1_HUMAN Isoform 2 of Alpha-aminoadipic semialdehyde dehydrogenase OS=Homo sapiens OX=9606 GN=ALDH7A1 |
| 416 | 25.08 | 5 | 5 | 14492 | 333 | >sp\|P20042\|IF2B_HUMAN Eukaryotic translation initiation factor 2 subunit 2 OS=Homo sapiens OX=9606 GN=EIF2S2 PE=1 SV=2 |
| 417 | 25.07 | 7 | 7 | 8102 | 558 | >sp\|P00367\|DHE3_HUMAN Glutamate dehydrogenase 1; mitochondrial OS=Homo sapiens OX=9606 GN=GLUD1 PE=1 SV=2 |
| 418 | 24.94 | 2 | 2 | 28590 | 329 | >sp\|O00151\|PDLI1_HUMAN PDZ and LIM domain protein 1 OS=Homo sapiens OX=9606 GN=PDLIM1 PE=1 SV=4 |
| 419 | 24.91 | 6 | 6 | 236 | 320 | >sp\|P08758\|ANXA5_HUMAN Annexin A5 OS=Homo sapiens OX=9606 GN=ANXA5 PE=1 SV=2 |
| 420 | 24.76 | 5 | 4 | 437 | 1042 | >sp\|P16615\|AT2A2_HUMAN Sarcoplasmic/endoplasmic reticulum calcium ATPase 2 OS=Homo sapiens OX=9606 GN=ATP2A2 PE=1 SV=1 |
| 421 | 24.75 | 4 | 4 | 10545 | 533 | >sp\|O15371-3\|EIF3D_HUMAN Isoform 3 of Eukaryotic translation initiation factor 3 subunit D OS=Homo sapiens OX=9606 GN=EIF3D |
| 422 | 24.66 | 5 | 5 | 35685 | 548 | >sp\|O43776\|SYNC_HUMAN Asparagine--tRNA ligase; cytoplasmic OS=Homo sapiens OX=9606 GN=NARS PE=1 SV=1 |
| 423 | 24.65 | 5 | 4 | 20020 | 563 | >sp\|Q15436-2\|SC23A_HUMAN Isoform 2 of Protein transport protein Sec23A OS=Homo sapiens OX=9606 GN=SEC23A |
| 424 | 24.57 | 4 | 4 | 26028 | 757 | >sp\|Q16891-4\|MIC60_HUMAN Isoform 4 of MICOS complex subunit MIC60 OS=Homo sapiens OX=9606 GN=IMMT |
| 425 | 24.56 | 6 | 6 | 35680 | 625 | >sp\|Q15046-2\|SYK_HUMAN Isoform Mitochondrial of Lysine--tRNA ligase OS=Homo sapiens OX=9606 GN=KARS |
| 426 | 24.50 | 4 | 4 | 39939 | 529 | >sp\|Q9Y5A9-2\|YTHD2_HUMAN Isoform 2 of YTH domain-containing family protein 2 OS=Homo sapiens OX=9606 GN=YTHDF2 |
| 427 | 24.50 | 3 | 3 | 27996 | 873 | >sp\|Q8WUM4-2\|PDC6I_HUMAN Isoform 2 of Programmed cell death 6-interacting protein OS=Homo sapiens OX=9606 GN=PDCD6IP |
| 428 | 24.47 | 4 | 4 | 19361 | 255 | >sp\|P09661\|RU2A_HUMAN U2 small nuclear ribonucleoprotein A' OS=Homo sapiens OX=9606 GN=SNRPA1 PE=1 SV=2 |
| 429 | 24.13 | 2 | 2 | 26361 | 739 | >sp\|O95202\|LETM1_HUMAN Mitochondrial proton/calcium exchanger protein OS=Homo sapiens OX=9606 GN=LETM1 PE=1 SV=1 |
| 430 | 23.83 | 2 | 2 | 7514 | 148 | >sp\|O60869\|EDF1_HUMAN Endothelial differentiation-related factor 1 OS=Homo sapiens OX=9606 GN=EDF1 PE=1 SV=1 |
| 431 | 23.81 | 3 | 3 | 35797 | 414 | >sp\|Q13148\|TADBP_HUMAN TAR DNA-binding protein 43 OS=Homo sapiens OX=9606 GN=TARDBP PE=1 SV=1 |

| | | | | | | |
|---|---|---|---|---|---|---|
| 432 | 23.72 | 3 | 3 | 1424 | 313 | >sp\|Q9UHD1-2\|CHRD1_HUMAN Isoform 2 of Cysteine and histidine-rich domain-containing protein 1 OS=Homo sapiens OX=9606 GN=CHORDC |
| 433 | 23.67 | 2 | 2 | 30760 | 634 | >sp\|Q86YP4-3\|P66A_HUMAN Isoform 3 of Transcriptional repressor p66-alpha OS=Homo sapiens OX=9606 GN=GATAD2A |
| 434 | 23.64 | 2 | 2 | 27872 | 534 | >sp\|Q8NF37\|PCAT1_HUMAN Lysophosphatidylcholine acyltransferase 1 OS=Homo sapiens OX=9606 GN=LPCAT1 PE=1 SV=2 |
| 435 | 23.53 | 4 | 4 | 34519 | 470 | >sp\|P52209-2\|6PGD_HUMAN Isoform 2 of 6-phosphogluconate dehydrogenase; decarboxylating OS=Homo sapiens OX=9606 GN=PGD |
| 436 | 23.53 | 3 | 3 | 2841 | 269 | >sp\|P35613-2\|BASI_HUMAN Isoform 2 of Basigin OS=Homo sapiens OX=9606 GN=BSG |
| 437 | 23.49 | 6 | 5 | 31475 | 943 | >sp\|O14974-5\|MYPT1_HUMAN Isoform 5 of Protein phosphatase 1 regulatory subunit 12A OS=Homo sapiens OX=9606 GN=PPP1R12A |
| 438 | 23.39 | 3 | 3 | 34145 | 377 | >sp\|P68032\|ACTC_HUMAN Actin; alpha cardiac muscle 1 OS=Homo sapiens OX=9606 GN=ACTC1 PE=1 SV=1 |
| 439 | 23.36 | 3 | 2 | 32763 | 440 | >sp\|Q9UKX7-2\|NUP50_HUMAN Isoform 2 of Nuclear pore complex protein Nup50 OS=Homo sapiens OX=9606 GN=NUP50 |
| 440 | 23.35 | 2 | 2 | 22483 | 649 | >sp\|P46063\|RECQ1_HUMAN ATP-dependent DNA helicase Q1 OS=Homo sapiens OX=9606 GN=RECQL PE=1 SV=3 |
| 441 | 23.34 | 5 | 4 | 22227 | 151 | >sp\|P62263\|RS14_HUMAN 40S ribosomal protein S14 OS=Homo sapiens OX=9606 GN=RPS14 PE=1 SV=3 |
| 442 | 23.30 | 5 | 5 | 32152 | 845 | >sp\|P46087-4\|NOP2_HUMAN Isoform 4 of Probable 28S rRNA (cytosine(4447)-C(5))-methyltransferase OS=Homo sapiens OX=9606 GN=NOP2 |
| 443 | 23.29 | 3 | 2 | 24477 | 609 | >sp\|O95831-3\|AIFM1_HUMAN Isoform 3 of Apoptosis-inducing factor 1; mitochondrial OS=Homo sapiens OX=9606 GN=AIFM1 |
| 444 | 23.13 | 5 | 5 | 23403 | 1288 | >sp\|Q9NTJ3\|SMC4_HUMAN Structural maintenance of chromosomes protein 4 OS=Homo sapiens OX=9606 GN=SMC4 PE=1 SV=2 |
| 445 | 23.11 | 3 | 3 | 20969 | 193 | >sp\|P61586\|RHOA_HUMAN Transforming protein RhoA OS=Homo sapiens OX=9606 GN=RHOA PE=1 SV=1 |
| 446 | 22.89 | 3 | 3 | 2781 | 213 | >sp\|P48047\|ATPO_HUMAN ATP synthase subunit O; mitochondrial OS=Homo sapiens OX=9606 GN=ATP5O PE=1 SV=1 |
| 447 | 22.82 | 3 | 3 | 19250 | 158 | >sp\|P62280\|RS11_HUMAN 40S ribosomal protein S11 OS=Homo sapiens OX=9606 GN=RPS11 PE=1 SV=3 |
| 448 | 22.81 | 3 | 2 | 17941 | 172 | >sp\|O14950\|ML12B_HUMAN Myosin regulatory light chain 12B OS=Homo sapiens OX=9606 GN=MYL12B PE=1 SV=2 |
| 449 | 22.78 | 3 | 3 | 21377 | 177 | >sp\|P62913-2\|RL11_HUMAN Isoform 2 of 60S ribosomal protein L11 OS=Homo sapiens OX=9606 GN=RPL11 |
| 450 | 22.78 | 3 | 2 | 22643 | 377 | >sp\|P55036\|PSMD4_HUMAN 26S proteasome non-ATPase regulatory subunit 4 OS=Homo sapiens OX=9606 GN=PSMD4 PE=1 SV=1 |
| 451 | 22.70 | 4 | 4 | 37907 | 1104 | >sp\|Q14157-5\|UBP2L_HUMAN Isoform 5 of Ubiquitin-associated protein 2-like OS=Homo sapiens OX=9606 GN=UBAP2L |
| 452 | 22.63 | 5 | 5 | 6195 | 802 | >sp\|Q99459\|CDC5L_HUMAN Cell division cycle 5-like protein OS=Homo sapiens OX=9606 GN=CDC5L PE=1 SV=2 |
| 453 | 22.53 | 4 | 4 | 34008 | 807 | >sp\|Q8NE71-2\|ABCF1_HUMAN Isoform 2 of ATP-binding cassette sub-family F member 1 OS=Homo sapiens OX=9606 GN=ABCF1 |
| 454 | 22.51 | 3 | 3 | 39068 | 172 | >sp\|P13693\|TCTP_HUMAN Translationally-controlled tumor protein OS=Homo sapiens OX=9606 GN=TPT1 PE=1 SV=1 |
| 455 | 22.48 | 5 | 4 | 11406 | 2079 | >sp\|P51610-4\|HCFC1_HUMAN Isoform 4 of Host cell factor 1 OS=Homo sapiens OX=9606 GN=HCFC1 |
| 456 | 22.39 | 3 | 3 | 41927 | 480 | >sp\|P31930\|QCR1_HUMAN Cytochrome b-c1 complex subunit 1; mitochondrial OS=Homo sapiens OX=9606 GN=UQCRC1 PE=1 SV=3 |
| 457 | 22.34 | 3 | 3 | 26503 | 489 | >sp\|O75439\|MPPB_HUMAN Mitochondrial-processing peptidase subunit beta OS=Homo sapiens OX=9606 GN=PMPCB PE=1 SV=2 |
| 458 | 22.34 | 2 | 2 | 18991 | 173 | >sp\|Q9Y5S9-2\|RBM8A_HUMAN Isoform 2 of RNA-binding protein 8A OS=Homo sapiens OX=9606 GN=RBM8A |
| 459 | 22.33 | 6 | 5 | 20644 | 280 | >sp\|Q15293-2\|RCN1_HUMAN Isoform 2 of Reticulocalbin-1 OS=Homo sapiens OX=9606 GN=RCN1 |
| 460 | 22.33 | 4 | 3 | 12503 | 209 | >sp\|P26583\|HMGB2_HUMAN High mobility group protein B2 OS=Homo sapiens OX=9606 GN=HMGB2 PE=1 SV=2 |
| 461 | 22.25 | 2 | 2 | 16994 | 1114 | >sp\|Q9BSJ8-2\|ESYT1_HUMAN Isoform 2 of Extended synaptotagmin-1 OS=Homo sapiens OX=9606 GN=ESYT1 |
| 462 | 22.18 | 4 | 3 | 26244 | 711 | >sp\|P49959-3\|MRE11_HUMAN Isoform 3 of Double-strand break repair protein MRE11 OS=Homo sapiens OX=9606 GN=MRE11 |
| 463 | 22.06 | 3 | 3 | 11533 | 144 | >sp\|P47813\|IF1AX_HUMAN Eukaryotic translation initiation factor 1A; X-chromosomal OS=Homo sapiens OX=9606 GN=EIF1AX PE=1 SV=2 |
| 464 | 22.04 | 3 | 3 | 16679 | 798 | >sp\|P05556\|ITB1_HUMAN Integrin beta-1 OS=Homo sapiens OX=9606 GN=ITGB1 PE=1 SV=2 |
| 465 | 22.01 | 5 | 5 | 21804 | 490 | >sp\|O76021\|RL1D1_HUMAN Ribosomal L1 domain-containing protein 1 OS=Homo sapiens OX=9606 GN=RSL1D1 PE=1 SV=3 |
| 466 | 21.96 | 2 | 2 | 27676 | 254 | >sp\|Q53H96-2\|P5CR3_HUMAN Isoform 2 of Pyrroline-5-carboxylate reductase 3 OS=Homo sapiens OX=9606 GN=PYCR3 |
| 467 | 21.95 | 5 | 5 | 37170 | 764 | >sp\|P47897-2\|SYQ_HUMAN Isoform 2 of Glutamine--tRNA ligase OS=Homo sapiens OX=9606 GN=QARS |
| 468 | 21.91 | 3 | 3 | 21880 | 83 | >sp\|P63220\|RS21_HUMAN 40S ribosomal protein S21 OS=Homo sapiens OX=9606 GN=RPS21 PE=1 SV=1 |
| 469 | 21.91 | 2 | 2 | 5553 | 202 | >sp\|Q9Y3Y2-4\|CHTOP_HUMAN Isoform 3 of Chromatin target of PRMT1 protein OS=Homo sapiens OX=9606 GN=CHTOP |
| 470 | 21.89 | 2 | 2 | 21948 | 244 | >sp\|Q9Y224\|RTRAF_HUMAN RNA transcription; translation and transport factor protein OS=Homo sapiens OX=9606 GN=RTRAF PE=1 SV=1 |
| 471 | 21.81 | 2 | 2 | 31046 | 516 | >sp\|P13674-3\|P4HA1_HUMAN Isoform 3 of Prolyl 4-hydroxylase subunit alpha-1 OS=Homo sapiens OX=9606 GN=P4HA1 |
| 472 | 21.78 | 2 | 2 | 30919 | 439 | >sp\|P04181\|OAT_HUMAN Ornithine aminotransferase; mitochondrial OS=Homo sapiens OX=9606 GN=OAT PE=1 SV=1 |
| 473 | 21.74 | 2 | 2 | 31374 | 341 | >sp\|P11177-3\|ODPB_HUMAN Isoform 3 of Pyruvate dehydrogenase E1 component subunit beta; mitochondrial OS=Homo sapiens OX=9606 GN |
| 474 | 21.56 | 2 | 2 | 21699 | 285 | >sp\|Q99729-3\|ROAA_HUMAN Isoform 3 of Heterogeneous nuclear ribonucleoprotein A/B OS=Homo sapiens OX=9606 GN=HNRNPAB |
| 475 | 21.52 | 2 | 2 | 36596 | 641 | >sp\|Q9Y2W2\|WBP11_HUMAN WW domain-binding protein 11 OS=Homo sapiens OX=9606 GN=WBP11 PE=1 SV=1 |
| 476 | 21.50 | 4 | 3 | 21918 | 130 | >sp\|P62244\|RS15A_HUMAN 40S ribosomal protein S15a OS=Homo sapiens OX=9606 GN=RPS15A PE=1 SV=2 |
| 477 | 21.49 | 2 | 2 | 24352 | 298 | >sp\|P12235\|ADT1_HUMAN ADP/ATP translocase 1 OS=Homo sapiens OX=9606 GN=SLC25A4 PE=1 SV=4 |
| 478 | 21.48 | 3 | 3 | 31683 | 187 | >sp\|O00746\|NDKM_HUMAN Nucleoside diphosphate kinase; mitochondrial OS=Homo sapiens OX=9606 GN=NME4 PE=1 SV=1 |
| 479 | 21.36 | 3 | 3 | 19293 | 326 | >sp\|Q9NQG5\|RPR1B_HUMAN Regulation of nuclear pre-mRNA domain-containing protein 1B OS=Homo sapiens OX=9606 GN=RPRD1B PE=1 SV=1 |
| 480 | 21.31 | 3 | 3 | 9255 | 321 | >sp\|P22087\|FBRL_HUMAN rRNA 2'-O-methyltransferase fibrillarin OS=Homo sapiens OX=9606 GN=FBL PE=1 SV=2 |
| 481 | 21.27 | 3 | 3 | 42194 | 534 | >sp\|O43242\|PSMD3_HUMAN 26S proteasome non-ATPase regulatory subunit 3 OS=Homo sapiens OX=9606 GN=PSMD3 PE=1 SV=2 |
| 482 | 21.19 | 2 | 2 | 36776 | 58 | >sp\|Q96IX5\|USMG5_HUMAN Up-regulated during skeletal muscle growth protein 5 OS=Homo sapiens OX=9606 GN=USMG5 PE=1 SV=1 |
| 483 | 21.16 | 4 | 4 | 22414 | 317 | >sp\|Q14257\|RCN2_HUMAN Reticulocalbin-2 OS=Homo sapiens OX=9606 GN=RCN2 PE=1 SV=1 |

| | | | | | | |
|---|---|---|---|---|---|---|
| 484 | 21.13 | 4 | 3 | 16422 | 218 | >sp\|Q9H910-3\|JUPI2_HUMAN Isoform 3 of Jupiter microtubule associated homolog 2 OS=Homo sapiens OX=9606 GN=JPT2 |
| 485 | 21.06 | 2 | 2 | 19143 | 145 | >sp\|Q6P1L8\|RM14_HUMAN 39S ribosomal protein L14; mitochondrial OS=Homo sapiens OX=9606 GN=MRPL14 PE=1 SV=1 |
| 486 | 21.05 | 3 | 2 | 24762 | 220 | >sp\|Q9BTT0-3\|AN32E_HUMAN Isoform 3 of Acidic leucine-rich nuclear phosphoprotein 32 family member E OS=Homo sapiens OX=9606 GN= |
| 487 | 21.01 | 5 | 5 | 7151 | 452 | >sp\|P55084-2\|ECHB_HUMAN Isoform 2 of Trifunctional enzyme subunit beta; mitochondrial OS=Homo sapiens OX=9606 GN=HADHB |
| 488 | 20.97 | 3 | 3 | 12737 | 224 | >sp\|P54819-5\|KAD2_HUMAN Isoform 5 of Adenylate kinase 2; mitochondrial OS=Homo sapiens OX=9606 GN=AK2 |
| 489 | 20.95 | 2 | 2 | 21801 | 215 | >sp\|P50914\|RL14_HUMAN 60S ribosomal protein L14 OS=Homo sapiens OX=9606 GN=RPL14 PE=1 SV=4 |
| 490 | 20.91 | 3 | 3 | 35745 | 992 | >sp\|P49588-2\|SYAC_HUMAN Isoform 2 of Alanine--tRNA ligase; cytoplasmic OS=Homo sapiens OX=9606 GN=AARS |
| 491 | 20.85 | 4 | 4 | 7215 | 225 | >sp\|P24534\|EF1B_HUMAN Elongation factor 1-beta OS=Homo sapiens OX=9606 GN=EEF1B2 PE=1 SV=3 |
| 492 | 20.84 | 4 | 4 | 12592 | 205 | >sp\|P04792\|HSPB1_HUMAN Heat shock protein beta-1 OS=Homo sapiens OX=9606 GN=HSPB1 PE=1 SV=2 |
| 493 | 20.78 | 2 | 2 | 28274 | 130 | >sp\|Q8WW12-2\|PCNP_HUMAN Isoform 2 of PEST proteolytic signal-containing nuclear protein OS=Homo sapiens OX=9606 GN=PCNP |
| 494 | 20.60 | 3 | 2 | 8021 | 1271 | >sp\|Q14203-6\|DCTN1_HUMAN Isoform 6 of Dynactin subunit 1 OS=Homo sapiens OX=9606 GN=DCTN1 |
| 495 | 20.58 | 2 | 2 | 23973 | 402 | >sp\|O60749-2\|SNX2_HUMAN Isoform 2 of Sorting nexin-2 OS=Homo sapiens OX=9606 GN=SNX2 |
| 496 | 20.52 | 2 | 2 | 9041 | 193 | >sp\|P21291\|CSRP1_HUMAN Cysteine and glycine-rich protein 1 OS=Homo sapiens OX=9606 GN=CSRP1 PE=1 SV=3 |
| 497 | 20.51 | 2 | 2 | 30793 | 524 | >sp\|Q13177\|PAK2_HUMAN Serine/threonine-protein kinase PAK 2 OS=Homo sapiens OX=9606 GN=PAK2 PE=1 SV=3 |
| 498 | 20.49 | 4 | 4 | 42160 | 504 | >sp\|Q9UMS4\|PRP19_HUMAN Pre-mRNA-processing factor 19 OS=Homo sapiens OX=9606 GN=PRPF19 PE=1 SV=1 |
| 499 | 20.47 | 4 | 3 | 19870 | 392 | >sp\|Q9NQC3-5\|RTN4_HUMAN Isoform 5 of Reticulon-4 OS=Homo sapiens OX=9606 GN=RTN4 |
| 500 | 20.47 | 2 | 2 | 12416 | 100 | >sp\|P05114\|HMGN1_HUMAN Non-histone chromosomal protein HMG-14 OS=Homo sapiens OX=9606 GN=HMGN1 PE=1 SV=3 |
| 501 | 20.45 | 3 | 3 | 32423 | 594 | >sp\|O00567\|NOP56_HUMAN Nucleolar protein 56 OS=Homo sapiens OX=9606 GN=NOP56 PE=1 SV=4 |
| 502 | 20.41 | 3 | 3 | 14142 | 537 | >sp\|Q9BVP2-2\|GNL3_HUMAN Isoform 2 of Guanine nucleotide-binding protein-like 3 OS=Homo sapiens OX=9606 GN=GNL3 |
| 503 | 20.36 | 4 | 3 | 37977 | 158 | >sp\|P63279\|UBC9_HUMAN SUMO-conjugating enzyme UBC9 OS=Homo sapiens OX=9606 GN=UBE2I PE=1 SV=1 |
| 504 | 20.34 | 5 | 3 | 11218 | 164 | >sp\|P33316-2\|DUT_HUMAN Isoform 2 of Deoxyuridine 5'-triphosphate nucleotidohydrolase; mitochondrial OS=Homo sapiens OX=9606 GN= |
| 505 | 20.33 | 4 | 4 | 4501 | 286 | >sp\|P52907\|CAZA1_HUMAN F-actin-capping protein subunit alpha-1 OS=Homo sapiens OX=9606 GN=CAPZA1 PE=1 SV=3 |
| 506 | 20.31 | 2 | 2 | 12637 | 218 | >sp\|P00492\|HPRT_HUMAN Hypoxanthine-guanine phosphoribosyltransferase OS=Homo sapiens OX=9606 GN=HPRT1 PE=1 SV=2 |
| 507 | 20.16 | 2 | 2 | 40291 | 243 | >sp\|Q9BVC6\|TM109_HUMAN Transmembrane protein 109 OS=Homo sapiens OX=9606 GN=TMEM109 PE=1 SV=1 |
| 508 | 20.14 | 5 | 5 | 29244 | 198 | >sp\|P32119\|PRDX2_HUMAN Peroxiredoxin-2 OS=Homo sapiens OX=9606 GN=PRDX2 PE=1 SV=5 |
| 509 | 20.06 | 3 | 3 | 31506 | 151 | >sp\|P60660-2\|MYL6_HUMAN Isoform Smooth muscle of Myosin light polypeptide 6 OS=Homo sapiens OX=9606 GN=MYL6 |
| 510 | 20.03 | 2 | 2 | 22245 | 190 | >sp\|Q9Y3D9\|RT23_HUMAN 28S ribosomal protein S23; mitochondrial OS=Homo sapiens OX=9606 GN=MRPS23 PE=1 SV=2 |
| 511 | 19.90 | 2 | 2 | 11978 | 205 | >sp\|P41236\|IPP2_HUMAN Protein phosphatase inhibitor 2 OS=Homo sapiens OX=9606 GN=PPP1R2 PE=1 SV=2 |
| 512 | 19.80 | 3 | 3 | 42013 | 205 | >sp\|P62820\|RAB1A_HUMAN Ras-related protein Rab-1A OS=Homo sapiens OX=9606 GN=RAB1A PE=1 SV=3 |
| 513 | 19.77 | 5 | 4 | 29256 | 140 | >sp\|P07737\|PROF1_HUMAN Profilin-1 OS=Homo sapiens OX=9606 GN=PFN1 PE=1 SV=2 |
| 514 | 19.66 | 2 | 2 | 16466 | 325 | >sp\|Q96CX2\|KCD12_HUMAN BTB/POZ domain-containing protein KCTD12 OS=Homo sapiens OX=9606 GN=KCTD12 PE=1 SV=1 |
| 515 | 19.60 | 4 | 3 | 20444 | 313 | >sp\|O43765\|SGTA_HUMAN Small glutamine-rich tetratricopeptide repeat-containing protein alpha OS=Homo sapiens OX=9606 GN=SGTA PE |
| 516 | 19.59 | 2 | 2 | 37006 | 327 | >sp\|O75436\|VP26A_HUMAN Vacuolar protein sorting-associated protein 26A OS=Homo sapiens OX=9606 GN=VPS26A PE=1 SV=2 |
| 517 | 19.59 | 3 | 3 | 40862 | 887 | >sp\|O14787-2\|TNPO2_HUMAN Isoform 2 of Transportin-2 OS=Homo sapiens OX=9606 GN=TNPO2 |
| 518 | 19.54 | 3 | 2 | 23241 | 114 | >sp\|O75368\|SH3L1_HUMAN SH3 domain-binding glutamic acid-rich-like protein OS=Homo sapiens OX=9606 GN=SH3BGRL PE=1 SV=1 |
| 519 | 19.50 | 3 | 2 | 7107 | 695 | >sp\|Q16643-3\|DREB_HUMAN Isoform 3 of Drebrin OS=Homo sapiens OX=9606 GN=DBN1 |
| 520 | 19.47 | 4 | 4 | 19553 | 520 | >sp\|P55809\|SCOT1_HUMAN Succinyl-CoA:3-ketoacid coenzyme A transferase 1; mitochondrial OS=Homo sapiens OX=9606 GN=OXCT1 PE=1 SV |
| 521 | 19.46 | 2 | 2 | 39058 | 280 | >sp\|P23193-2\|TCEA1_HUMAN Isoform 2 of Transcription elongation factor A protein 1 OS=Homo sapiens OX=9606 GN=TCEA1 |
| 522 | 19.46 | 5 | 3 | 21288 | 140 | >sp\|P62829\|RL23_HUMAN 60S ribosomal protein L23 OS=Homo sapiens OX=9606 GN=RPL23 PE=1 SV=1 |
| 523 | 19.44 | 3 | 3 | 22075 | 916 | >sp\|Q15424-4\|SAFB1_HUMAN Isoform 4 of Scaffold attachment factor B1 OS=Homo sapiens OX=9606 GN=SAFB |
| 524 | 19.40 | 3 | 3 | 36158 | 1083 | >sp\|O00267-2\|SPT5H_HUMAN Isoform 2 of Transcription elongation factor SPT5 OS=Homo sapiens OX=9606 GN=SUPT5H |
| 525 | 19.38 | 2 | 2 | 28430 | 233 | >sp\|Q9BVG4\|PBDC1_HUMAN Protein PBDC1 OS=Homo sapiens OX=9606 GN=PBDC1 PE=1 SV=1 |
| 526 | 19.32 | 2 | 2 | 36060 | 610 | >sp\|O76094-2\|SRP72_HUMAN Isoform 2 of Signal recognition particle subunit SRP72 OS=Homo sapiens OX=9606 GN=SRP72 |
| 527 | 19.08 | 6 | 4 | 21921 | 125 | >sp\|P62851\|RS25_HUMAN 40S ribosomal protein S25 OS=Homo sapiens OX=9606 GN=RPS25 PE=1 SV=1 |
| 528 | 19.07 | 2 | 2 | 42257 | 246 | >sp\|P60900\|PSA6_HUMAN Proteasome subunit alpha type-6 OS=Homo sapiens OX=9606 GN=PSMA6 PE=1 SV=1 |
| 529 | 19.00 | 5 | 3 | 12623 | 747 | >sp\|Q1KMD3\|HNRL2_HUMAN Heterogeneous nuclear ribonucleoprotein U-like protein 2 OS=Homo sapiens OX=9606 GN=HNRNPUL2 PE=1 SV=1 |
| 530 | 18.98 | 3 | 3 | 31758 | 491 | >sp\|P43490\|NAMPT_HUMAN Nicotinamide phosphoribosyltransferase OS=Homo sapiens OX=9606 GN=NAMPT PE=1 SV=1 |
| 531 | 18.96 | 3 | 3 | 10744 | 104 | >sp\|P84090\|ERH_HUMAN Enhancer of rudimentary homolog OS=Homo sapiens OX=9606 GN=ERH PE=1 SV=1 |
| 532 | 18.96 | 2 | 2 | 17243 | 209 | >sp\|O75822-3\|EIF3J_HUMAN Isoform 3 of Eukaryotic translation initiation factor 3 subunit J OS=Homo sapiens OX=9606 GN=EIF3J |
| 533 | 18.83 | 2 | 1 | 16609 | 1042 | >sp\|Q68E01-2\|INT3_HUMAN Isoform 2 of Integrator complex subunit 3 OS=Homo sapiens OX=9606 GN=INTS3 |
| 534 | 18.77 | 2 | 2 | 26828 | 774 | >sp\|Q02809-2\|PLOD1_HUMAN Isoform 2 of Procollagen-lysine;2-oxoglutarate 5-dioxygenase 1 OS=Homo sapiens OX=9606 GN=PLOD1 |

#### Score distribution

808 peptide matches above score 32, 1064 (15 reverse) matches above 20
Top score for reverse: 45, top score for reverse with > 9 aa's: 39
Charge distribution: z=+1: 0, z=+2: 776, z=+3: 32, z=+4: 0

#### Mass measurement errors

**Precursors**

*Before recalibration:*
Median precursor m/z error (observed minus true): 0.0001 Da, 0.2 ppm
Median precursor accuracy (absolute value of error): 0.0002 Da, 0.4 ppm
Precursors measured too high: 436 Too low: 239

*After recalibration:*
Median precursor m/z error (observed minus true): 0.0000 Da, 0.0 ppm
Median precursor accuracy (absolute value of error): 0.0002 Da, 0.4 ppm
Precursors measured too high: 364 Too low: 342

Off-by-one errors (nominal mass is +1 isotope): 9.6% (75/781)

**Fragments**

*Before recalibration:*
Median fragment m/z error (observed minus true): 0.0792 Da, 125.3 ppm
Median fragment accuracy (absolute value of error): 0.0797 Da, 127.8 ppm
Fragments (within ±0.5 Da) measured too high: 3997 Too low: 197

*After recalibration:*
Median fragment m/z error (observed minus true): 0.0159 Da, 23.9 ppm
Median fragment accuracy (absolute value of error): 0.0313 Da, 48.8 ppm
Fragments (within ±0.5 Da) measured too high: 2416 Too low: 1781

#### Cysteine

Cysteine set to a fixed modification of +57
Completeness: 52.8% (38/72) peptides C[+57], 45.8% (33/72) C[+0], 1.4% (1/72) C[+71] (from gel)
Carbamidomethylation artifacts (H,K,N-terminus[+57/+114]): 0.2% (2/888) of peptides
DTT artifacts (C[+209]): 0.7% (6/892) of peptides
Disulfide bridge (unmodified_C[-2]): 0.0% (0/5) of peptides containing at least two C's

#### Nonspecific cleavage

Cleavage sites (C-side): RK
Missed cleavage: 19.7% (143/726) of semitryptic peptides contain an internal K or R not followed by P
Semitryptic peptides (% of tryptic and semitryptic): 5.1% (37/726) ragged-N, 0.8% (6/726) ragged-C
Nontryptic peptides (% of all peptides): 0.0% (0/717)

#### Oxidation

Oxidized methionine (M[+16]): 18.5% (55/298) of peptides containing M[+0] or M[+16]
Side chain loss from oxidized methionine (M[-48]): 0.0% (0/241) of peptides containing M[+0] or M[-48]
Doubly oxidized methionine (M[+32]): 0.0% (0/199) of peptides containing one M[+32] or M[+0]
Oxidized histidine and tryptophan (H[+16], W[+16]): 0.5% (1/190) (1 H, 0 W) of peptides containing H or W
Doubly oxidized tryptophan (W[+32]): 0.0% (0/52) of peptides containing W but not M
Triply oxidized cysteine (C[+48]): 0.0% (0/46) of peptides containing C

#### Chemical modifications

Deamidated asparagine or glutamine: 9.4% (75/797) (65 N, 10 Q) of peptides containing N and/or Q
Amidated aspartic or glutamic acid: 0.2% (2/839) of peptides containing D and/or E
Pyro-glu N-terminus (Q[-17], E[-18], C[+57][-17]): 7.1% (6/85) (6 -17, 0 -18) of peptides with N-terminal Q, E, or camC
Sodiation: 0.0% (0/1019) of all peptides (0 on E and D)
Carbamylation (N-terminus[+43], R[+43], K[+43]): 0.1% (1/1022) (0 N-term) of peptides
Carbamylated methionine (M[+43]): 0.0% (0/298) of peptides containing M
Formaldehyde (N-terminus[+12], W[+12]): 0.0% (0/1019) of peptides
Acetaldehyde (N-terminus[+26], H[+26], K[+26]): 0.0% (0/1019) (0 N-term) of peptides
N-terminal methylation/dimethylation (N-terminus[+14/+28]): 0.8% (8/1019) (8 +14, 0 +28) of peptides
Peptide (not protein) N-terminal acetylation (N-terminus[+42]): 0.0% (0/1019) of peptides
Cysteine propionamide (C[+71]): 2.2% (1/46) of peptides containing C
Methyl ester (E[+14]): 0.3% (2/588) of peptides containing E
Formylation (S[+28], T[+28]): 0.2% (1/611) of peptides containing S or T

#### Posttranslational modifications

Hydroxyproline: 1.6% (6/370) of peptides containing P
Phosphorylation: 0.0% (0/727) (0 S, 0 T, 0 Y) of peptides containing S, T, or Y
Beta-elimination (S[-18], T[-18]): 0.0% (0/608) peptides containing S or T
Dimethylation (K[+28], R[+28]): 1.1% (11/983) (11 K, 0 R) of peptides containing K or R
Methylation (K[+14], H[+14], N[+14], R[+14]): 0.6% (6/990) (0 K) of peptides containing K, H, N, or R
Acetylation (or guanidination or trimethylation) (K[+42]): 0.7% (4/591) of peptides containing K
Protein N-terminal acetylation: 50.0% (3/6) of N-terminal peptides

#### Computational information

Spectrum file: D:\Luci\Moggridge et al data\ch_06Fe...s_1.raw_Preview_181130_173846\objs\ch_06Fe...e-yes_1.mgf
Protein database: D:\Luci\PD Databases\Schulz_CP_Homo_sapiens_proteomeUP000005640reveiwed_Download_20180420_proteins_20303.fasta
Presets: Cysteine 57, Lysine 0, Arginine 0, N-terminus 0, C-terminus 0
CID fragmentation
Digestion after RK
First pass is fully tryptic digestion
No wildcard search
Read in 21688 spectrum/charge combinations: z=+1 0, z=+2 15186, z=+3 5783, z=+4 719
Precursor mass from 756 to 3000 Da
Read 42313 proteins
Scored 344315466 candidate/spectrum/charge combinations

Preview version: v2.13.17
Output folder: D:\Luci\Moggridge et al data\ch_06Fe...s_1.raw_Preview_181130_173846

Click to open **parameters** used for this run.

Note: The Details page reports the rates of modification on eligible peptides, whereas the Summary page reports potential gains in the total number of identifications. Denominators in the percentages may also vary from search to search due to "second-order" effects such as multiply modified peptides and corrections for hits to decoys.

| Summary | Detail |
|---|---|

www.proteinmetrics.com

PROTEIN METRICS
