## Supplementary Table S4 for "S-Trap eliminates cell culture media polymeric surfactants for effective proteomic analysis of mammalian cell bioreactor supernatants"

Summary of experimental details for each of the comparisons presented in this work

| **Figure** | **Biological samples compared** | **Replicates** | **Method and lysis buffer** |
|---|---|---|---|
| 1 | Organism: CHO cells<br>Supernatant from bioreactor at day 13 of fed batch, media was CD CHO + EfficientFeed B + Vitamin K + 1g/L Pluronic F68 | 1 replicate | **FASP** (6 M GdCl, 50 mM Tris pH 8, 10mM DTT, Amicon Ultra 0.5mL Centrifugal Filters 10 kDa cut-off) |
| 2 | Organism: CHO cells<br>Supernatant from bioreactor at day 12 of fed batch, media was CD CHO + EfficientFeed B + Vitamin K + 1g/L Pluronic F68 | 2 technical replicates | **FASP** (6 M GdCl, 50 mM Tris pH 8, 10mM DTT, Amicon Ultra 0.5mL Centrifugal Filters 30 kDa cut-off)<br>**SP3** (1% SDS, 50 mM Hepes pH 7.4, 10 mM DTT, 1X Roche PICs)<br>**Precipitation** (6 M GdCl, 50 mM Tris pH 8, 10mM DTT)<br>**S-Trap** (5% SDS, 50 mM Tris pH 7.55, 10 mM DTT) |
| 3 | Organism: *Saccharomyces cerevisiae*<br>Whole cell extract | 2 technical replicates | **SP3** (1% SDS, 50 mM Hepes pH 7.4, 10 mM DTT, 1X Roche PICs)<br>**Precipitation** (1% SDS, 50 mM Hepes pH 7.4, 10 mM DTT, 1X Roche PICs)<br>**S-Trap** (5% SDS, 50 mM Tris pH 7.55, 10 mM DTT) |
